# The role of the right language network and the multiple-demand network in verbal semantics: Insights from an Activation Likelihood Estimation Meta-analysis of 561 Functional Neuroimaging Studies

**DOI:** 10.1101/2025.04.05.647349

**Authors:** Eszter Demirkan, Francesca M. Branzi

## Abstract

Language processing has been traditionally associated with a network of fronto-parietal and temporal regions in the left hemisphere. Nevertheless, the ‘right language network’ (frontal, temporal, and parietal regions homologous to the left language network) and the ‘Multiple-Demand Network’ (MDN) are often involved in verbal semantic processing as well, however their role remain poorly understood. This is in part due to the inconsistent engagement of these latter two networks across linguistic tasks. To explore the factors driving networks recruitment of right language network and MDN during verbal semantic processing, we conducted a large-scale Activation Likelihood Estimation meta-analysis of neuroimaging studies. We examined whether the right language network is influenced by verbal stimulus type (sentences/narratives versus single words/word pairs) and whether this may be due to differences in semantic control demands and/or the presence of social content in the stimuli. Additionally, we investigated whether MDN recruitment depends on external task demands rather than semantic control demands. Our main findings revealed greater engagement of the right language network during semantic processing of sentence/narrative stimuli, with distinct regions reflecting different functions: increased semantic control demands recruit the right inferior frontal gyrus. Instead, social content processing during a semantic task engages the right Anterior Temporal Lobe, as well as the right posterior middle temporal gyrus. Finally, semantic processing engages the MDN, but only when external task (rather than semantic) demands increase.

## Introduction

Language processing has been traditionally associated with a network of fronto-parietal and temporal regions in the left hemisphere (*henceforth* ‘left language network’) (Fedorenko et al., 2024; Friederici & Gierhan, 2013; Price, 2012). Yet, there is increasing evidence that right homologue brain regions (*henceforth* ‘right language network’) support language and verbal semantic processing as well (Lindell, 2006). Not only healthy individuals recruit right frontal, parietal and temporal brain regions during verbal semantic processing (e.g., (Branzi, Humphreys, et al., 2020; Hodgson et al., 2021); the ‘right language network’ plays a crucial role in language recovery after left-hemisphere brain damage (Crinion & Price, 2005; Nardo et al., 2017). These findings align with evidence that short-term disruption of left-hemisphere processing, induced by neurostimulation, prompts the recruitment of the right language network to support efficient semantic processing (Binney & Lambon Ralph, 2015; Hartwigsen et al., 2013; J. Jung & Lambon Ralph, 2016). All this evidence suggests that the right hemisphere possesses significant language processing capabilities.

Nevertheless, the function of the ‘right language network’ in language processing is not well understood (*see* Fedorenko et al., 2024). One main reason is that the right language network seems not as consistently involved in language processing as the left language network. For instance, the right hemisphere homologues of left hemisphere language areas seem to be rarely recruited in tasks that require single or word-pair semantic processing (Badre et al., 2005; Graessner et al., 2021; Whitney et al., 2009). Conversely, other types of tasks, such as sentence and/or narrative processing tasks, which require integration of multiple meanings, tend to recruit more extensively the right language network (Branzi, Humphreys, et al., 2020; Silbert et al., 2014; Xu et al., 2005).

Although the literature reviewed above suggests that the right language network supports meaning processing, particularly during sentence/narrative processing tasks, the evidence remains inconclusive. The studies reviewed above differ not only in the type of semantic stimuli, but also in terms of sample sizes, task demands, and analyses employed. These idiosyncrasies make it difficult to draw definitive conclusions.

Therefore, as an initial step to address this question, i.e., whether the right language network is particularly involved in tasks that require processing meaning across multiple words and/or sentences, we adopted a meta-analytic approach instead of conducting a single study. A meta-analytic method is particularly suitable for addressing this question as it allows us to establish which results are replicable and generalisable across multiple studies, while also overcoming the limitations of individual studies, which are often underpowered (Button et al., 2013) and susceptible to the effects of unique design and analytic choices.

The right language network may be particularly involved in sentence/narrative processing tasks due to semantic control demands (Stefaniak et al., 2020). We reasoned that the need to buffer information over time and compute complex semantic meaning across short but also longer time scales may increase control demands, and therefore determine the recruitment of the right language network (see Branzi et al., 2020; Branzi & Lambon Ralph, 2023). This hypothesis is in line with evidence showing that increasing semantic control demands results in increased neural activation in the right anterior temporal lobe (ATL), right posterior middle temporal gyrus (pMTG), right inferior frontal (IFG) and right angular gyrus (AG) (Branzi, Humphreys, et al., 2020; Hodgson et al., 2022; Humphreys & Lambon Ralph, 2017; J. Y. Jung et al., 2021; Quillen et al., 2021; Rice et al., 2018). In healthy young individuals, the right language network activity is generally not observed, unless semantic control demands are increased (J. Y. Jung et al., 2021; Rice et al., 2018). However, the evidence is mixed, with studies reporting different or contrasting results (Quillen et al., 2021; Rice et al., 2018). Thus, whether the right language network supports verbal semantic processing particularly when semantic control demands increase (e.g., Rice et al., 2018; Stefaniak et al., 2020) remains an open question.

Another non-mutually exclusive possibility is that the right language network, or at least parts of it, does not support core semantic functions (e.g., meaning processing and retrieval) but rather, social cognition processes, which would be assimilated during the language tasks only when they require those specific functions. Sentences and narratives often involve human characters, which may engage social cognition. Thus, language comprehension may require inference-making processes about people’s mental states and/or the processing of other types of ‘Social Cues’ (e.g., prosody). Accordingly, various studies have shown that social inferences about other people’s mental states (or ‘Theory of Mind-ToM’ processing) as compared to non-social inferences (e.g., mechanical inference) engage brain areas within the right language network, such as the right vAG/temporoparietal junction (TPJ) and the right ATL (Saxe et al., 2004; Saxe & Wexler, 2005). Besides social inference-making processes, also the perception of Social Cues engages parts of the right language network, including the right ventral AG (vAG/TPJ) and the right pMTG (Hodgson et al., 2022; Saxe et al., 2004; Turker et al., 2023).

A recent large Activation Likelihood Estimation (ALE) meta-analysis of functional neuroimaging studies has revealed some neural overlap between the ALE maps for ToM and semantic cognition in the above-mentioned areas (Balgova et al., 2024). However, it is unclear if this is because semantic processes are integral to performing ToM tasks (Balgova et al., 2024) or rather because some stimuli (i.e., sentences/narratives) contained in the semantic cognition meta-analysis indeed engaged some ToM and social cognition processing.

To date whether the right hemisphere homologues of left hemisphere language regions can perform similar — albeit weaker — language computations as their dominant left homologues (Stefaniak et al., 2020) or whether, instead, they support different cognitive functions, such as those related to social cognition is unknown. Understanding the contribution of the right hemisphere to verbal semantic processing is important not only to delineate the functional organisation of the language network in healthy individuals but also to understand language recovery following brain damage in neurological patients. Thus, the primary goal of this large-scale ALE meta-analysis of functional neuroimaging studies is to clarify whether the recruitment of the right language network during verbal semantic tasks is modulated by (1) the type of stimuli (single-words/word-pairs *versus* sentences/narratives) and if this is due to semantic control demands (high *versus* low control demands). Furthermore, we aim to establish whether (2) the right language network engaged during verbal semantic cognition overlaps with the neural network engaged during social and ToM cognition, particularly when stimuli are likely to engage these processes (e.g., during semantic tasks with sentences or narratives).

An additional goal of this study is (3) to address the role of the Multiple-Demand Network or MDN (Duncan, 2010; Fedorenko et al., 2013) in language processing and the extent to which its recruitment depends on task demands. Recent studies have shown that the MDN, a set of domain-general brain regions typically activated during a diverse range of executively demanding language and non-language cognitive tasks (Duncan, 2010; Fedorenko et al., 2013), does not support core language operations (Blank & Fedorenko, 2017; Branzi & Lambon Ralph, 2023). Nevertheless, these conclusions have been mainly drawn from studies that have employed low demand tasks, i.e., passive tasks such as reading and listening sentences. Thus, it is necessary to determine whether these results generalise also to tasks that require a linguistic decision to be made (e.g., whether a stimulus is a word or pseudoword, concrete or abstract, etc).

Our first hypothesis is that, independently from the perceptual modality (visual or auditory), the right language network supports the same lexical-semantic functions as the left hemisphere. Still, it may be recruited particularly when semantic control demands increase (Stefaniak et al., 2020). If true, upregulation of activity within the right language network should be observed only when semantic control demands increase, independently from the stimulus type (single-words/word-pairs or sentences/narratives). However, as hinted above, we expect semantic tasks with sentences/narratives to recruit the right language network more extensively as compared to single-words/word-pairs. This is because meaning processing is computed over time and across multiple words.

The brain areas that we expect to be modulated by semantic control demands include the right pMTG, the right IFG and the right dorsal AG/intraparietal sulcus (dAG/IPS) (Lambon Ralph et al., 2017). Instead, we do not expect increased semantic demands to extensively modulate activity within the MDN (Duncan, 2010; Fedorenko et al., 2013). To test these hypotheses, we examined whether semantic control demands modulated the recruitment of the right language network in both sentences/narratives and single word type/word pairs tasks, separately. To verify if the brain areas involved in semantic control overlapped with the MDN we examined the spatial overlap between the clusters identified in the above analyses with the MDN mask derived from (Fedorenko et al., 2013).

Our second hypothesis is that some brain regions within the right language network support social cognition processing during language tasks. Particularly, the right ATL and the right vAG/TPJ are often implicated in social cognition processing (Diveica et al., 2021; Olson et al., 2007; Saxe et al., 2004; Zahn et al., 2007). Accordingly, neural activity in these regions should be observed only when the task may require social cognition processing, that is, when verbal semantic stimuli involve the processing of information about ‘people and/or social words that convey information about relationships with people and can inform our understanding of their actions’ (*see* Pexman et al., 2023). Therefore, to establish whether the brain regions activated during language tasks reflect social cognition processing, we examined the neural overlap between a contrast reflecting semantic cognition measured during sentence/narrative tasks with social elements (i.e., containing information about people), but excluding sentence/narrative tasks with no social elements (i.e., containing descriptions of nature), and a contrast reflecting ToM processing measured during non-verbal tasks. We reasoned that any observed neural overlap between these two contrasts would not reflect similarities due to the type of stimuli (e.g., verbal input), but rather social cognition processing. Furthermore, we also examined an overlap between a contrast reflecting semantic cognition measured during sentence/narrative tasks with social elements and a contrast reflecting the processing of Social Cues.

Critically, for both formal conjunction analyses we expected neural overlap in the right vAG/TPJ and the right ATL, i.e., brain regions previously associated with social cognition (Diveica et al., 2021; Hodgson et al., 2022; Saxe & Wexler, 2005), during sentence and/or narrative processing with social content, but not during verbal semantic tasks including single-words/word-pairs as stimuli without social-related content.

Finally, other brain regions might be recruited during language tasks. These brain regions may overlap with the MDN (Duncan, 2010). Even though we included semantic contrasts against active baselines (see methods), not all the active baseline/control tasks are necessarily designed to control for task demands. Thus, it is still possible that the recruitment of these brain regions is due to increased demands imposed by the task at hand, rather than control imposed at the lexical and semantic levels. For instance, tasks that involve deciding between different response options may require inhibitory control processes that are not required during more passive tasks such as reading or listening tasks with no explicit instruction to make a semantic decision. If true, then activity in brain areas overlapping with the MDN may be observed in tasks that involve increased executive control demands (e.g., tasks requiring a semantic decision), irrespective of the type of stimuli (sentences/ narratives or single words/word-pairs). Accordingly, this study tackled these targeted questions through the largest ALE meta-analysis, to date, of functional neuroimaging semantic studies in healthy individuals (n= 561).

## 2. Materials and methods

There were several steps to address the questions of the present study. First, two ALE meta-analyses were conducted to identify brain regions associated with general verbal semantic cognition: one for tasks involving multi-item processing (e.g., sentences and narratives) and another one for tasks involving single-word or word-pair stimuli. Then, we computed two ALE meta-analyses to examine which brain regions were sensitive to increased semantic control demands, and whether these regions differed depending on the type of stimuli, i.e., sentences/narratives and single-word/word-pair stimuli. Then, to verify if the brain areas involved in semantic control overlapped with the MDN, we examined the spatial overlap between the clusters identified in the above analyses with the MDN (Fedorenko et al., 2013).

In a second step, two ALE meta-analyses were conducted to reveal the brain regions reflecting social cognition processing, including non-verbal ToM and the processing of verbal and non-verbal Social Cues (e.g., prosody and biological motion). Then, through formal conjunction analysis, we examined whether the brain regions sensitive to multi-item semantic processing (sentences/narratives) (see above) were also sensitive to social cognition processing measured in non-verbal ToM and Social Cues processing.

In a third step, we examined whether the neural network supporting verbal semantic cognition in sentences/narratives and single-words/word-pairs was modulated by external task demands. To do so, we conducted separate ALE meta-analyses and identified the neural networks underlying verbal semantic cognition across tasks with low and high control demands. Finally, we examined the extent to which these regions overlapped with the MDN (Duncan, 2010; Fedorenko et al., 2013).

### 2.1. Studies selection

All meta-analyses were based on experimental studies published in peer-reviewed journals in English, describing task-based activation coordinates reported in Montreal Neurological Institute (MNI) or Tailarach coordinate systems, as a result of univariate analyses from fMRI or PET studies, and studying only young (< 40 years old) healthy adults. Coordinates from multiple contrasts within the same studies coming from the same set of participants were combined into one study, except in cases where the contrasts differed in a crucial aspect examined in the study (type of stimuli, modality, etc) (Müller, et al., 2018).

For all analyses, coordinates were obtained from previous meta-analyses (see details below). Additionally, the Web of Science (www.webofscience.com) was used to identify new studies for each meta-analysis, employing the same search terms as those used in the original meta-analyses (see details below). In line with previous meta-analyses (Diveica et al., 2021; Hodgson et al., 2022; Jackson, 2021; Turker et al., 2023), we only included coordinates derived from studies using whole-brain analyses and discarded coordinates derived from studies focussing on region-of-interest analyses or small-volume correction analyses since these types of analyses violate a key assumption of coordinate-based meta-analyses (Eickhoff et al., 2012; Müller, Cieslik, et al., 2018).

While the coordinates for each ALE meta-analysis and the search for new studies were based on results and selection criteria of existing meta-analysis studies, we adjusted the list of studies (and coordinates) to make the different meta-analyses optimally comparable to each other (**see Tables 1-4 in Supplementary Materials**). In detail, for the general verbal semantic cognition, verbal semantic control, and non-verbal social cognition ALE meta-analyses, we only included studies in which the experimental condition was contrasted against an active baseline. Instead, we did not include coordinates taken from studies where the experimental condition of interest was contrasted against the rest/passive baseline which required lower cognitive demands than the experimental condition. This approach is beneficial not only for controlling methodological differences across various types of stimuli (sentences/narratives *versus* single-words/word-pairs) or cognitive domains (semantic *versus* social), but also for ensuring that the observed neural networks reflect the processes of interest rather than task-related differences between experimental conditions and baseline. Many brain regions associated with semantic cognition and social cognition overlap with the Default Mode Network (DMN) (Andrews-Hanna et al., 2010; Buckner et al., 2008; Raichle et al., 2001). The DMN is a resting-state network that typically shows reduced activity during demanding tasks but also increased activity during tasks requiring internally oriented cognition (Konu et al., 2020; Smallwood et al., 2021; Vatansever et al., 2017). Therefore, a baseline that does not match the experimental condition in terms of difficulty and the degree of internally *versus* externally oriented processing can introduce confounding for the interpretation of meta-analytical results. The total number and the type of studies for each meta-analysis are listed in **Table 1.**

**Table 1.**
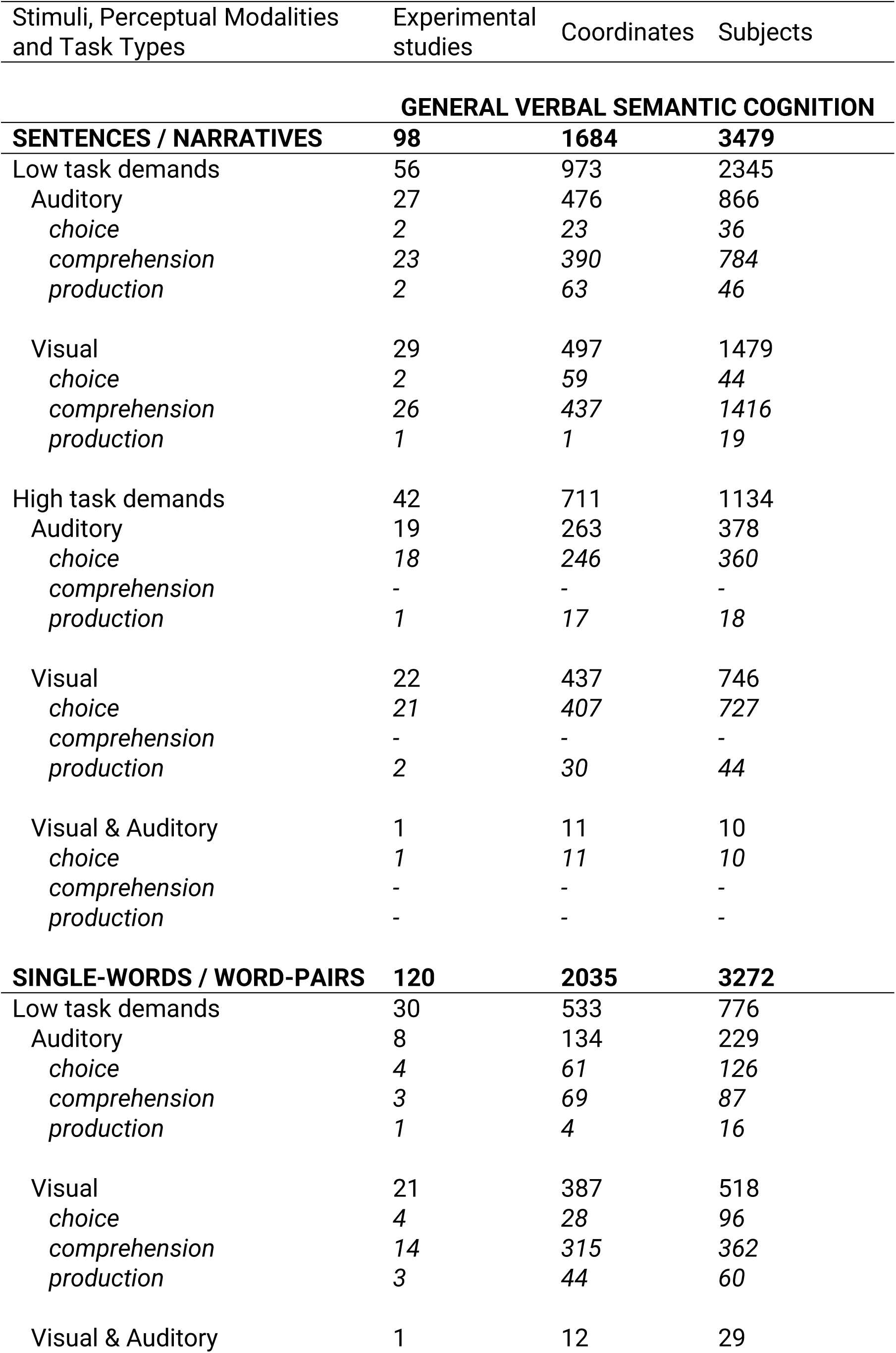
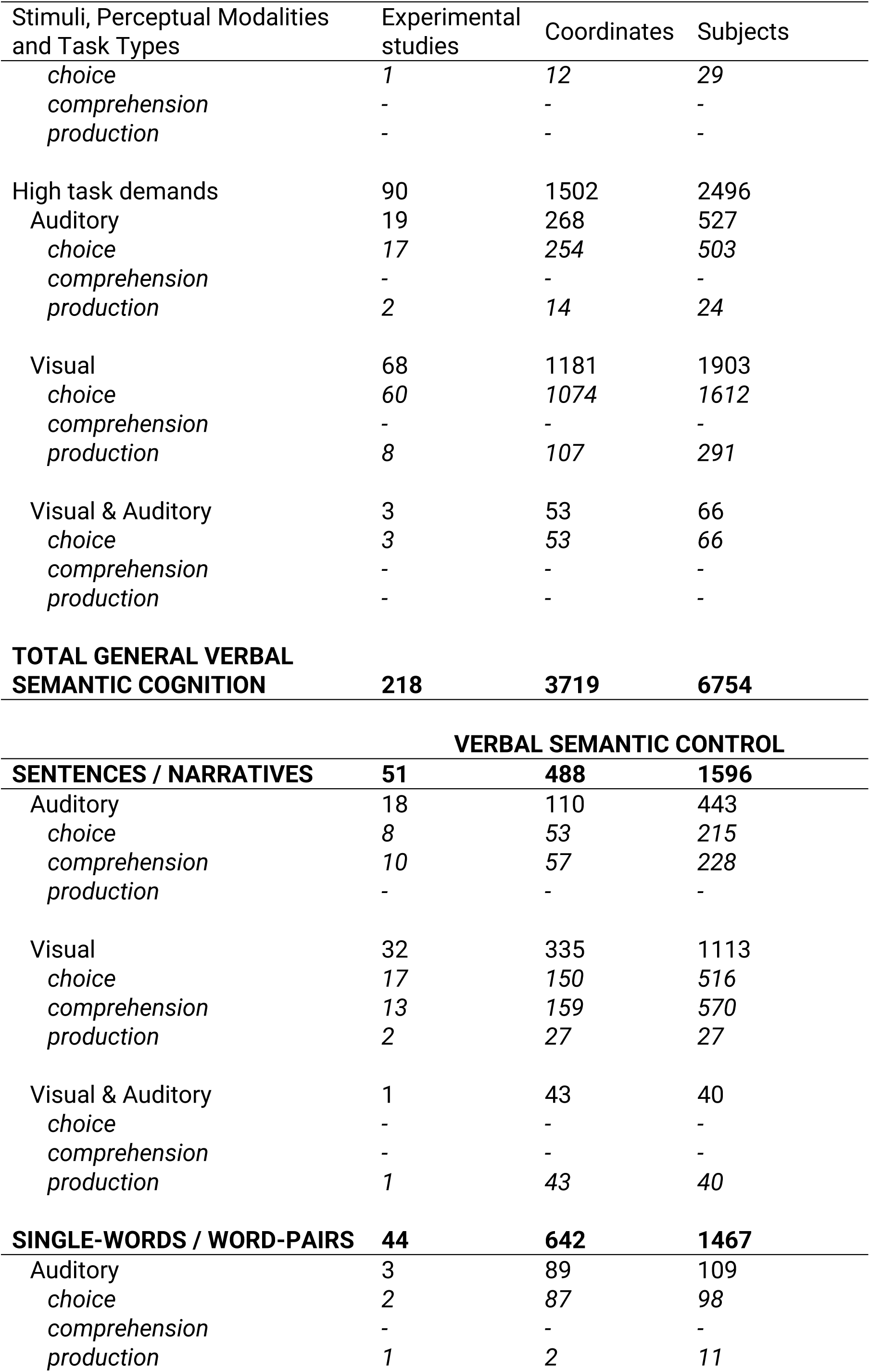
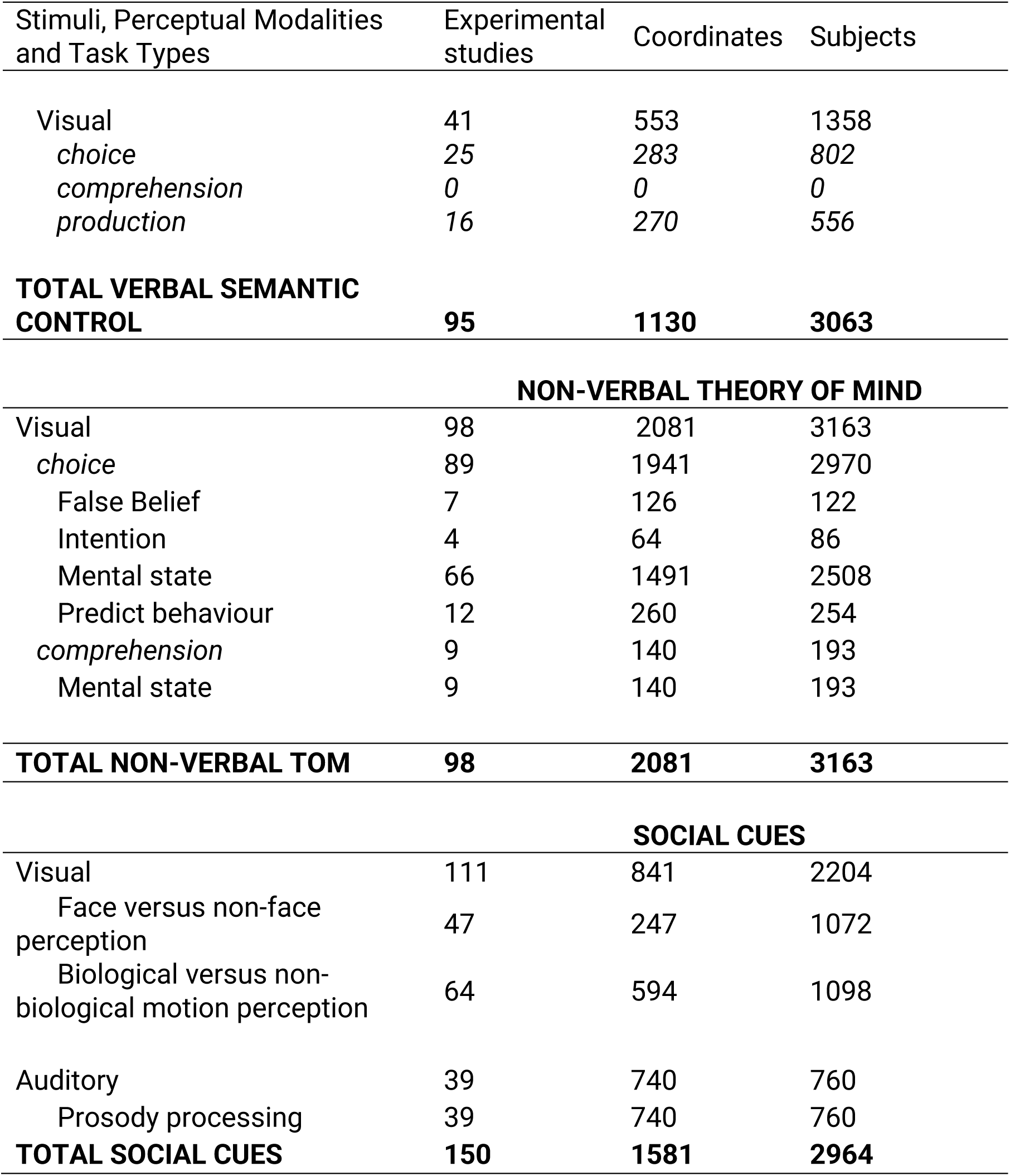
Number of studies, coordinates, and subjects included in each ALE meta-analysis.

| Stimuli, Perceptual Modalities and Task Types | Experimental studies | Coordinates | Subjects |
| --- | --- | --- | --- |
| **GENERAL VERBAL SEMANTIC COGNITION** |  |  |  |
| **SENTENCES / NARRATIVES** | **98** | **1684** | **3479** |
| Low task demands | 56 | 973 | 2345 |
| Auditory | 27 | 476 | 866 |
| *choice* | 2 | 23 | 36 |
| *comprehension* | 23 | 390 | 784 |
| *production* | 2 | 63 | 46 |
| Visual | 29 | 497 | 1479 |
| *choice* | 2 | 59 | 44 |
| *comprehension* | 26 | 437 | 1416 |
| *production* | 1 | 1 | 19 |
| High task demands | 42 | 711 | 1134 |
| Auditory | 19 | 263 | 378 |
| *choice* | 18 | 246 | 360 |
| *comprehension* | - | - | - |
| *production* | 1 | 17 | 18 |
| Visual | 22 | 437 | 746 |
| *choice* | 21 | 407 | 727 |
| *comprehension* | - | - | - |
| *production* | 2 | 30 | 44 |
| Visual & Auditory | 1 | 11 | 10 |
| *choice* | 1 | 11 | 10 |
| *comprehension* | - | - | - |
| *production* | - | - | - |
| **SINGLE-WORDS / WORD-PAIRS** | **120** | **2035** | **3272** |
| Low task demands | 30 | 533 | 776 |
| Auditory | 8 | 134 | 229 |
| *choice* | 4 | 61 | 126 |
| *comprehension* | 3 | 69 | 87 |
| *production* | 1 | 4 | 16 |
| Visual | 21 | 387 | 518 |
| *choice* | 4 | 28 | 96 |
| *comprehension* | 14 | 315 | 362 |
| *production* | 3 | 44 | 60 |
| Visual & Auditory | 1 | 12 | 29 |
| *choice* | 1 | 12 | 29 |
| *comprehension* | - | - | - |
| *production* | - | - | - |
| High task demands | 90 | 1502 | 2496 |
| Auditory | 19 | 268 | 527 |
| *choice* | 17 | 254 | 503 |
| *comprehension* | - | - | - |
| *production* | 2 | 14 | 24 |
| Visual | 68 | 1181 | 1903 |
| *choice* | 60 | 1074 | 1612 |
| *comprehension* | - | - | - |
| *production* | 8 | 107 | 291 |
| Visual & Auditory | 3 | 53 | 66 |
| *choice* | 3 | 53 | 66 |
| *comprehension* | - | - | - |
| *production* | - | - | - |
| **TOTAL GENERAL VERBAL SEMANTIC COGNITION** | **218** | **3719** | **6754** |
| **VERBAL SEMANTIC CONTROL** |  |  |  |
| **SENTENCES / NARRATIVES** | **51** | **488** | **1596** |
| Auditory | 18 | 110 | 443 |
| *choice* | 8 | 53 | 215 |
| *comprehension* | 10 | 57 | 228 |
| *production* | - | - | - |
| Visual | 32 | 335 | 1113 |
| *choice* | 17 | 150 | 516 |
| *comprehension* | 13 | 159 | 570 |
| *production* | 2 | 27 | 27 |
| Visual & Auditory | 1 | 43 | 40 |
| *choice* | - | - | - |
| *comprehension* | - | - | - |
| *production* | 1 | 43 | 40 |
| **SINGLE-WORDS / WORD-PAIRS** | **44** | **642** | **1467** |
| Auditory | 3 | 89 | 109 |
| *choice* | 2 | 87 | 98 |
| *comprehension* | - | - | - |
| *production* | 1 | 2 | 11 |
| Visual | 41 | 553 | 1358 |
| *choice* | 25 | 283 | 802 |
| *comprehension* | 0 | 0 | 0 |
| *production* | 16 | 270 | 556 |
| **TOTAL VERBAL SEMANTIC CONTROL** | **95** | **1130** | **3063** |
| **NON-VERBAL THEORY OF MIND** |  |  |  |
| Visual | 98 | 2081 | 3163 |
| *choice* | 89 | 1941 | 2970 |
| False Belief | 7 | 126 | 122 |
| Intention | 4 | 64 | 86 |
| Mental state | 66 | 1491 | 2508 |
| Predict behaviour | 12 | 260 | 254 |
| *comprehension* | 9 | 140 | 193 |
| Mental state | 9 | 140 | 193 |
| **TOTAL NON-VERBAL TOM** | **98** | **2081** | **3163** |
| **SOCIAL CUES** |  |  |  |
| Visual | 111 | 841 | 2204 |
| Face versus non-face perception | 47 | 247 | 1072 |
| Biological versus non-biological motion perception | 64 | 594 | 1098 |
| Auditory | 39 | 740 | 760 |
| Prosody processing | 39 | 740 | 760 |
| **TOTAL SOCIAL CUES** | **150** | **1581** | **2964** |

### 2.2. Verbal Semantic Cognition

The experiments assessing verbal semantic cognition have employed many different tasks and designs. These often vary substantially in terms of cognitive load, computations, and modalities involved. For instance, in some tasks, both the experimental task and control task might require participants to passively comprehend a word or sentence, while instead, other tasks might require participants to generate several words according to instructions. These differences may affect the recruitment of language areas as well as the MDN (Diachek et al., 2020; Hu et al., 2023) and result in an artificial divergence in the neural networks supporting semantic cognition during sentence/narrative processing *versus* single-word/word-pairs processing. Therefore, to ensure that studies included in each semantic ALE meta-analysis did not differ substantially in terms of cognitive load, computations, and modalities involved, i.e., aspects that may affect the results, we screened and classified semantic studies according to stimulus modality (visual, auditory, or audio-visual) and task type. Task types were the following: performing a choice (requiring a decision between two or more options usually accompanied by a button press), passive comprehension (reading or listening), or cue-based production (both external: semantic fluency tasks, or internal: think about an item belonging to a semantic category). We also categorised studies according to task demands, in line with (Stefaniak et al., 2021) (see **Table 1 and Section 2.2.3 for details**) to address the role of the MDN (Duncan, 2010; Fedorenko et al., 2013) in verbal semantic processing and particularly the extent to which its recruitment depends on external task demands.

#### 2.2.1. General Verbal Semantic Cognition in words versus sentences

These ALE meta-analyses were based on a selection of studies reported in (Jackson, 2021) measuring the neural basis of verbal semantic cognition through the comparison of neural activity during verbal semantic tasks against non-semantic (or less semantic) tasks (active baseline). We excluded tasks explicitly measuring ToM (e.g., False Belief tasks) and/or requiring participants to think about people, such as tasks in which participants had to rate adjectives by how much they described them or perform verbal tasks involving the recognition of famous people. Thus, none of the tasks included in the verbal semantic cognition ALE meta-analyses explicitly required social cognition and/or ToM processing, e.g., the semantic tasks were not centred around giving a response to a character’s thoughts, perspectives, emotions, or mental states (156 studies met the criteria, yielding 2367 coordinates).

Examples of contrasts included in the verbal semantic ALE meta-analyses refer to comparisons between processing words *versus* pseudowords, generating words that matched the probe based on semantic association *versus* rhyme (phonology), and listening to meaningful sentences *versus* scrambled/non-meaningful sentences. We also updated the list of studies based on search words used by (Jackson, 2021), i.e., using the search terms [‘semantic’, ‘comprehension’, or ‘conceptual knowledge’], combined with [‘PET or fMRI’], excluding [‘patient’, ‘priming’, ‘disorder’, ‘dementia’, ‘ageing’, ‘bilingual’, ‘meta-analysis’ or ‘multivariate’], with a publication date range between 19/06/2019 – 05/01/2025. Therefore, 58 additional studies with 1345 coordinates were added to the studies sourced from (Jackson, 2021).

To identify the semantic neural network during sentence *versus* word processing, we separated these experiments assessing semantics according to whether the semantic task included sentences/narratives or single-words/word-pairs. To determine whether meaning processing activates neural regions similar to those involved in non-verbal ToM and Social Cue processing, specifically when stimuli contain social elements, we examined which stimuli were imbued with social content and which were not. Stimuli with a social element were defined as words or sentences describing people’s relationships, people’s mental states, and words or sentences that could inform our understanding of people’s actions and intentions (e.g. Pexman et al., 2023). For studies where the stimuli were not openly available, we emailed the authors and only included studies where we could ensure the stimuli fit the criteria. Given the nature of our stimuli categories, all tasks involving a social element were tasks with sentences/narratives as stimuli. Since only a few studies containing sentences did not include social aspects (e.g., describing a natural phenomenon), we eliminated them from our analyses. In total, we analysed 98 studies reporting 1684 coordinates involving tasks with a social element, and 120 tasks reporting 2035 coordinates with no social element. We also determined the type of task and modality for each task. The full list of studies is reported in **Supplementary Table 1**.

Finally, to ensure that any observed differences between semantic processing of sentences/narratives and single-words/word-pairs were not influenced by the type of task (choice, comprehension, production), we conducted the same analyses as above (verbal semantic cognition in sentences/narratives and in single-words/word-pairs) on a subset of studies, focusing only on tasks that required participants to make a choice. This control analysis accounted for differences in task types (and the likely cognitive demands) that varied substantially between sentence/narrative tasks and single-word/word-pair tasks. Furthermore, to minimise potential confounding effects related to the stimulus modality, we created this subset so that within both stimuli types (sentences/narratives and single-words/word-pairs), there were almost equal numbers of studies (and coordinates) involving auditory *versus* visual modalities. In detail, the sentences/narratives subset of choice tasks comprised 20 auditory studies with 268 coordinates and 19 visual studies with 253 coordinates. Instead, the single-words/word-pairs subset of choice tasks comprised 21 auditory studies with 325 coordinates, and 20 visual studies with 336 coordinates.

#### 2.2.2. Verbal Semantic Control (hard versus easy semantic task/condition) in sentences versus words

These ALE meta-analyses included studies that manipulated the amount of verbal semantic control required during the task and directly contrasted a harder verbal semantic condition against an easier verbal semantic condition. Coordinates were partly derived from (Jackson, 2021), including only studies with verbal stimuli (80 studies, 874 coordinates). We then searched the literature for new studies with a publication range from 19/06/2019 to 01/05/2025. The inclusion and exclusion criteria were based on (Jackson, 2021), as were the search words: [‘semantic’, ‘comprehension’, or ‘conceptual knowledge’], combined with [‘PET’ or ‘fMRI’], combined with [‘selection’, ‘retrieval’, ‘inhibition’, ‘control’, ‘elaboration’, ‘fluency’, ‘ambiguity’, ‘metaphor’ or ‘idiom’], excluding [‘patient’, ‘priming’, ‘disorder’, ‘dementia’, ‘ageing’, ‘bilingual’, ‘meta-analysis’ or ‘multivariate’]. In short, we included studies that manipulated the difficulty of semantic tasks and, therefore, the cognitive load on semantic control by either (1) focusing on a meaning that is less frequently used, e.g., requiring a weaker association or subordinate homonyms; (2) increasing the number of competitors or distractors when there is an answer required, or decreasing the semantic distance between target and distractors; (3) requiring processing a semantic violation or subordinate homonyms; (4) increasing the unpredictability or surprisal factor of semantic elements by not giving enough context; or (5) requiring flexible switching between different meanings or instructions (Jackson, 2021). This search yielded 17 more studies, reporting 265 coordinates. To determine whether some brain areas could be differentially modulated by increased semantic control demands depending on the type of stimuli, we split this dataset similarly as above, according to whether the stimuli corresponded to sentences/narratives or single-words/word-pairs **(Supplementary Table 2)**.

#### 2.2.3. General Verbal Semantic Cognition and task demands

*(high versus low demands)* As mentioned above, within the general verbal semantic cognition ALE meta-analysis, we separated studies according to task difficulty. In short, all types of tasks (choice, comprehension, and production) were categorised as ‘high task demands’ if they required the participant to perform a semantic decision or production of *>* 1 word. This ranged from participants differentiating between words and pseudowords, making semantic relatedness judgements, and producing words or thinking about them according to criteria based on meaning, or grammar. A task was categorised ‘low task demands’ if there was no linguistic decision to perform or required a very simple identity match or a low-level perceptual decision: passive comprehension tasks fell into this category, as well as low-level/perceptual choice and matching tasks in which the experimental task was not semantic in nature (font size, or whether two items were identical or not), and production tasks with single item naming or repetition. In total, we examined 86 studies yielding 1506 coordinates that were categorised as ‘low task demands’, and 132 studies yielding 2213 coordinates that fell into the ‘high task demands’ category. The full list of studies is reported in **Supplementary Table 2.**

### 2.3. Social cognition

We examined verbal semantic tasks separately according to stimuli with *versus* without social elements (sentences/narratives *versus* single-words/word-pairs). However, the comparison between sentence/narrative tasks *versus* single-word/word-pairs tasks may have resulted in brain activations reflecting non-social factors, such as syntax processing, or an increased cognitive load on working memory and attention. Therefore, to establish which brain regions active during semantic processing of sentences/narratives reflected the processing of social elements (e.g., identifying a character within the narrative, and performing mentalisations based on the characters’ actions and intentions, etc.) we examined the similarities between brain areas active during narrative/sentence semantic tasks and different types of social cognition tasks including non-verbal mentalisation and the processing of Social Cues. To do so, we computed two separate ALE meta-analyses, one reflecting non-verbal ToM and the other one reflecting Social Cue processing. The list of all studies contributing to social cognition contrasts is reported in **Supplementary Tables 3 and 4.**

#### 2.3.1. Non-verbal ToM

The ALE meta-analysis for non-verbal ToM was partly sourced from (Diveica et al., 2021), discarding studies with verbal stimuli. We included all the categories defined in their ALE meta-analysis (ToM, Trait inference, Empathy, and Moral reasoning). The tasks required participants to perform inferences about a character’s intents, traits, or emotional state via non-verbal stimuli. For example, some tasks presented participants with a cartoon story and required them to select the final image based on the intentions of the cartoon characters. Other tasks involved observing facial expressions to determine the emotion being displayed. These two tasks were categorised as ‘intention’, and ‘mental state’, respectively (see **Table 1**). Additionally, some studies presented the false belief paradigm, in which the perspective or knowledge of the participant differs from the perspective of a character. The participant is then required to disassociate from their perspective, take the character’s perspective (such as the classic Sally-Anne task (Baron-Cohen et al., 1985), and make inferences about the character’s mental state. Finally, some experiments presented non-human entities, such as cartoon shapes, which were moving either randomly or as if with a human goal (helpful, protective, etc). In these tasks, participants were required to determine the type of behaviour (‘predict behaviour’) (see **Table 1**).

As with the verbal semantic cognition ALE meta-analyses, we discarded studies that used a passive baseline. Therefore, coordinates reported in the current study were sourced from experiments in which the tasks were contrasted with an active baseline that matched the general cognitive demands of the experimental condition/task, but required no mentalisation (e.g., making physical inferences, or perception of non-human images). This resulted in 76 studies with 1398 coordinates. We extended this dataset with new studies using the same search words and criteria as described in (Diveica et al., 2021). Thus, the Web of Science was searched with the key words [‘PET’ or ‘fMRI’], combined with [‘theory of mind’, ‘ToM’, ‘mentalising’, ‘mentalizing’, ‘social judgement’, ‘social evaluation’, ‘social attribution’, ‘trait inference’, ‘impression formation’, ‘empath∗’, ‘morality’, ‘moral’, ‘moral decision making’, ‘moral emotion’, ‘harm’, or ‘guilt’], with a publication date ranging from 01/03/2020 to 05/01/2025. This resulted in in 22 additional studies with 679 coordinates in addition to those sourced from (Diveica et al., 2021) (76 studies with 1398 coordinates), resulting in a total of 98 studies with 2081 coordinates. The full list of studies is reported in **Supplementary Table 3.**

#### 2.3.2. Social Cues

This ALE meta-analysis included other types of social cognition tasks that required processing stimuli with social valence but were not explicitly ToM tasks. Therefore, in this ALE meta-analysis, we included studies of biological motion, face perception, and changes in prosody. These tasks presented participants with cues they would observe in a social interaction (motion, tone of voice), without directly requiring them to perform ToM or a semantic task. We included studies of biological motion, face perception, and prosody tasks. The studies were partly sourced from (Hodgson et al., 2022), particularly the category labelled ‘Biological motion’. To avoid examining the effects of face/emotion processing and mentalising along with biological motion processing, we only included studies in which actors’ faces were not visible (40 studies, 151 coordinates). Since (Hodgson et al., 2022) only examined studies up to 2010, we searched the literature using the same criteria (see Grosbras et al., 2012), and we updated the studies ranging from 19/05/2010 to 05/01/2025. Search terms on the Web of Science and criteria included ‘fMRI’ or ‘PET’, as well as ‘biological motion’, ‘action observation’, and ‘human movement’. As explained before, we excluded studies where stimuli included faces that were visible rather than blurred. This yielded 20 studies with 383 coordinates.

We then assessed ‘face perception’ and included 54 studies (158 coordinates) from the ALE meta-analysis of (Hodgson et al., 2022). Studies examining the perception of emotions (n=13) were excluded from this ALE meta-analysis. In fact, processing of emotional faces and making inferences about them were classified as ToM. Thus, these studies were included in the ALE meta-analysis of non-verbal ToM instead (see above). We then updated the dataset following the same search words and criteria from (Hodgson et al., 2022; Müller, Höhner, et al., 2018): ‘fMRI’, ‘PET’, ‘face’, ‘facial’, ‘emotion’, ‘expression’, ‘neutral’, ‘localizer’, between 01/01/2016 to 05/01/2025. We included all studies that contrasted images or videos of faces with a non-face or scrambled counterpart. Instead, we discarded studies that contrasted assessing emotional *versus* neutral faces. We did so to avoid including activation coordinates during emotional processing, as tasks containing emotional processing would be already included in our ToM contrast. This new search yielded 6 new studies with 135 coordinates.

Finally, we included studies assessing auditory prosody processing. Our dataset was based on the prosody ALE meta-analysis recently published by (Turker et al., 2023). These were tasks including judging whether presented sentences had the same melody/intonation, whether two auditorily presented words differed in their pitch, or passive listening of auditory material in the form of meaningless sound envelopes. In total, 38 studies (677 coordinates) were selected from this ALE meta-analysis. We then performed a search with the terms ‘functional magnetic resonance imaging’/’positron emission tomography’ and ‘prosody’, from 01/04/2021 to 05/01/2025. This new search resulted in 2 additional studies with 63 coordinates. The full list of studies is reported in **Supplementary Table 4.**

### 2.4. Data Analysis

All coordinate-based ALE meta-analyses were performed via the revised ALE algorithm (Eickhoff et al., 2009, 2012; Turkeltaub et al., 2012), using Ginger ALE version 3.0.2 (http://brainmap.org/ale). All analyses were performed in the MNI152 space. Before running the analyses, all coordinates reported in Tailarach space were converted to MNI using the Lancaster transform (‘icbm2tal’) via GingerALE (Laird et al., 2010; Lancaster et al., 2007). GingerALE treats each activation foci as a spatial probability distribution, centred at the given coordinate. It then creates a map in which each voxel of the brain is given a value that signifies the probability of one of these distributions falling within the voxel, resulting in the ALE value (Turkeltaub et al., 2002). Permutation processes ensure differentiating between noise and true convergence of foci, and the process accounts for the spatial uncertainty caused by the between-subjects and inter-laboratory nature of meta-analytic approaches (Eickhoff et al., 2009). Importantly, following the recommendation by (Müller, Cieslik, et al., 2018), we only computed ALE meta-analyses in which the number of included experiments exceeded the recommended 20.

For each ALE meta-analysis, we obtained an ALE-map which was thresholded using a priori cluster-forming threshold of *p <* .001 (uncorrected) and then conservative cluster-level family-wise error (FWE) correction with a *p*-value of .001, with 10,000 permutations. As explained in (Eickhoff et al., 2012), this method creates a random set of foci with the same number of experiments and subjects as the provided dataset, repeated 10,000 times, and assesses for each found cluster how likely it is to obtain a cluster of the same size from these randomly relocated foci.

We then performed subtraction and conjunction analyses to reveal differences and similarities between the obtained ALE-maps. The conjunction images were generated using the voxel-wise minimum value (Nichols et al., 2005) of the included ALE maps to highlight shared activation. Instead, contrast images were generated by subtracting one ALE map from another ALE map. For both analyses, we used a priori cluster-forming threshold of *p <* .001 (uncorrected), with a minimum cluster volume of 200 mm3, which was estimated using a permutation approach with 10,000 repetitions (as in Diveica et al., 2021). This algorithm pools both datasets and randomly divides the experiments in line with the original numbers in each group. ALE maps are then assigned for these two newly formed sets of coordinates, and the difference of ALE values between the two sets is calculated for each voxel in the brain. With 10,000 permutations, a null distribution is created, against which the differences in ALE values between the original groups of observed data are compared against (Eickhoff et al., 2011). Since there is still no established method to correct ALE difference maps for multiple comparisons, we set a conservative significance threshold of *p* < .001, as conventional (Eickhoff et al., 2011). Thus, we reported only clusters that survived to the threshold of *p* < .001, and the cluster-size threshold of 200 mm^3^.

All coordinates are reported in the MNI space. For identifying clusters, the software package Statistical Parametric Mapping (SPM12) was used, with the software Matlab 2023b. Within SPM12, we utilised the Anatomy toolbox version 3 (Eickhoff et al., 2005, 2006, 2007). For each local maxima, we reported the most likely cytoarchitectonic area, via the Cytoarchitecture / Julich-Brain Atlas feature (Zilles et al., 2002; Zilles & Amunts, 2010), for increased accuracy. The cytoarchitectonic information for each one of the foci was assigned based on the Maximum Probability Map, along with the quantified probability for the most likely histological area found at this location. To identify the labels for coordinates reported in the Results, we used the Harvard-Oxford microanatomical atlas, also via the Anatomy Toolbox (SPM).

#### 2.4.1. General Verbal Semantic Cognition - words versus sentences

First, we performed individual ALE meta-analyses with single-words/word-pairs and sentences/narratives studies *versus* their respective non-semantic (or less semantic) baselines to identify brain regions involved in the semantic processing of these different stimulus types. To examine the brain regions that were significantly different between these two stimulus types, we performed a subtraction analysis. To provide a more comprehensive overview of the general verbal semantic network, we also conducted another ALE meta-analysis including both sentences/narratives and single-word/word-pairs, which is reported in the **Supplementary Figure 1**.

#### 2.4.2. Verbal Semantic Control - words versus sentences

To reveal the brain regions involved in verbal semantic control processes and how might they differ based on stimuli type, we performed individual ALE meta-analyses with semantic control tasks involving single-words/word-pairs on the one side and sentences/narratives on the other side. To provide a more comprehensive overview of the verbal semantic control network, we also conducted another ALE meta-analysis including both sentences/narratives and single-word/word-pairs, which is reported in the **Supplementary Figure 2.**

#### 2.4.3. Social Cognition and Neural Overlap between General Verbal Semantic Cognition

To determine whether the right language network activated during verbal semantic tasks involving sentences and narratives (as opposed to single words or word pairs) reflects the processing of social content, we first conducted two ALE meta-analyses: one to identify brain regions active during non-verbal ToM and another to pinpoint regions involved in processing verbal and non-verbal Social Cues, which would accompany social interactions, but do not require direct mentalisation. Then, we performed one conjunction analysis to identify the brain areas reliably implicated in both verbal semantic processing of sentences/narratives (containing social elements) as well as in non-verbal ToM processing. With this, we aimed to identify clusters active during social inferences, independent from modality, in the presence of instructions to perform mentalisation (non-verbal ToM tasks), and in the absence of such instructions (verbal tasks with sentences/narratives with a social element).

Furthermore, we explored also the overlap between verbal semantic cognition and the processing of non-verbal and verbal Social Cues. Thus, we conducted an additional conjunction analysis between semantic processing in sentence/narrative tasks and Social Cue processing measured in verbal and non-verbal tasks. Finally, to ensure the overlap observed with the sentence/narrative tasks was due to the social content of the stimuli rather than verbal semantic processing, likely engaged – at least to some extent – in both semantic and social cognition ALE meta-analyses, we performed the same conjunction analyses between social cognition tasks and verbal semantic tasks but focusing only on single-words/word-pairs (without social content).

#### 2.4.4 General Verbal Semantic Cognition and task demands (high versus low demands)

To isolate areas responsive to high *versus* low task demands during verbal semantic processing, we sorted the coordinates of the general verbal semantic cognition ALE meta-analysis according to task demands. Then we performed separate ALE meta-analyses for low and high demand tasks. To pinpoint whether the MDN is particularly activated because of task demands (e.g., tasks requiring a semantic decision) we examined the overlap between the individual ALE meta-analyses (high and low demand tasks) with the MDN mask identified by (Fedorenko et al., 2013).

##### Data Availability statement

All data analysed in this study are included in Supplementary Materials. The results of all the analyses (thresholded masks) are available online on the Open Science Framework project page (https://osf.io/nd8ye/?view_only=067a2cc490c2426781a0e2ad3b0c6073).

## 3. Results

The full list of activation clusters and local maxima foci are reported in **Supplementary Tables 5, 6, 7, and 8**.

### 3.1. General Verbal Semantic Cognition - words versus sentences

The first goal of our study was to establish whether the right language network was differently recruited depending on the type of stimuli (single-words/word-pairs versus sentences/narratives). Therefore, after completing an ALE meta-analysis with all tasks assessing verbal semantic cognition (semantic > non-semantic contrast) (see **Supplementary Figure 1A**), we examined if this neural network for verbal semantic cognition fractionated depending on the stimuli type **(Figure 1A and 1B)**. We determined this via two separate ALE meta-analyses, one with tasks involving semantic stimuli with sentences/narratives > non-semantic (or less semantic) task, and one with single-words/word-pairs > non-semantic (or less semantic) task.

**Figure 1.**
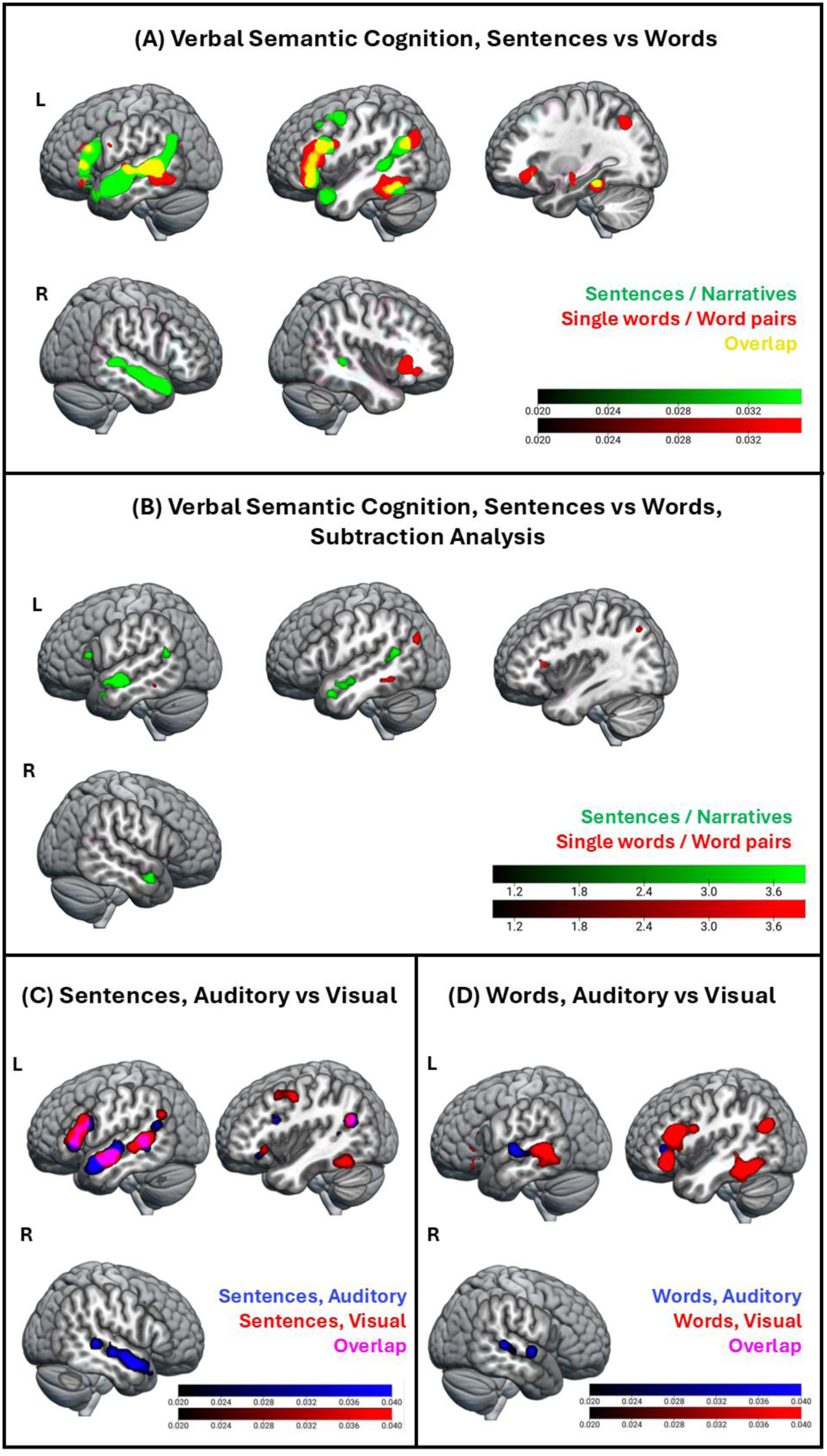
The Neural Network of General Verbal Semantic Cognition split by stimuli type and modality. **(A)** Brain regions reliably activated during verbal semantic cognition (semantic *versus* non-semantic or less semantic tasks) split by stimulus type (red: Single words/Word pairs; green: Sentences/Narratives) overlaid. Colour bar: ALE-values; cluster forming threshold: *p* < .001 (uncorrected); cluster extent correction: FWE *p* < .001. **(B)** Brain regions differentially engaged depending on stimulus type (red: Single words/Word pairs vs Sentences/Narratives; green: Sentences/Narratives vs Single words/Word pairs), calculated via t-test/subtraction analysis. Colour bar: Z-scores, cluster forming threshold*: p* <.001 (uncorrected), minimum cluster volume: 200 mm^3^. **(C)** Brain regions reliably activated during verbal semantic cognition (semantic *versus* non-semantic or less semantic tasks) for Sentences/Narratives split by the perceptual modality of the task, overlaid. Colour bar: ALE-values; cluster forming threshold: *p* < .001 (uncorrected); cluster extent correction: FWE *p* < .001. **(D)** Brain regions reliably activated during verbal semantic cognition (semantic *versus* non-semantic or less semantic tasks) for Single words/Word pairs split by the perceptual modality of the task, overlaid. Colour bar: ALE-values; cluster forming threshold: *p* < .001 (uncorrected); cluster extent correction: FWE *p* < .001.

On the one hand, results revealed that semantic processing of sentences/narratives recruited bilaterally the middle and superior ATL as well as the mid- and posterior portions of the temporal lobe, including pMTG. This activity in the right anterior and middle temporal lobes (as well as in the most anterior portions of the left ATL) was seen only in response to sentences/narratives processing.

Semantic processing of single-words/word-pairs stimuli activated the right Insula. In the left hemisphere, some activation in the left pMTG as well as a more inferior portion of the left posterior temporal gyrus (pITG) was observed during single-word/word-pairs tasks. The left dorsal and ventral IFG (Pars Triangularis and Pars Orbitalis, respectively) showed activation during semantic processing of both multi-item and single-word/word-pair stimuli. Finally, the left AG fractionated depending on the type of stimuli, with sentences/narratives recruiting vAG/TPJ, and single-words/word-pairs stimuli activating more dorsal and posterior portions of the left AG. Activation coordinates for all contrasts involving verbal semantic cognition are reported in **Supplementary Table 5**.

Some brain areas showed overlap between semantic processing of multi-item and single-word/word-pair tasks. Therefore, we performed a subtraction analysis to determine whether these overlapping brain regions were recruited to different extents depending on the stimuli (*see* **Figure 1B**). We found that the right ATL (together with the left ATL) was more strongly recruited during semantic tasks with sentences/narratives as compared to tasks with single-words/word-pairs. The left TPJ, as well as the left IFG Pars Triangularis also showed a significantly greater activation for semantic processing during sentences/narratives *versus* single-words/word-pairs. Semantic processing of single-words/word-pairs did not recruit regions of the right language network to a larger extent than processing of sentences/narratives. In the left hemisphere, the pITG and the Lateral Occipital Cortex seemed to be recruited more during single-words/word-pairs tasks.

An inspection of the semantic cognition dataset revealed an unequal distribution of perceptual modality between sentences/narratives and single-word/word-pair tasks (**see Table 1**). While sentence/narrative tasks included a similar number of auditory and visual tasks, resulting in comparable coordinate counts, single-word/word-pairs tasks were predominantly in the visual modality, with more than three times as many studies (and coordinates) from visual tasks compared to auditory ones. Therefore, to ensure the observed differences between sentences/narratives *versus* single-word/word-pairs were not due to the differences in perceptual modality, we examined the recruitment of brain regions based on perceptual modality. We performed four individual ALE meta-analyses, dividing the verbal semantic tasks based on both stimulus type and perceptual modality (**Figures 1C and 1D**).

The results revealed that semantic processing of visual stimuli did not activate large clusters in the right hemisphere, across both sentence and word task types. Semantic processing of auditory stimuli, instead, activated posterior areas of the right temporal lobe in both sentences/narratives and single-word/word-pair tasks. This result is not fully consistent with the results reported above (i.e., single-words/word-pairs *versus* non-semantic or less semantic tasks) which did not show any activation in the right hemisphere (**see Figure 1**). This discrepancy might be due to the imbalance between auditory and visual tasks in single-word/word-pairs experiments. As these tasks were predominantly presented in the visual modality, any activation linked to the auditory properties of the stimuli may have been too subtle to be detected in the ALE meta-analysis. However, the right ATL was consistently activated only for tasks with sentences/narratives as stimuli, indicating that neural activity in this brain area is primarily modulated by the presence of sentence/narrative stimuli.

The differences between sentences/narratives *versus* single-words/word-pairs reported above remained when we controlled for the type of task (**Supplementary Figure 1B**). In detail, we analysed a subset of the sentences/narratives and single-words/word-pairs datasets, focusing only on studies where participants were required to make a semantic choice (as opposed to passive comprehension, or answer generation). Even when the analyses were limited to choice tasks only, the results were consistent with those obtained when all tasks were included in the analysis – in detail, the right (and left ATL) only showed activation for sentences/narratives processing but not for single-words/word-pairs. This suggests that the type of task did not influence the differential activations observed between sentences/narratives *versus* single-words/word-pairs processing.

### 3.2. Semantic control - words versus sentences

To examine whether the right language network supports the same semantic functions as the left hemisphere but is recruited only when lexical demands increase, we conducted two ALE meta-analyses, one including only semantic tasks with single-word/word-pair stimuli and another one including only semantic tasks with sentence/narrative stimuli. The ALE meta-analyses included activation coordinates derived from hard > easy semantic task or condition contrasts (see **Figure 2A**). For completeness, in the **Supplementary Materials** we report the results of an ALE meta-analysis including both data sets (see **Supplementary Figure 2**). Activation coordinates for verbal semantic control are reported in **Supplementary Table 6**.

**Figure 2.**
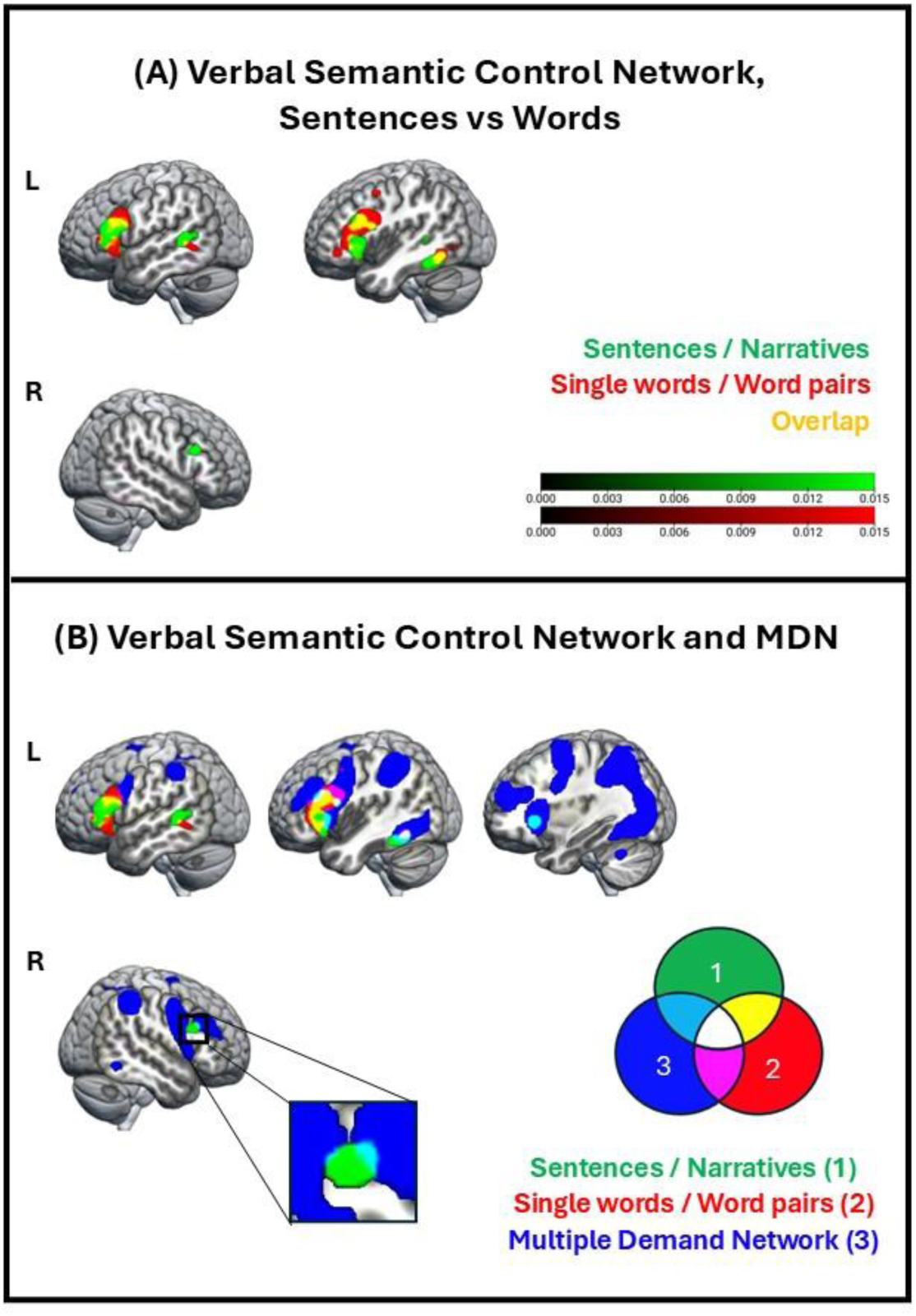
Verbal Semantic Control. **(A)** Brain regions reliably activated during all hard > easy semantic tasks or conditions, split by stimulus type (red: single-words/word-pairs, green: sentences/narratives) overlaid. Colourbar: ALE-values; cluster forming threshold: *p* < .001 (uncorrected); cluster extent correction: FWE *p* < .001. **(B)** The verbal semantic control network, split by stimulus type (red: single-words/word pairs, green: sentences/narratives), and the Multiple-Demand Network (MDN) identified by Fedorenko et al. (2013) (blue), overlaid. Colour codes are explained in the Venn diagram.

In line with our hypotheses, tasks or conditions with higher semantic control, as compared to those with lower semantic control, engaged the right IFG Pars Triangularis and Opercularis. This effect was only observed during tasks requiring higher semantic combinatorial demands, i.e., tasks using sentences/narratives as stimuli. Instead, increased semantic control demands in tasks using single words/word pairs did not activate the right hemisphere (**Figure 2A**). The right IFG is part of the right language network and was not observed in the general verbal semantic cognition results (see **Figure 1A / Supplementary Figure 1**).

For all types of stimuli, increased semantic control demands engaged left frontal regions, particularly the IFG Pars Triangularis and Opercularis. The posterior temporal lobe including left pITG and pMTG, seemed more engaged during hard *versus* easy semantic processing, across both stimulus types. However, left pSTG was modulated by semantic control demands only during sentences/narratives processing.

Finally, and in line with our hypotheses, the brain regions modulated by semantic control in both the right and left hemispheres showed a minimal overlap with the MDN. We found some overlap in the left Insula, the left and right IFG Pars Triangularis, and in the left IFG Pars Opercularis (**Figure 2B**).

### 3.3. Social Cognition and its Neural Overlap with General Verbal Semantic Cognition

One ALE meta-analysis revealed that non-verbal ToM processing engaged the ATL, the middle temporal gyrus (MTG), the pMTG and the vAG/TPJ in both hemispheres. Additionally, bilateral IFG, including Pars Triangularis and Orbitalis, as well as more posterior and dorsal portions of the frontal lobe, were also sensitive to non-verbal mentalisation (**Figure 3, left side**). The second ALE meta-analysis revealed that processing Social Cues recruited a similar bilateral network as in non-verbal ToM, including pMTG and vAG/TPJ, except for the ATL. Whilst the ATL was involved bilaterally in non-verbal ToM, the processing of non-verbal Social Cues did not engage the ATL. Another difference between the two meta-analyses was that the right IFG was engaged only by non-verbal ToM processing. Activation coordinates for all ALE meta-analyses involving social cognition are reported in **Supplementary Table 7**.

**Figure 3.**
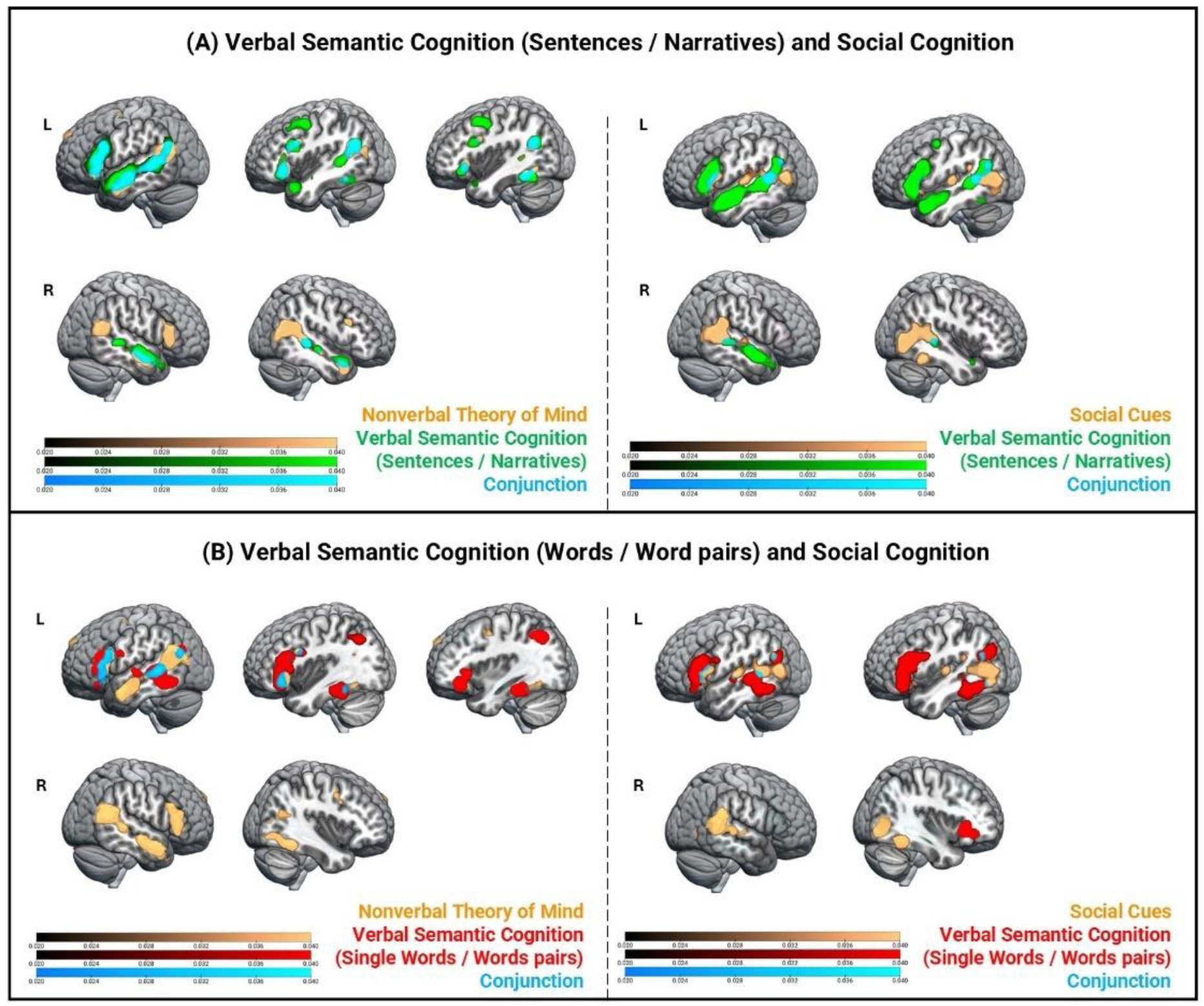
Neural overlap between Social Cognition and General Verbal Semantic Cognition. **(A)** Brain regions activated during social cognition (brown; left panel – non-verbal Theory of Mind, right panel – Social Cues, against respective active baselines), general verbal semantic cognition (semantic *versus* non-semantic or less semantic tasks) with sentences/narratives (green), and their neural overlap calculated via conjunction analysis (cyan), overlaid; Colourbar: ALE-values; cluster forming threshold: *p* < .001 (uncorrected); cluster extent correction: FWE *p* < .001. Conjunction: ALE-values, *p* < .001 (uncorrected), minimum cluster volume: 200 mm^3^. **(B)** Brain regions activated during social cognition (brown; left panel – Theory of Mind, right panel – Social Cues, against respective active baselines), general verbal semantic cognition (semantic *versus* non-semantic or less semantic tasks) with single-words/word-pairs (red), and their neural overlap calculated via conjunction analysis (cyan), overlaid; Colourbar: ALE-values; cluster forming threshold: *p* < .001 (uncorrected); cluster extent correction: FWE *p* < .001. Conjunction: ALE-values, *p* < .001 (uncorrected), minimum cluster volume: 200 mm^3^.

The first conjunction analysis was conducted to examine whether the right language network engaged during meaning processing in sentences/narrative stimuli would be similarly recruited also during non-verbal ToM processing (**Figure 3A, left)**. The results revealed several brain regions commonly recruited, independently from the modality (verbal or non-verbal) and the type of tasks (semantic task with sentences/narratives and mentalisation task). In the right hemisphere, the ATL, MTG and pMTG were similarly engaged during the processing of meaning in sentences and narratives and non-verbal mentalisation. Unexpectedly, we did not observe joint activation of the right vAG/TPJ, as this area was only activated during non-verbal ToM tasks. Along the same lines, the right IFG was only activated during non-verbal ToM tasks. In the left hemisphere, a wider portion of the language network showed joint activation, including the IFG Pars Orbitalis, Pars Triangularis and Pars Opercularis, as well as the MTG, vAG/TPJ, and pITG.

The second conjunction analysis was conducted to examine whether the right language network during meaning processing in sentences/narrative stimuli would be similarly recruited also during the perception of Social Cues, i.e., perception of social stimuli but without explicit instructions to make inferences about characters or perform mentalisation (**Figure 3A, right**). Within the right language network, only the pMTG showed joint activation. Instead, in the left hemisphere, processing of both social cues and meaning in sentences/narratives activated the IFG Pars Opercularis and Pars Triangularis, as well as the pMTG.

To ensure that the observed neural overlap between verbal semantic cognition and social cognition—particularly in the right ATL—was driven by similarities in social cognitive processes rather than semantic processing, we also conducted a conjunction analysis between non-verbal ToM processing and verbal semantic cognition using only single-word/word-pair stimuli (**Figure 3B, left**). Because the latter should engage semantic processing but not social cognition, any neural overlap would likely suggest that activation of the right language network, and especially the right ATL, is not specific to social cognition. As expected, the results showed no joint neural activation in the right hemisphere, suggesting that the right ATL may have a role in mentalisation.

A broader portion of the left hemisphere showed joint activation, including the IFG (Pars Opercularis and Pars Triangularis) and the pMTG – key brain regions of the semantic network (Lambon Ralph et al., 2017). Additionally, also the left vAG/TPJ which has been associated to both semantic processing (Binder, 2009) and mentalisation (Molenberghs et al., 2016; Schurz et al., 2014; Diveica 2021) showed joint activation.

We then performed a conjunction analysis to reveal the neural overlap between the brain regions involved the processing of Social Cues and those involved in semantic tasks with single-words or word-pairs (**Figure 3B, right)**. The results revealed no neural overlap in the right hemisphere. In the left hemisphere, the neural overlap was observed in vAG/TPJ and brain regions of the semantic network (Lambon Ralph, et al, 2017), including the left IFG (Pars Triangularis and Pars Opercularis), as well as the left pMTG and pITG.

Finally, for completeness, we also performed a conjunction analysis between non-verbal ToM and Social Cues processing. The results are reported in **Supplementary Figure 3.** This conjunction analysis aimed to reveal the brain areas that are active during perception of social content in the stimuli, independently from explicit task instructions to perform mentalisation on the stimuli. The right pMTG showed joint activation, alongside with its left analogue. Another brain region that was similarly engaged in both social cognition tasks was the left IFG Pars Opercularis.

### 3.4. Verbal Semantic Cognition – task demands

After examining the effect of increased semantic control we also examined the effect of overall task difficulty (Stefaniak et al., 2021) on the neural network of general verbal semantic cognition. The results from the four ALE meta-analyses revealed a minimal overlap between the language network and the MDN (Fedorenko et al., 2013) (**Figure 4A**). Activity in brain areas overlapping with the MDN was observed only when semantic tasks loaded on external cognitive control demands (e.g., tasks requiring a semantic decision). However, this effect was not independent of the type of stimuli as we initially predicted. High task demands modulated activity in brain areas overlapping with the MDN only when the semantic task involved single-words/word-pairs. This modulation was observed in the Insula in the right hemisphere, and in the pITG and IFG Pars Opercularis and Triangularis in the left hemisphere. The peak coordinates for all ALE meta-analyses on high and low task demands are reported in **Supplementary Table 8.**

**Figure 4.**
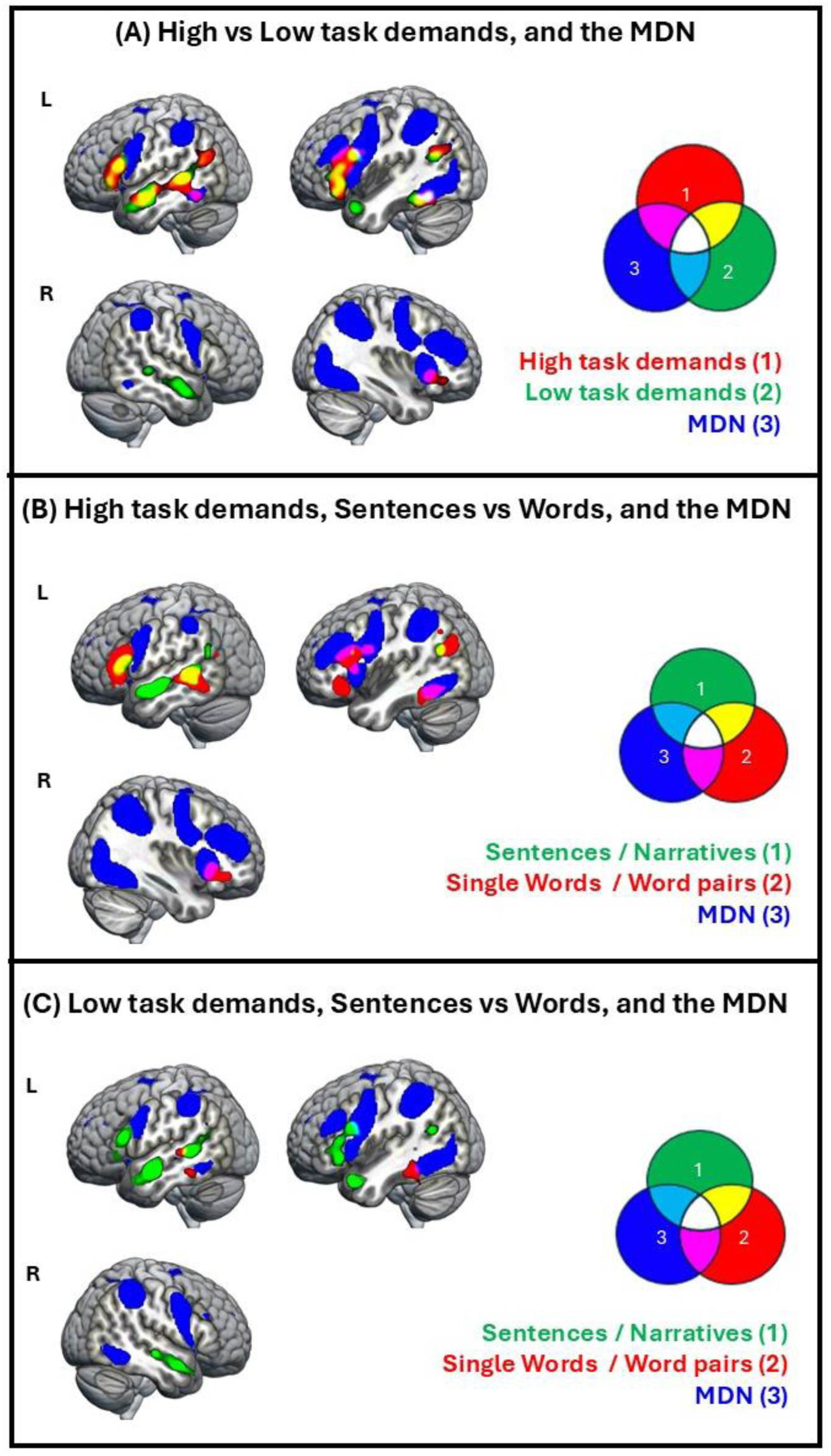
Brain areas activated during high *versus* low task demands in verbal semantic tasks, overlaid with the Multiple-Demand Network (MDN). **(A)** Brain regions active during semantic tasks with high task demand (red), low task demand (green), (cluster forming threshold: *p* < .001 (uncorrected); cluster extent correction: FWE *p* < .001), and the MDN identified by Fedorenko et al., (2013), overlaid. **(B)** Brain regions active during semantic tasks with high task demands, split by stimuli type (sentences/narratives – green, single-words/word-pairs – red), (cluster forming threshold: *p* < .001 (uncorrected); cluster extent correction: FWE *p* < .001), and the MDN, overlaid. **(C)** Brain regions active during semantic tasks with low task demands, split by stimuli type (sentences/narratives – green, single-words/word-pairs – red), (cluster forming threshold: *p* < .001 (uncorrected); cluster extent correction: FWE *p* < .001), and the MDN, overlaid. Colours codes of overlaps are explained in the Venn diagrams.

## 4. Discussion

In this ALE meta-analysis study, we investigated for the first time whether the right language network is involved during verbal semantic processing as an additional resource for increased semantic demands, and/or in response to separate, social cognition processing demands. In addition, we examined whether brain regions active during verbal semantic processing show overlap with the MDN or reflect domain-specific responses to semantic and mentalising tasks. A second main aim of this study was to establish whether MDN recruitment depends on external task demands rather than semantic control demands. The key findings on the functional specialisation of the right language network and MDN for verbal semantic cognition are discussed below, along with the discussion of additional findings outside the territories of the right language network and the MDN.

### 4.1. Verbal semantic processing engages the right language network differentially for sentences/narratives *versus* single-words/word-pairs

We demonstrated that the right language network is recruited differentially during semantic processing depending on whether the stimuli consist of single-words/word-pairs or sentences/narratives. In detail, the right ATL, MTG and pMTG were engaged during the semantic processing of sentences/narratives processing, but did not respond to single-words/word-pairs. Whilst the detailed interpretation of this result— namely, whether increased engagement of the right language network during sentence and narrative processing reflects heightened semantic control demands, involvement in social cognition, or both—will be discussed in depth in the following sections, it is worth noting that this finding aligns with previous studies showing that sentence and narrative processing often engages both the left and right language networks (Branzi et al., 2020; Branzi et al., 2023; Xu et al., 2005).

The differential recruitment of the right temporal lobe between sentences/narratives *versus* single-words or word-pairs did not depend on differences in the type of baselines. In fact, since the rest baseline may reduce the likelihood of observing ATL semantic-related activation, studies that employed rest as a baseline for semantic contrasts were excluded from both datasets.

Furthermore, we also demonstrated that the differential recruitment of the right temporal lobe between sentences/narratives *versus* single-words/word-pairs did not depend on differences in the type of tasks associated with sentences/narratives *versus* single-words or word-pairs. When we computed the same analysis but on a subset of the sentences/narratives and single-words/word-pairs datasets, focusing only on studies where participants were required to make a semantic choice, the results were consistent with those obtained when all tasks were included.

Therefore, the differential recruitment of the right temporal lobe between sentences/narratives *versus* single-words or word-pairs, which is unlikely to be due to differences in baseline conditions and/or tasks, aligns with our hypothesis that sentences and narratives would lead to more extensive recruitment of the right language network.

It is important to note that the recruitment of the right temporal lobe for meaning processing in sentences/narratives was particularly evident with auditory stimuli. Instead, semantic tasks including sentence/narrative stimuli presented in visual (written modality) activated mainly the left ATL. The left bias for written stimuli found in this study replicates previous results of auditory (bilateral) *versus* written (left) word processing (Marinkovic et al., 2003; Rice et al., 2015; Visser & Lambon Ralph, 2011). Accordingly, a recent ALE meta-analysis revealed that auditory stimuli during semantic processing engaged the ATL bilaterally, while visual stimuli engaged primarily the left ATL (Balgova et al., 2024).

Balgova et al. (2024) proposed that the recruitment of bilateral ATL during the processing of auditory stimuli could be understood in terms of general processing effort. This is because auditory stimuli are mainly sentences that require both rapid processing of individual words and combinatorial meaning, which could work the semantic system more vigorously than other stimuli. Be that as it may, here we show a substantial difference in the recruitment of the right temporal lobe between written and auditory sentences which does not fully support this hypothesis.

Therefore, one possibility is that the right ATL is more strongly engaged during auditory stimuli because spoken verbal stimuli involve human voice recognition, allowing listeners to automatically infer details and activate social knowledge about a speaker’s identity, such as their sex and age. Therefore, as compared to written stimuli, auditory stimuli may represent a more inherently social format, which may lead to the recruitment of the right ATL.

Finally, we found that the right Insula responded to semantic processing during single-words/word-pairs but not to sentence/narrative stimuli. The right Insula is part of the MDN (Duncan 2010; Fedorenko et al., 2013). In language tasks, this brain region has been associated to executive control processes such as inhibition of the non-target meaning of ambiguous words (Mason & Just, 2007) and update of semantic representation following semantic violations (Branzi, Humphreys, et al., 2020; Branzi & Lambon Ralph, 2023). Accordingly, in the present study we found that the right Insula responded to both semantic and non-semantic (external/task) demands, consistent with the proposal that it may reflect domain-general control processes (Duncan, 2010; Fedorenko et al., 2013).

The observation that the right Insula is active only when semantic tasks require response suppression and conflict resolution (i.e., tasks with high external task demands) indicates that this brain region is unlikely to play a central role in core language functions such as lexical/semantic retrieval and combinatorial semantics. Instead, it may be engaged to reduce interference during response selection, particularly when competition between target and non-target semantic representations is artificially heightened by task demands.

### 4.2. Semantic control modulates activity in the right IFG

Our first hypothesis was that the right language network would support the same functions as their homologue regions in the left hemisphere, and areas within the right language network, particularly the right pMTG, the right IFG and the right dAG/IPS would be recruited when semantic control demands increase (Lambon Ralph et al., 2017).

Accordingly, we observed the recruitment of the right IFG Pars Opercularis and Pars Triangularis in response to increased semantic control demands during tasks including sentences/narratives as stimuli. These results align with recent studies that have shown increased activity in the right IFG as a function of an increase in semantic control demands during integration of meaning across multiple stimuli (sentence processing) (Branzi, Humphreys, et al., 2020; Silbert et al., 2014). The neural activity measured in the right IFG Pars Opercularis and Pars Triangularis was accompanied by a similar but more pronounced difficulty-related effect in the left IFG, in line with previous reports (Jung et al., 2021; Krieger-Redwood et al., 2015; Lambon Ralph et al., 2017; Quillen et al., 2021; Wu & Hoffman, 2024).

Interestingly, increased activity in the right IFG was not observed for semantic control during single word/word pairs tasks. This discrepancy may arise because, compared to single-word or word-pair tasks, combining meanings across sentences likely imposes greater semantic control demands, for which left IFG engagement alone may not be sufficient to optimize performance. In line with the hypothesis that right IFG may be recruited only when semantic control demands are especially high, the right IFG Pars Opercularis/Pars Triangularis was not activated in the general verbal semantic cognition ALE meta-analysis.

The additional involvement of right IFG, relative to the left IFG, during semantic control is in accord with the ‘variable neuro-displacement’ hypothesis (Jung et al., 2021; Stefaniak et al., 2020). This proposes that cognitive systems are formed with dynamic, spare processing capacity, which balances energy consumption against performance requirements and can be resilient to changes in performance demands. In the context of the present data, healthy language processing would be supported by both left and right language networks, with a bias towards the left system (see results from the general verbal semantic cognition ALE meta-analysis). The right language areas would be downregulated unless the system is under high pressure (increase of semantic control demands during sentences/narratives) to optimise performance.

Contrary to our hypotheses, we found no evidence of involvement of the right pMTG and the right dAG/IPS in response to increased semantic control demands. Although few studies have found the right dAG/IPS activity to be modulated by semantic control demands (Branzi et al., 2020; Jung et al., 2021), many others have failed to show consistent involvement of this region in any aspect of semantic cognition (Jackson, 2021). This observation does not uniquely apply to the right dAG/IPS but also to the left dAG/IPS (Jackson, 2021).

Since dAG/IPS is part of the MDN (Duncan, 2010; Fedorenko et al., 2013), the occasional identification of dAG/IPS during semantic tasks may be due to domain-general control requirements imposed by the task at hand rather than semantic control. Accordingly, in our study we show that increased external task demands, rather than semantic control demands, modulate neural activity in left dAG/IPS.

In the present study, we found that the social content of the stimuli rather than semantic control demands upregulated the neural activity measured in the right pMTG. This result will be discussed in greater detail in the next section. However, it is worth noting that, while previous studies have shown that the right pMTG responds to increased semantic control demands (Branzi et al., 2020; Jung et al., 2021), this study and earlier reports suggest that such evidence is inconsistent (Jackson, 2021). The activity observed in the right pMTG, modulated by semantic control, may be due to the semantic tasks involving stimuli with social content (Branzi et al., 2020; Jung et al., 2021), which could have engaged social cognition processes.

Importantly and in line with our hypothesis, only a minimal overlap between the MDN and the semantic control network was observed. This overlap concerned the left dorsal IFG. These results align with previous data showing some involvement of MDN during semantic tasks (Branzi, Humphreys, et al., 2020; Branzi & Lambon Ralph, 2023; Hodgson et al., 2021). Here, we demonstrate that this brain region is engaged in semantic tasks only when lexical semantic competition between representations is artificially heightened due to task design choices, rather than reflecting fundamental verbal semantic processes.

### 4.3. Semantic control beyond the right hemisphere: Stimulus type modulates the recruitment of the left posterior temporal lobe

We observed that the semantic control network fractionates depending on the type of stimuli not only across the left *versus* right dimension but also the dorsal v*ersus* ventral dimension. In the left posterior middle temporal lobe, the pMTG and pITG exhibited a graded shift in their preference for controlling meaningful stimuli, with the pMTG being more involved in sentence and narrative processing, while the pITG showed a stronger preference for tasks with single words and/or word pairs.

Previous studies have demonstrated that pITG responds particularly to hard phonological and semantic tasks (Hodgson et al., 2021) and is implicated in the extended MDN (Assem et al., 2020). It is unclear if this region reflects shared control processes for language subdomains or rather across a broad set of linguistic and non-linguistic domains.

Our results suggest that this brain region may reflect some sort of language-specific control. In fact, in our study the left pITG was (i) recruited during semantic processing of word stimuli irrespective of the external task demands and (ii) found sensitive to semantic control demands. Accordingly, Hodgson et al. (2021) found left pITG implicated in both phonological and semantic control, but not in nonverbal working memory. This, along with the fact that the left pITG has been found in MDN assessments which did not specifically exclude verbal stimuli (Assem et al., 2020, Fedorenko et al., 2013), leaves open the possibility that left pITG might support language-specific control processes.

However, left pITG has been also implicated also in non-verbal executive function tasks, such as task-switching (Lemire-Rodger et al., 2019; Schumacher et al., 2019). Therefore, it may reflect a domain-general function shared with task-switching paradigms, such as for instance inhibition of the competing (semantic or non-semantic) representations. This function may be particularly required in tasks with single-words and/or word-pairs which often include target words presented concurrently with competing semantic distractors that need to be inhibited to perform the semantic task.

### 4.4. The right temporal lobe supports both verbal semantics, but only for stimuli with social content

Sentences and narratives typically provide an acting/feeling character(s) and ‘schema or situation models’, i.e., mental representations that summarise the interaction among entities and the environment in a scene or event (Ranganath & Ritchey, 2012). In this context, it is likely that, besides semantic processing, language comprehension engages social cognition to support mentalisation and the processing of social cues.

Accordingly, we hypothesised that some parts of the right language network previously associated with social cognition, i.e., right vAG/TPJ, the right ATL (Olson et al., 2013; Saxe et al., 2004; Saxe & Wexler, 2005; Turker et al., 2023), and the right pMTG (e.g., Hodgson et al., 2022) would be engaged during semantic processing, but only with stimuli with social content. We tested this hypothesis by examining the neural overlap between verbal semantic cognition [measured in social (sentences and/or narrative tasks) and non-social (single-words/word-pairs) stimuli, as separately] and social cognition (measured in non-verbal ToM and in Social Cue processing tasks, as separately).

Contrary to our hypotheses, we did not find any overlap in the right vAG/TPJ between the social cognition networks (non-verbal ToM and Social Cue processing) and the verbal semantic network. In line with previous results (Balgova et al., 2024; Balgova et al., 2022; Hodgson, 2022), the right vAG/TPJ seems to respond specifically to social cognition. However, the overlap between the non-verbal ToM networks and the verbal semantic network was observed in the left vAG/TPJ, but only for sentences/narratives.

The laterality of vAG/TPJ involvement in social cognition is unclear. Neuroimaging evidence has shown that ToM engages this area bilaterally (Diveica et al., 2021). Yet, some have argued that the selectivity of this region for ToM is in the right hemisphere (Saxe & Wexler, 2005). Others, instead, have argued that selectivity is in the left hemisphere (Aichhorn et al., 2006, 2009). In semantic cognition, activation of this region is generally left lateralised for verbal stimuli (Branzi et al., 2021; Branzi, Humphreys, et al., 2020; Branzi & Lambon Ralph, 2023; Humphreys et al., 2024; Humphreys & Lambon Ralph, 2015). Our results further demonstrate this is the case, but only when semantic cognition involves sentences/narratives, i.e., stimuli with social content. This may indicate that left vAG/TPJ during semantic tasks has a role in social and mental inference-making processes.

Additionally, and in accord with our hypotheses, we found that the right ATL (including the temporal pole) was similarly engaged in non-verbal ToM processing and verbal semantic processing, but only when semantic processing involved stimuli with social content (verbal sentences/narratives). Instead, while the semantic processing of single-words/word-pairs engaged the left ATL (anterior STG), no activity was observed in the right ATL during semantic processing of single-words/word-pairs.

Finally, we observed activity in the right pMTG in response to (1) nonverbal tasks with explicit instructions to perform ToM, (2) verbal semantic tasks including sentences/narratives possibly resulting in participants performing mentalisation about the character(s), and finally, (3) tasks requiring the processing of Social Cues, but with no explicit instructions to perform mentalisation. All in all, these results indicate that the right but not the left ATL and pMTG are specifically recruited when social cognition is likely involved in the task.

Note that the similar recruitment of the right temporal lobe during both semantic and social cognition cannot be explained by the similarity in the type of stimuli employed. In fact, semantic cognition tasks and social cognition tasks included substantially different stimuli and, with exception of prosody processing, presented in different formats (verbal *versus* non-verbal).

The similar engagement of right temporal lobe regions in verbal semantic tasks and non-verbal ToM tasks might be because both engage semantic processing (see Balgova et al., 2024; Binney & Ramsey, 2020). According to this hypothesis, a recent ALE meta-analysis study of functional neuroimaging studies revealed a great overlap between the ALE maps for ToM and semantic cognition, after controlling for critical aspects such as baseline type, stimulus format and type of modality (Balgova et al., 2024). Interestingly, however, the overlap was mostly observed in brain areas that in our study were uniquely involved in sentences and/or narratives, e.g., the bilateral ATL including the temporal pole and anterior superior temporal sulcus, as well as right MTG, and left vAG/TPJ, all key regions for semantic processing (Binder et al., 2009).

Thus, our results indicate that at least another explanation for the overlap between ToM and semantic cognition may be viable. That is, processing of stimuli such as sentences and narratives may engage social cognition besides semantic processing. In fact, we found that only sentence and narrative tasks—likely involving a social cognition component—showed neural overlap (conjunction) with ToM tasks and tasks measuring social cue recognition in brain regions typically associated with social cognition. In contrast, single-word and word-pair tasks did not exhibit this overlap.

Therefore, although semantic cognition is integral to social cognition (Binney & Ramsey, 2020) and some neural overlap between these two domains can be explained as such, the neural overlap observed in the right ATL (anterior MTG), and the right pMTG may reflect similar engagement of social cognition, rather than semantic processes. This aligns with studies on both healthy participants and patients, which have shown that the right ATL may be particularly tuned to social cognition (Gainotti, 2015; Pobric et al., 2016; Rankin et al., 2006; Rice et al., 2015). For example, the execution of empathic tasks and emotion processing seem to be more impaired by atrophy of the right ATL (Papinutto et al., 2016; Rankin et al., 2006; Seeley et al., 2005). Accordingly, the right *versus* the left ATL presents stronger connectivity with the orbital frontal cortex via the uncinate fasciculus (Papinutto et al., 2016), i.e., brain structures known to be important for social cognition (Oishi et al., 2015; Rolls & Grabenhorst, 2008; Von Der Heide et al., 2013).

We propose that the ATL—including both STG and MTG—and particularly the right ATL—contributes to both nonverbal ToM tasks and semantic processing tasks with sentences by providing conceptual information to constrain inferences about the intentions and actions of others (Binney & Ramsey, 2020). This includes, for example, information about an actor’s state of mind, independently from whether an explicit descriptor is present. Instead, the right pMTG may reflect control processes to regulate and select conceptual information relevant for making social inferences.

### 4.5. Increased external task demands during verbal semantic processing recruit the MDN

Using language requires applying cognitive operations (e.g., retrieval, prediction, combinatorial processes) similar to those used in other cognitive domains. This has led to the hypothesis that the brain contains domain-general circuits responsible for carrying out these operations, with language relying on these circuits (Abutalebi & Green, 2007; Fitch & Martins, 2014; Green & Abutalebi, 2013; Hasson et al., 2018; Koechlin & Jubault, 2006). Whilst there is evidence for domain-general executive processes and MDN engagement in diverse linguistic phenomena including ambiguity processing (Mcmillan et al., 2013; Novais-Santos et al., 2007; Rodd et al., 2005), high surprisal (Shain et al., 2020) and grammatical violations (Kuperberg et al., 2003; Mollica et al., 2020; Nieuwland, 2012), some researchers have questioned the importance of domain-general executive processes and the MDN to support core language operations (for review, see Fedorenko & Shain, 2021).

First, our results are in line with previous reports by showing that some MDN brain regions are active during semantic cognition (Hodgson 2021; Branzi et al., 2023; Branzi et al., 2020). These brain areas include the dorsomedial prefrontal cortex (including supplementary motor and pre-supplementary motor area) (Branzi et al., 2022; Branzi, Martin, et al., 2020; Geranmayeh et al., 2014), the left pITG (Assem et al., 2020; Duncan, 2010), the right Insula, the left IFG Pars Opercularis and Triangularis bordering with the Premotor Cortex.

Second, and in accordance with proposals/evidence that the MDN does not support core language computations (Blank & Fedorenko, 2017; Branzi & Lambon Ralph, 2023; Diachek et al., 2020), our results indicate that rather than being modulated by semantic control demands specifically, MDN regions are modulated by external task demands (e.g., inhibiting concurrent distractors and making choices between multiple conflictual options) which often accompany core language demands (lexical retrieval, combinatorial semantics, etc).

In detail, we found that high task demands modulated activity in the right Insula, the left pITG and the left IFG Pars Opercularis and Triangularis. This result is in accord with previous studies that have shown that unless language processing is associated with a secondary task, the MDN is not recruited during verbal semantic processing. For instance, Wright et al. (2011) showed that some frontal MDN areas are engaged during a lexical decision task, but not passive listening to the same materials. Along the same lines, Diachek et al. (2020) compared the MDN’s engagement in language experiments that involved passive comprehension (visual or auditory) with its engagement in experiments that involved an additional task such as answering comprehension questions or determining semantic associations. The language network responded equally strongly during tasks with and without an additional task. Instead, the MDN was recruited only in the presence of an additional task.

Interestingly, the effect of external task demands on the recruitment of the DMN was not fully independent of the type of stimuli as we initially predicted. In fact, except for the left pITG, the overlap between verbal semantic processing and the MDN was only seen in high demand tasks with single-words/word-pairs as stimuli. This may be a result of overall task type differences between single-word/word-pair tasks and sentence/narrative tasks. Even if we carefully split the data in high versus low task demands following the same criteria for both types of stimuli (narratives/sentences and single-words/word-pairs), there may still be some differences in the external demands imposed by tasks typically associated with these stimuli. For example, sentences/narratives associated with a secondary choice task often included a yes/no question about the sentence or narrative, while single-word/word-pair tasks requiring a choice more often included a probe word, a target word, and one to three distractors. The presence of multiple-choice options likely puts more demands on domain-general control processes and the MDN (Assem et al., 2020; Duncan, 2010) to perform target-nontarget discrimination, shifts in attention and inhibition of non-target responses.

Overall, our results indicate that the verbal semantic network may include regions specifically involved in semantic representation (e.g., the ATL) and control (such as the ventral IFG and pMTG) of meaningful representations, as well as regions associated with domain-general control processes, including the left dorsal IFG, left pITG, and right Insula, which may only be recruited when external task demands are heightened.

## Limitations

The use of a meta-analysis approach allowed us to shed light on the broader organisation of the language network, and especially the role of the right language network for semantic processing, while also uncovering previously hidden relationships between different semantic, social cognition and domain-general executive control domains. While meta-analyses have the advantage of synthesising information from a wide range of studies and cognitive domains, they lack precision in spatial resolution. This may represent a limitation, especially when examining regions bordering each other. Therefore, the hypotheses on the role of the right language network proposed in the present study should be tested within individual participants.

## Conclusions and Future Directions

Our findings provide new insights into the relationship between semantic and social cognition, revealing that the recruitment of the right temporal lobes during semantic processing is influenced by the type of stimuli and likely their socialness. One possible explanation for these observations is that verbal semantic processing involving social stimuli may engage ToM processing and/or other aspects of social cognition and therefore activate the semantic network along with additional regions in the right hemisphere more tuned for stimuli and/or tasks that involve social cognition. Alternatively, these regions may belong to a single, widely distributed yet functionally integrated semantic network, with systematic variation in the engagement of specific nodes across hemispheres driven by task-related or stimulus-related factors, such as input modality and the social nature of the semantic stimuli. Our data are compatible with both explanations. Future studies will have to employ within-subjects designs and assess the factors that drive the recruitment of the right hemisphere within and across social and semantic cognition domains.

Furthermore, our findings revealed that the right IFG supports semantic control in sentence and narrative processing. Interestingly, non-verbal ToM processing also engaged the right IFG. One possible explanation for this result, in line with the semantic control results, is that ToM tasks, overall, may place greater semantic control demands compared to typical semantic tasks. In other words, ToM tasks could be classified as ‘hard semantic conditions’. Nonverbal ToM experiments often include tasks in which participants are asked to switch between their own and a different character’s visual, cognitive, and emotional perspective. Social stimuli that describe character’s mental states or intentions (Pexman et al., 2023), independently from modality, will likely activate a wide range semantic associations coupled with different perspectives, possibly creating incongruency with the observer’s own perspective. This might increase the overall demand for semantic control. That said, it remains unclear whether the right IFG activity observed during ToM processing reflects the same control processes as those involved in the semantic domain (Binney & Ramsey, 2020). Future research will need to address this question using within-subject designs to determine whether the distinction between knowledge representation and cognitive control observed in the semantic domain also applies to the social domain.

---

**Supplementary Figure 1.**
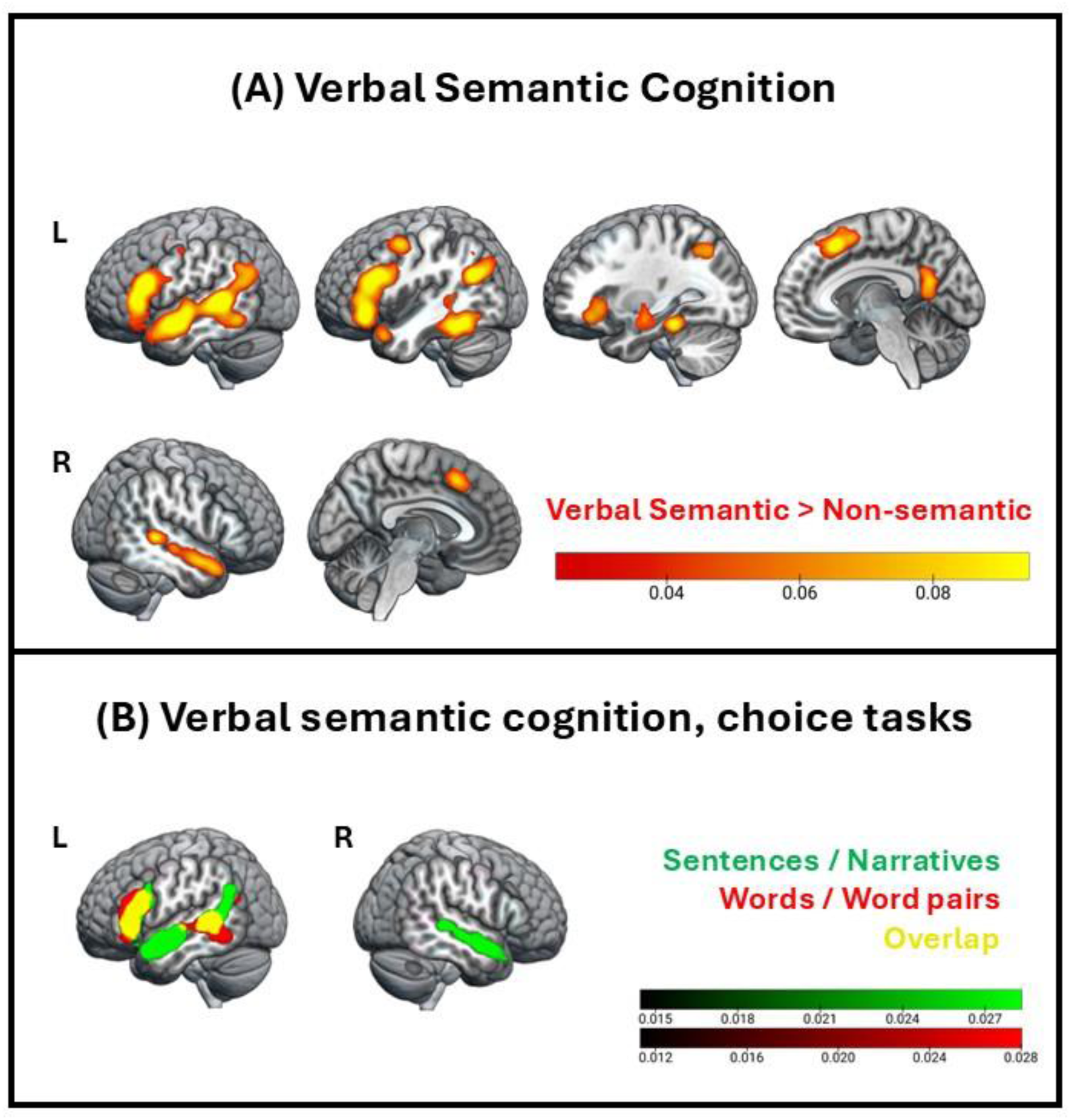
Semantic cognition. **(A)** Brain regions activated during all verbal semantic > non-semantic (or less semantic) tasks. Colour bar: ALE-values; cluster forming threshold: p < .001; cluster extent correction: FWE p < .001. **(B)** Brain regions activated during verbal semantic processing, only including tasks in which participants had to perform a choice, split by stimuli type (green: sentences/narratives, red: word-pairs/single words, yellow: overlapping brain regions). Colour bar: Z-scores, p <.001, minimum cluster volume: 200 mm.3

**Supplementary Figure 2.**
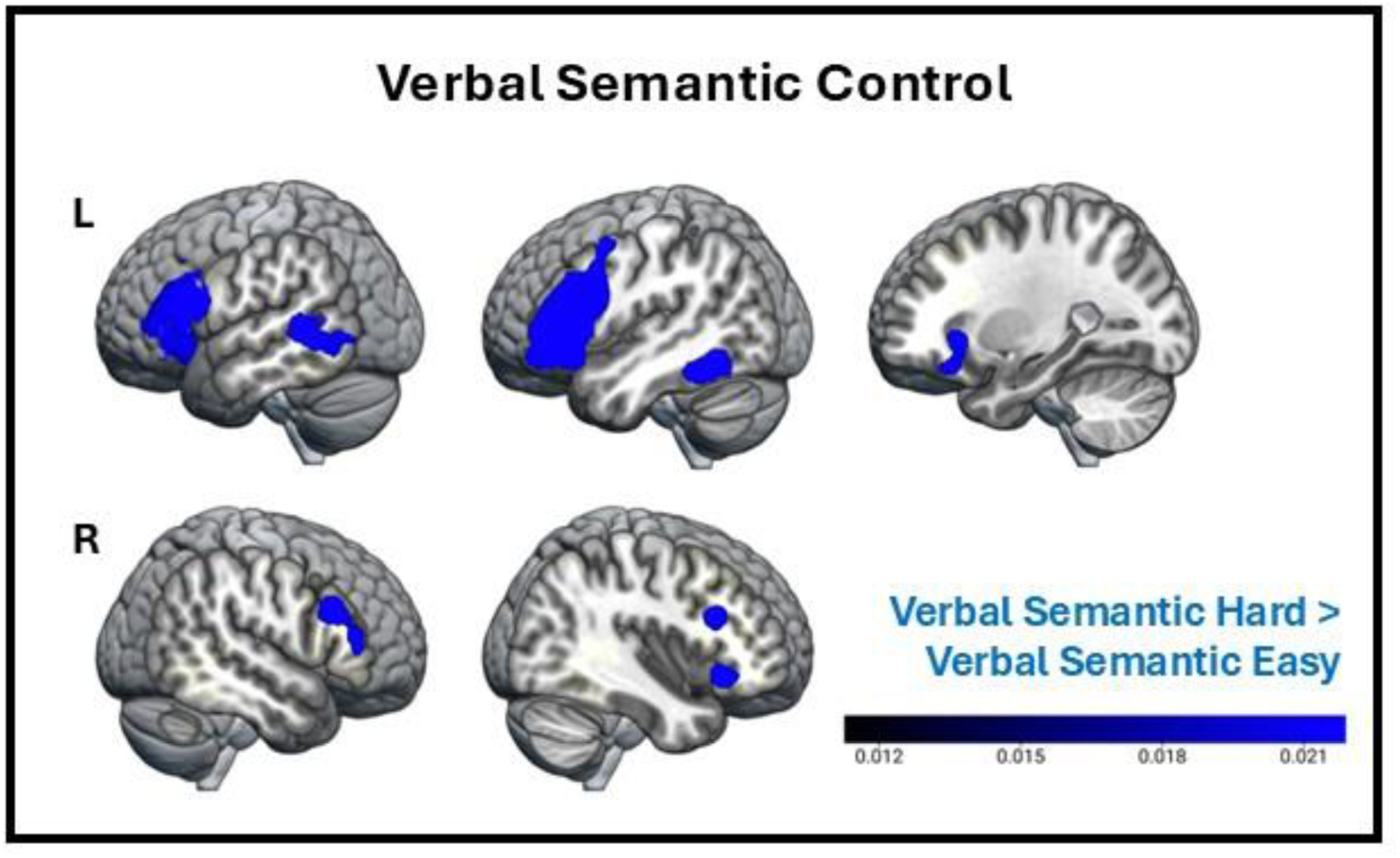
Brain regions reliably activated during all hard > easy semantic tasks or conditions. Colourbar: ALE-values; cluster forming threshold: *p* < .001 (uncorrected); cluster extent correction: FWE *p* < .001.

**Supplementary Figure 3.**
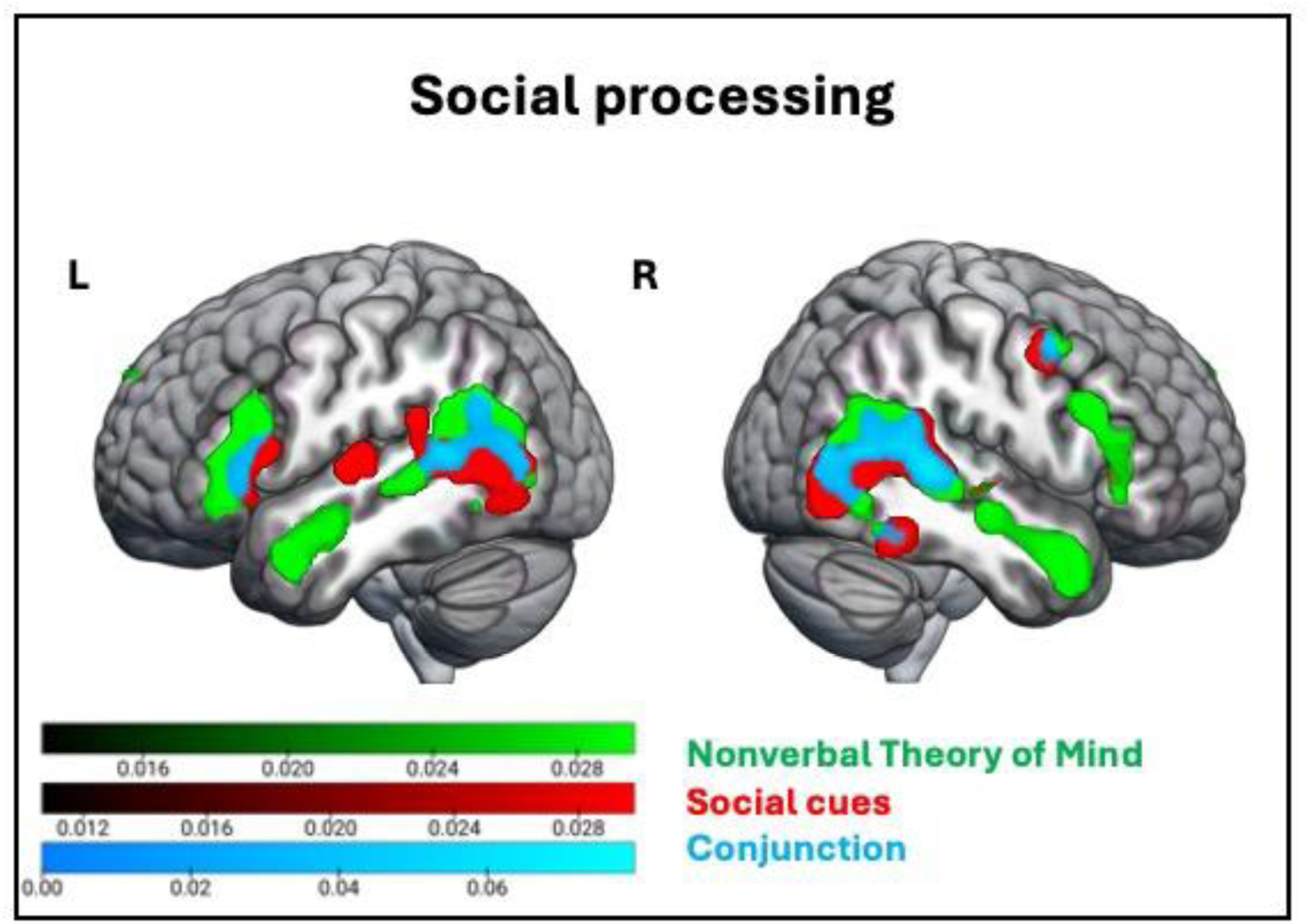
Social processing. Individual ALE-analyses and conjunction analyses of nonverbal Theory of Mind processing (green), Social cues processing (red), and their conjunction (blue). Colourbar: ALE-values; cluster forming threshold: p < .001 (uncorrected); cluster extent correction: FWE p < .001. Conjunction: ALE-values, p <.001, minimum cluster volume: 200 mm^3^.

**Supplementary Table 1.**
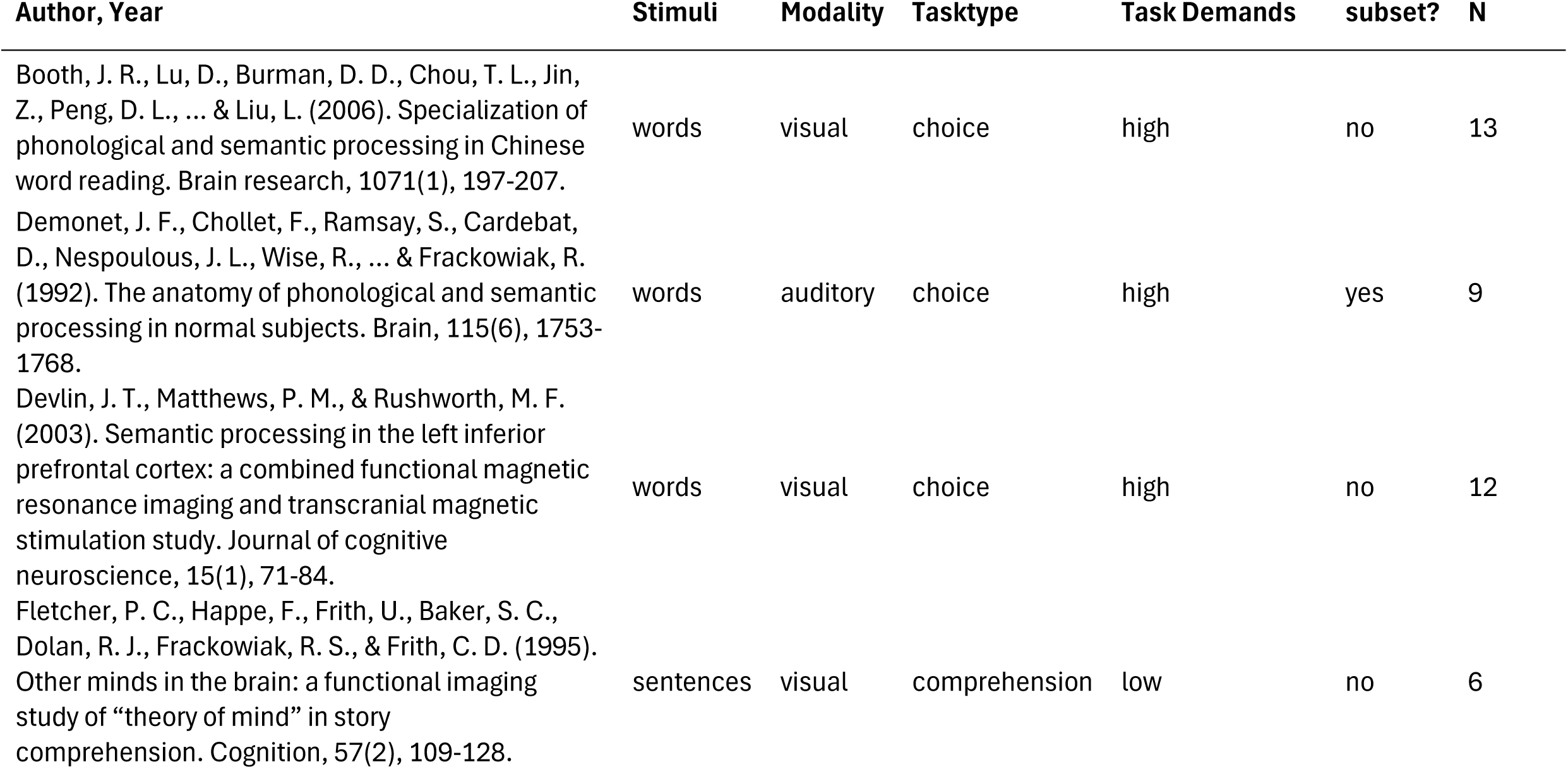
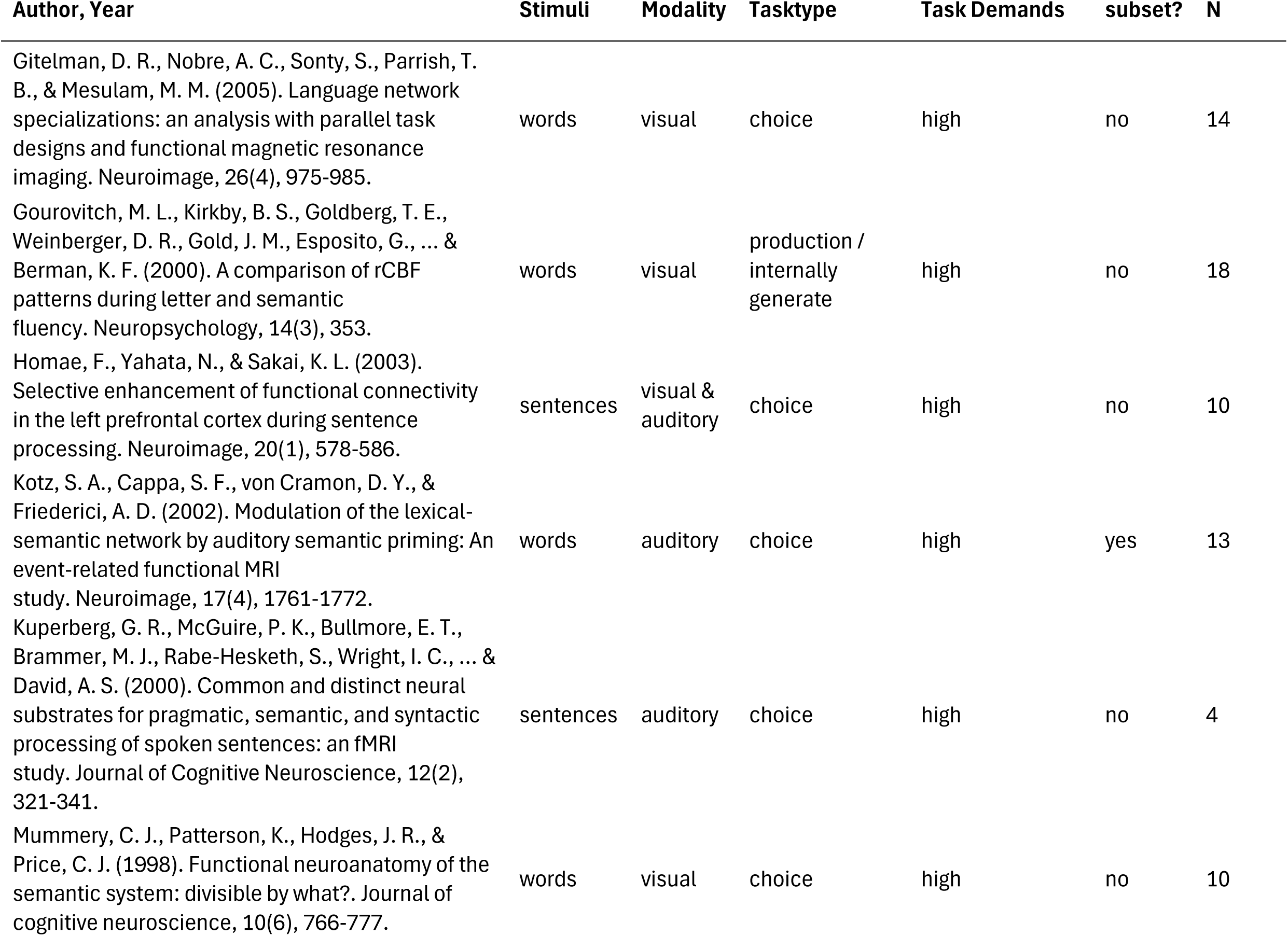
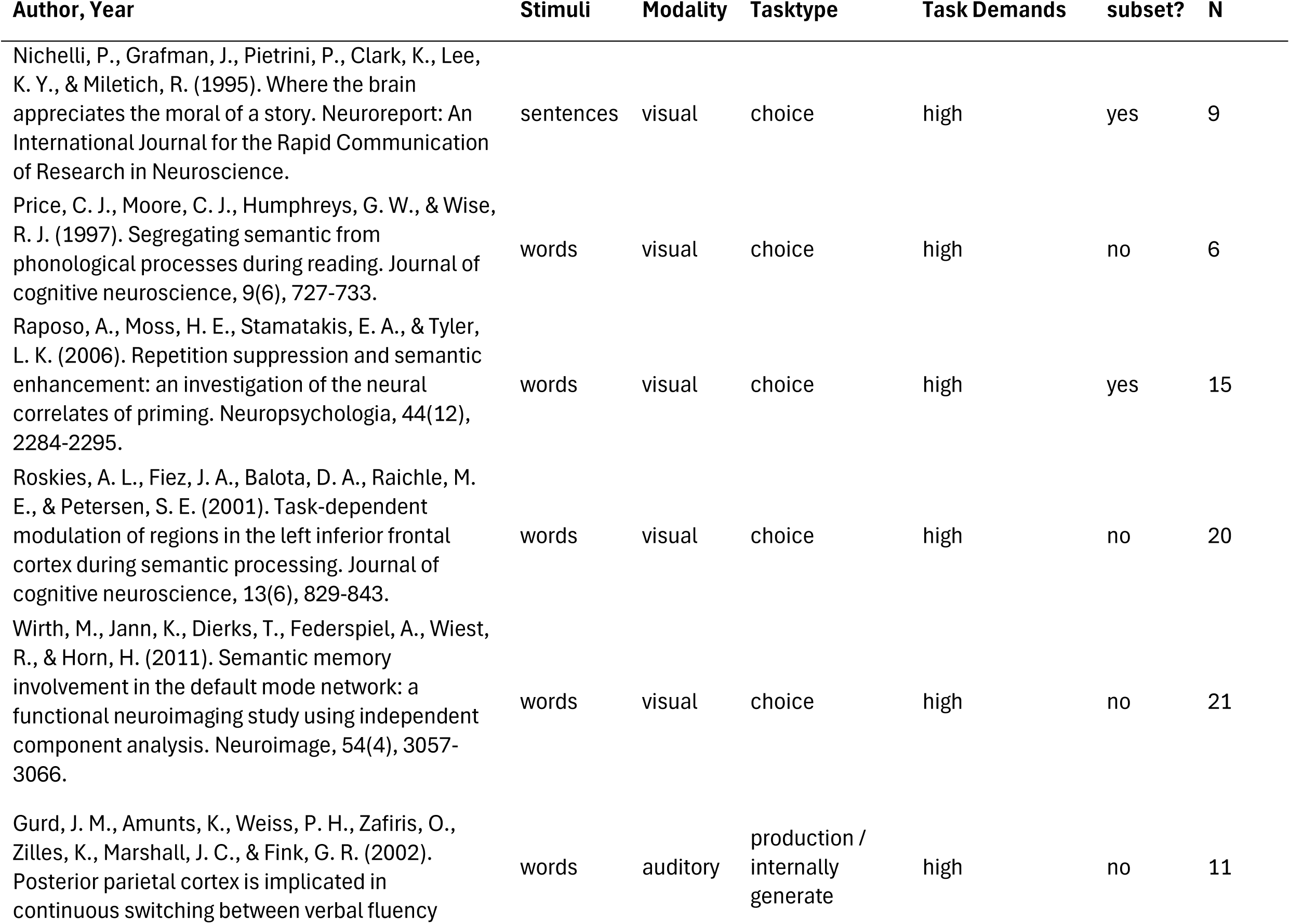
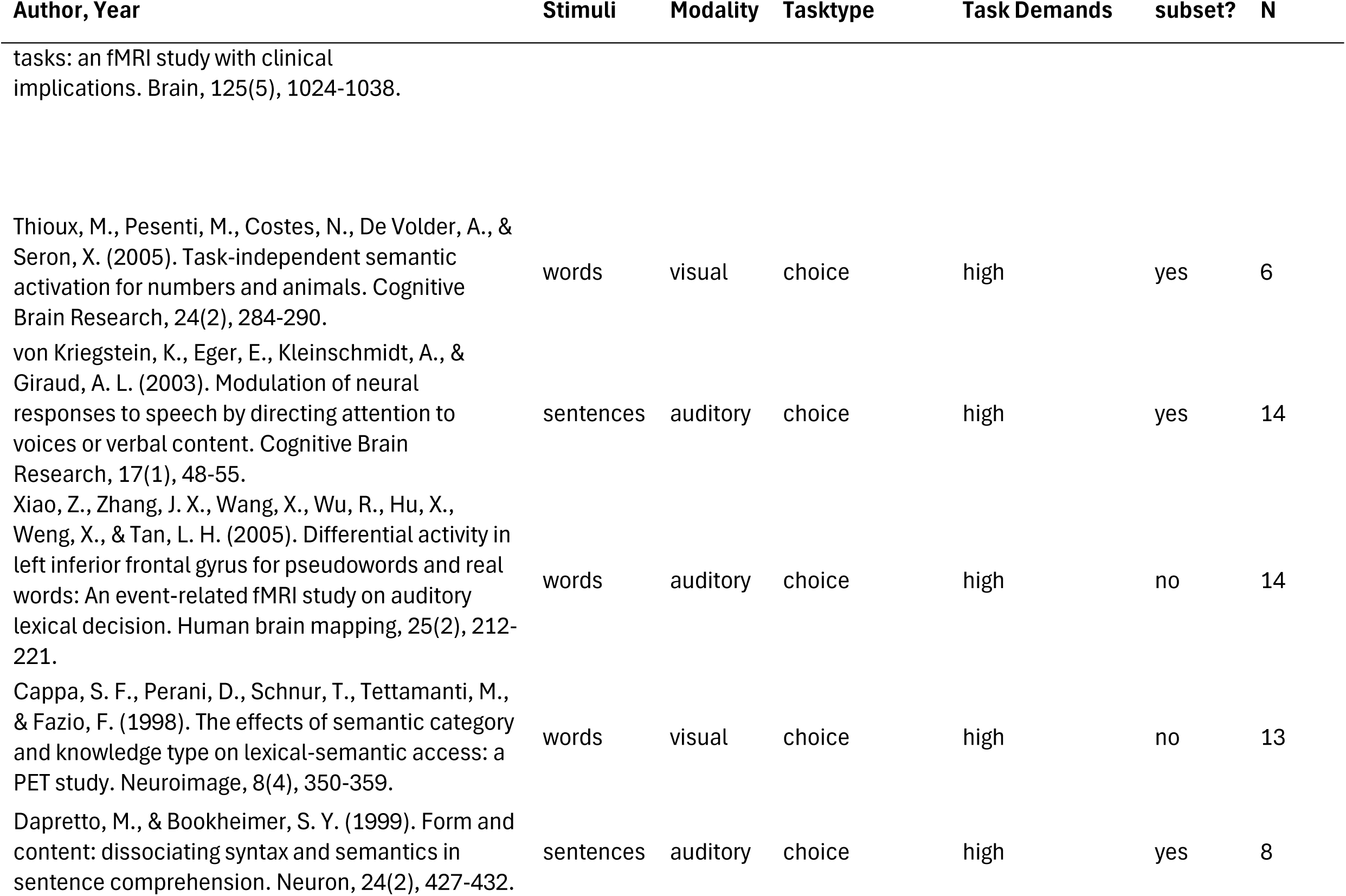
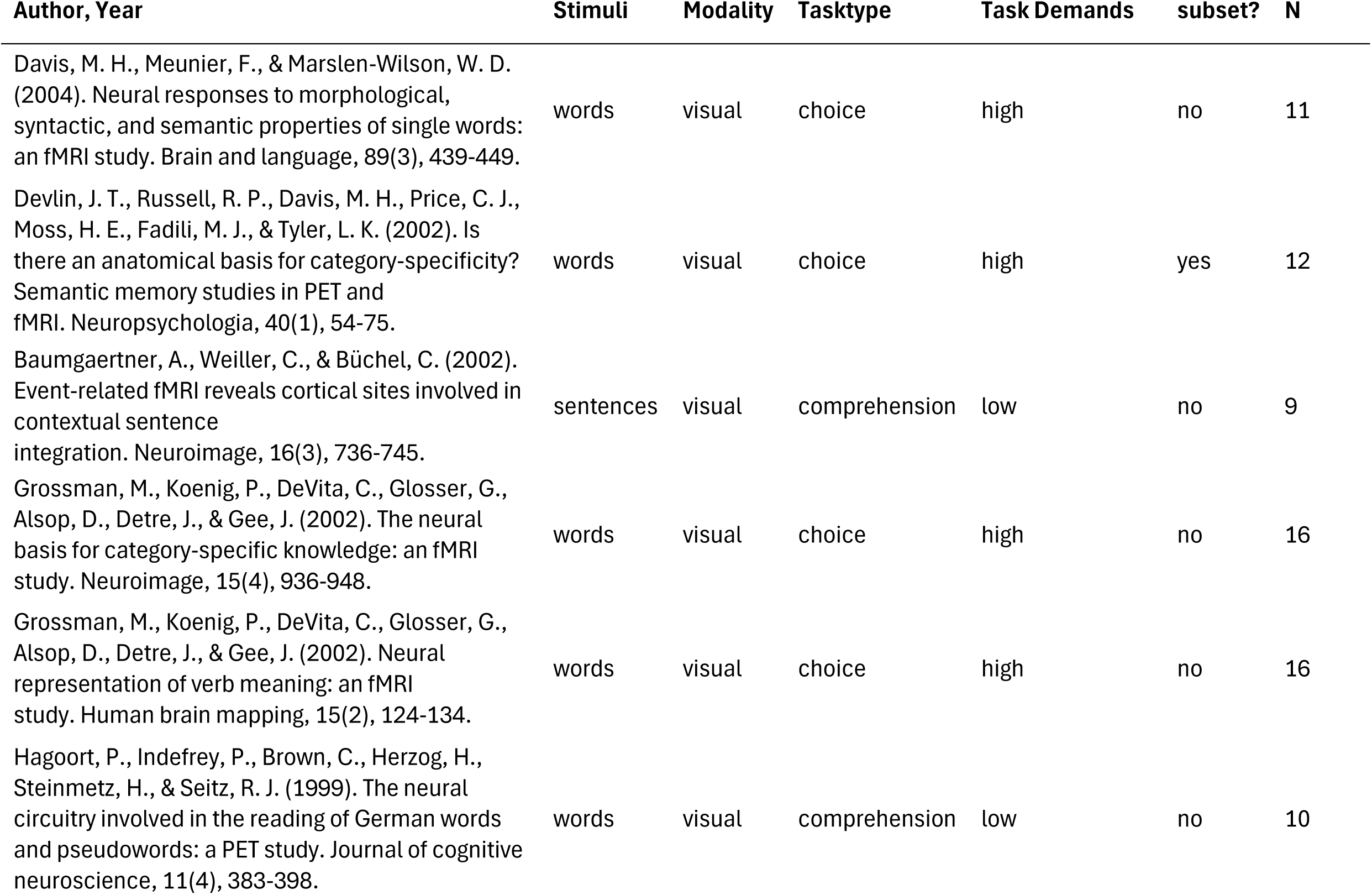
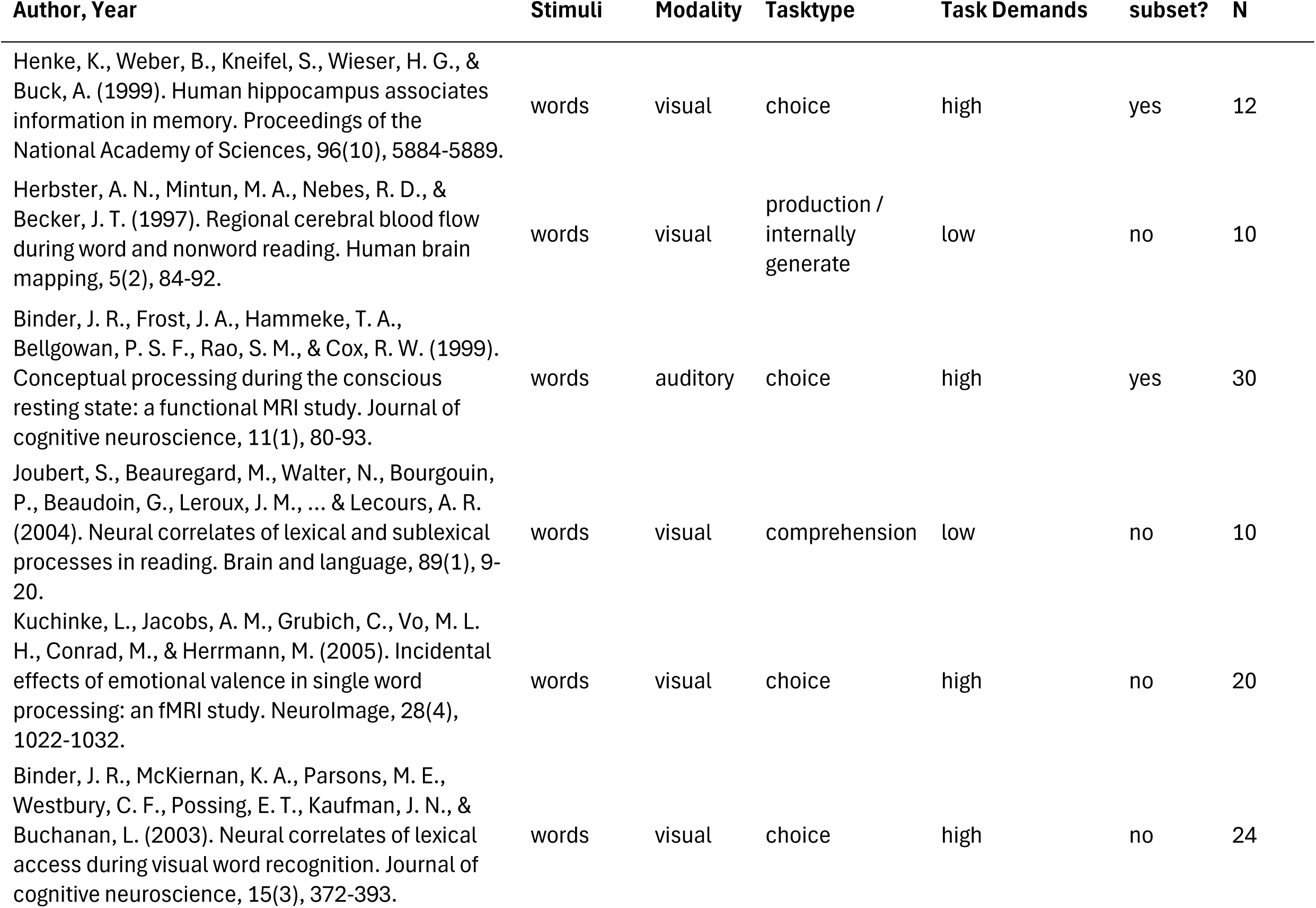
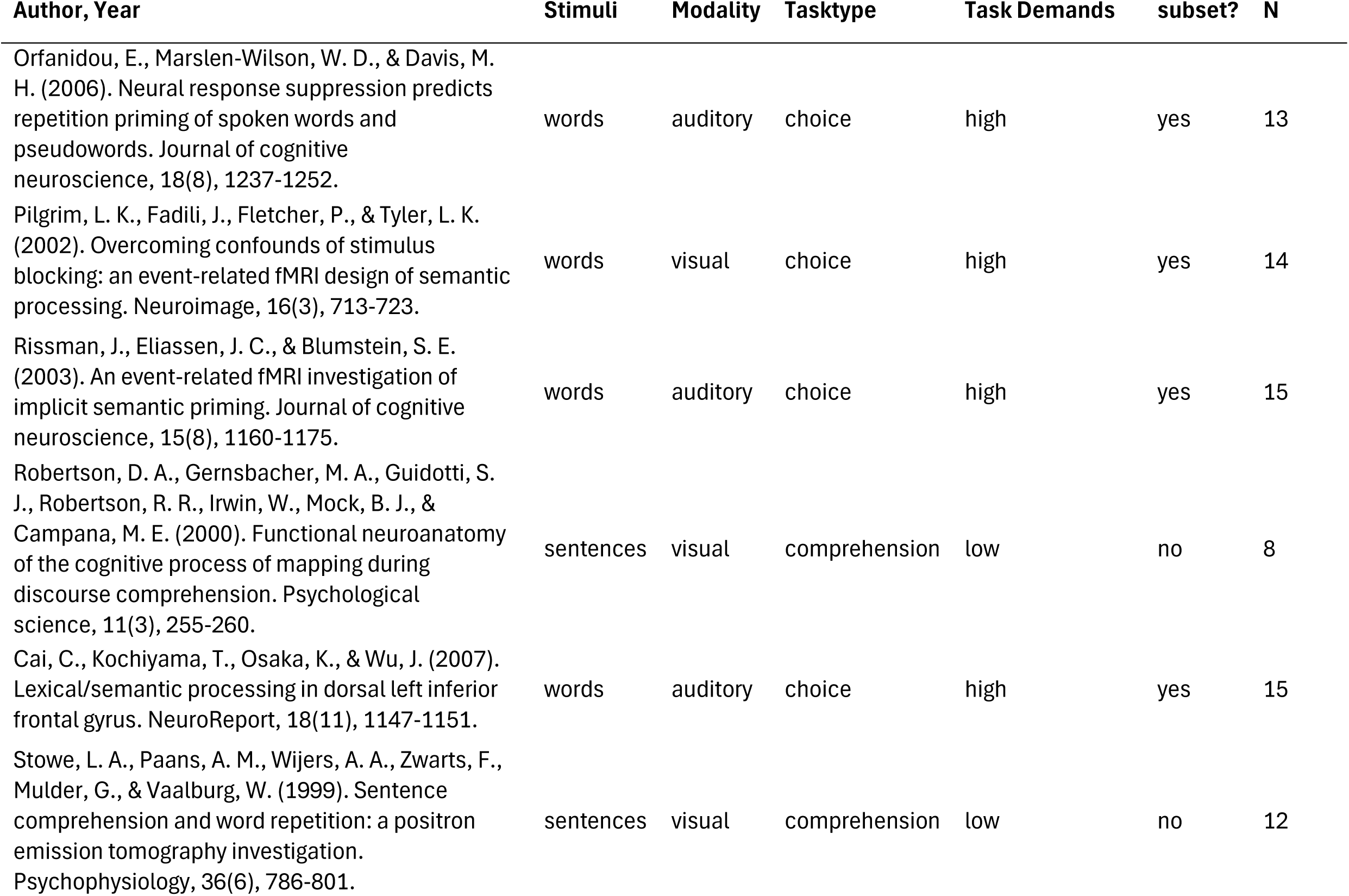
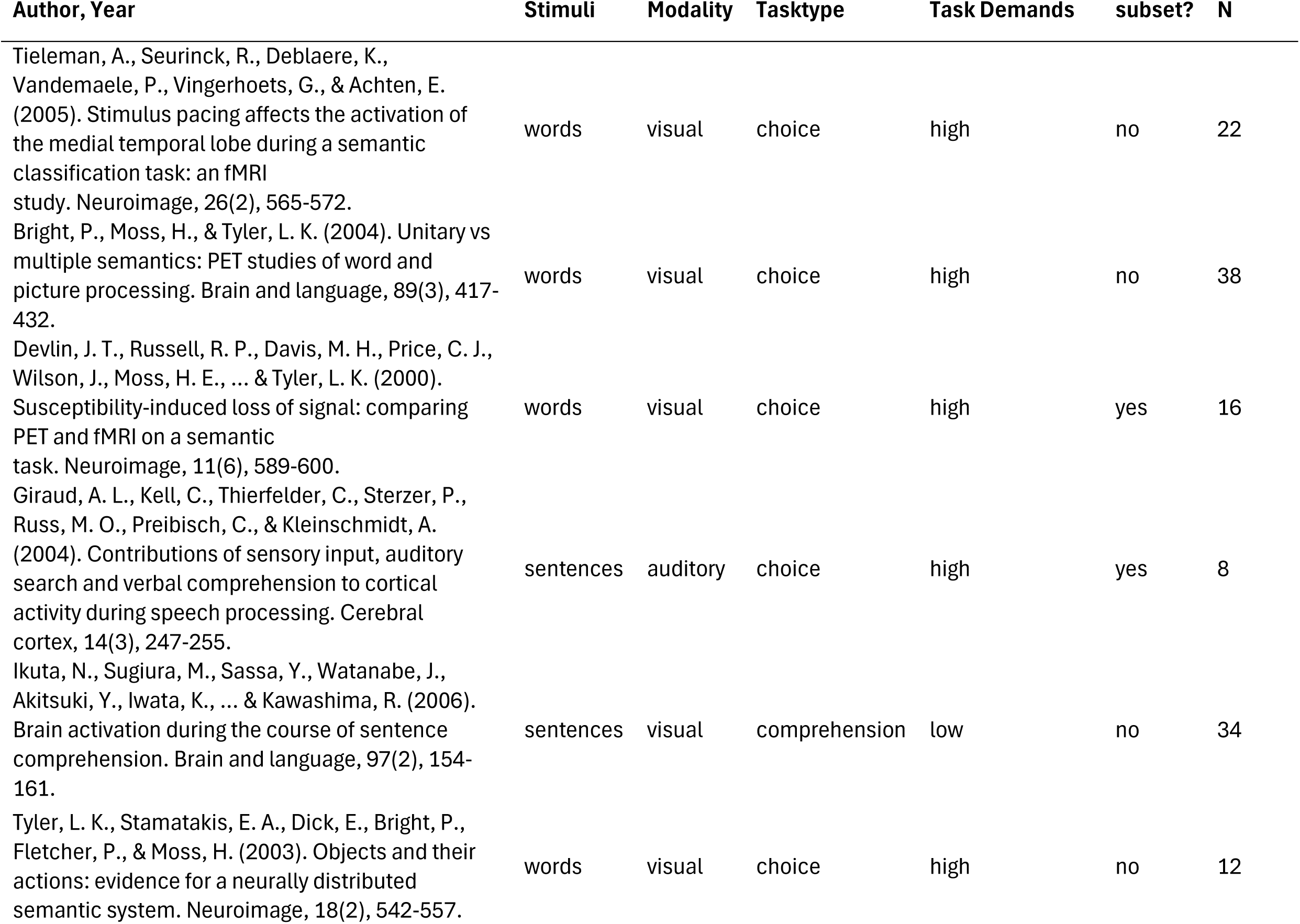
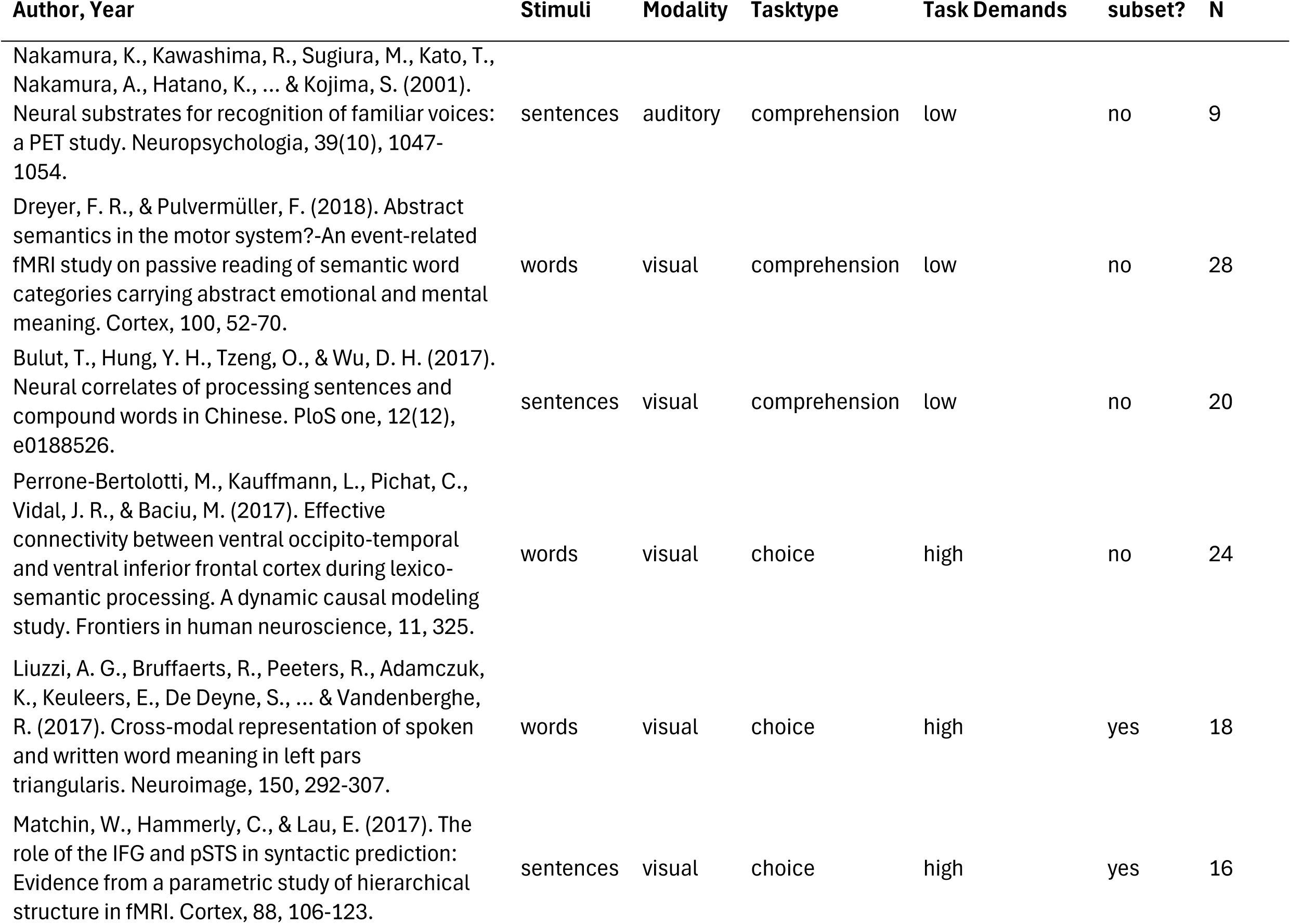
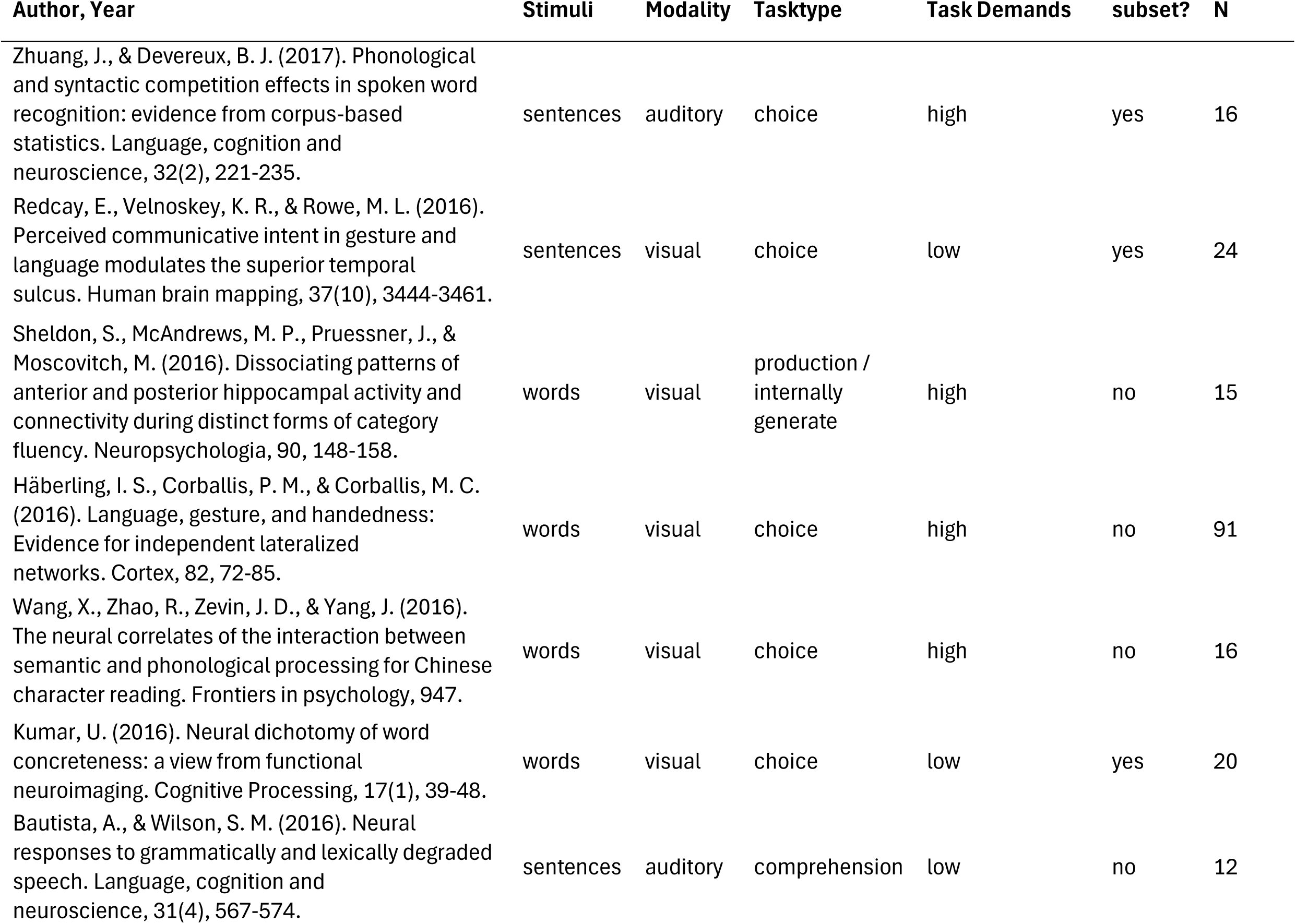
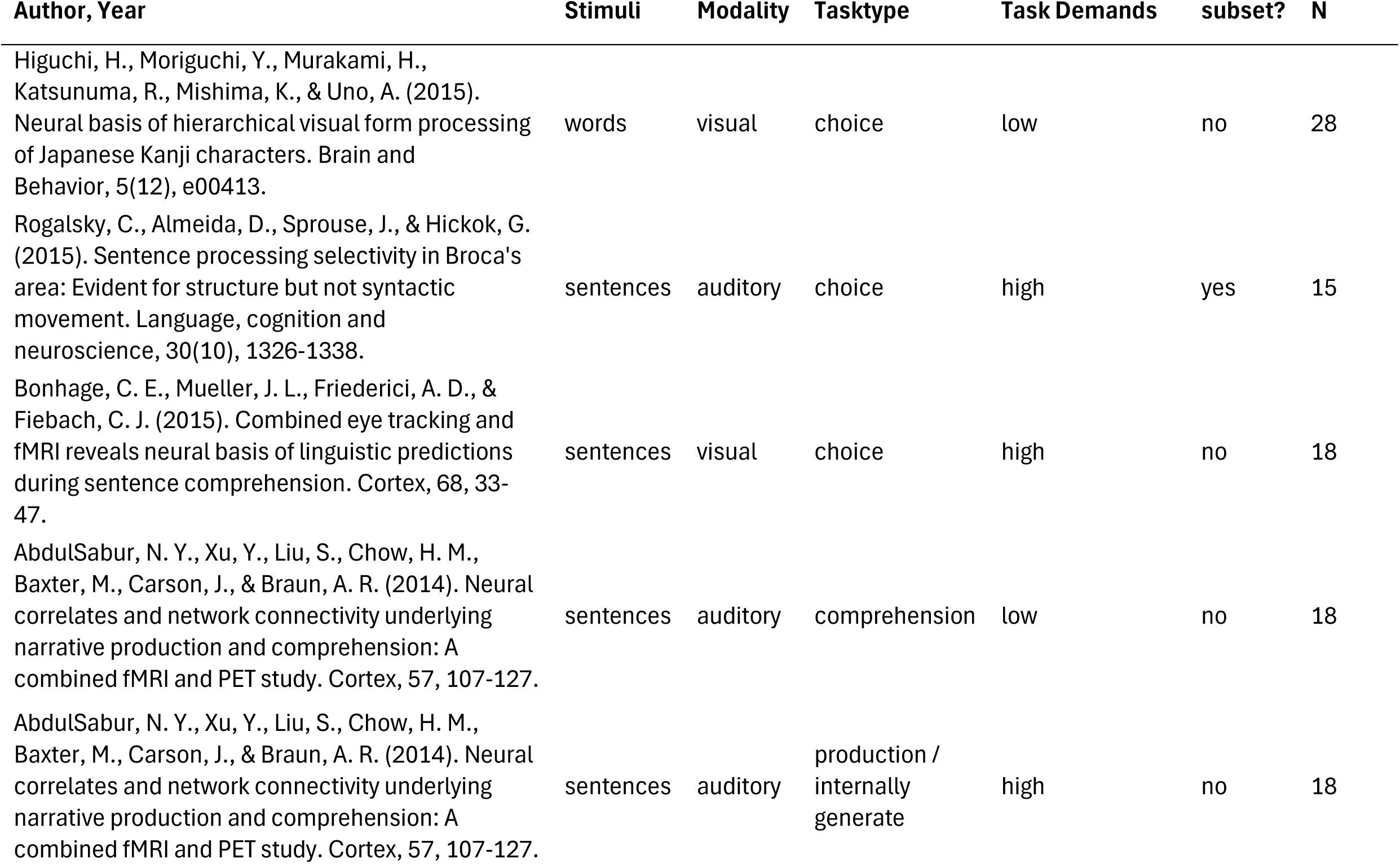
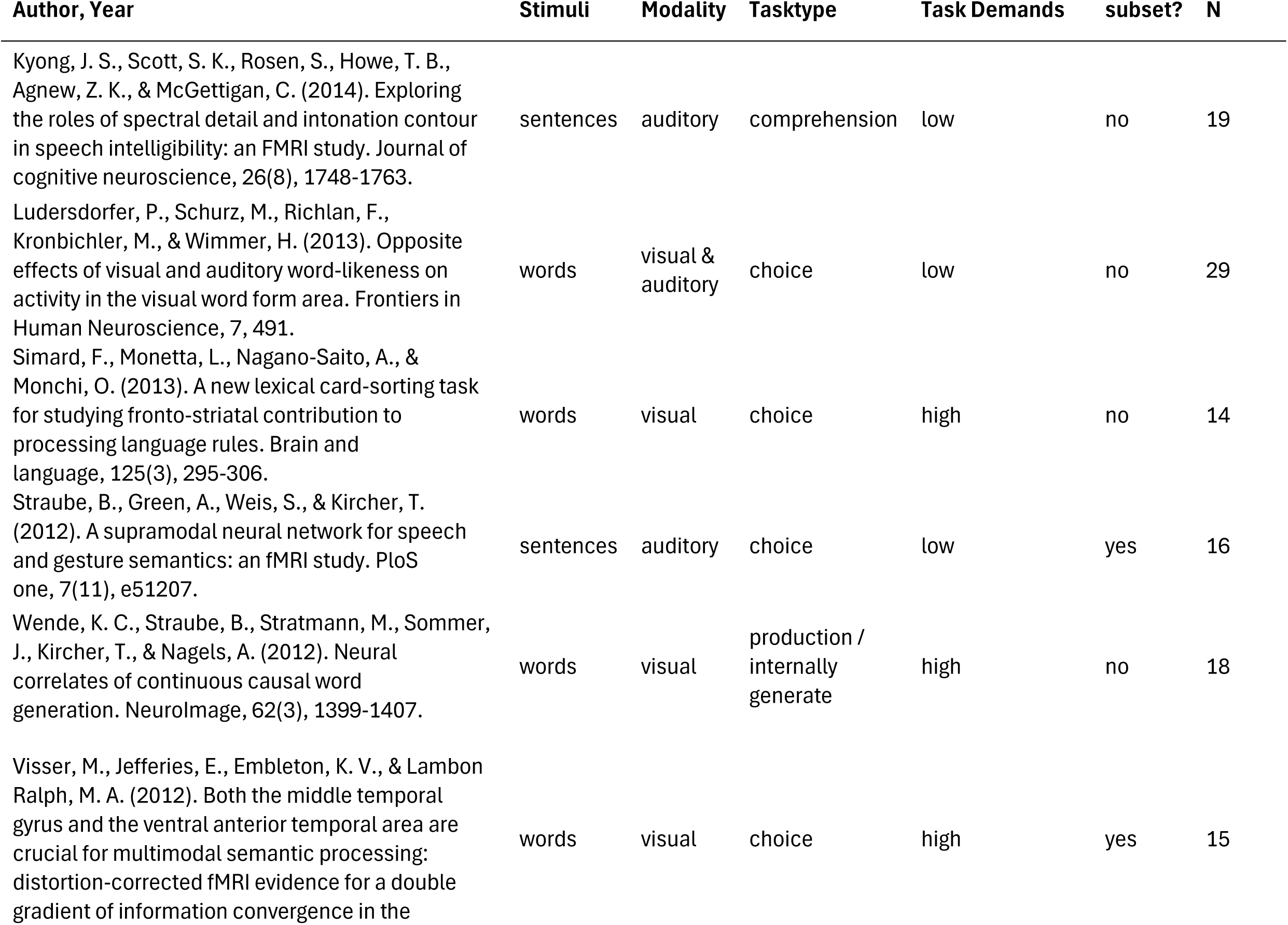
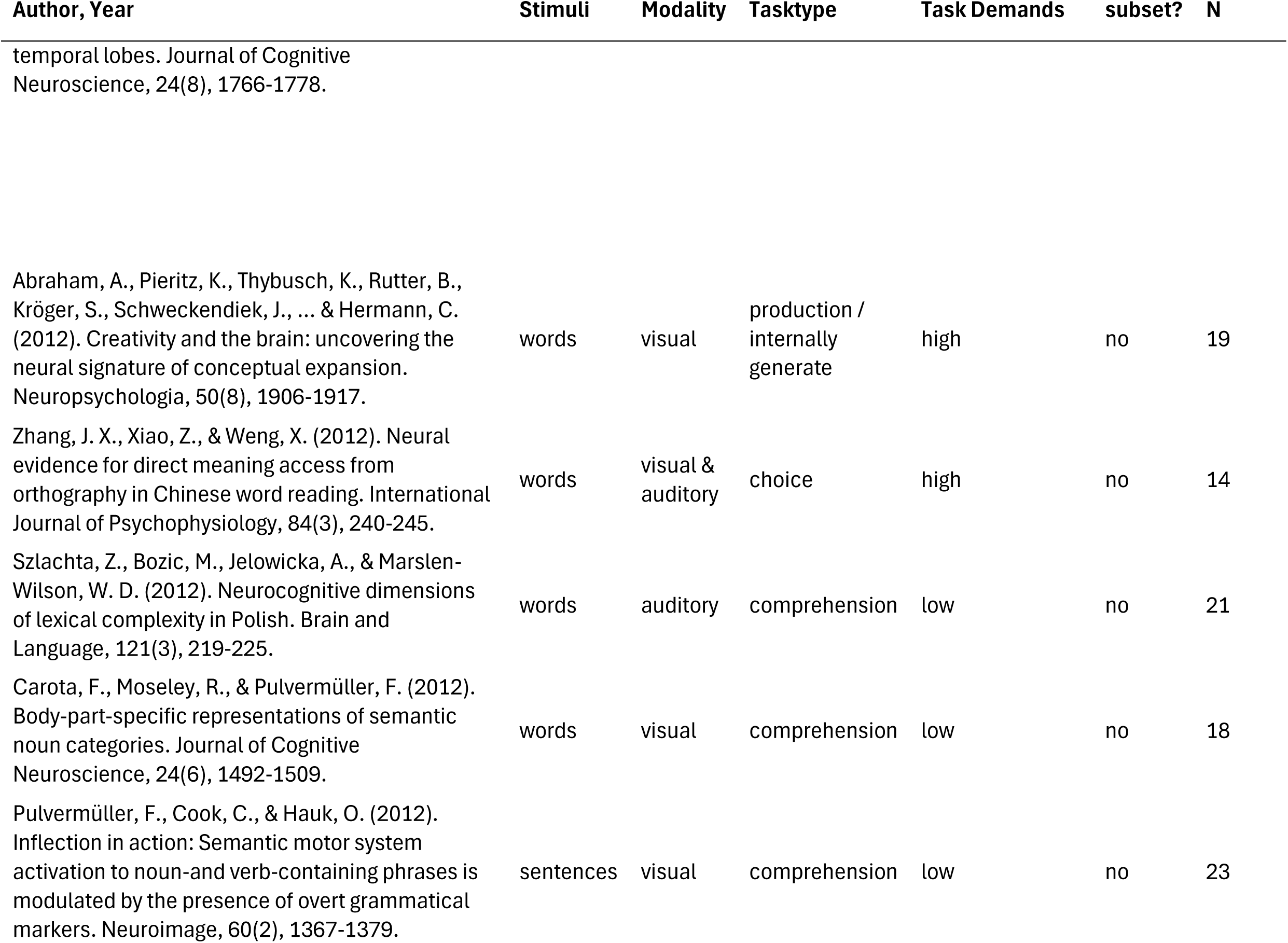
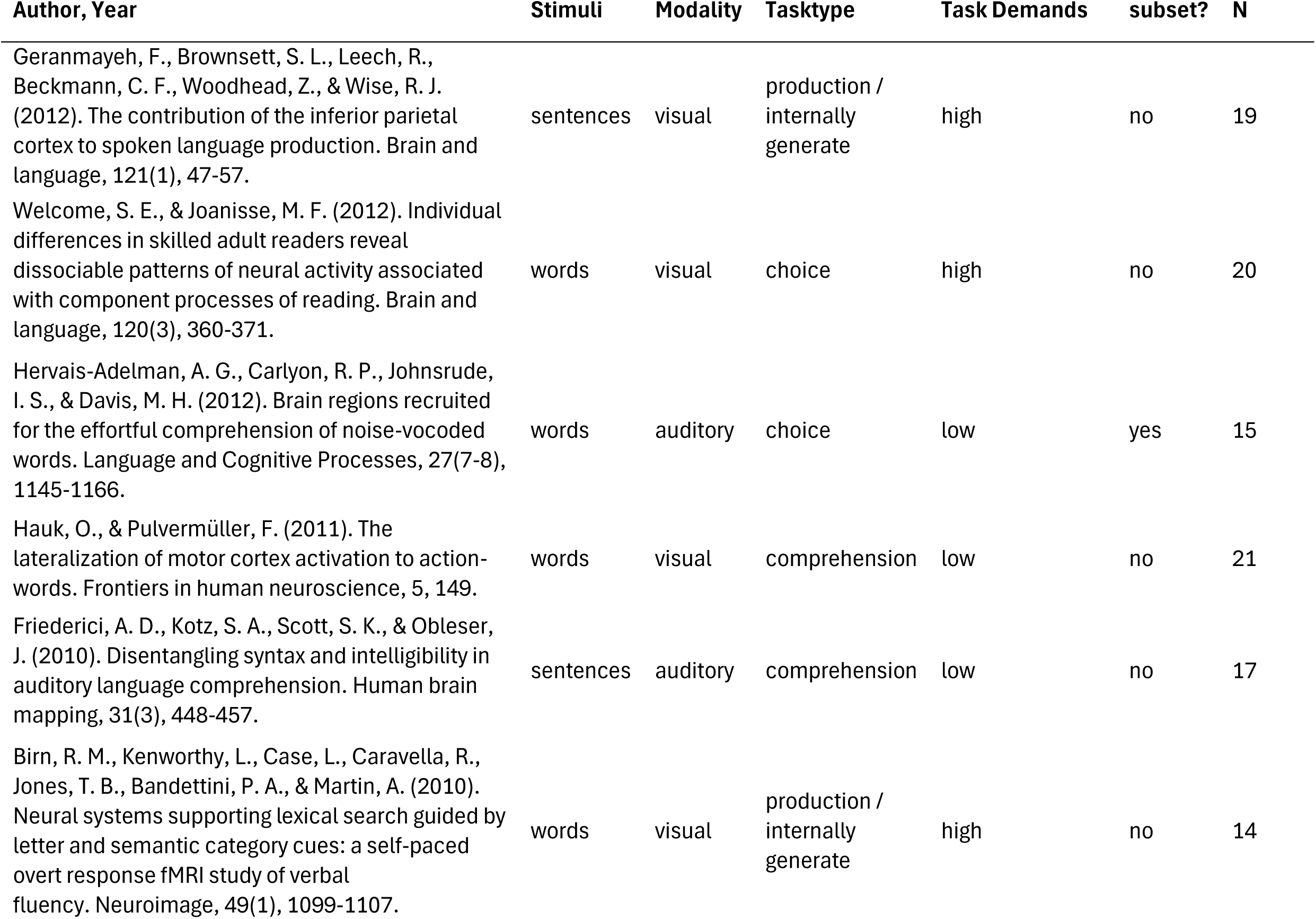
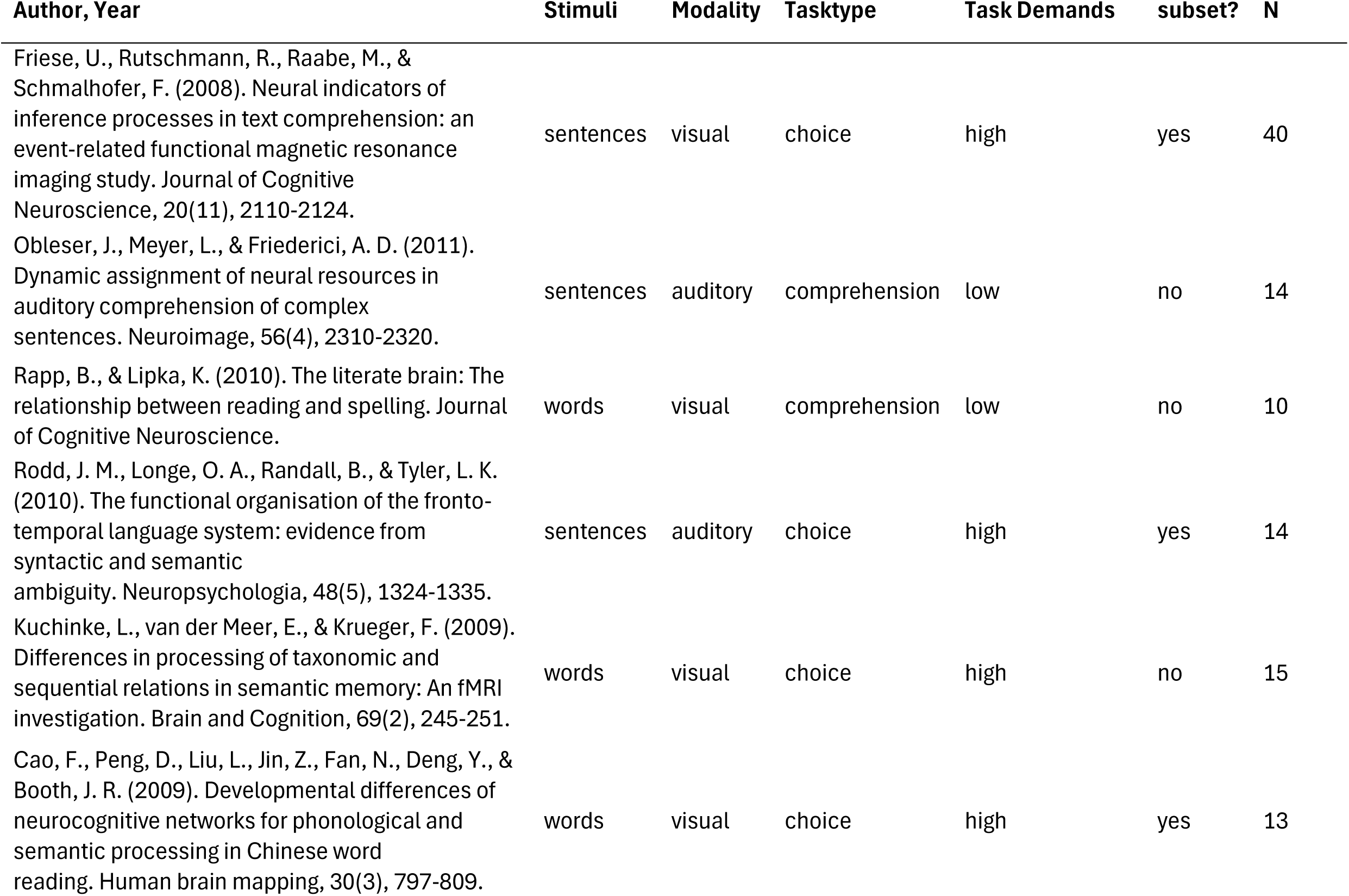
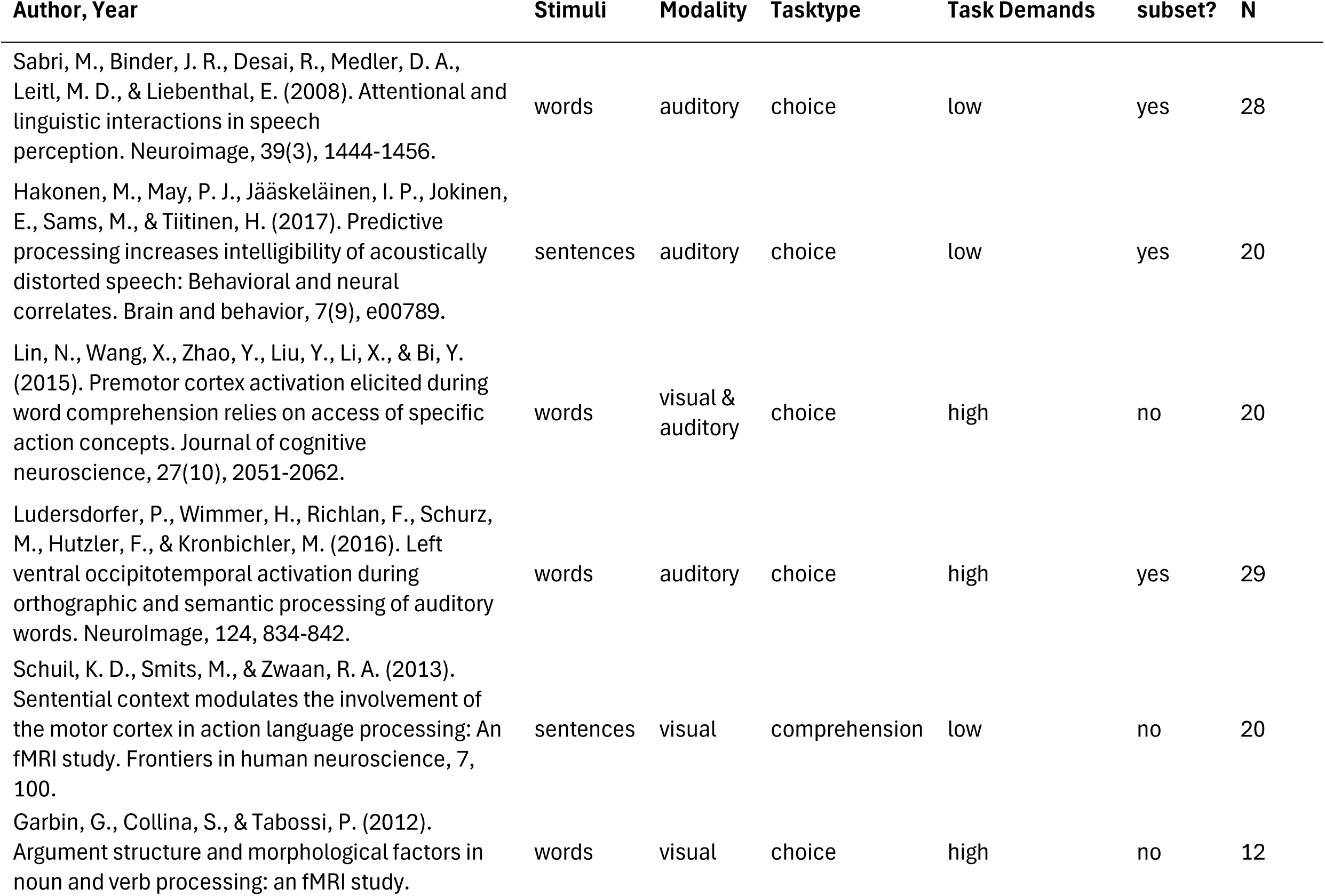
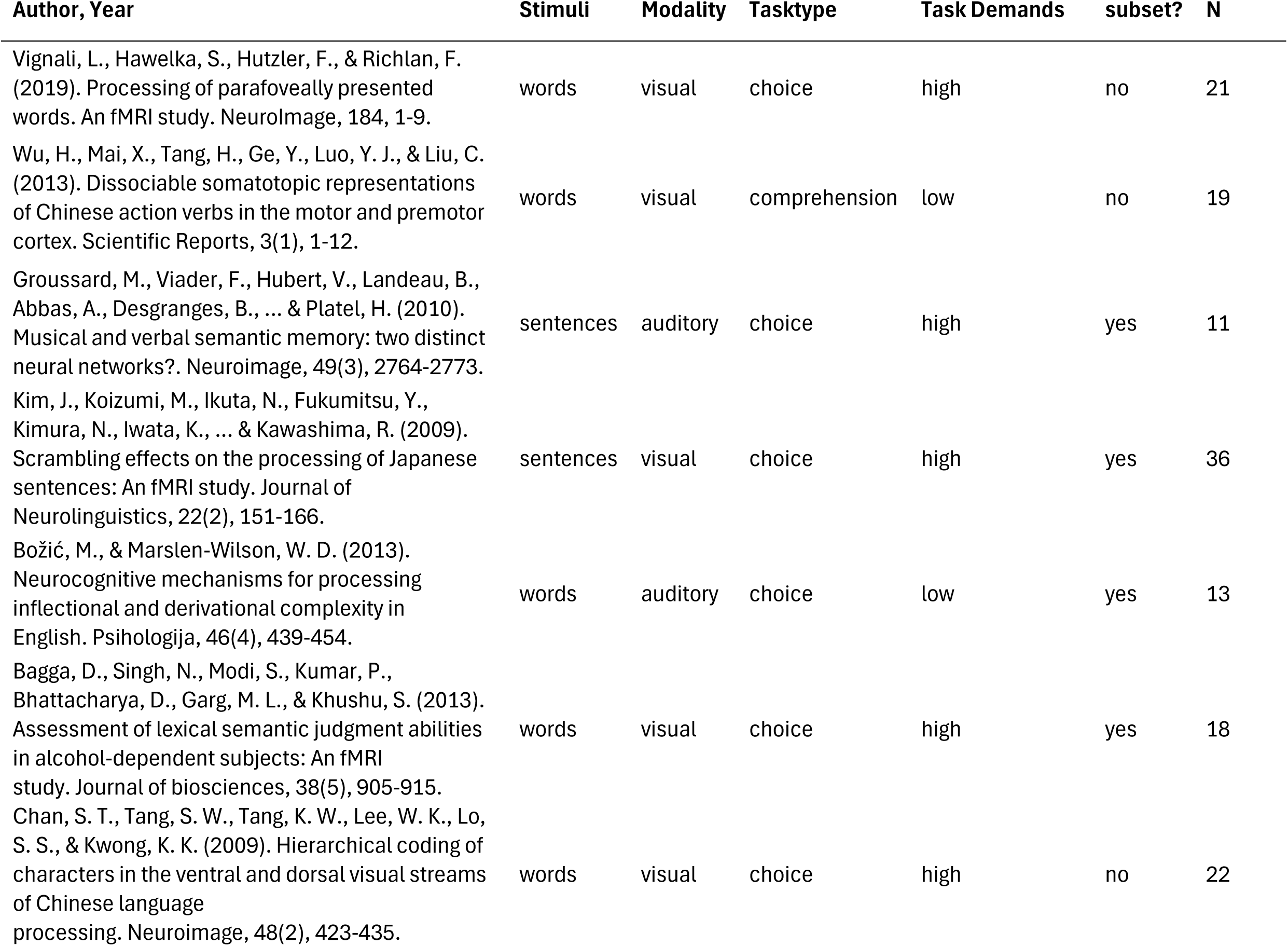
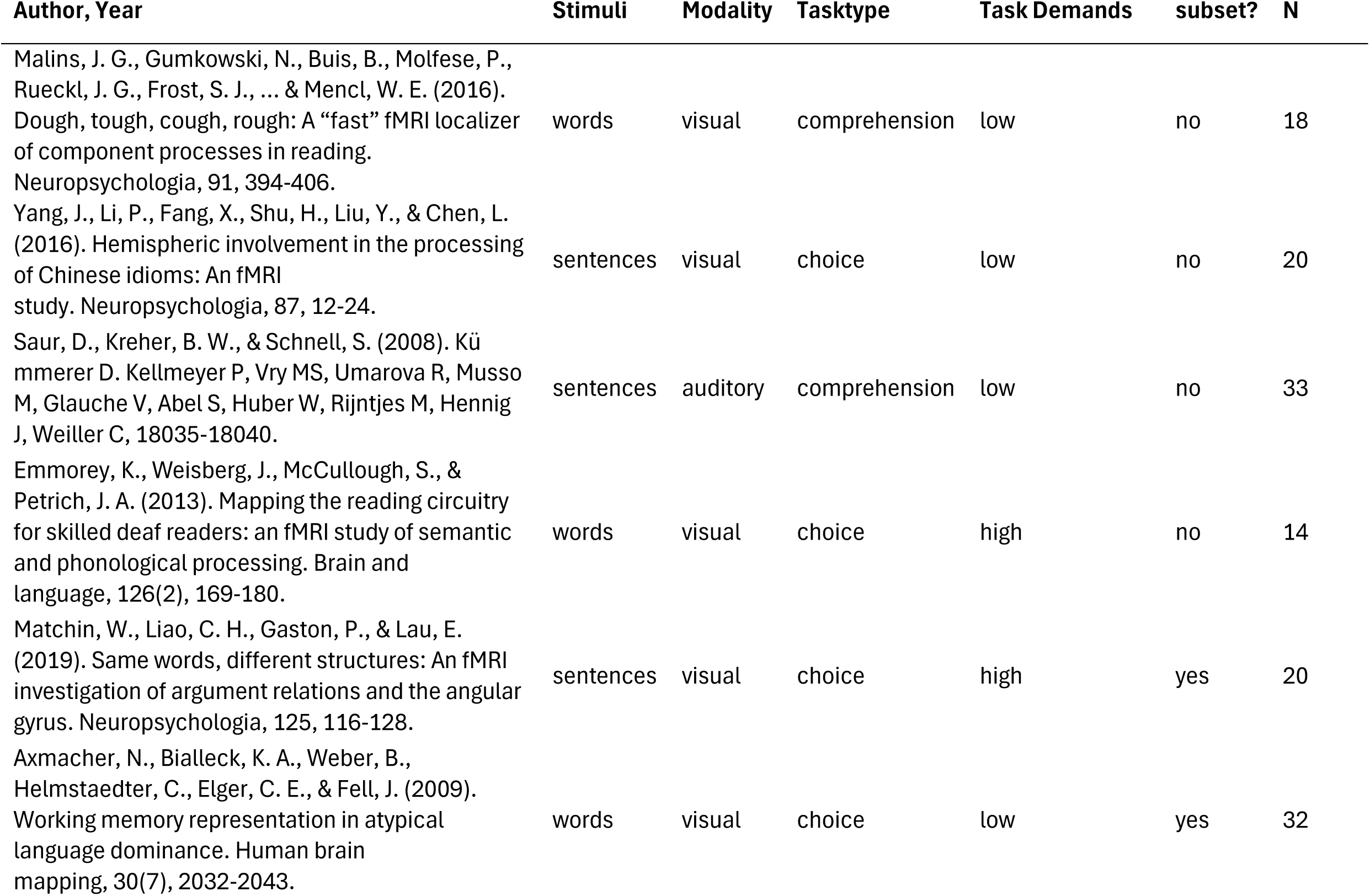
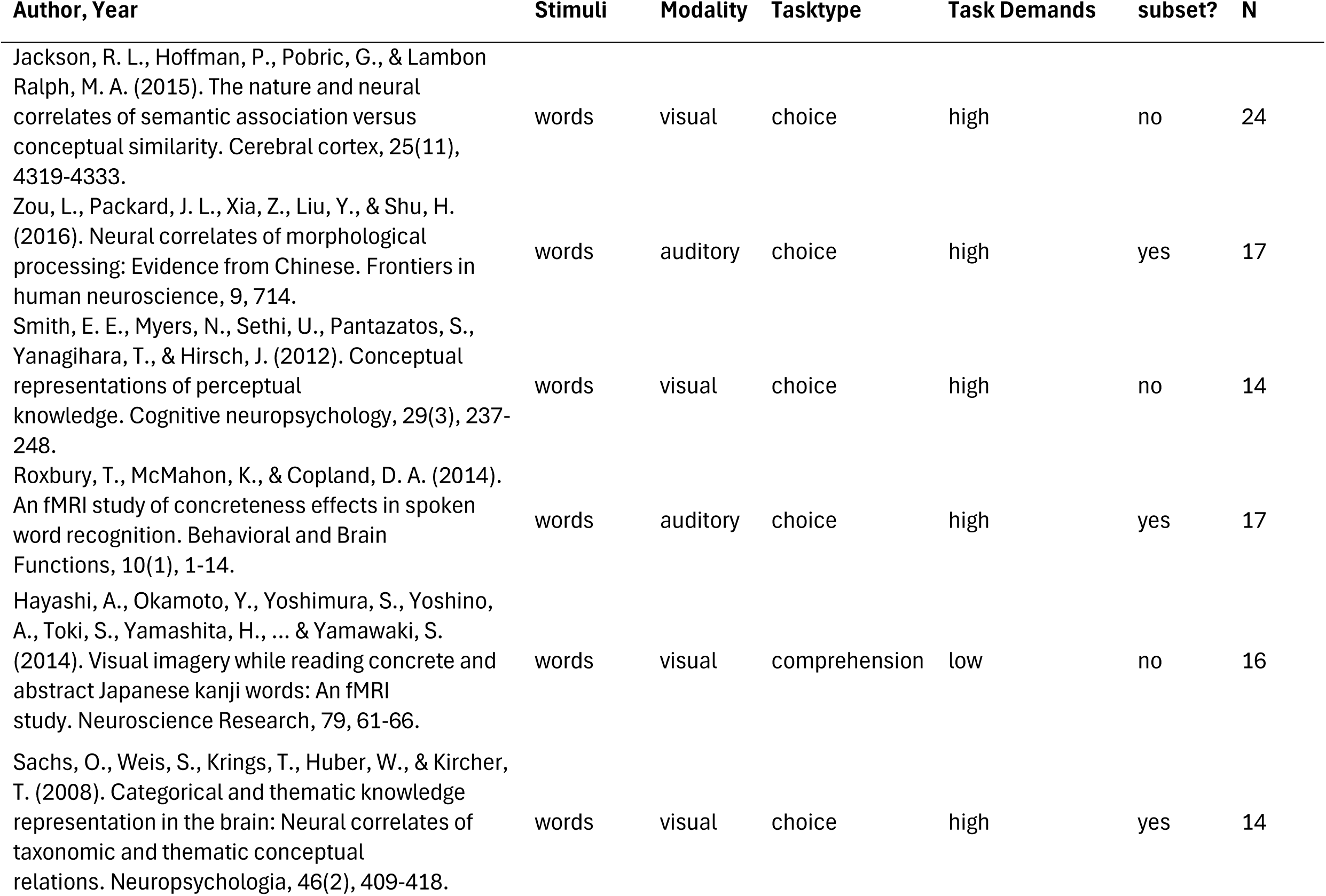
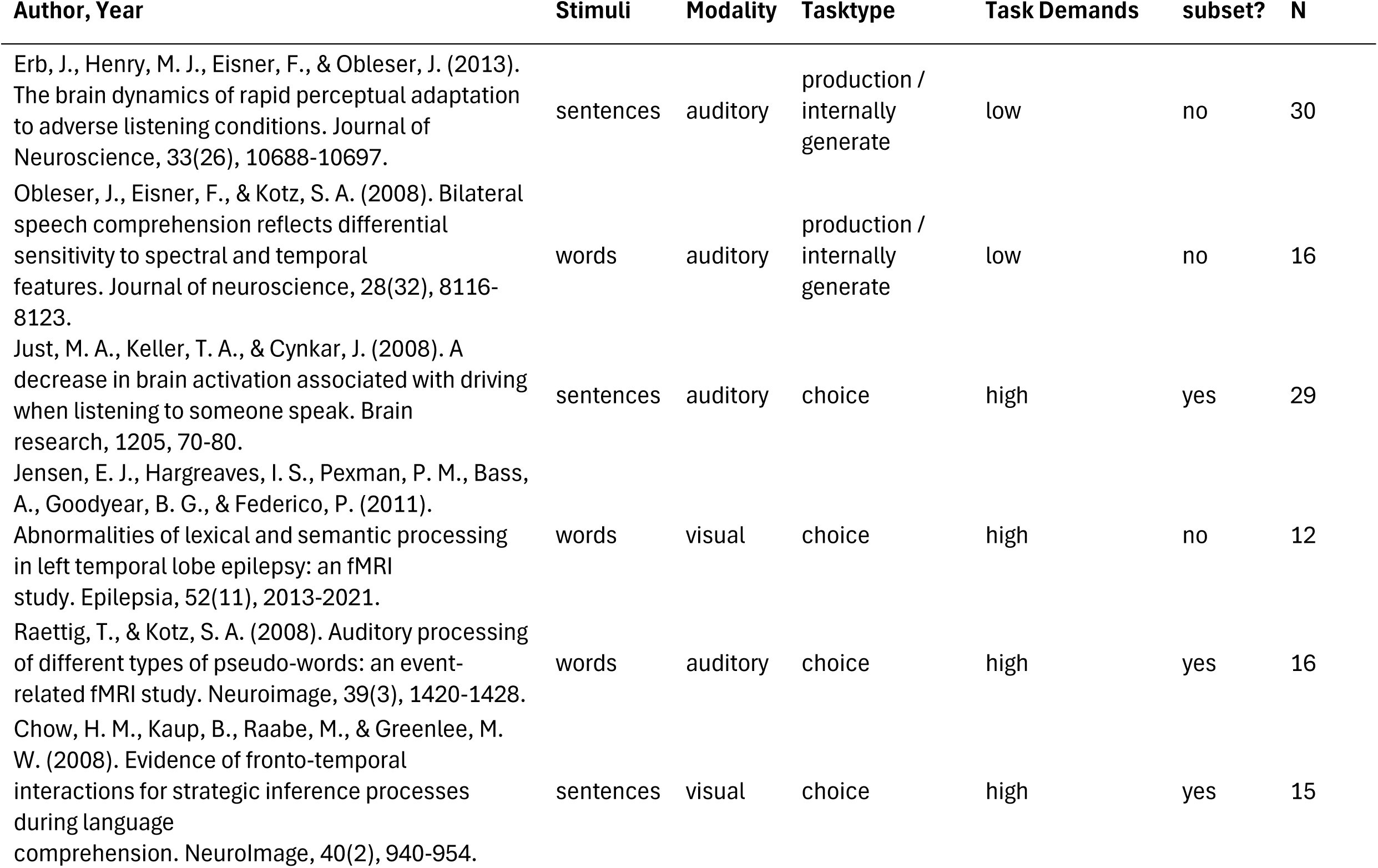
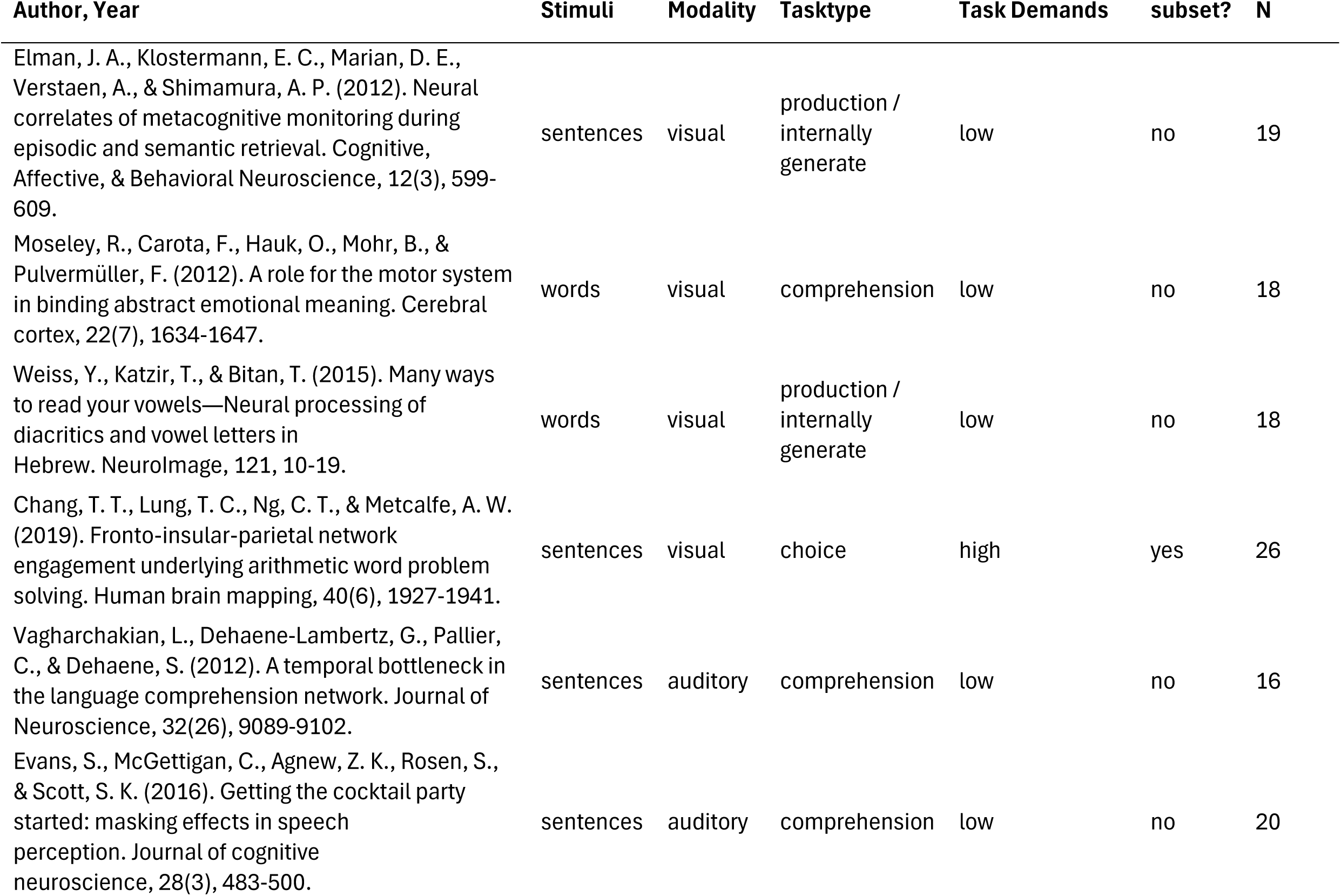
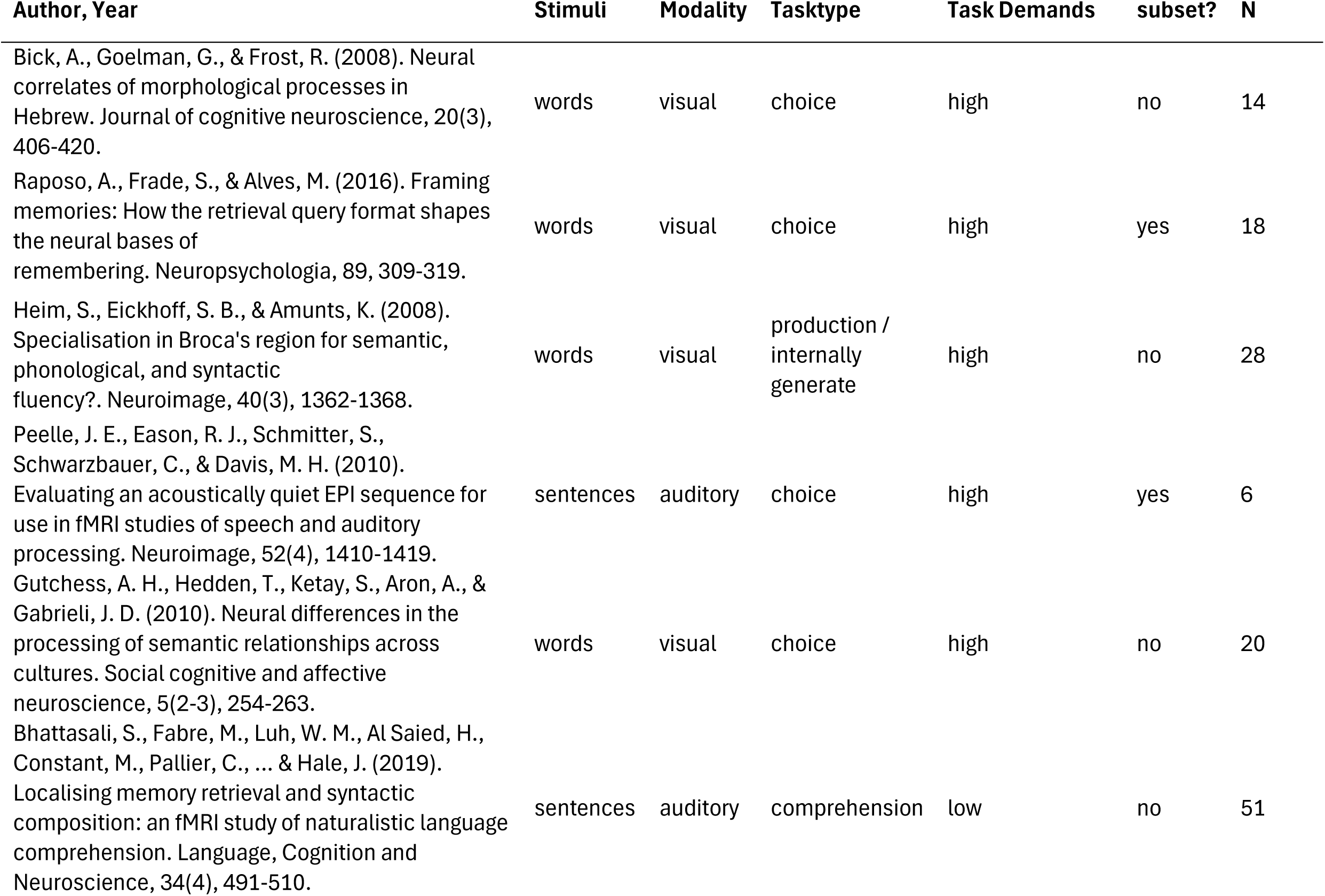
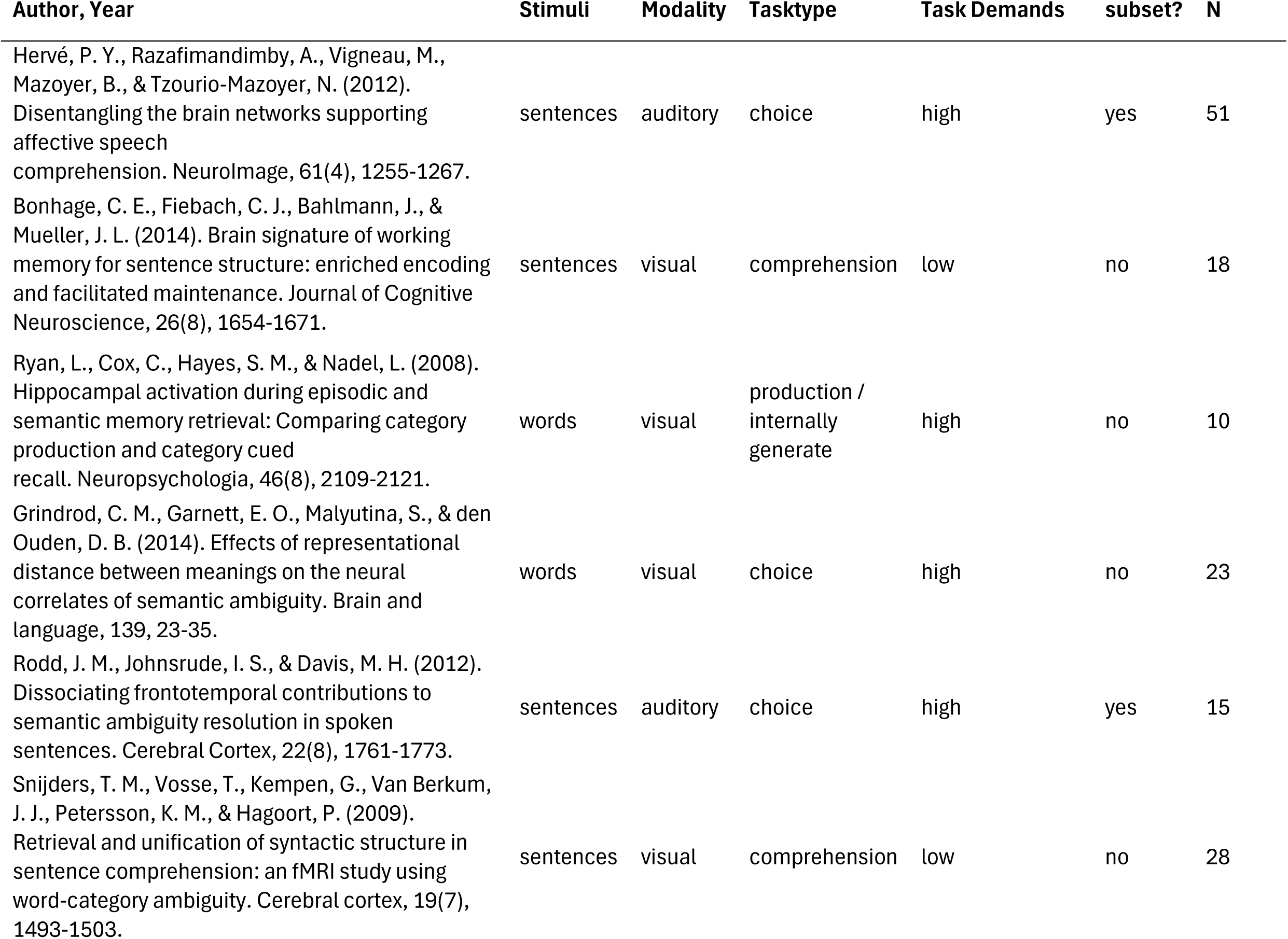
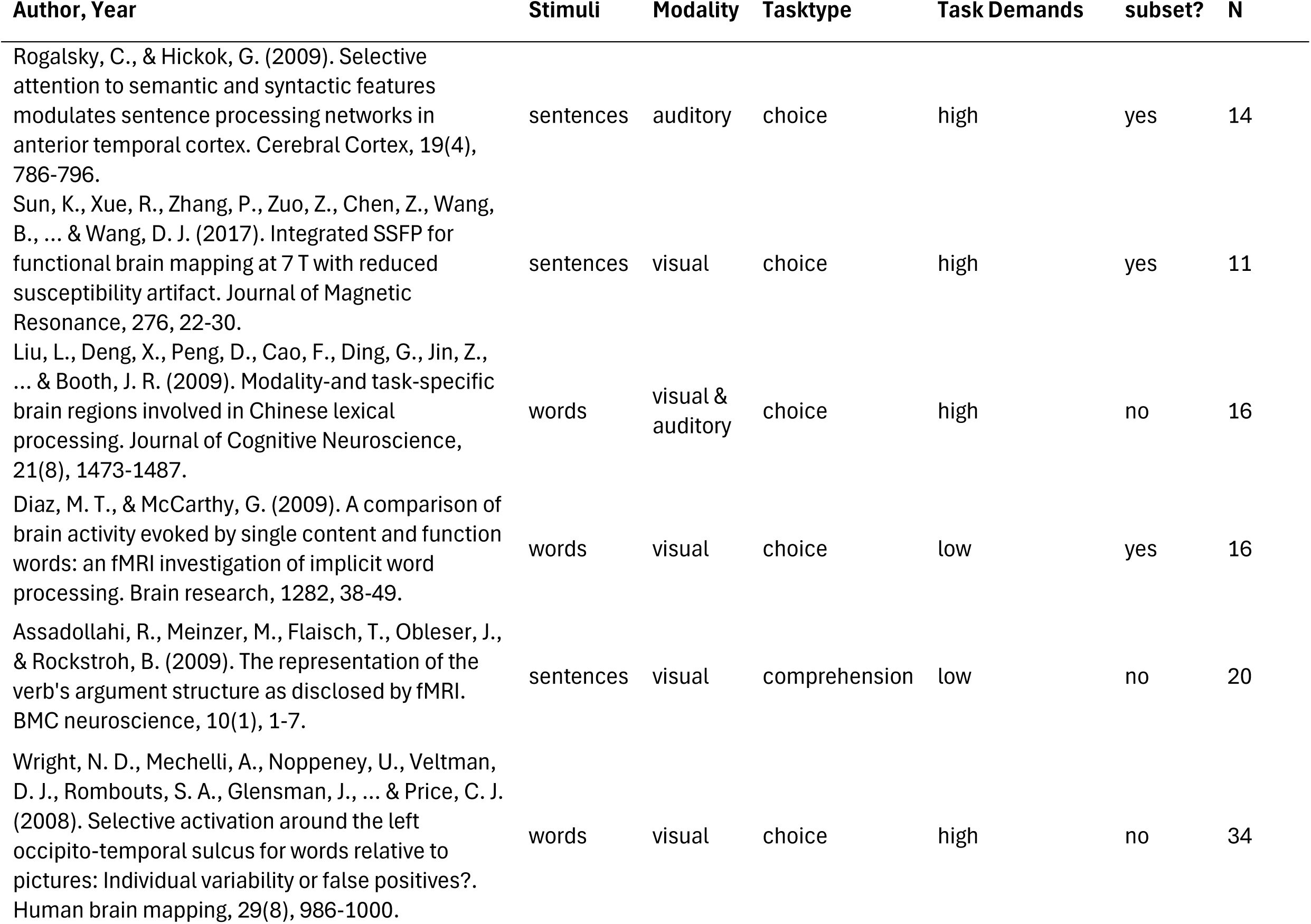
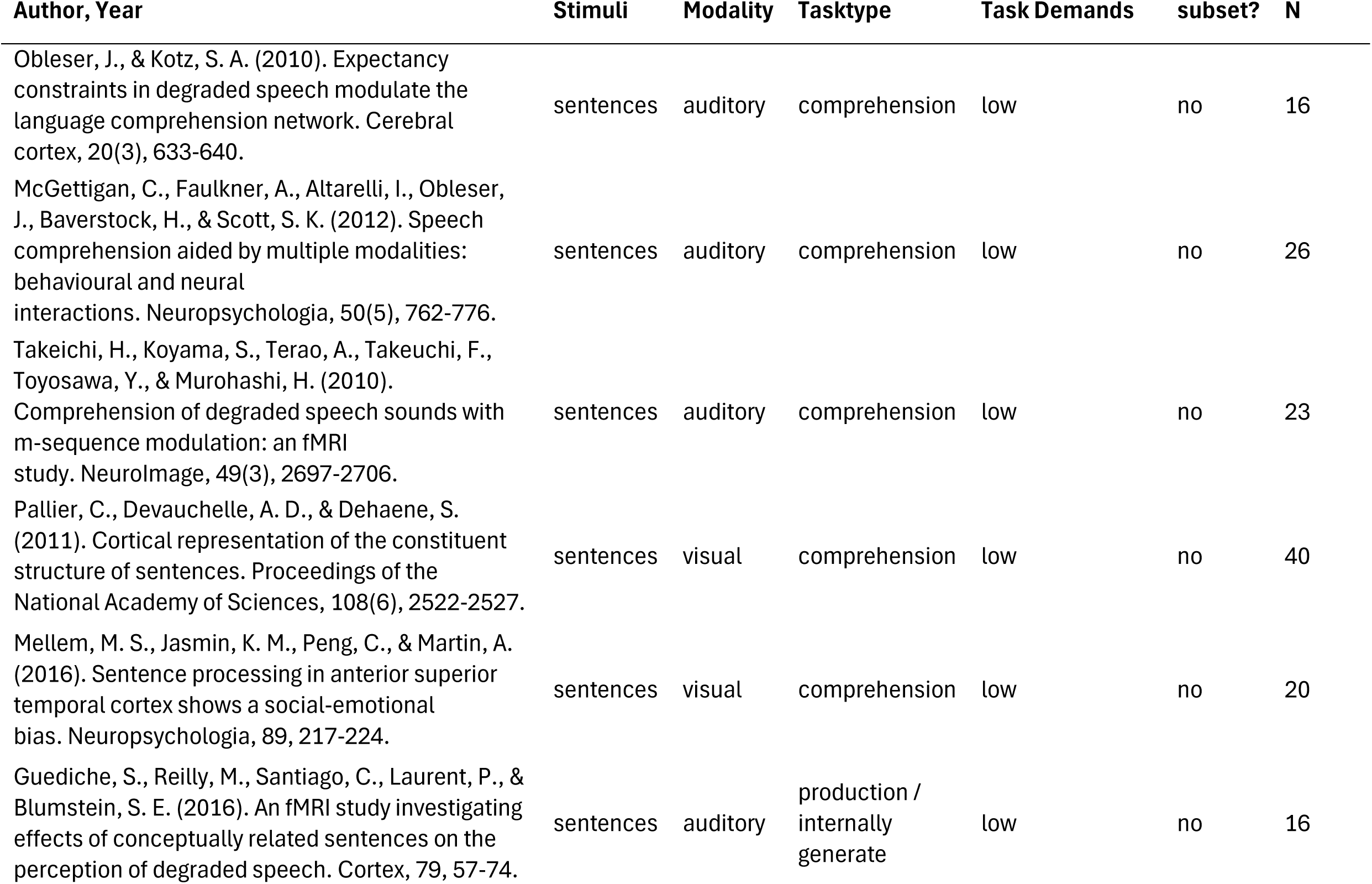
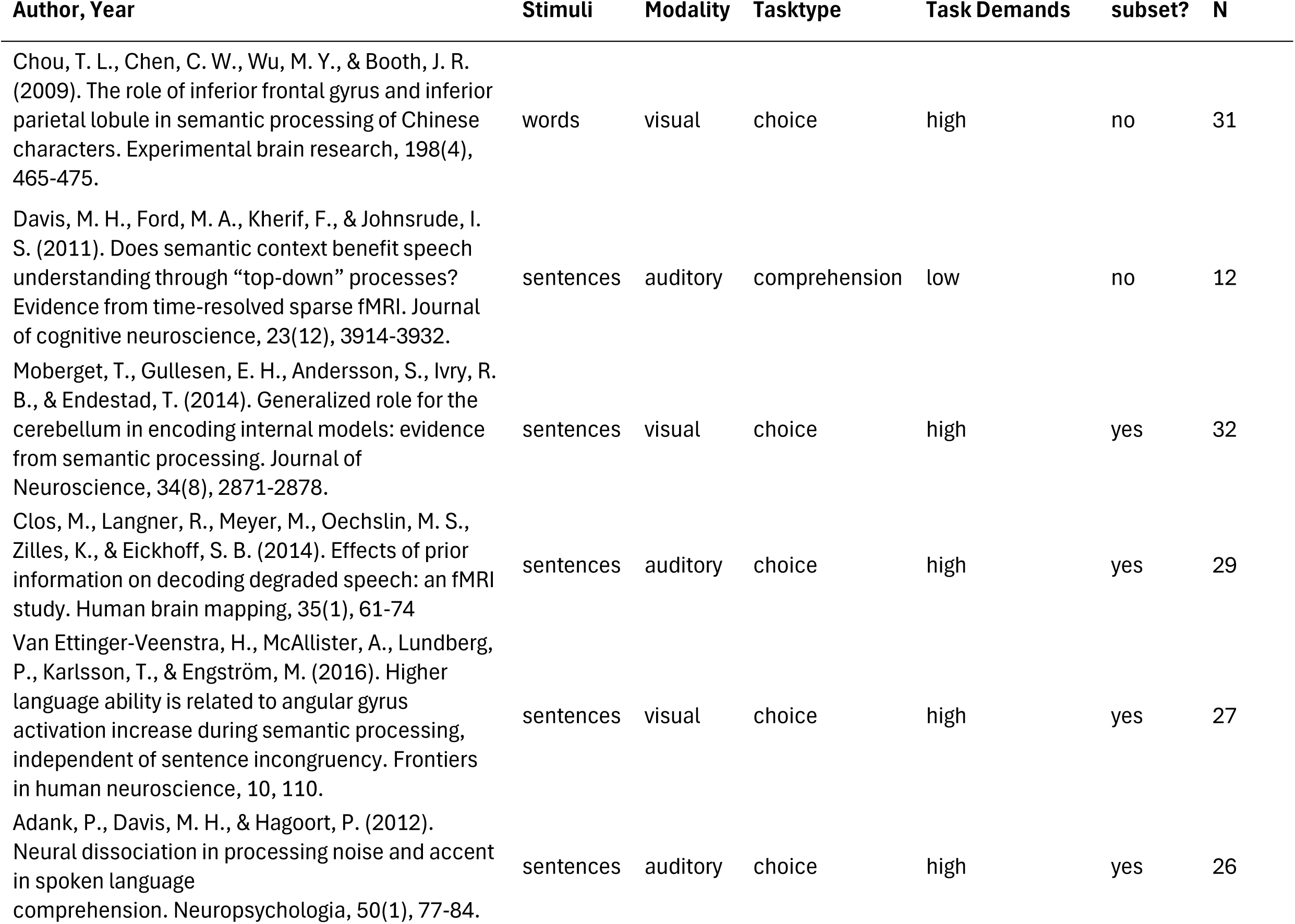
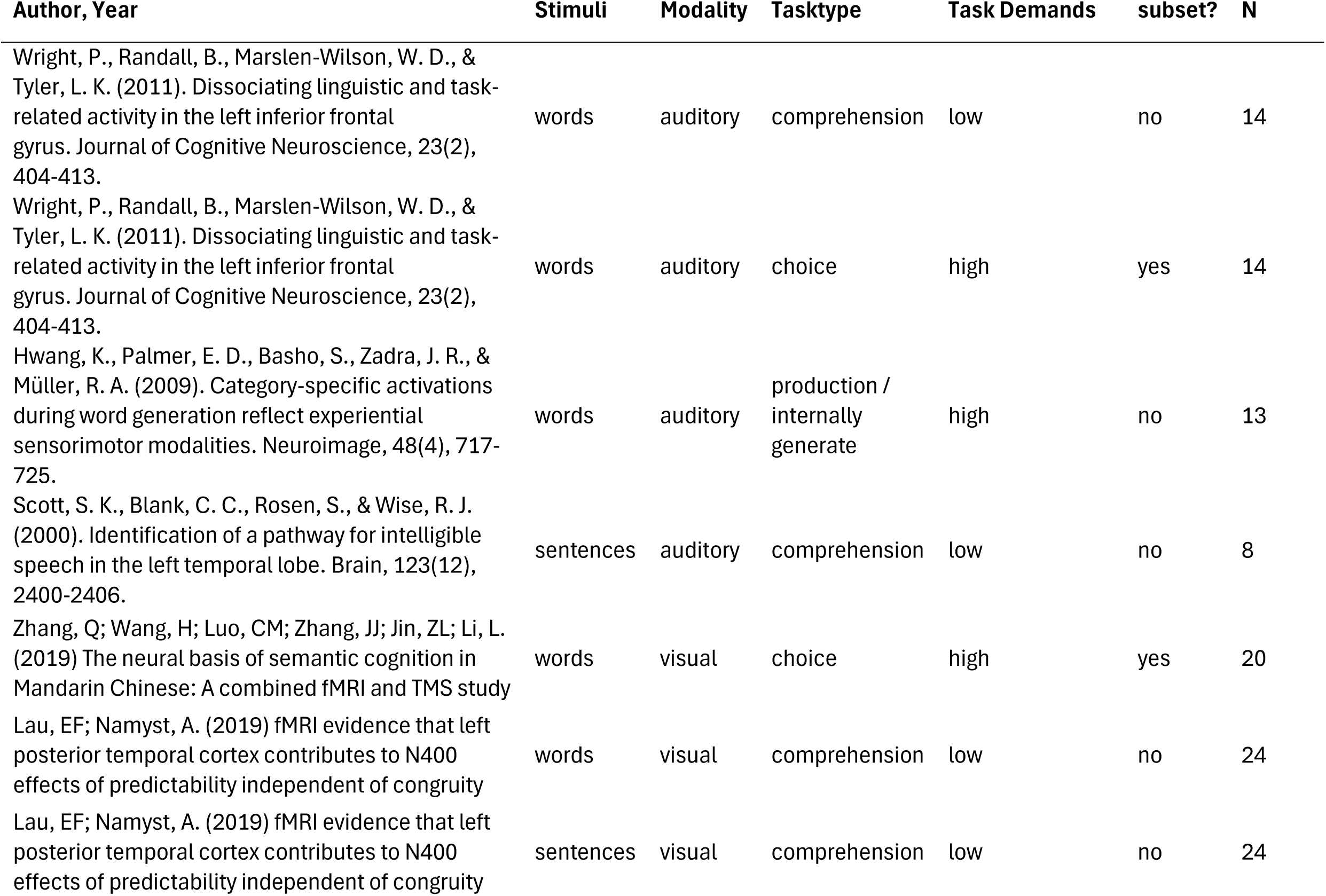
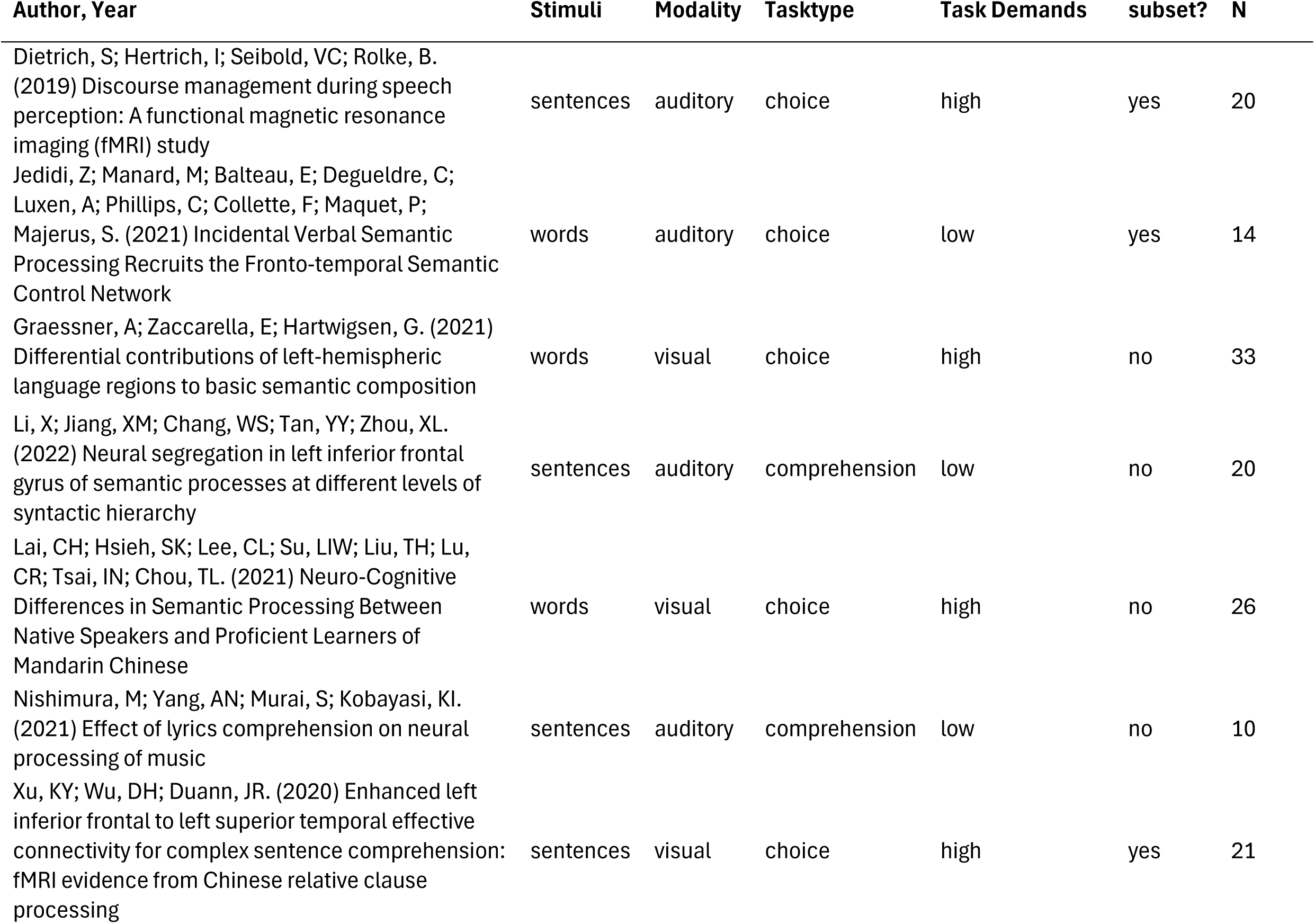
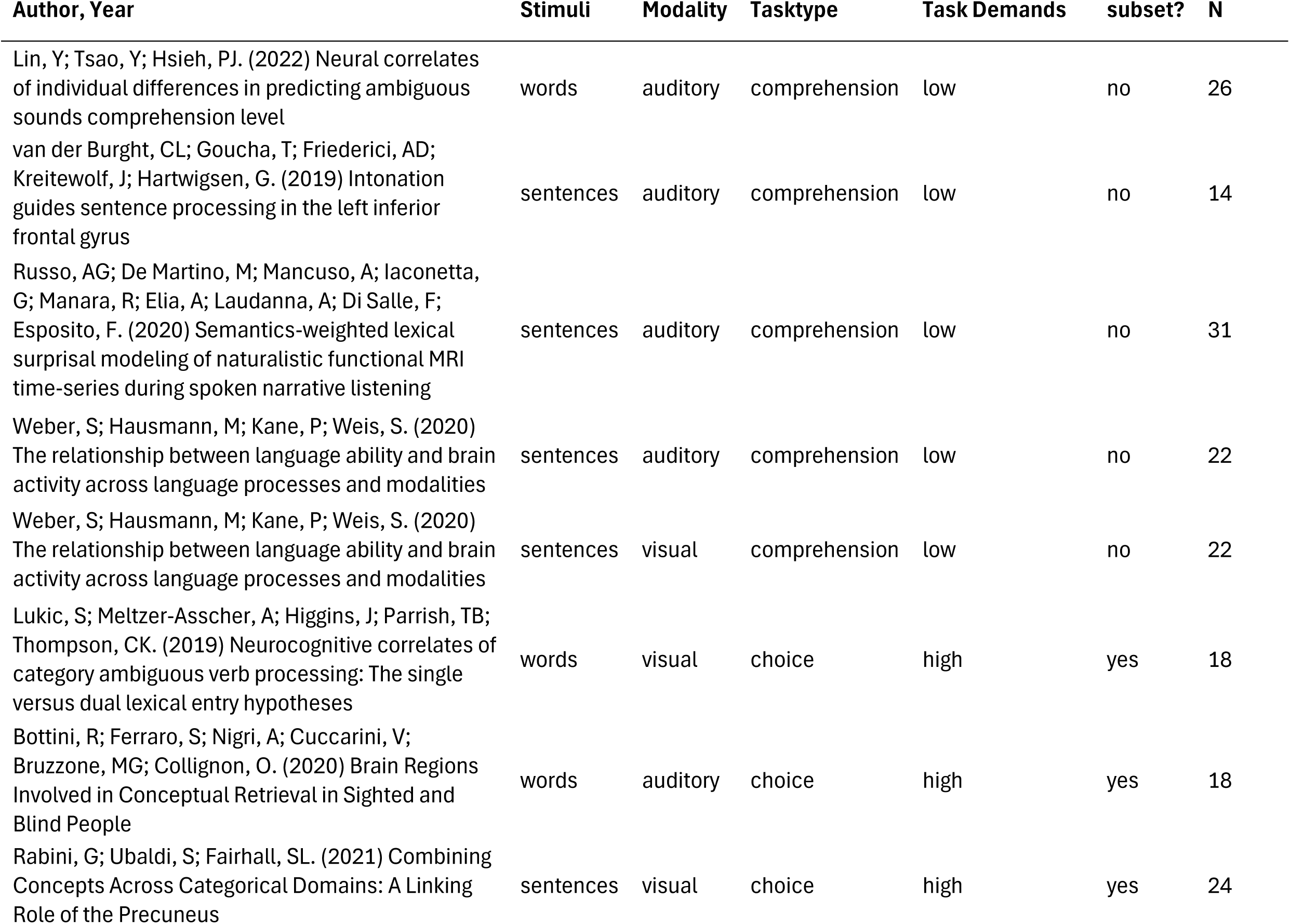
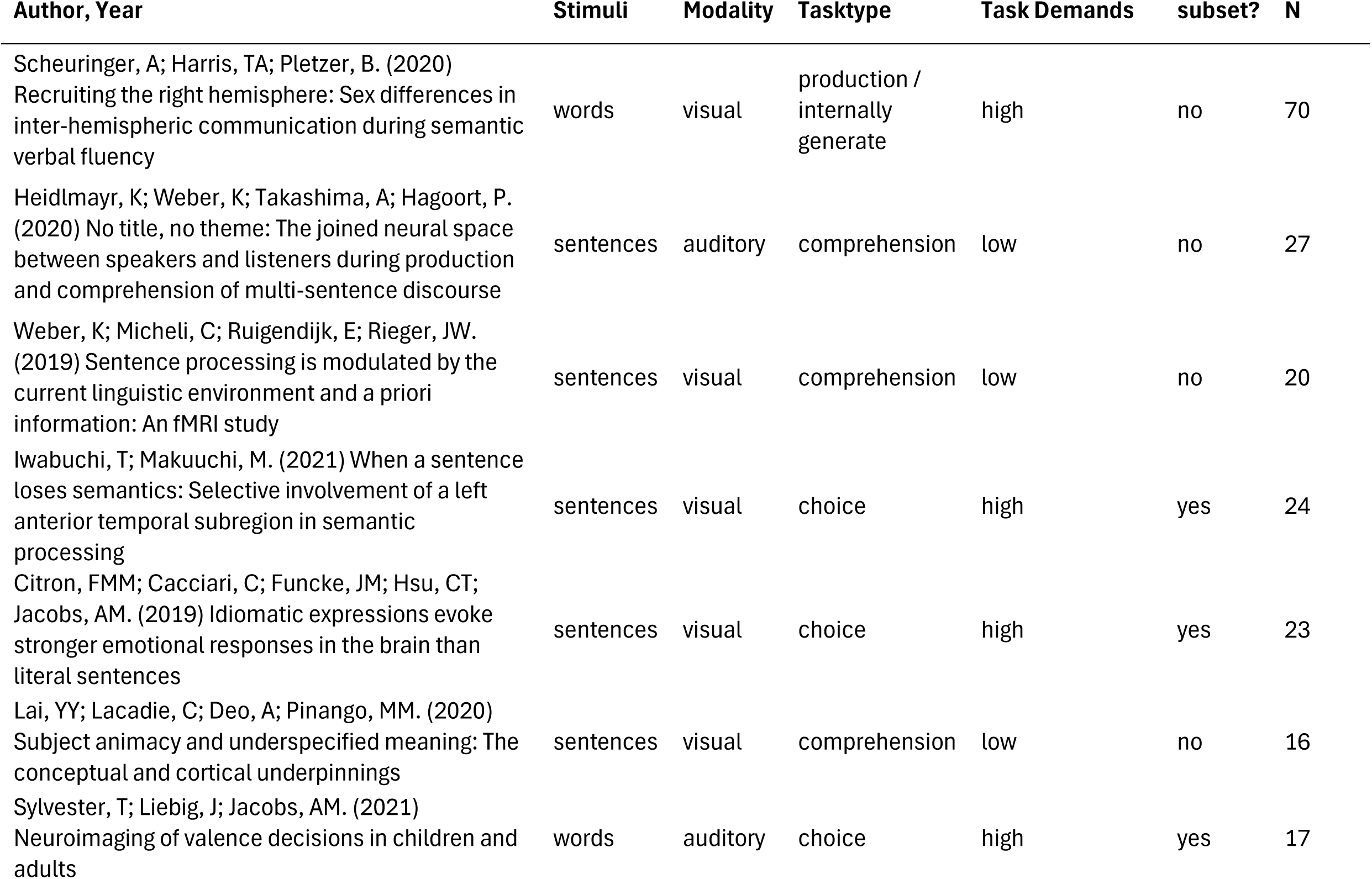
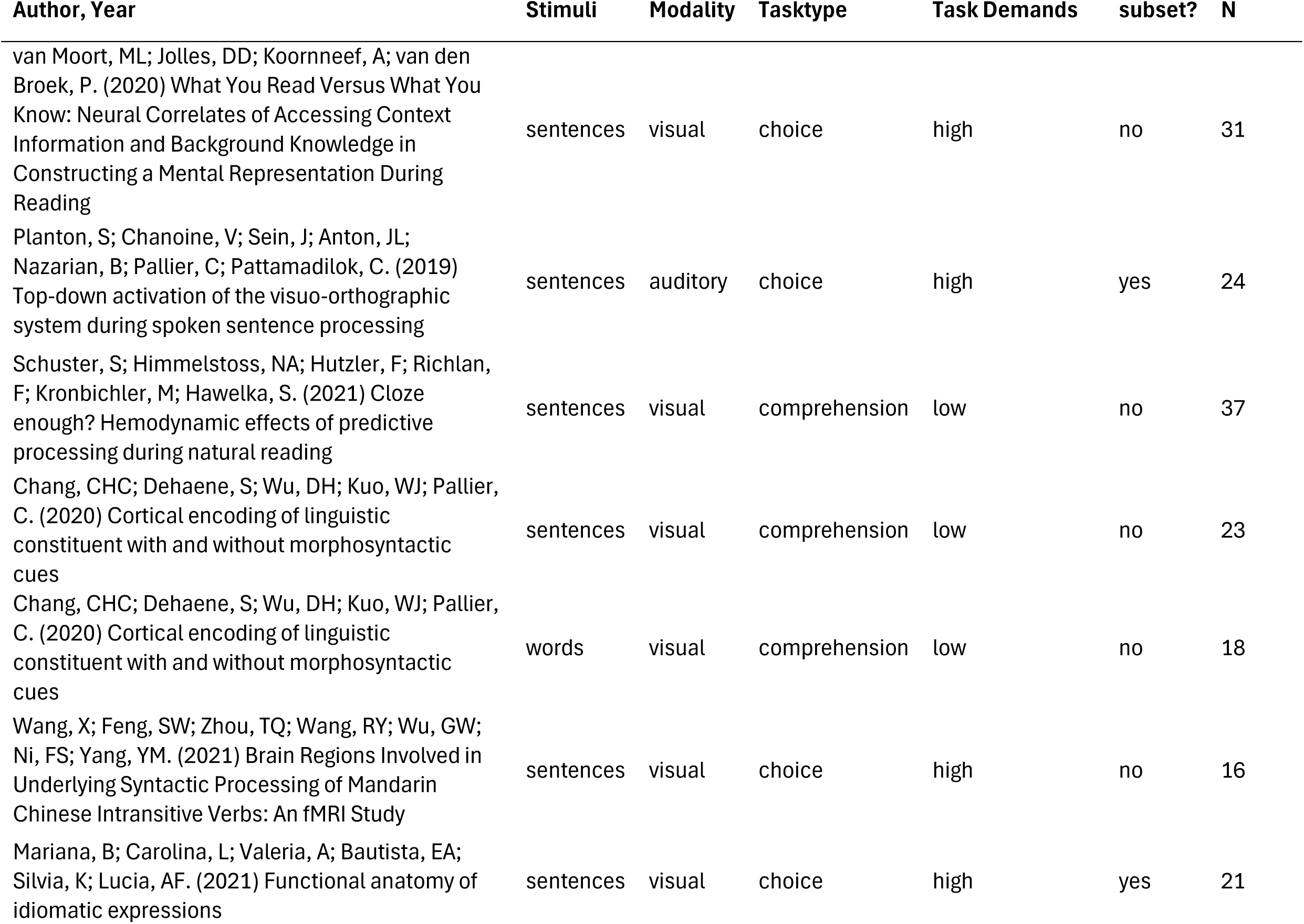
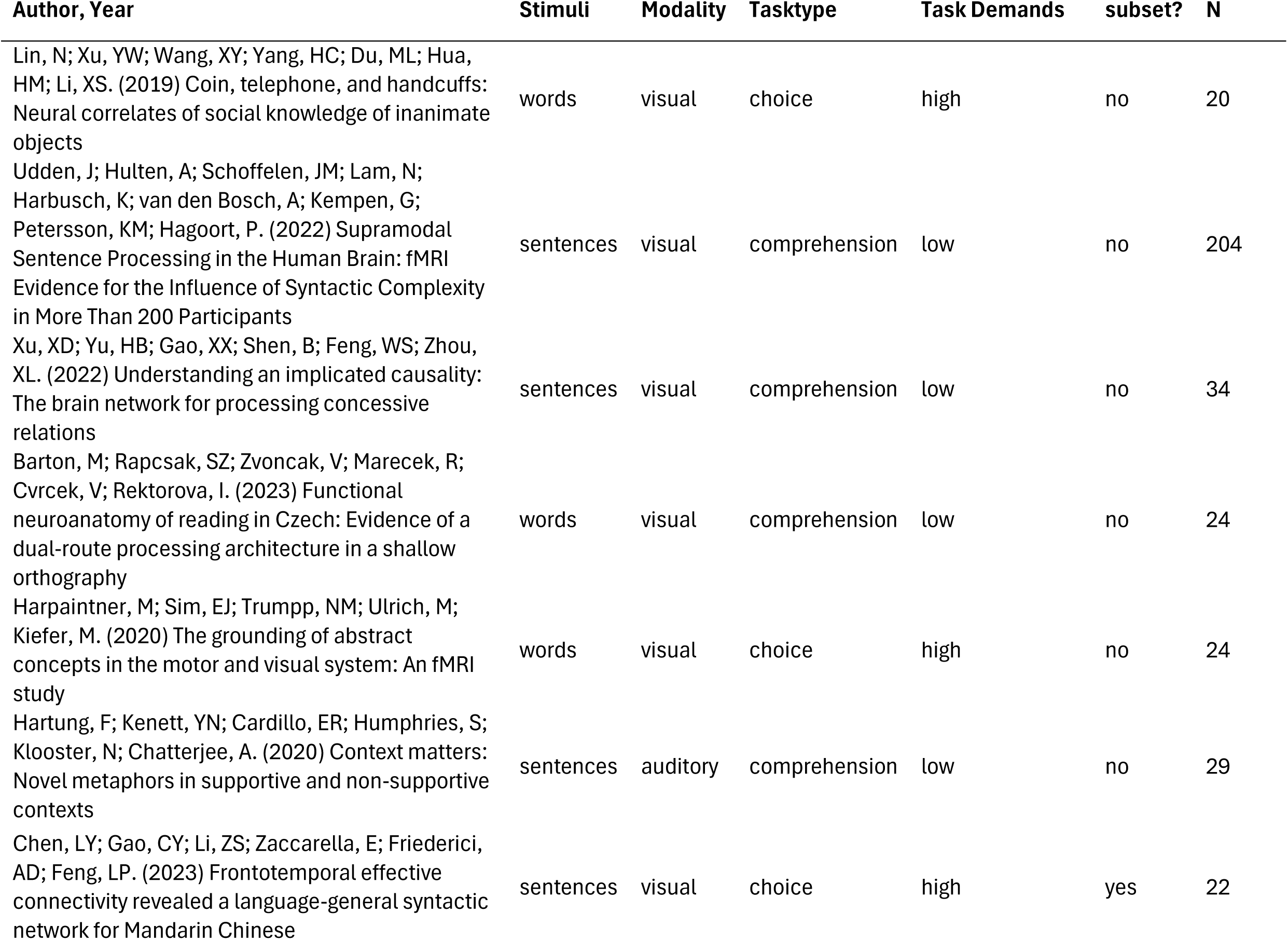
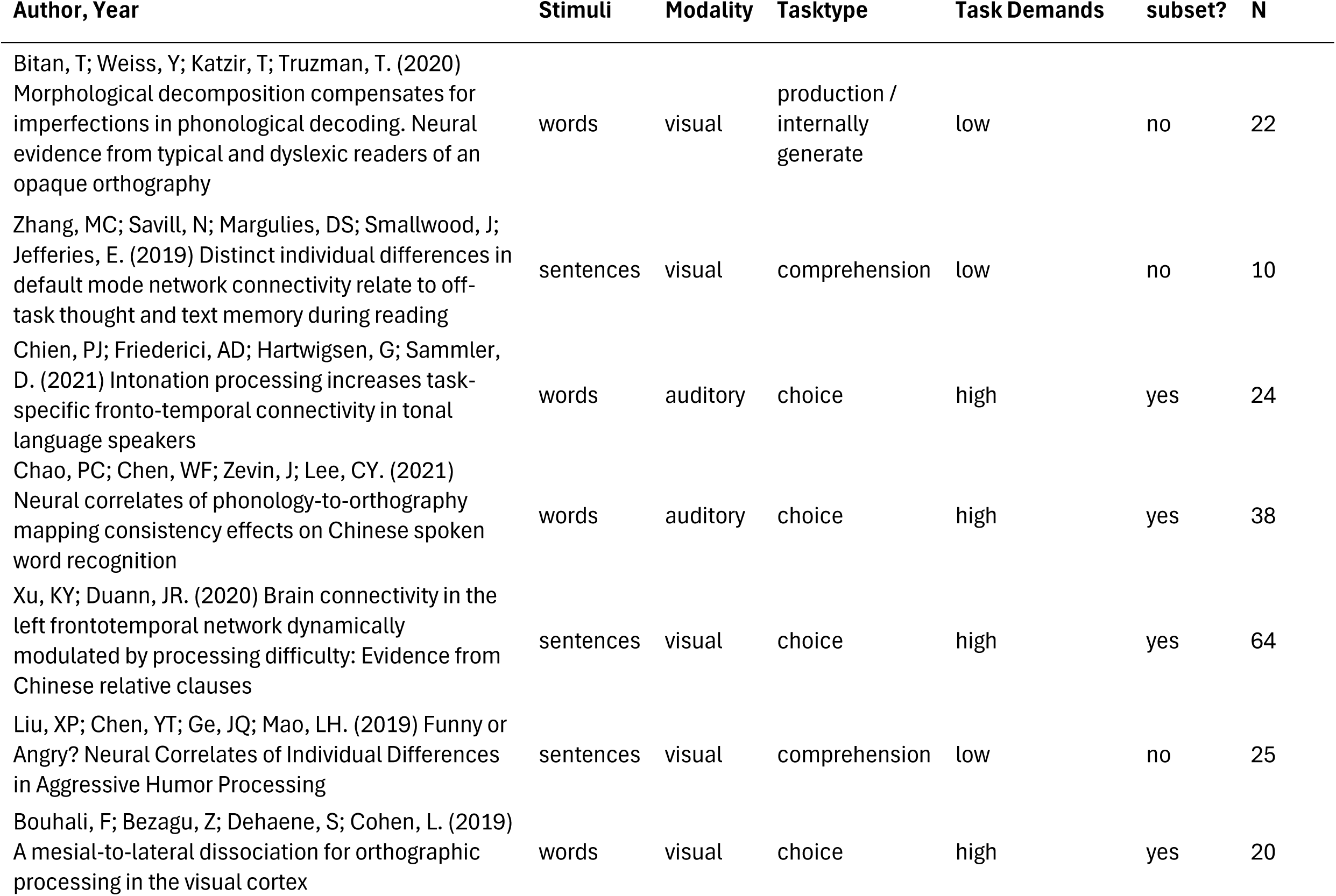
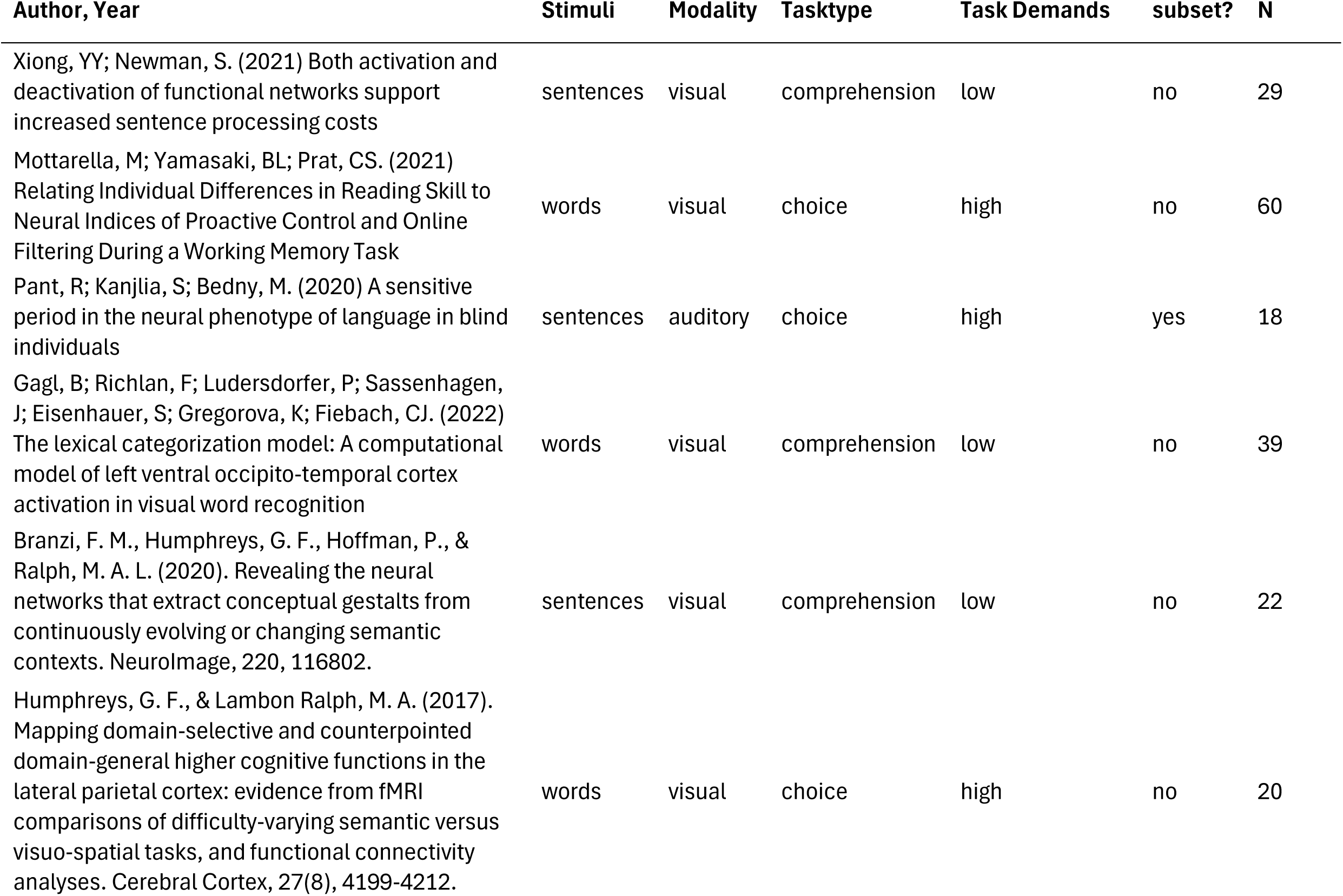
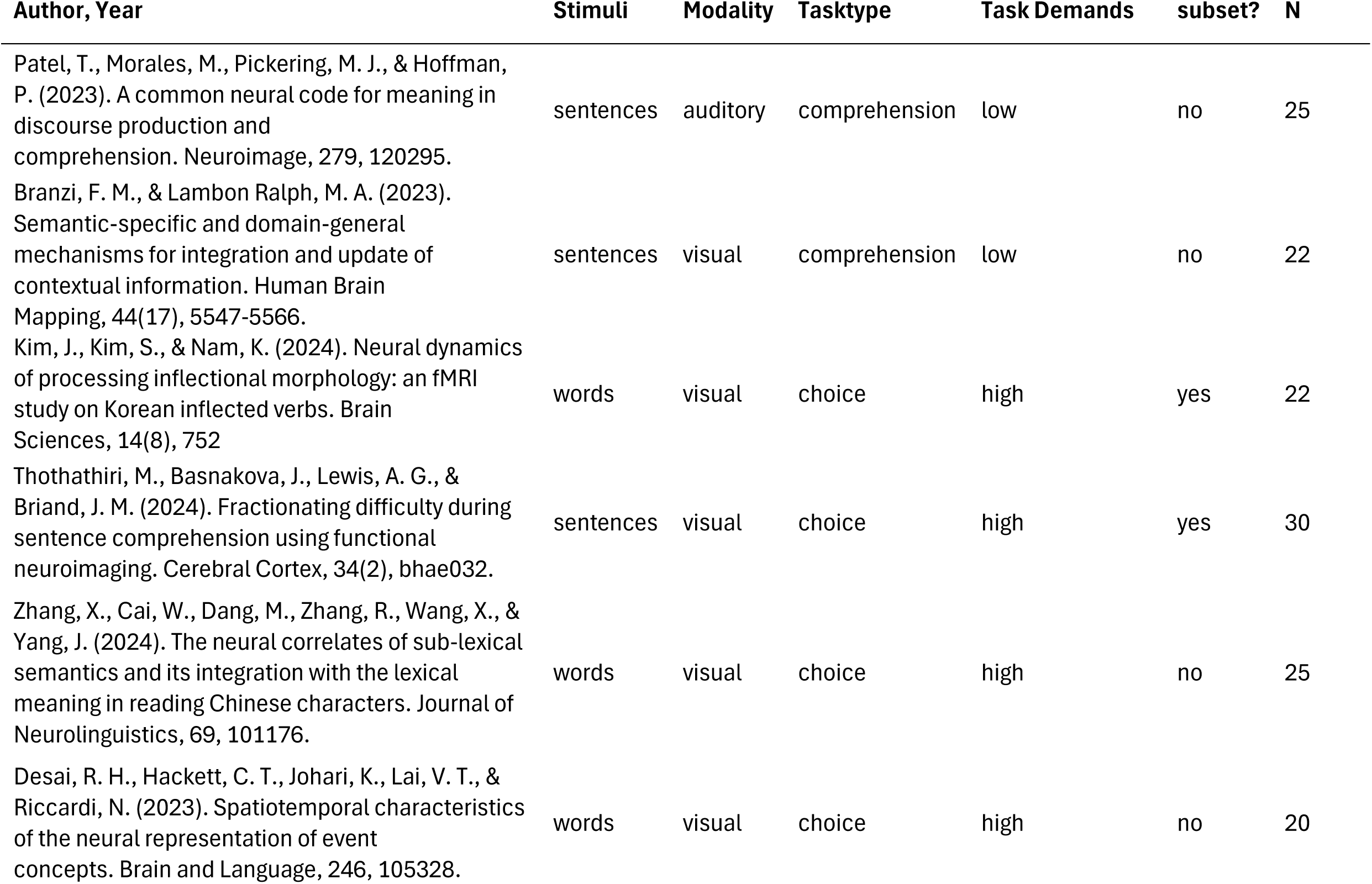
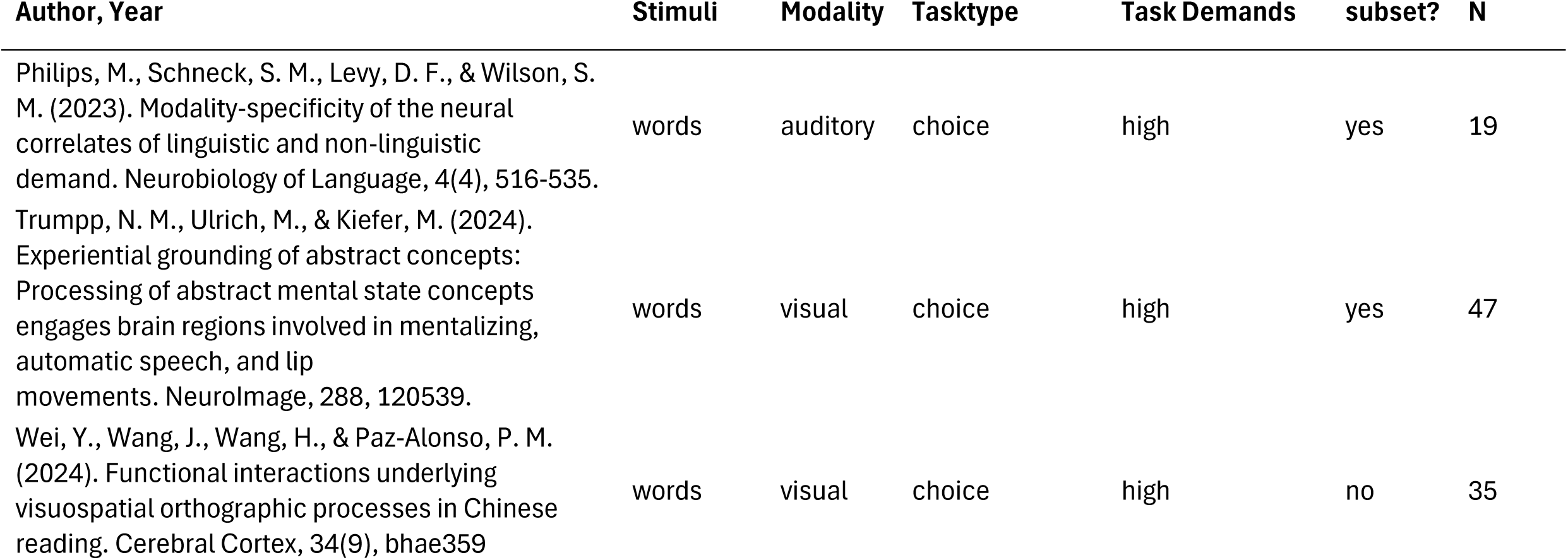
List of studies included in the **Semantic Cognition** meta-analysis. Studies have been categorised based on type of **stimuli** (Single words/Word pairs or Sentences/Narratives); **modality** (visual or auditory or both), **type of task** (choice or comprehension or production/internal generation (thinking about the right answer)); **task demands** (high task demands or low task demands), and whether the study was included in the **subset** analysed in the choice tasks contrasts. Number of participants is provided (N).

**Supplementary Table 2.**
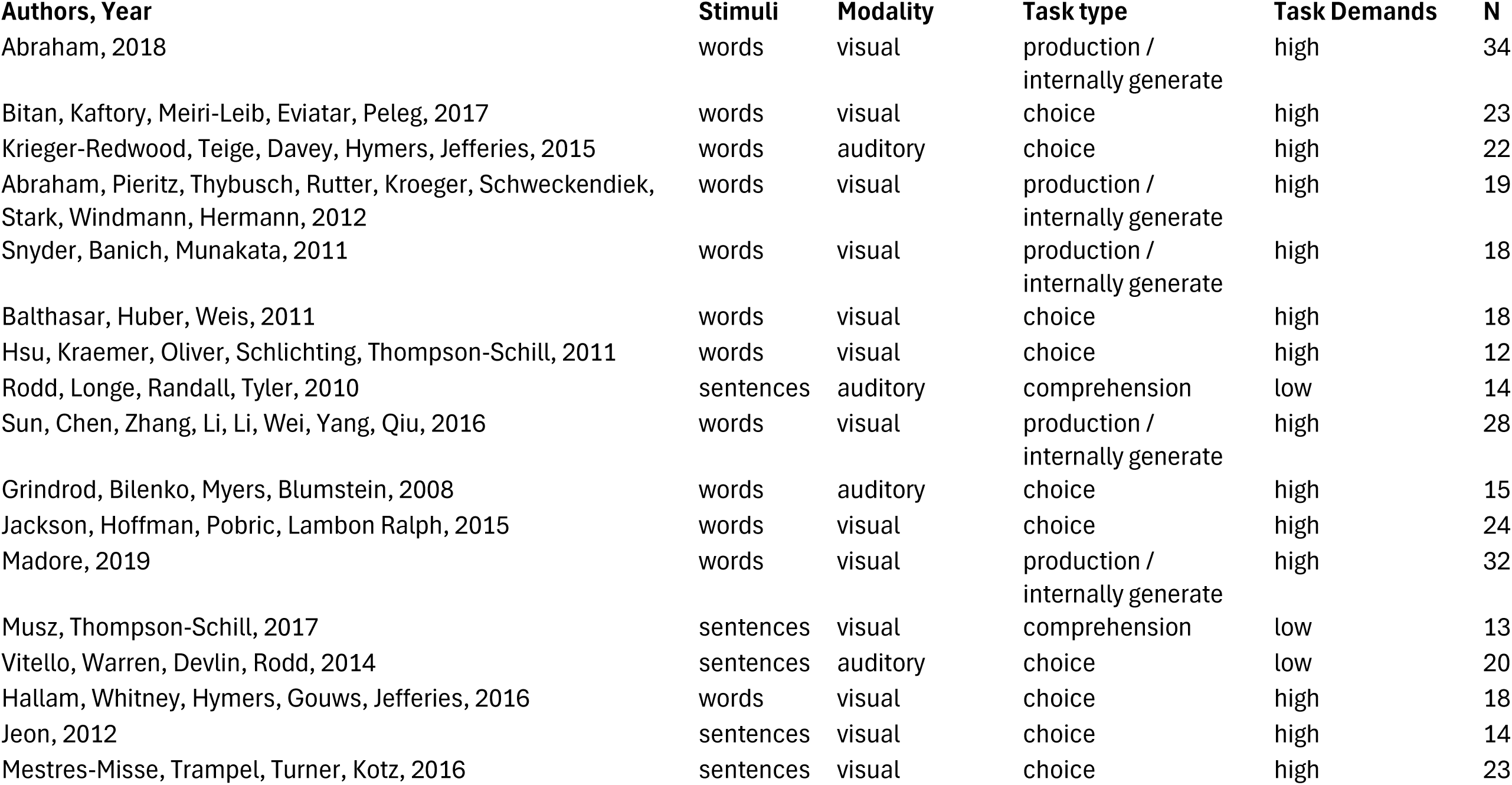
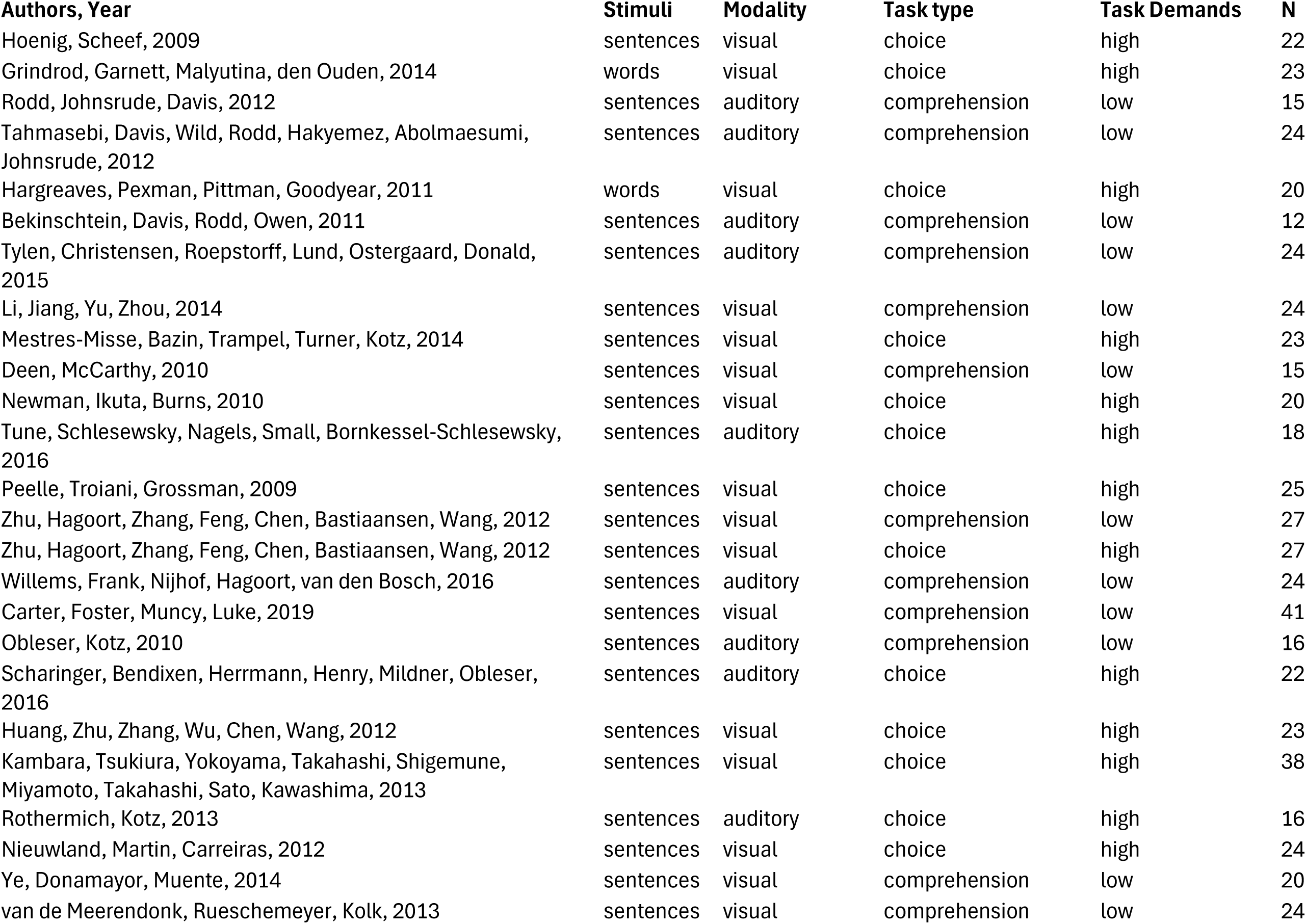
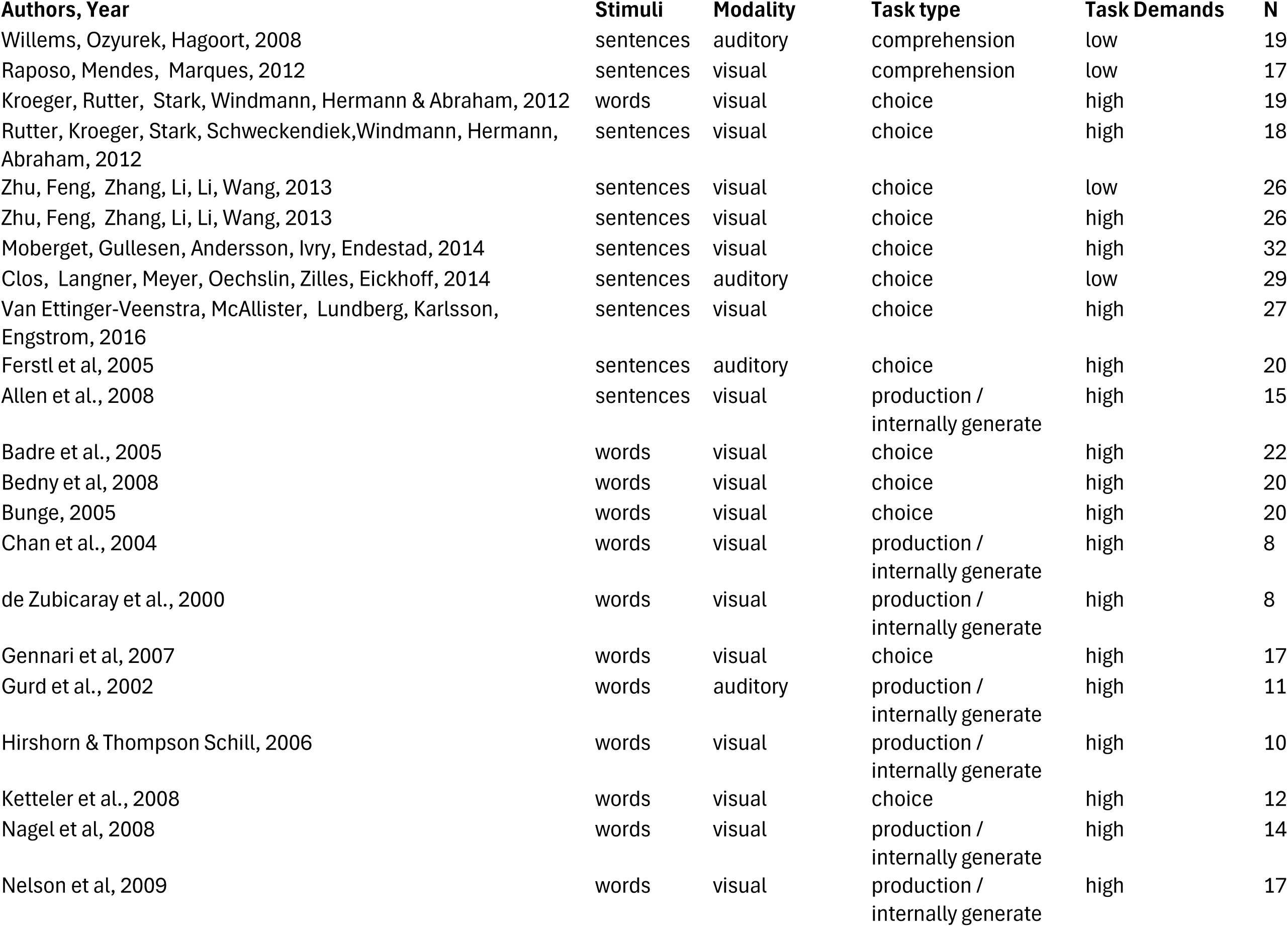
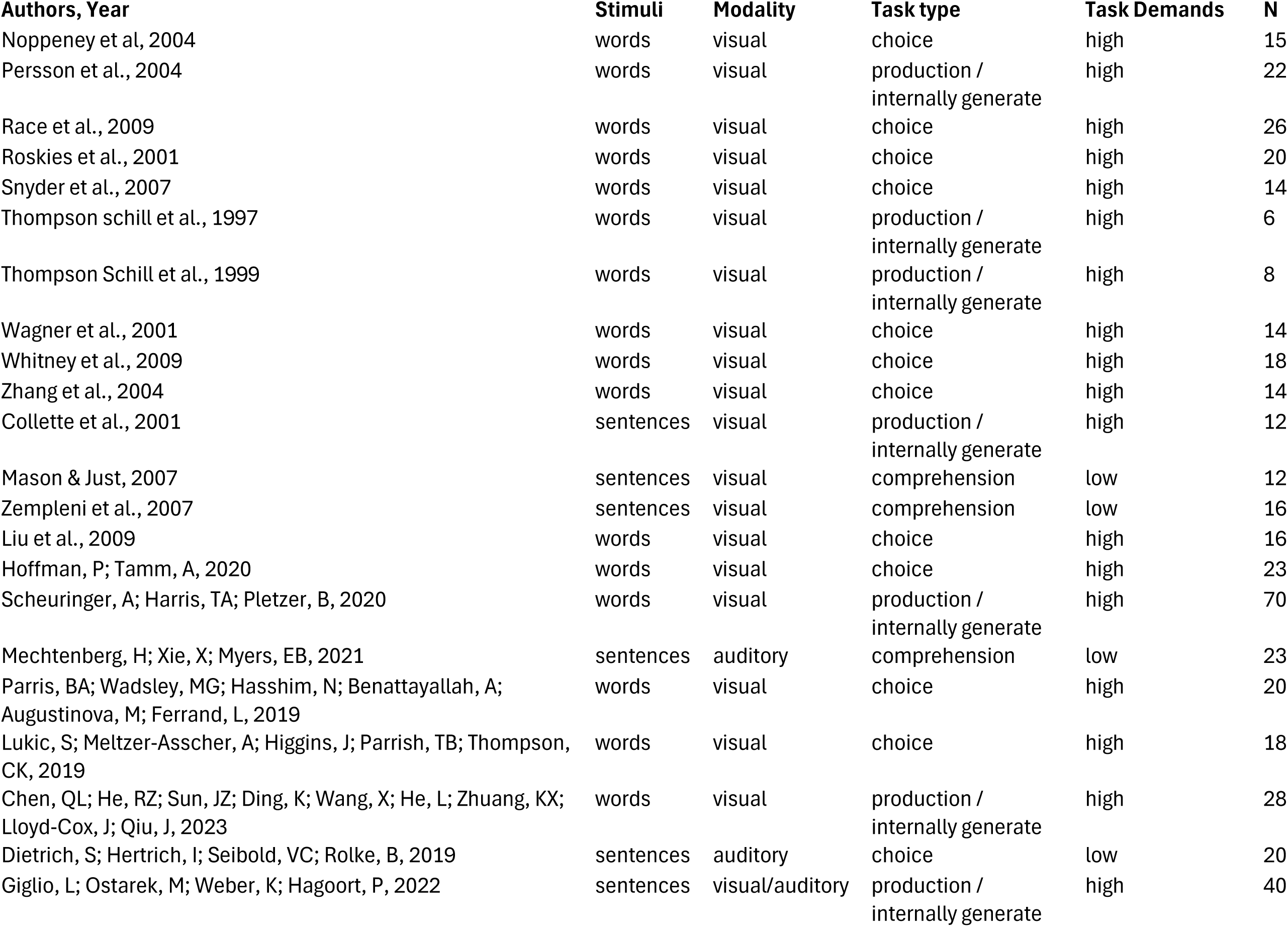
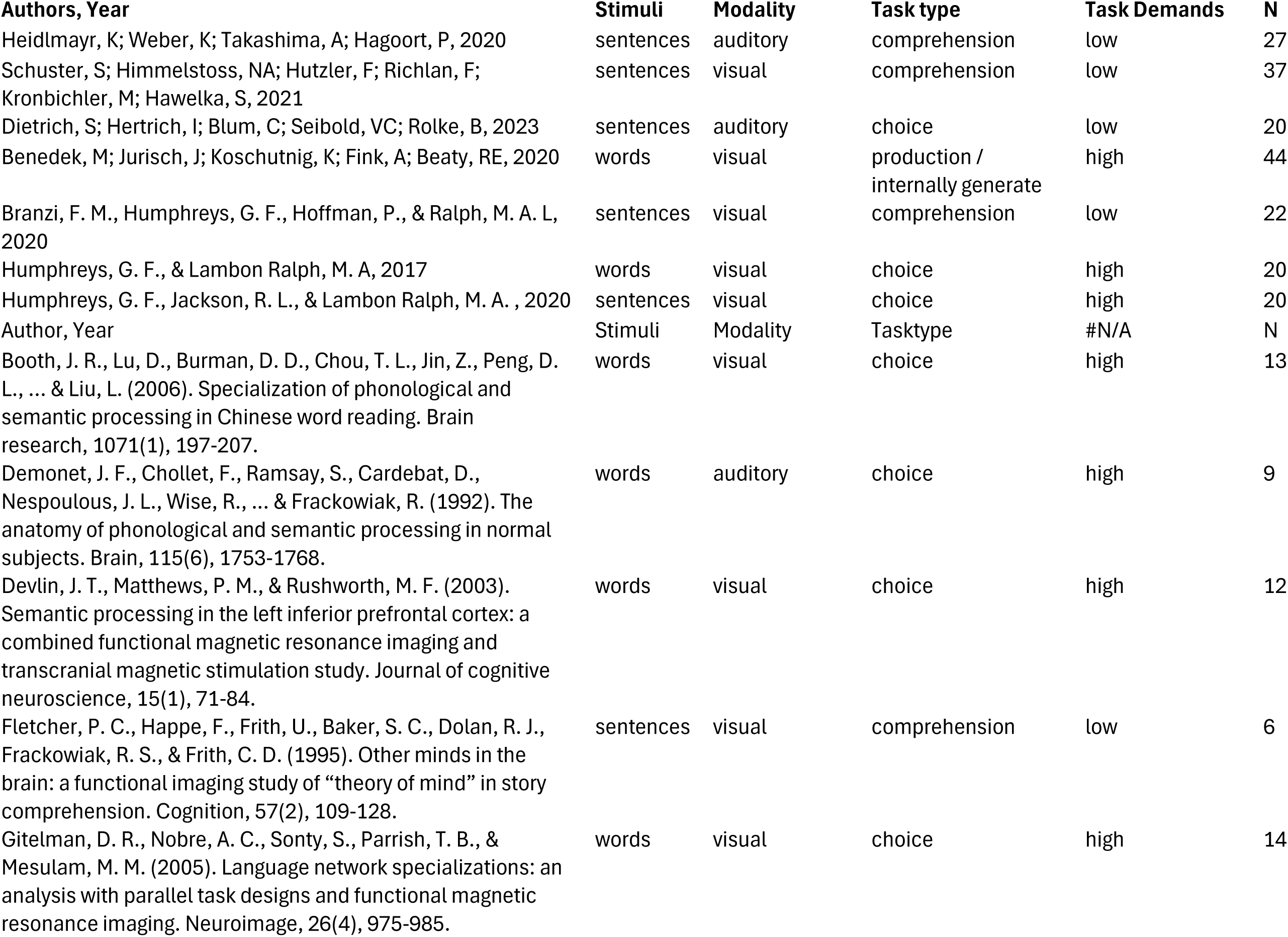
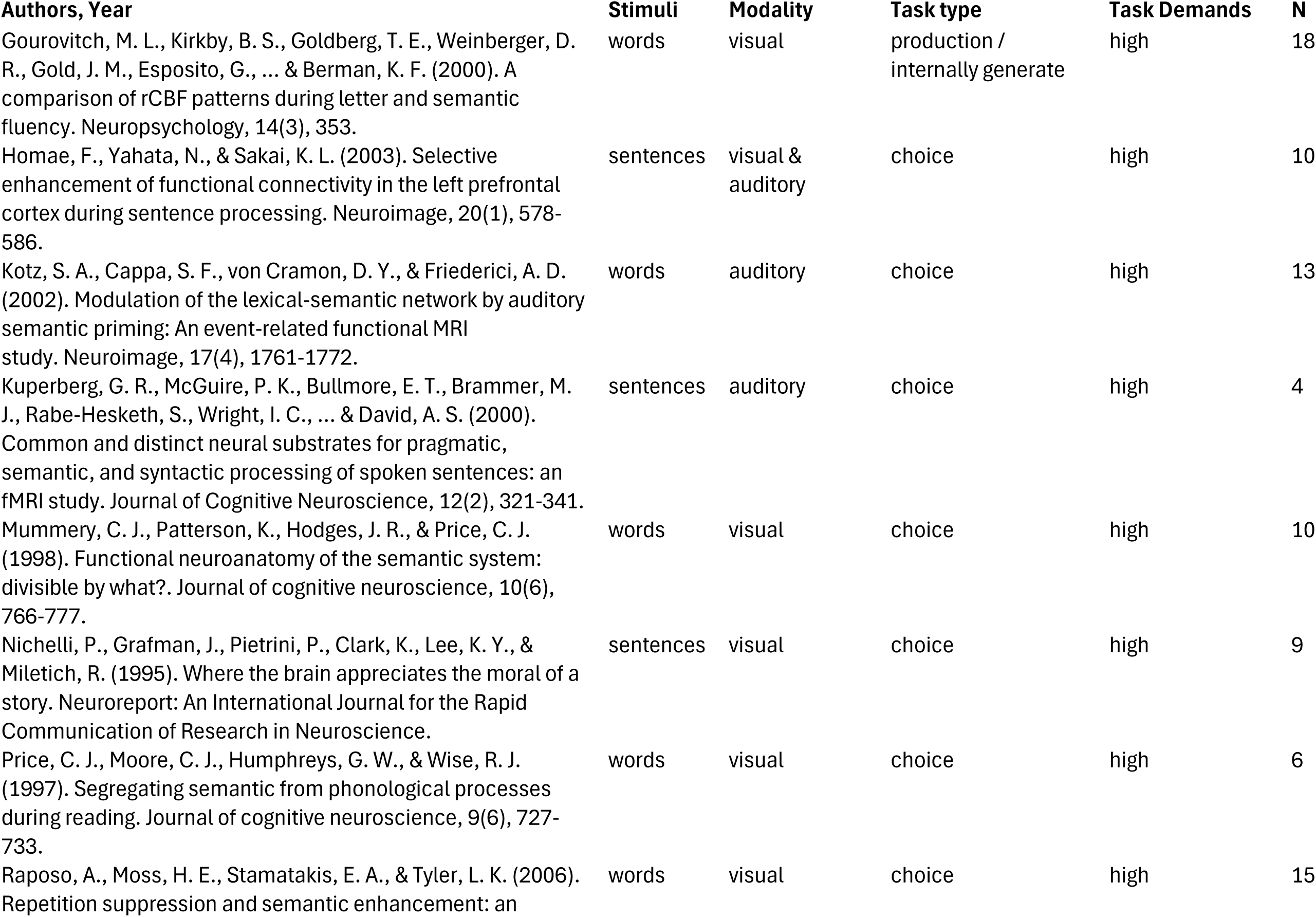
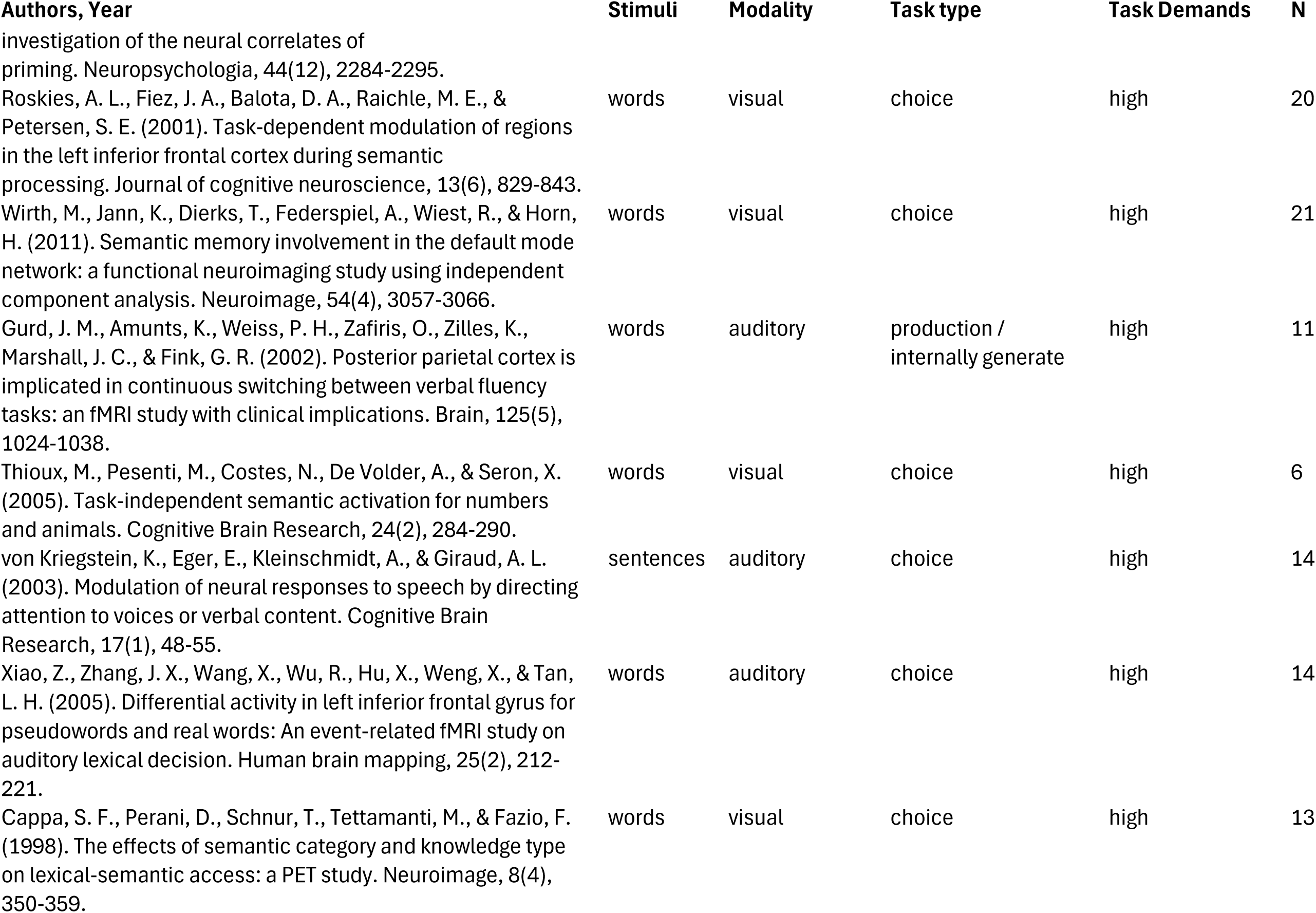
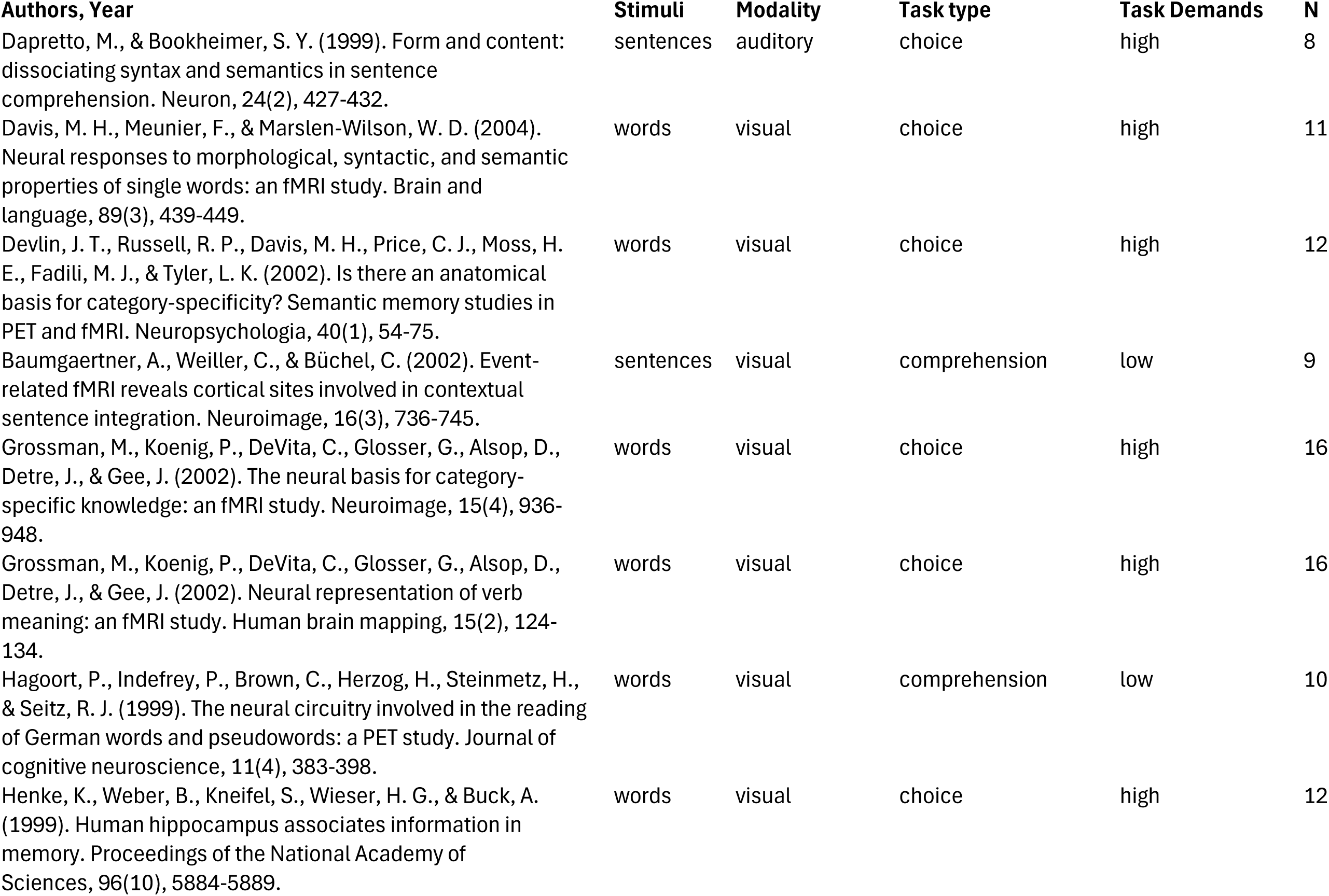
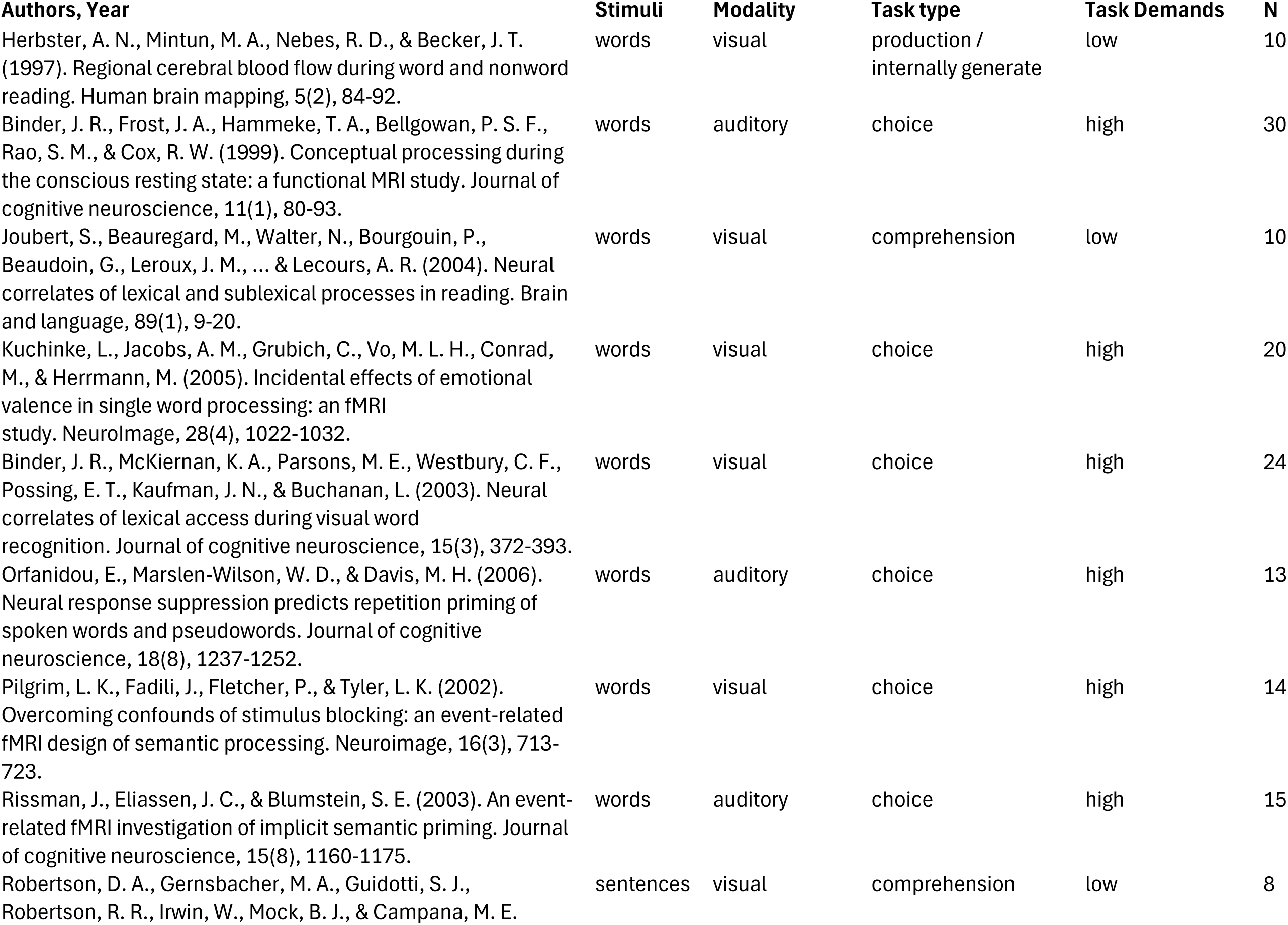
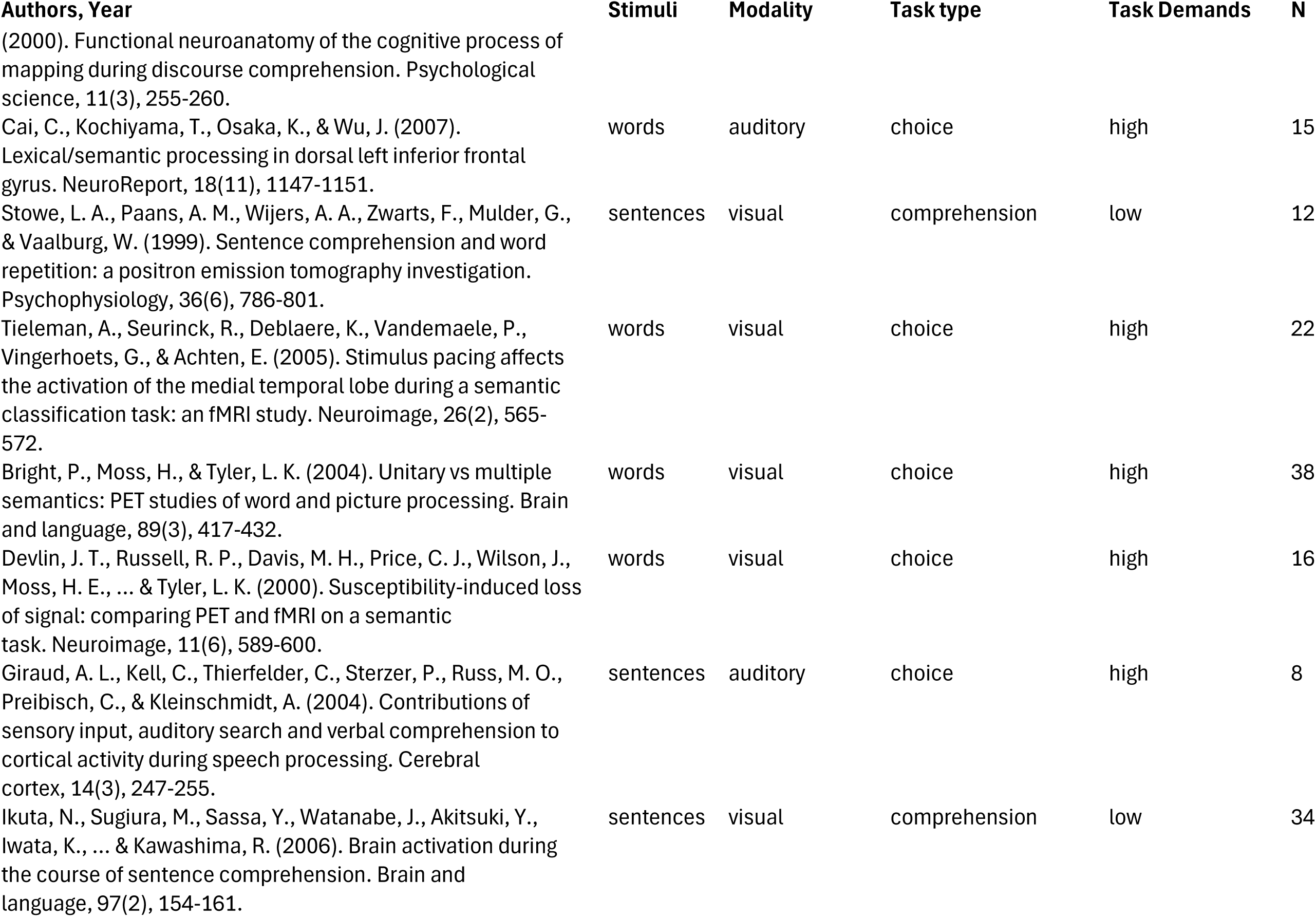
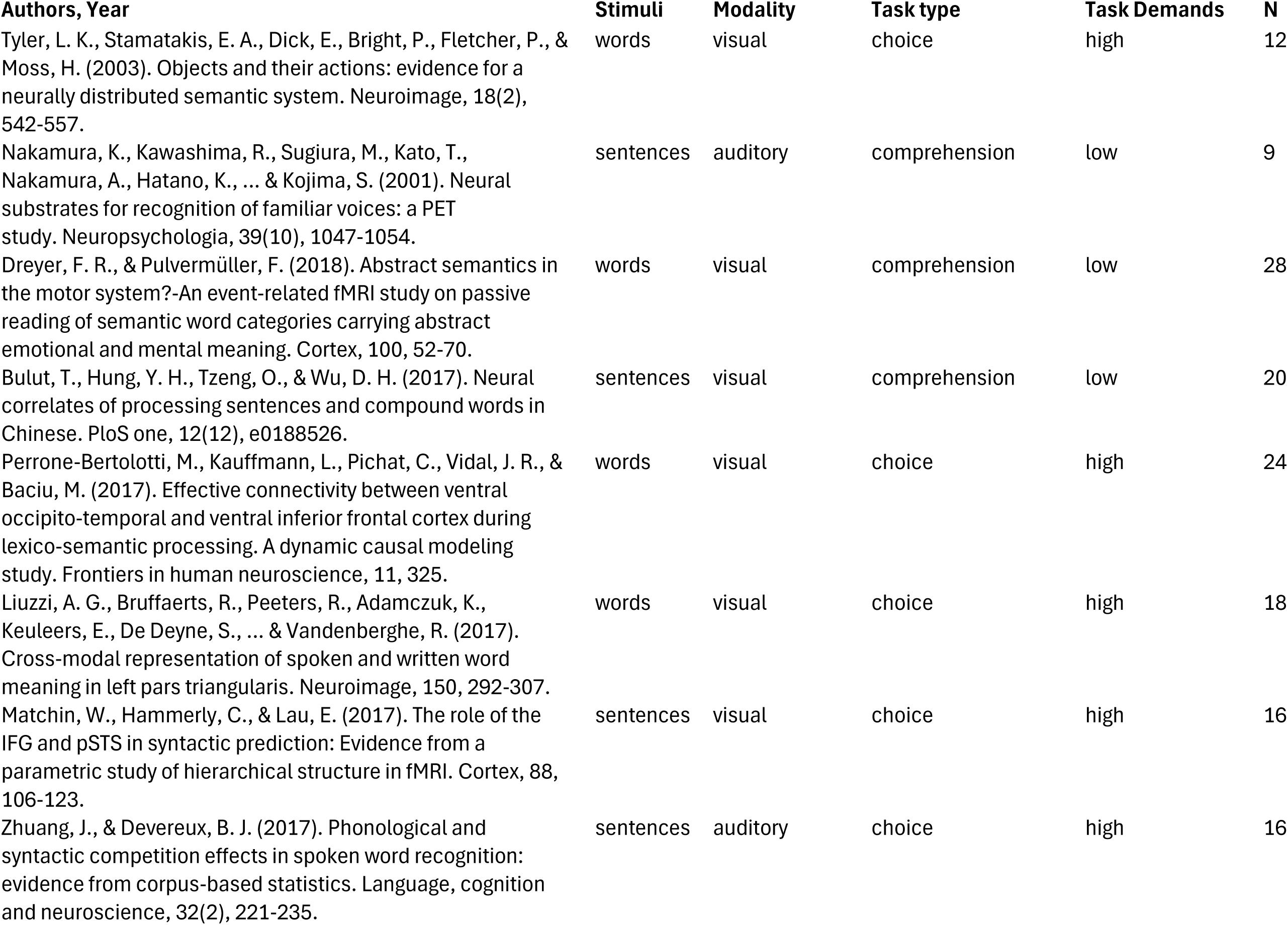
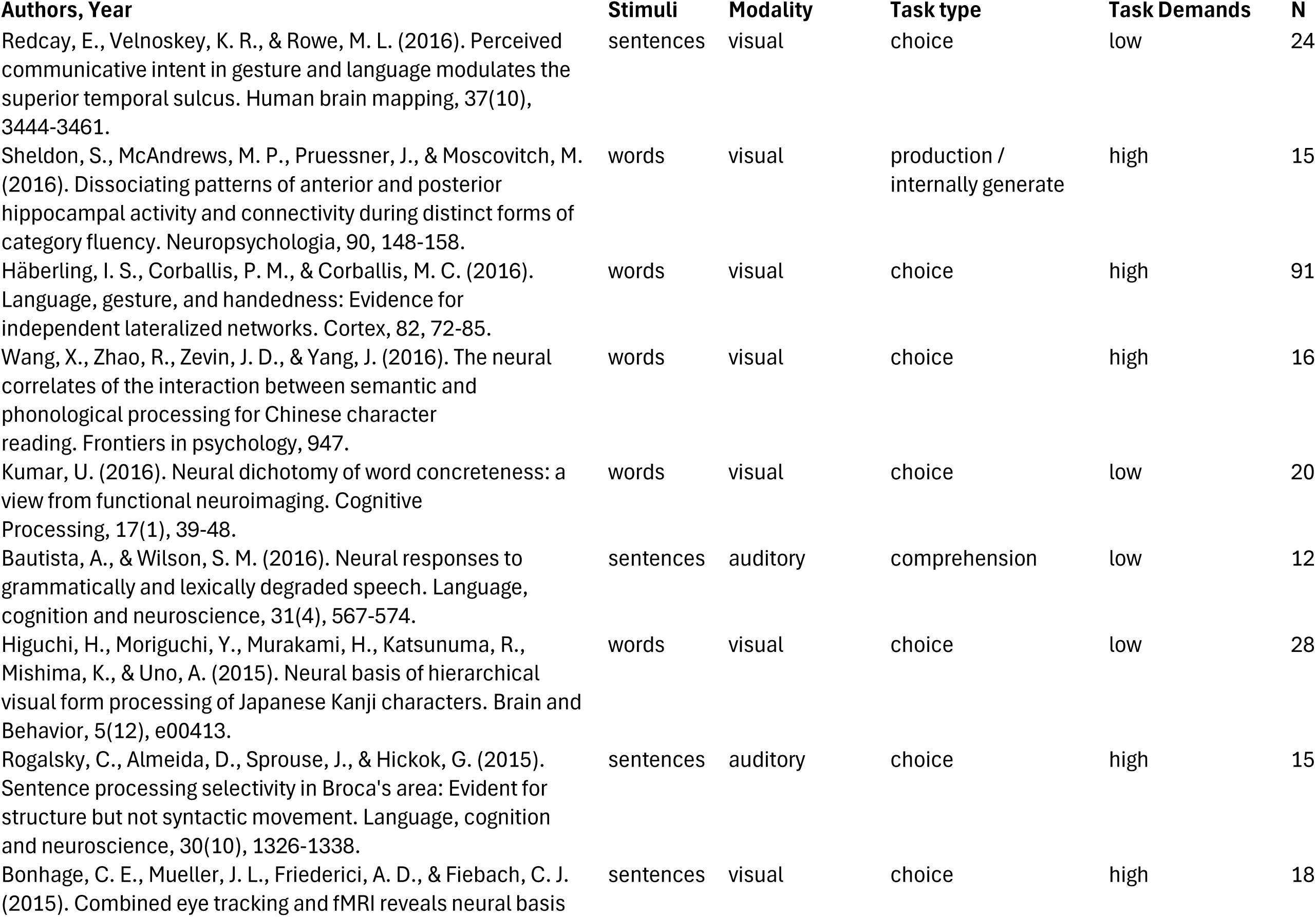
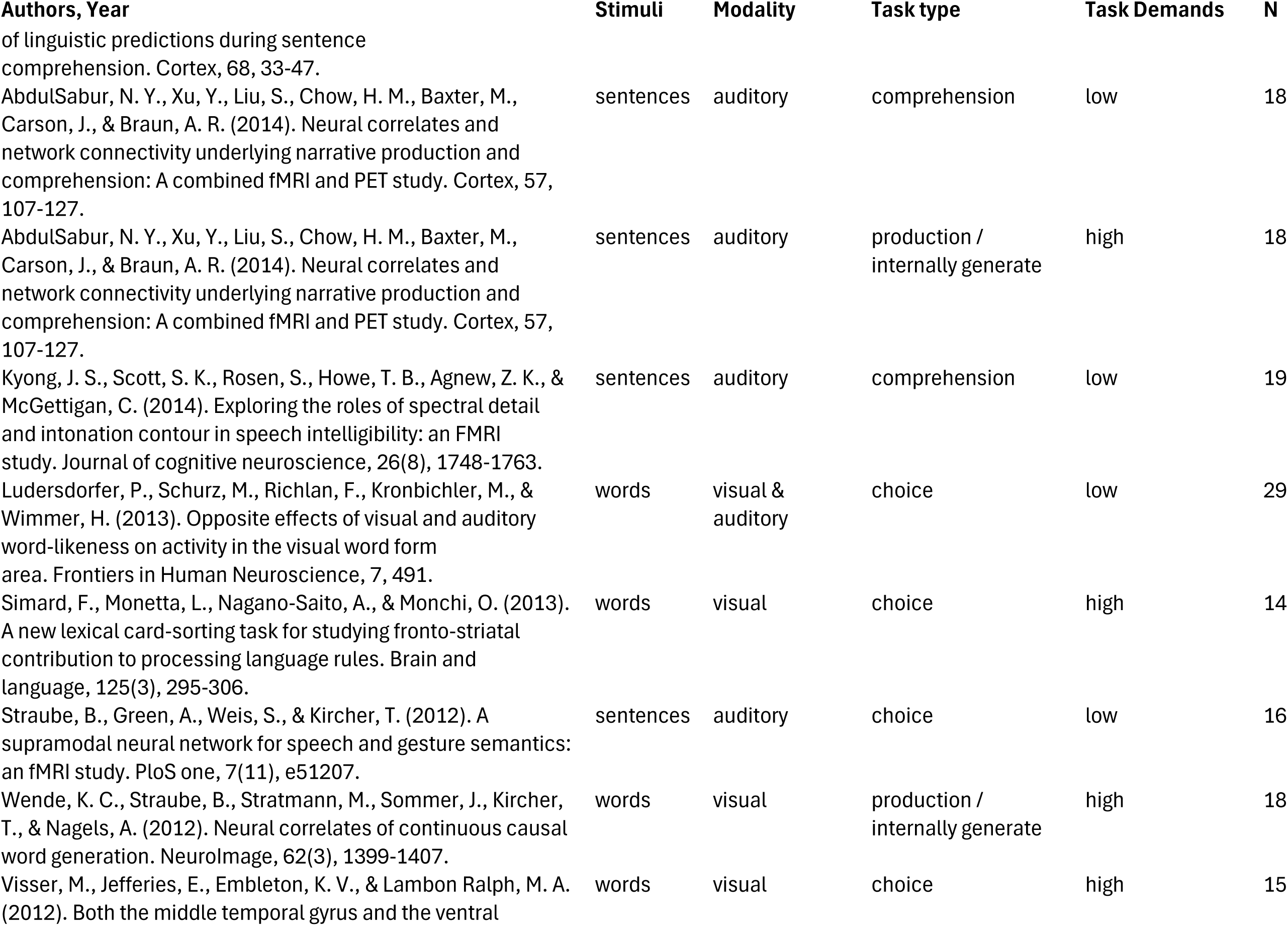
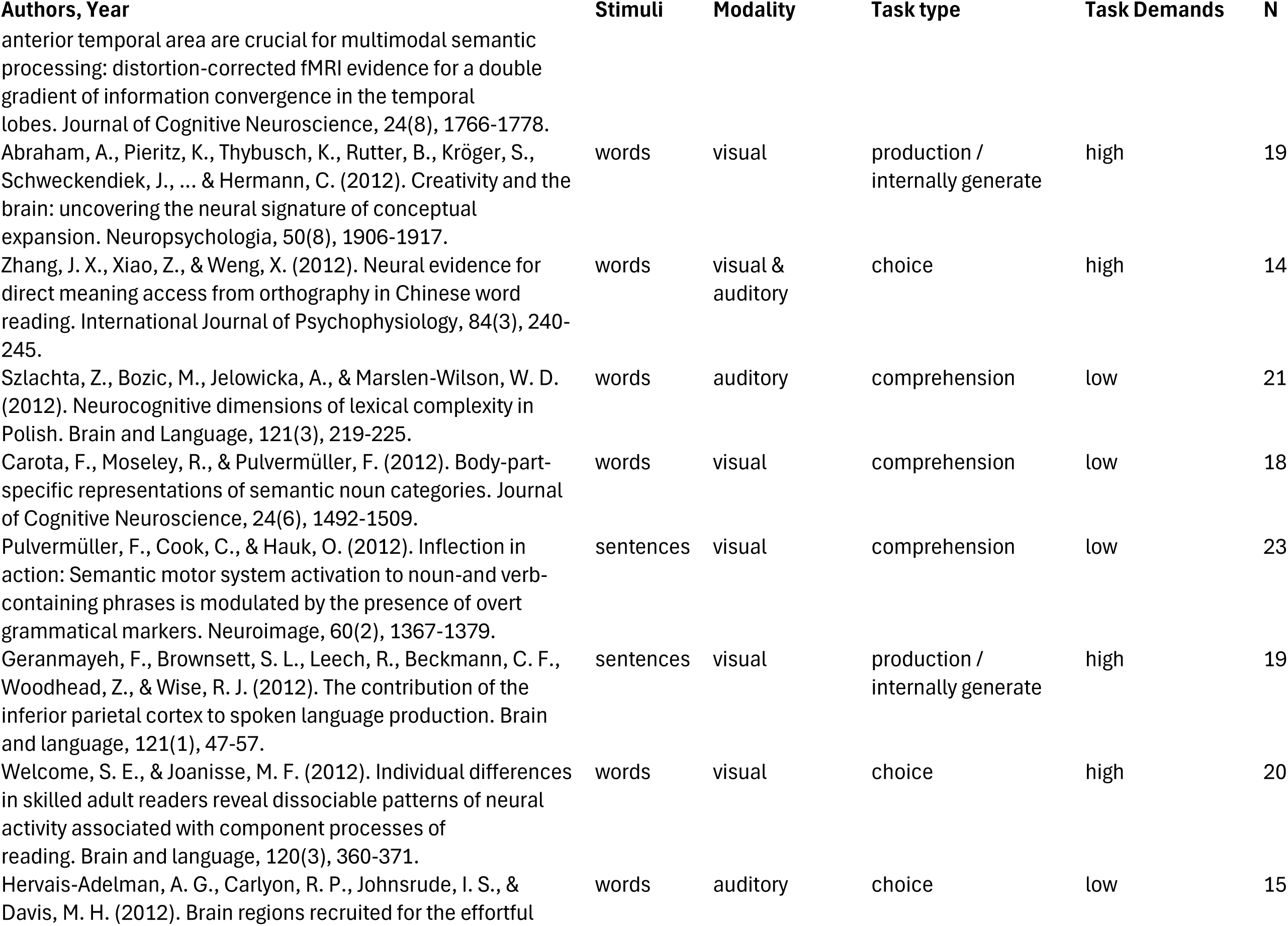
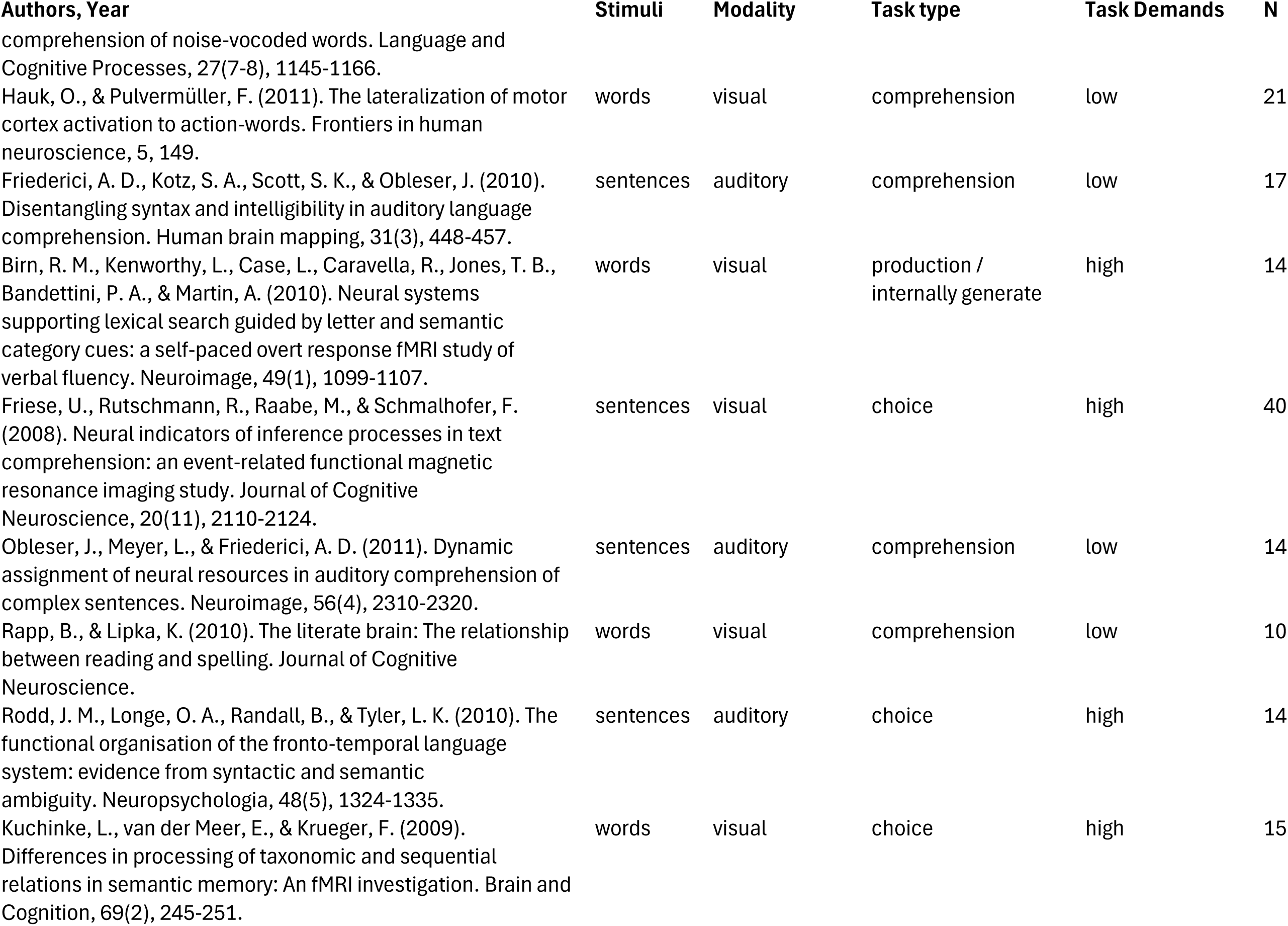
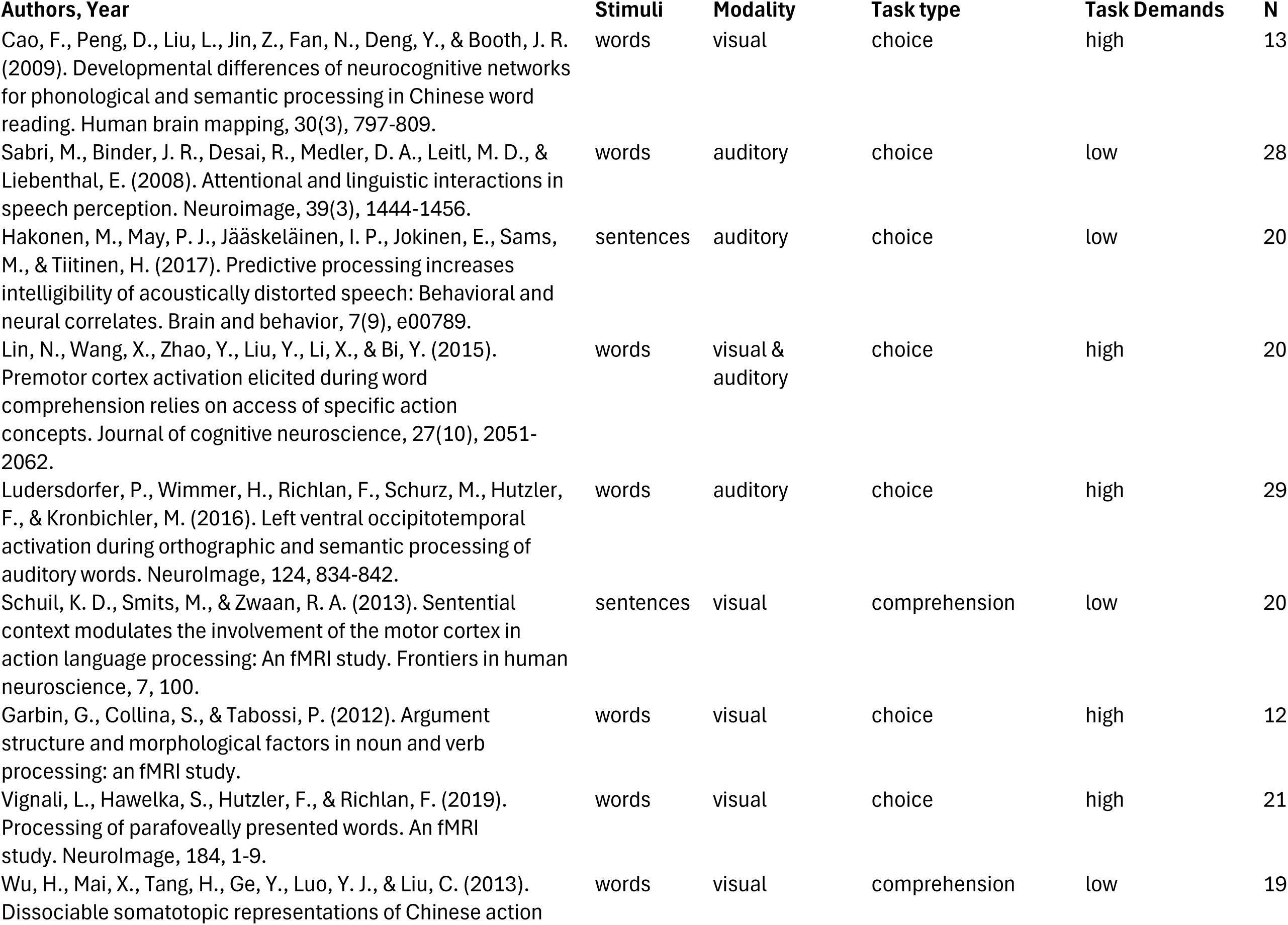
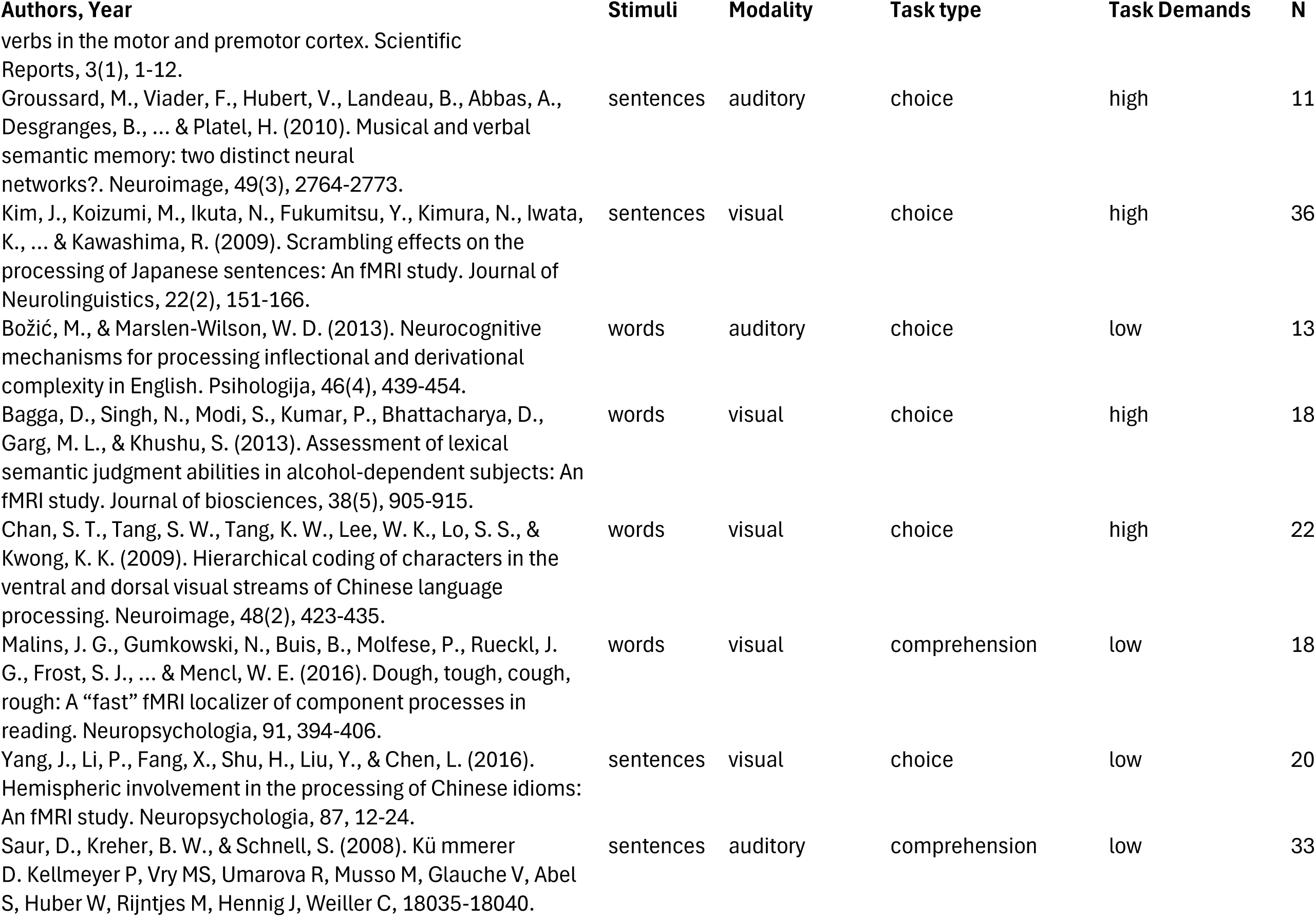
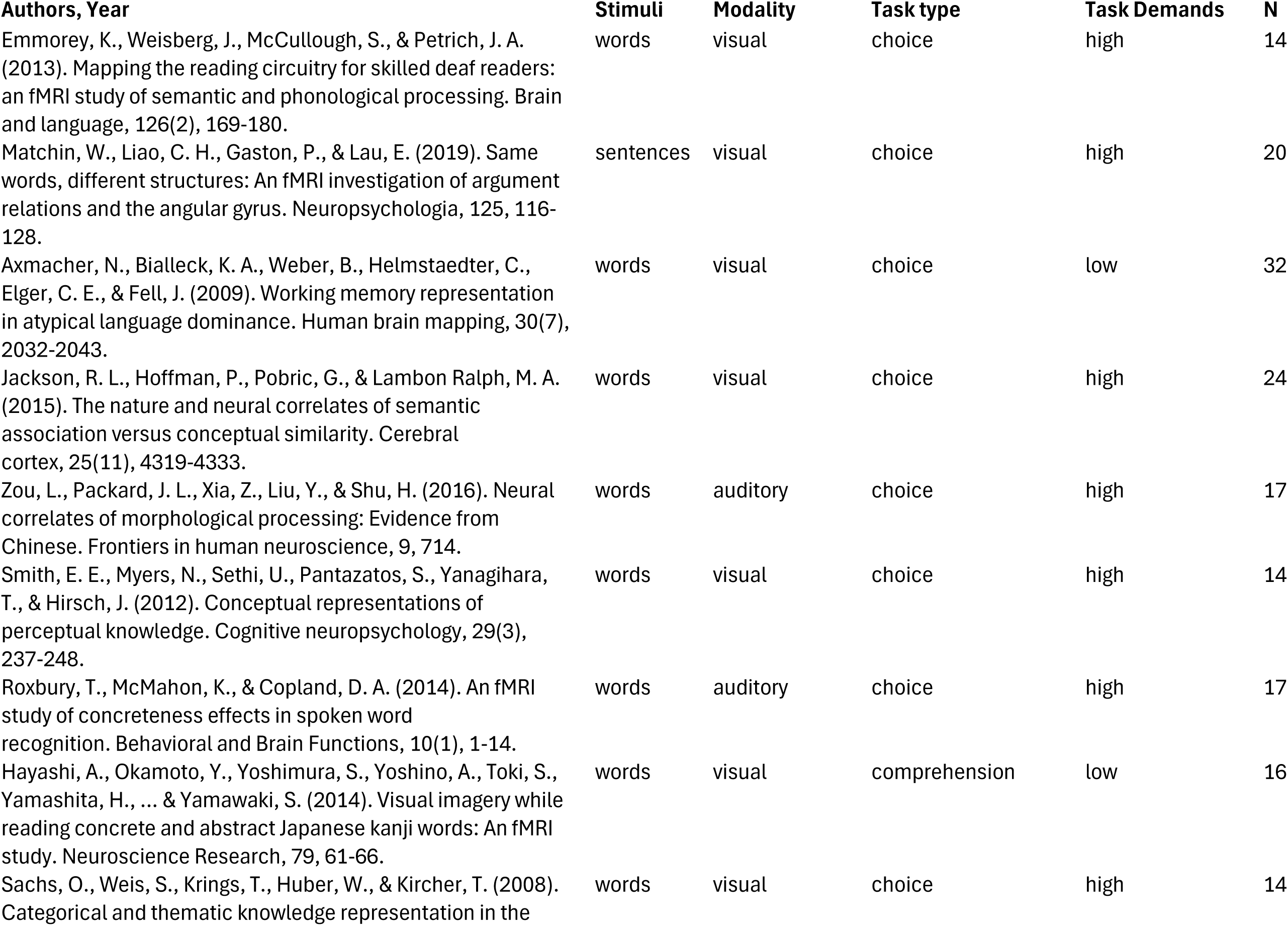
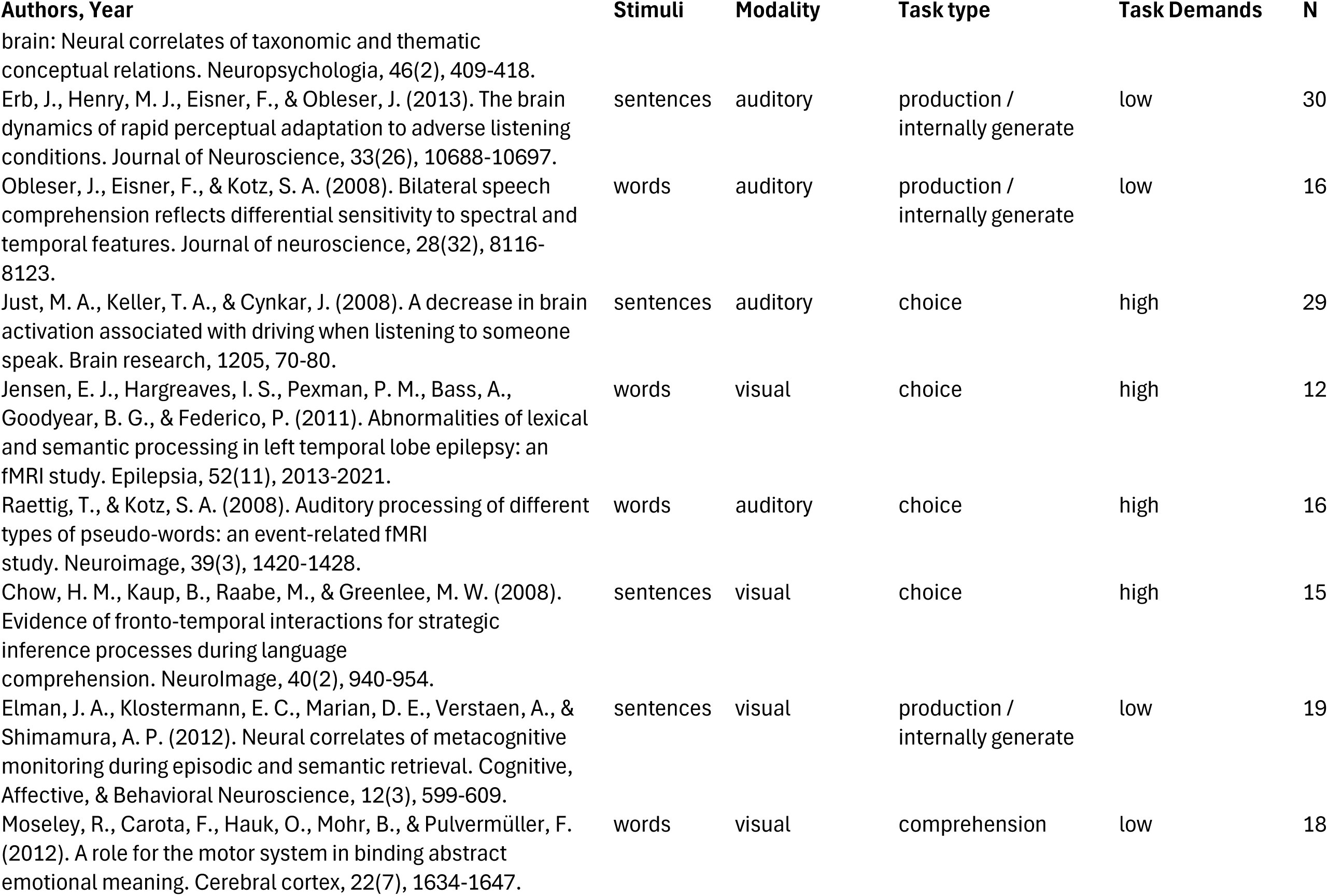
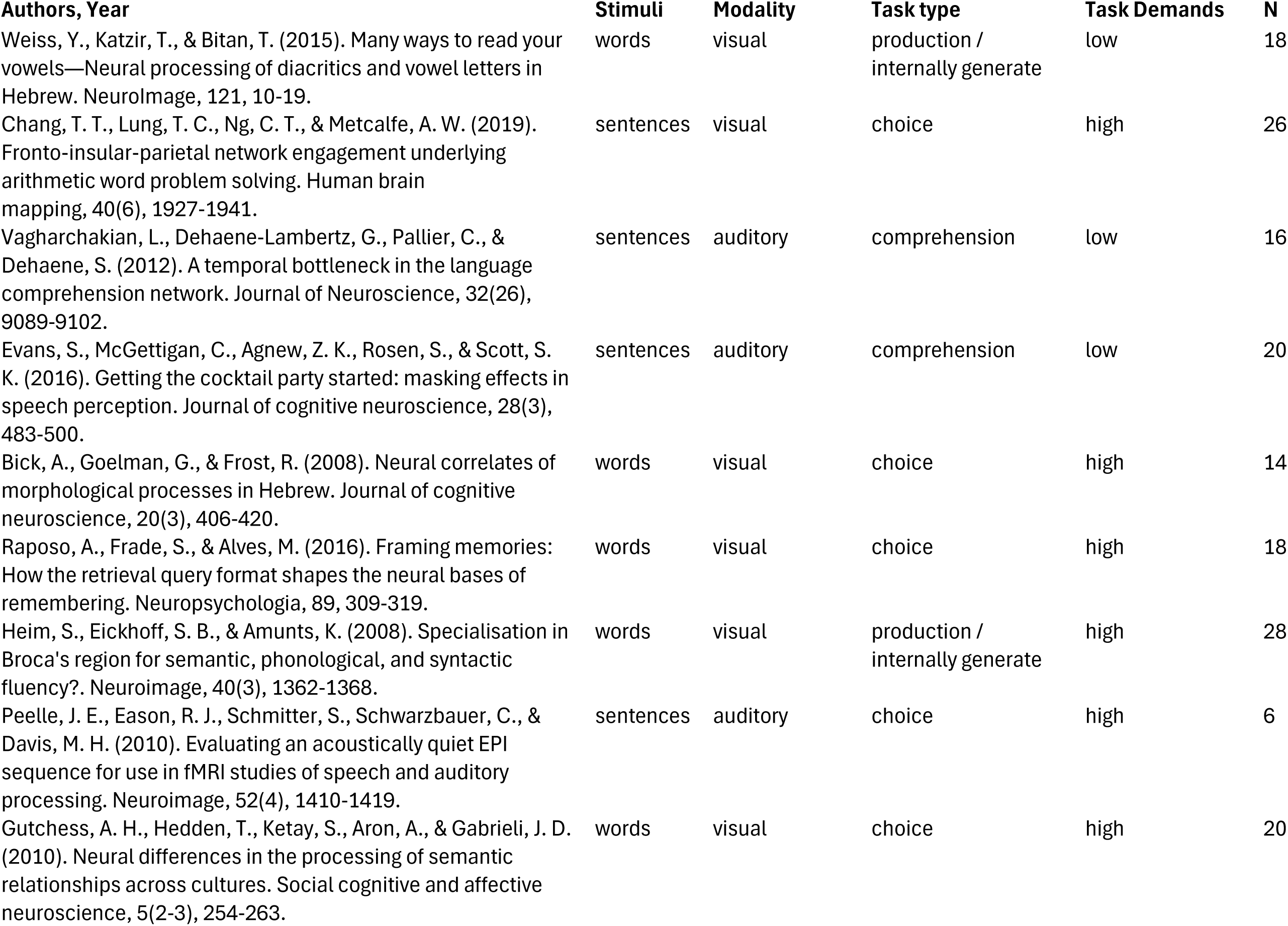
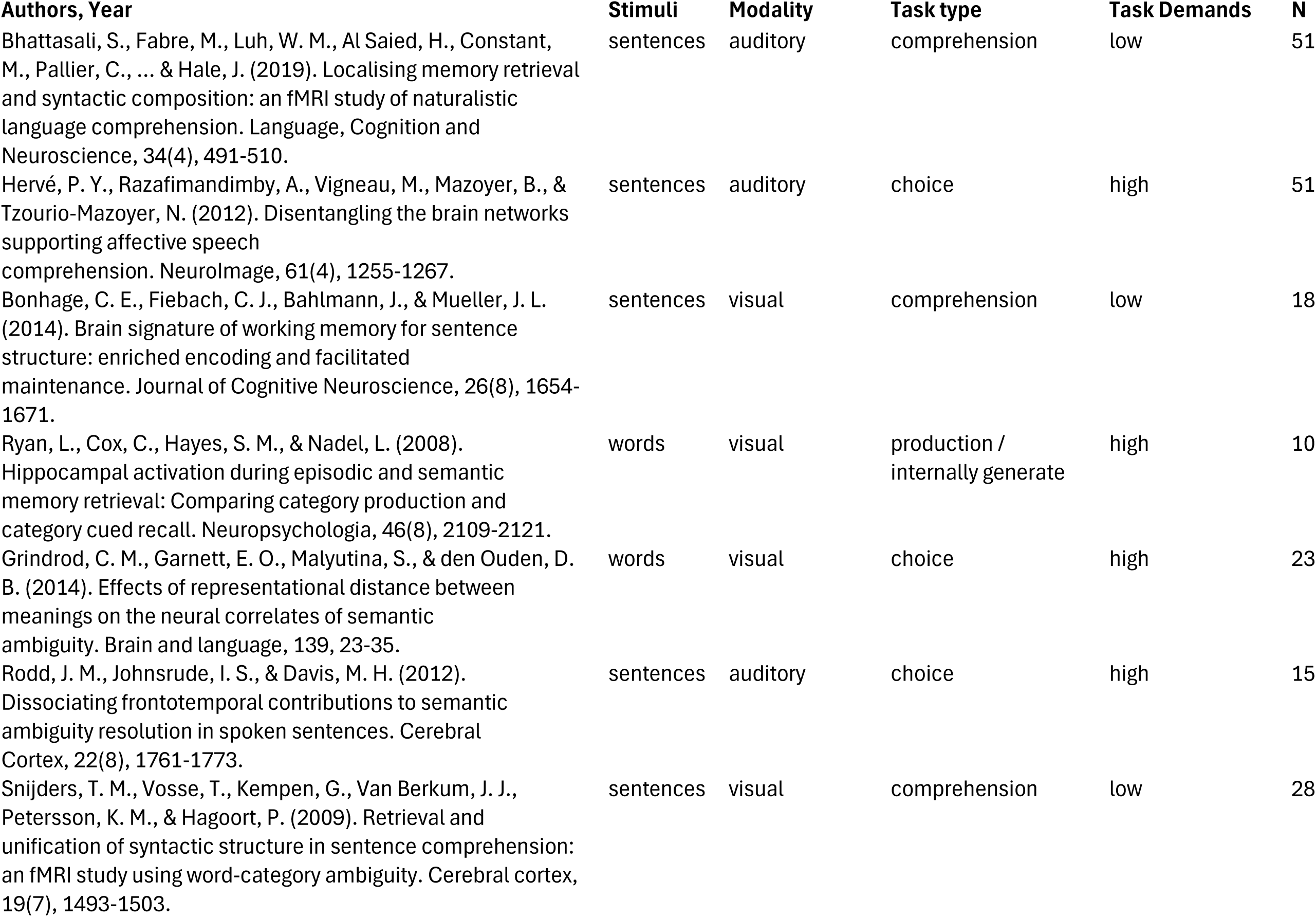
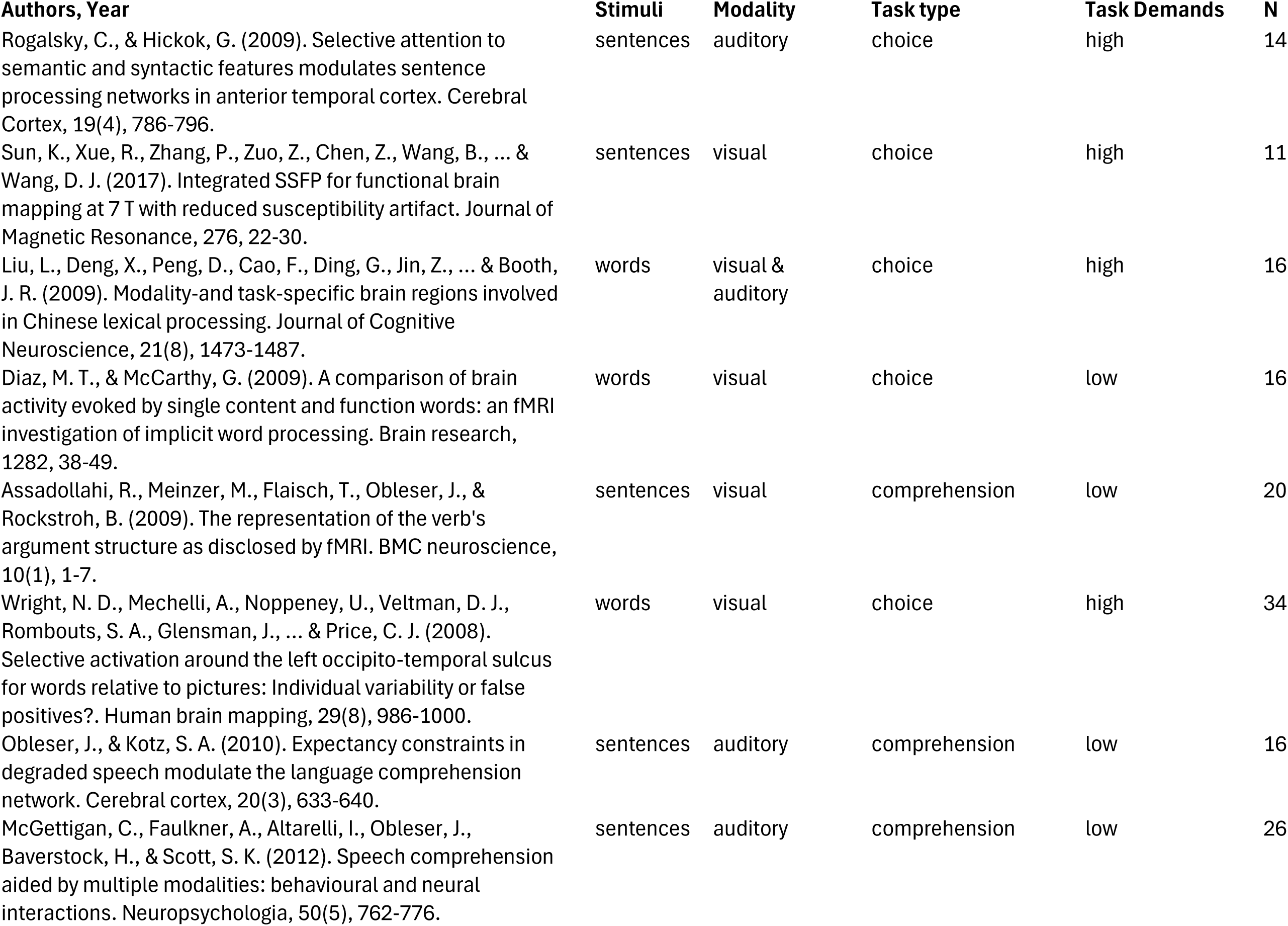
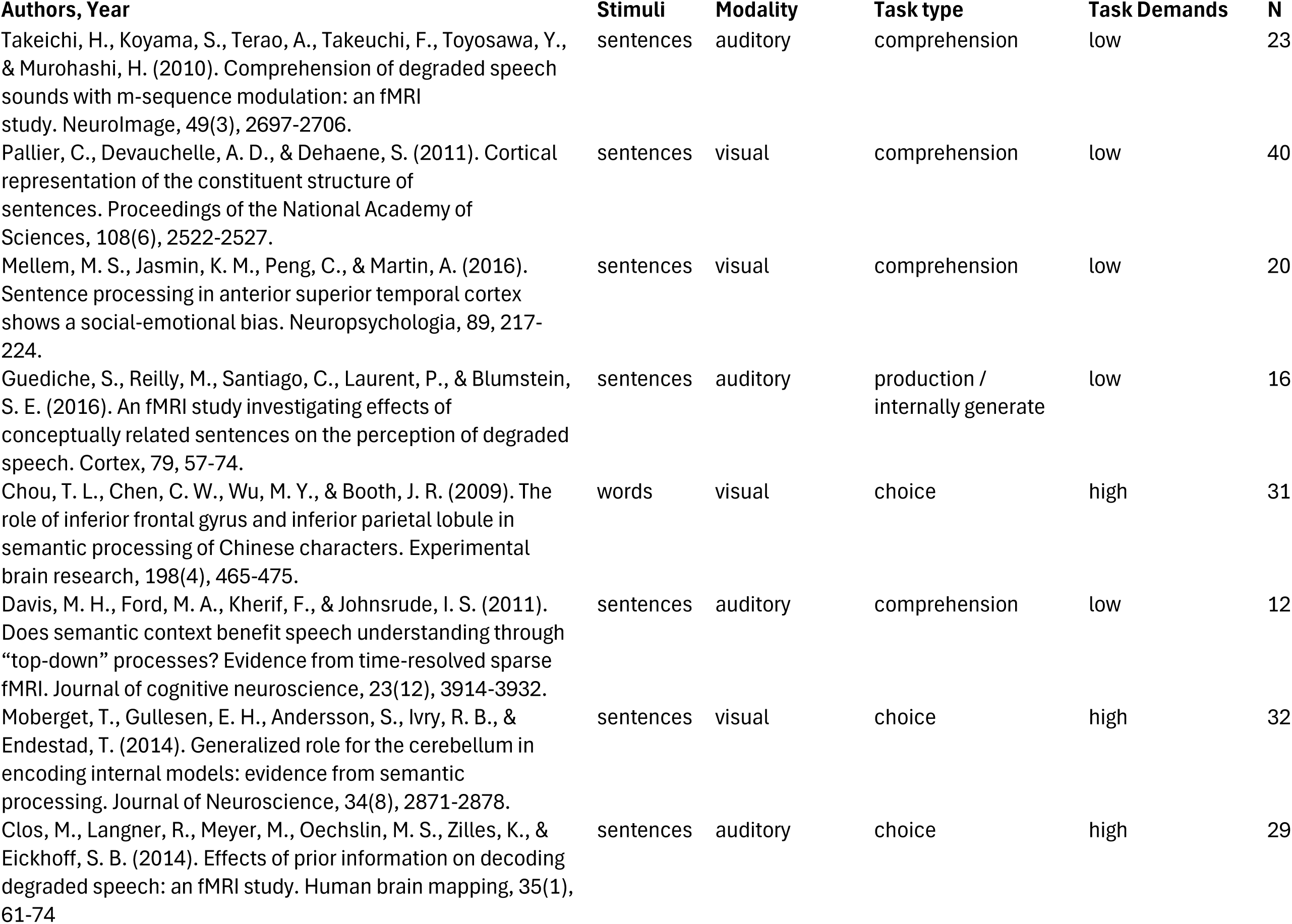
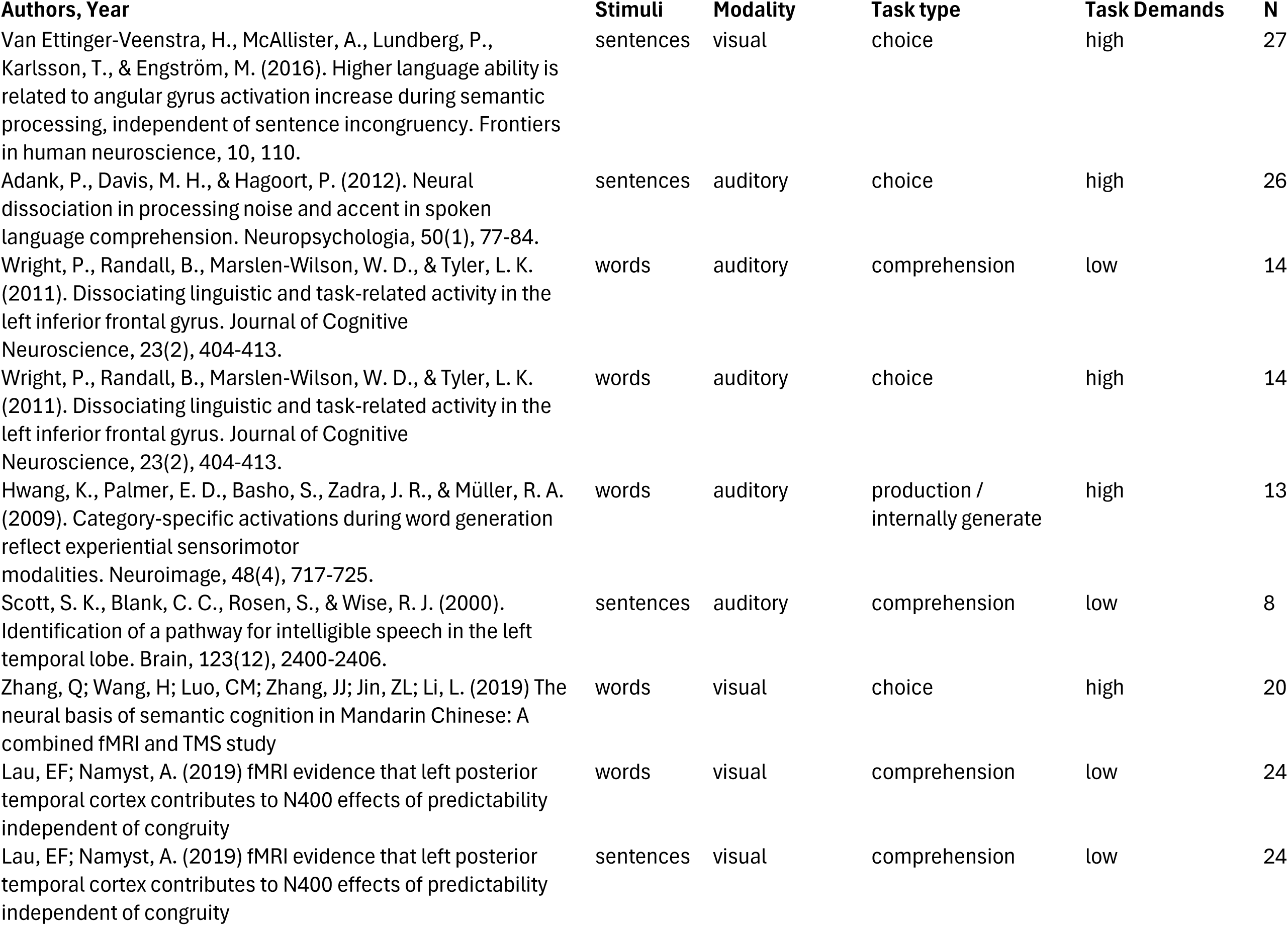
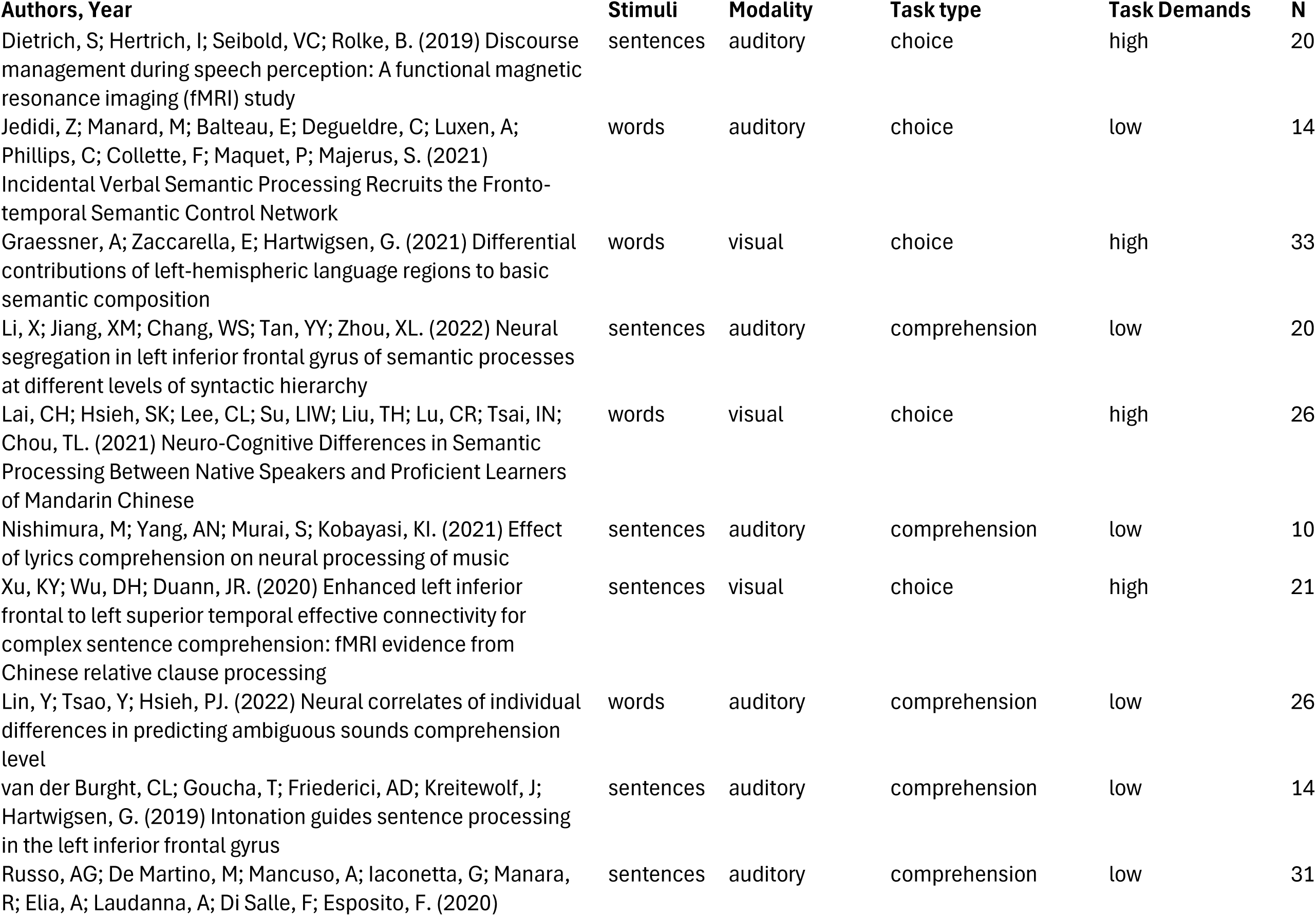
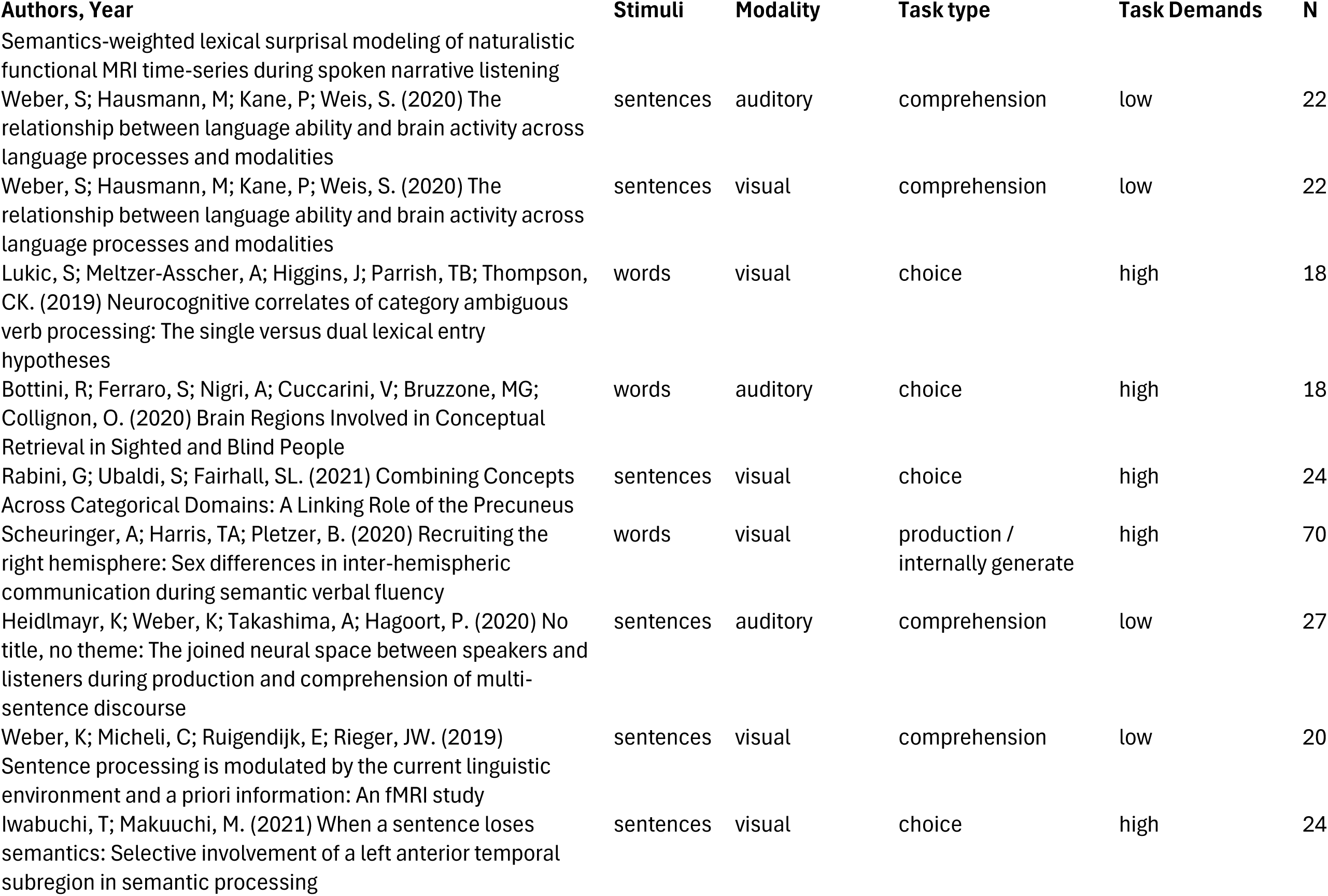
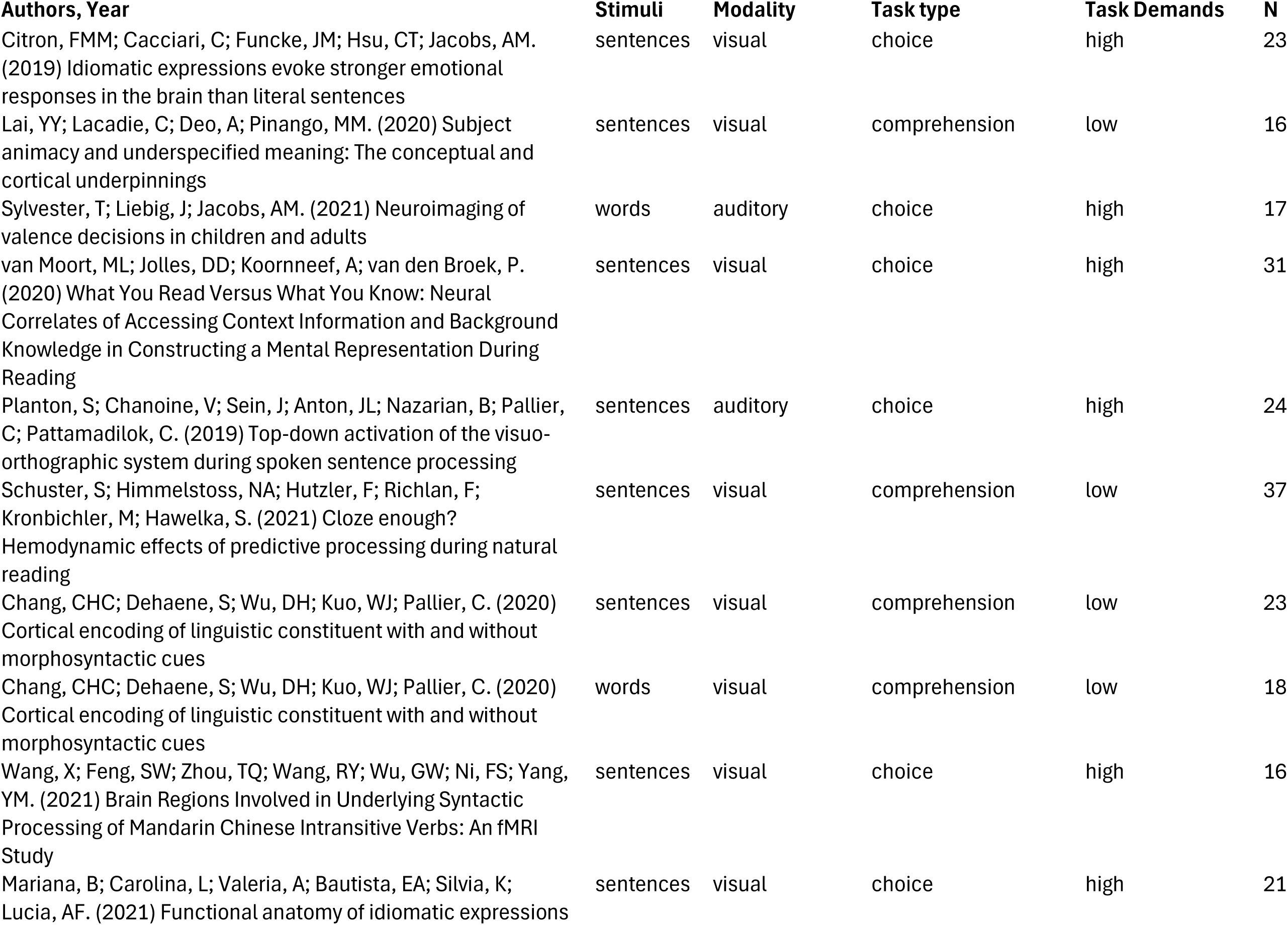
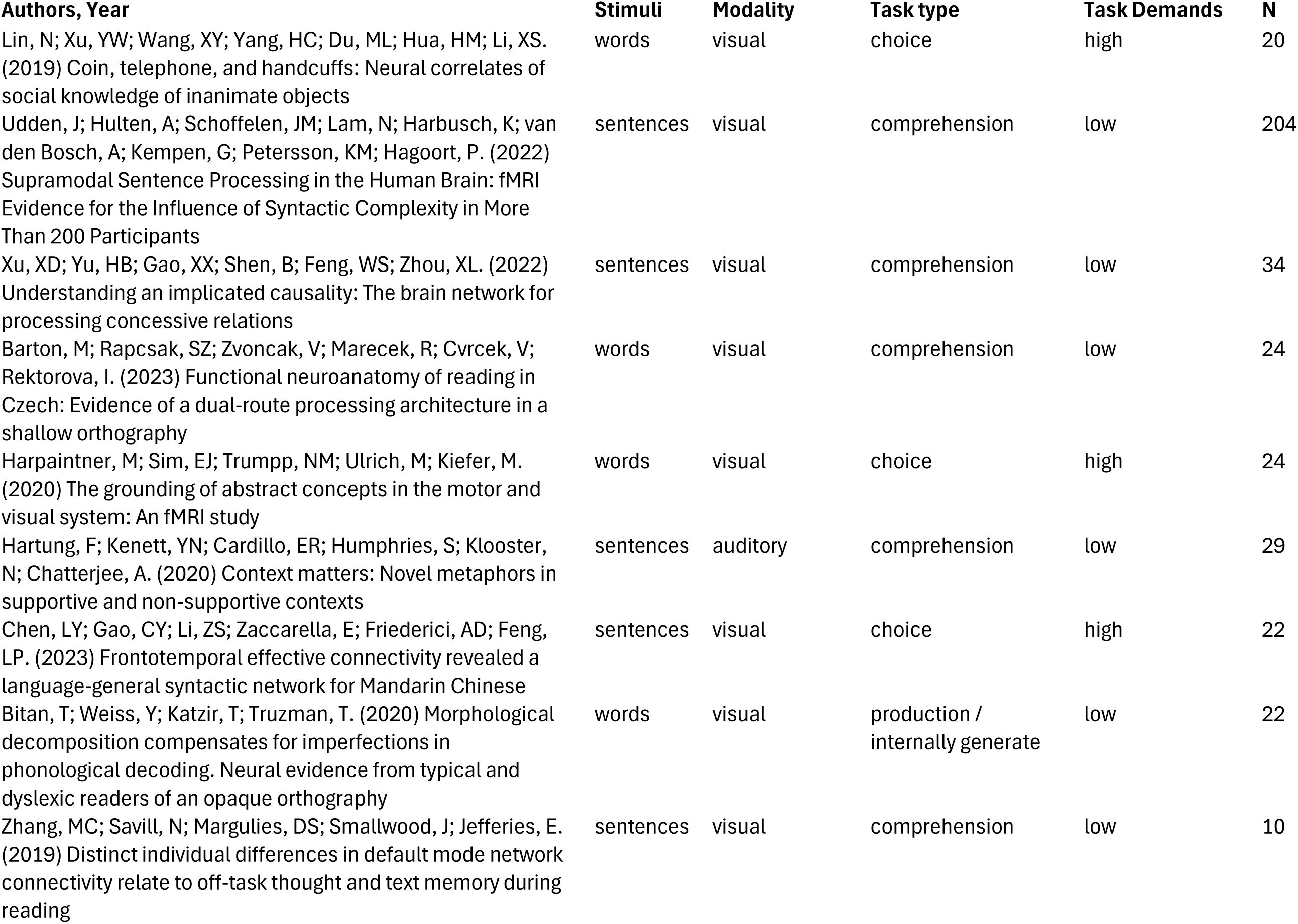
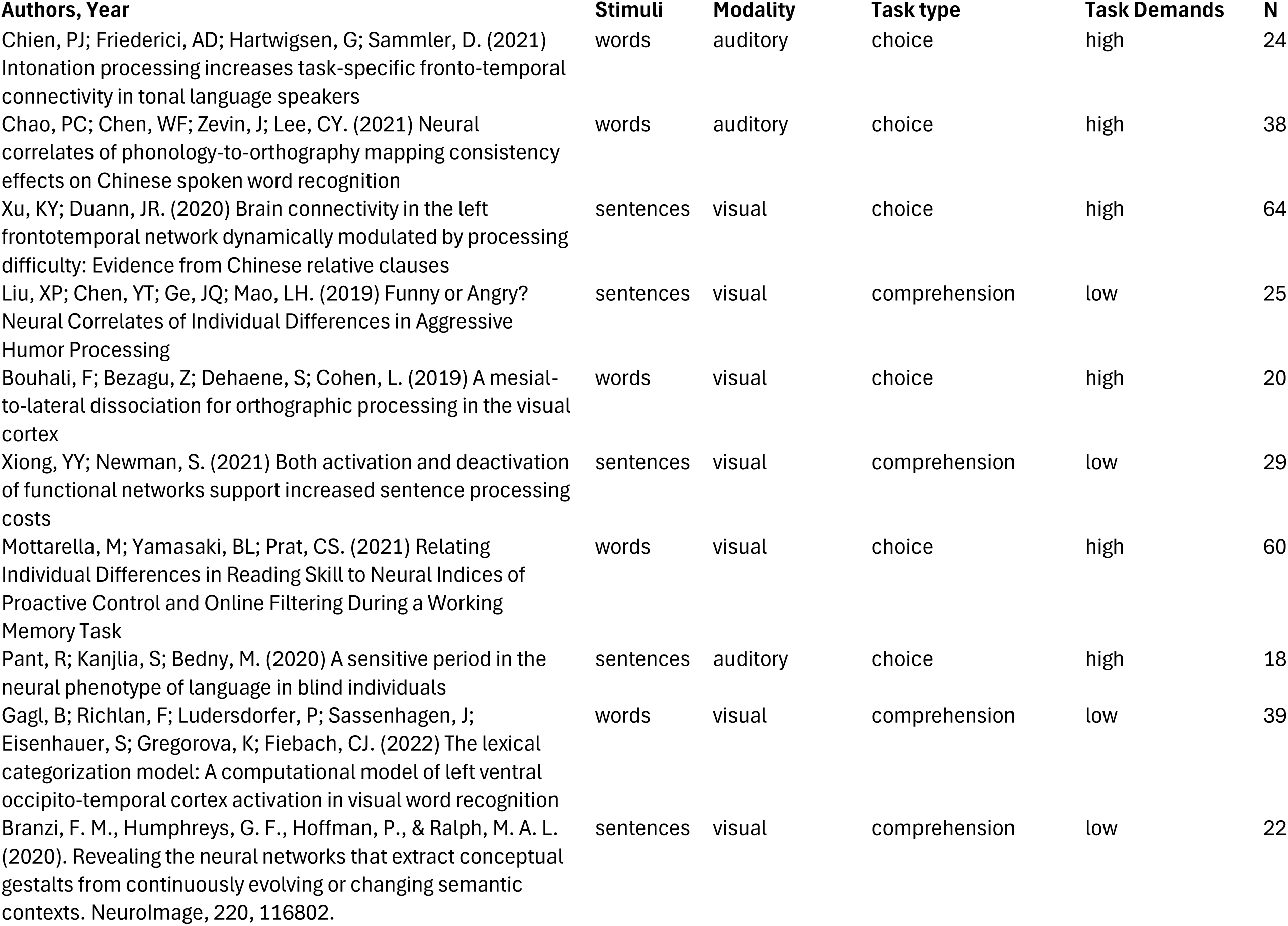
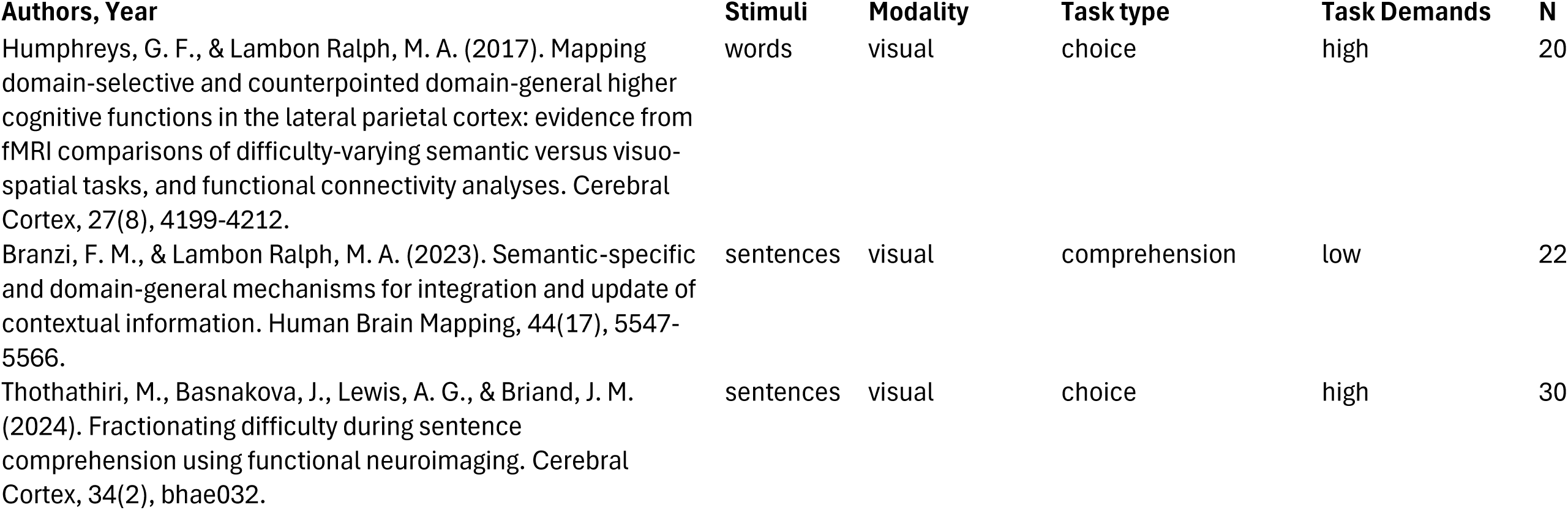
List of studies included in the Semantic Control meta-analysis. Studies have been categorised based on type of stimuli (Single words/Word pairs or Sentences/Narratives); modality (visual or auditory or both), type of task (choice or comprehension or production/internal generation (thinking about the right answer)); and task demands (high task demands or low task demands). Number of participants is provided (N).

**Supplementary Table 3.**
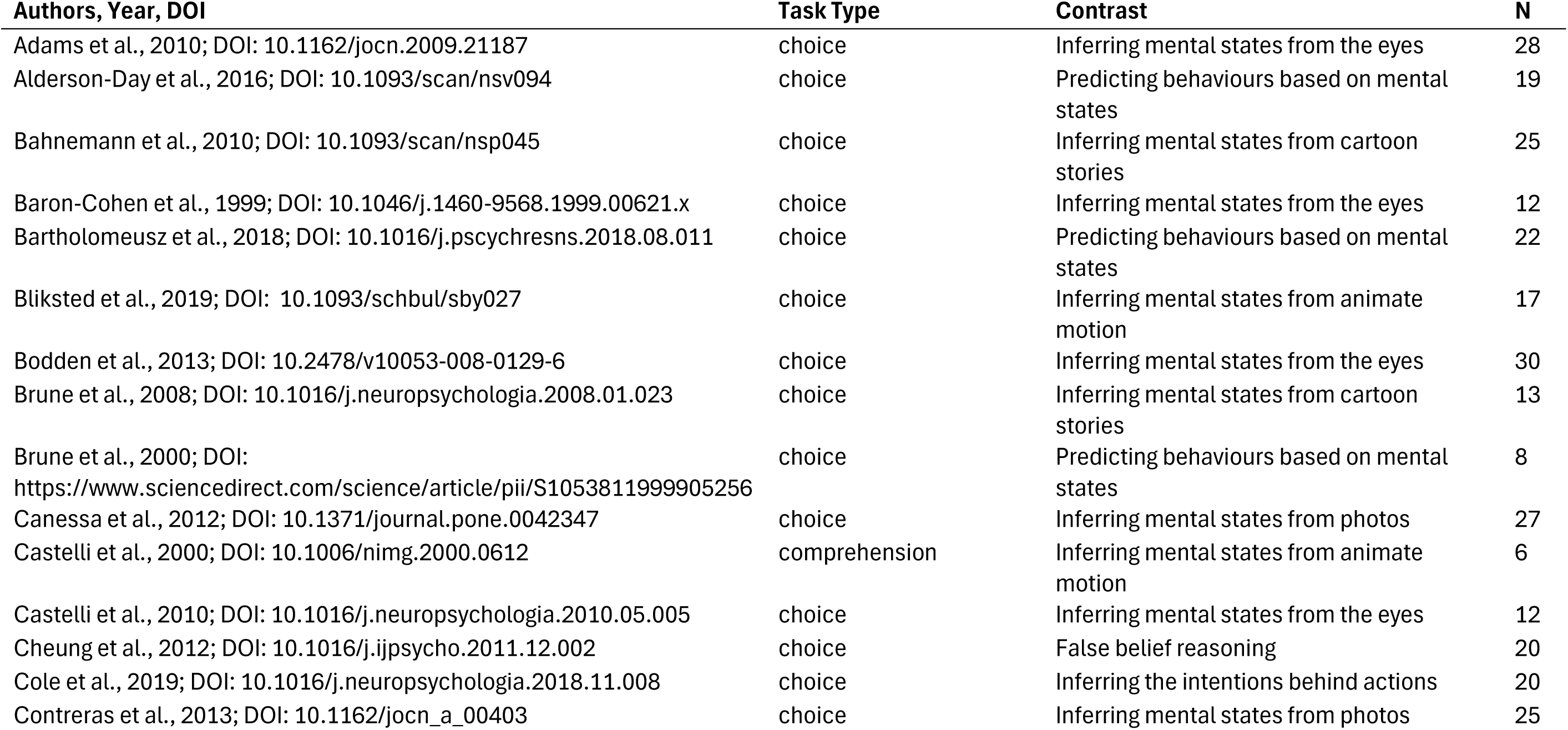
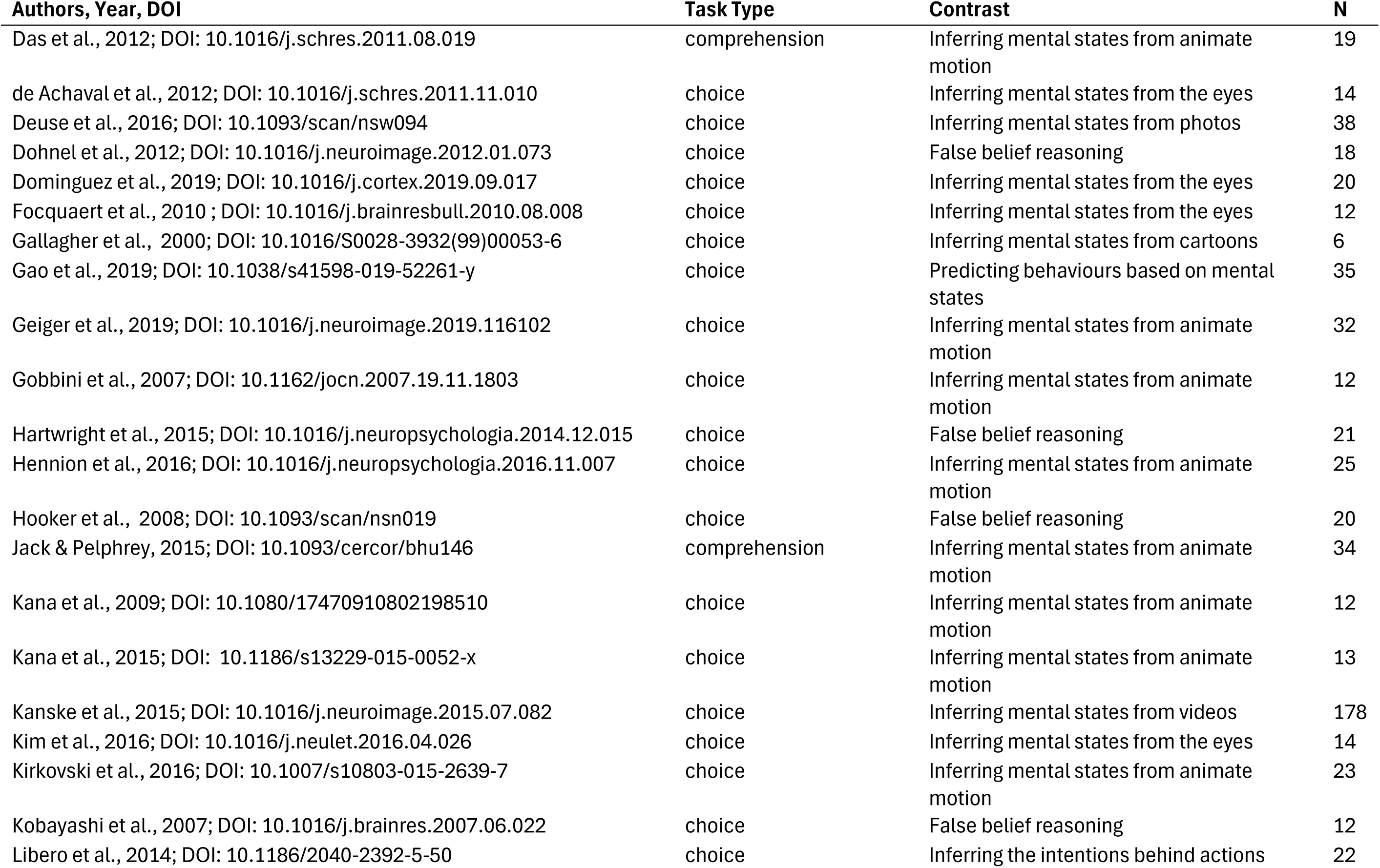
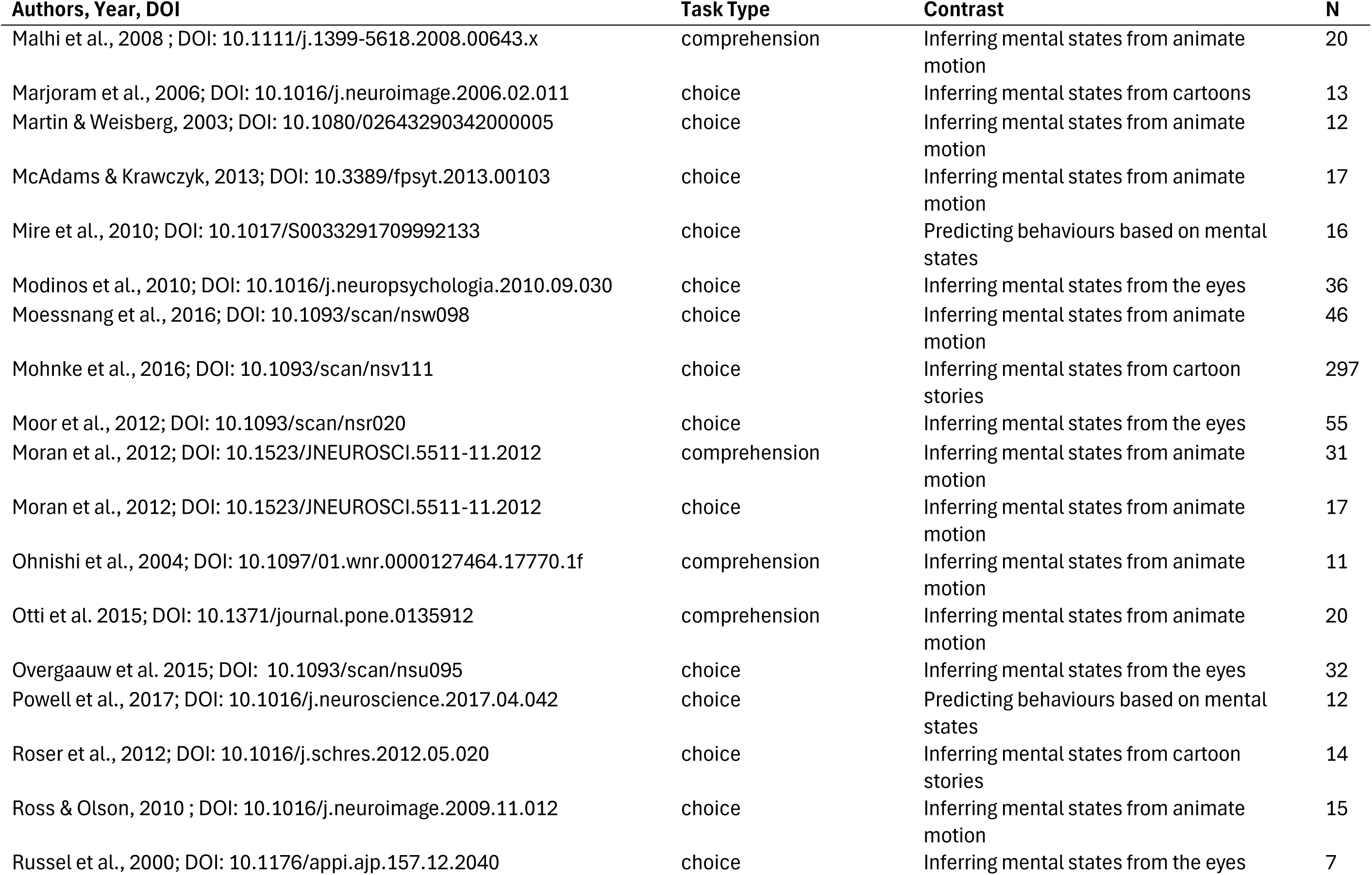
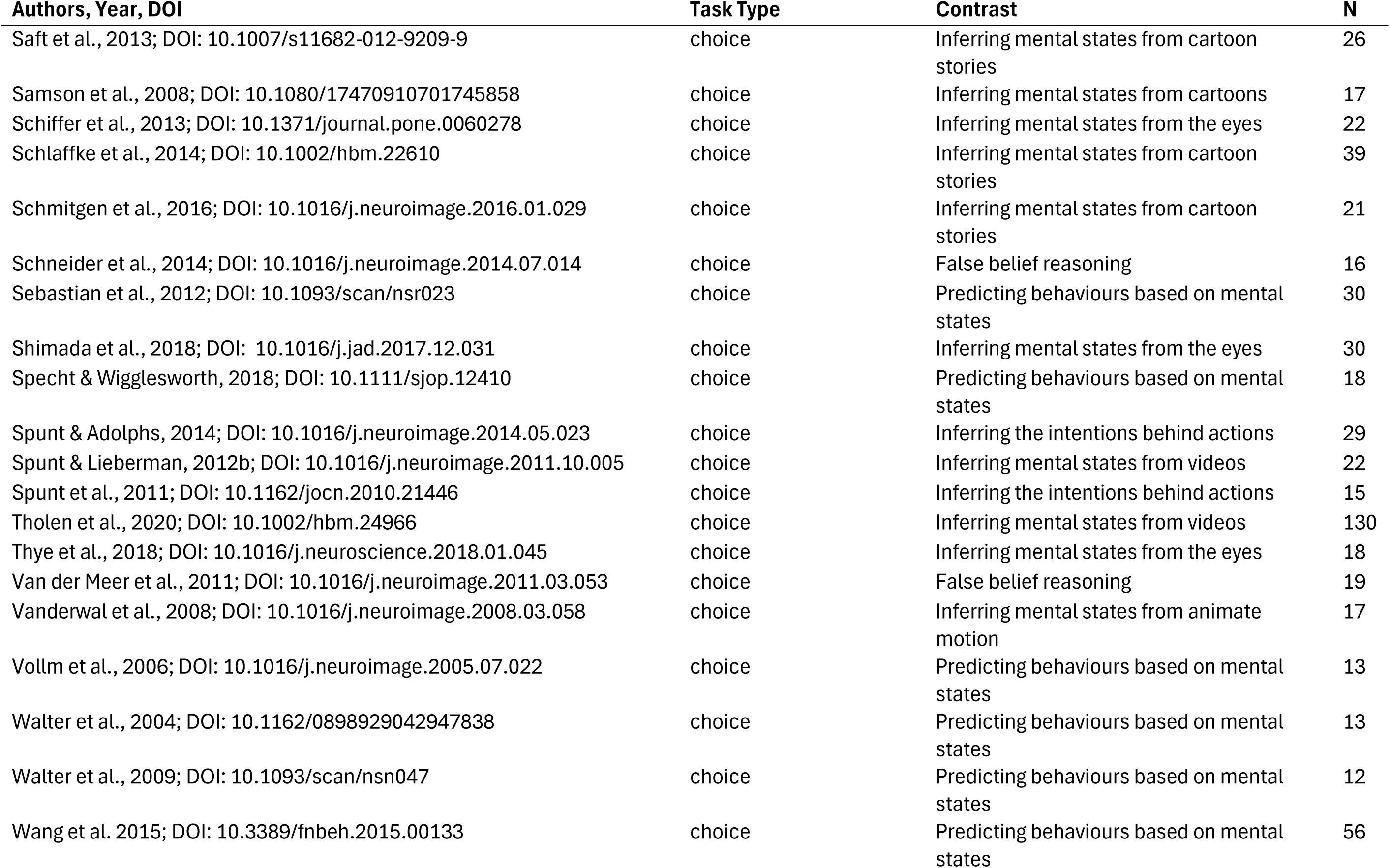
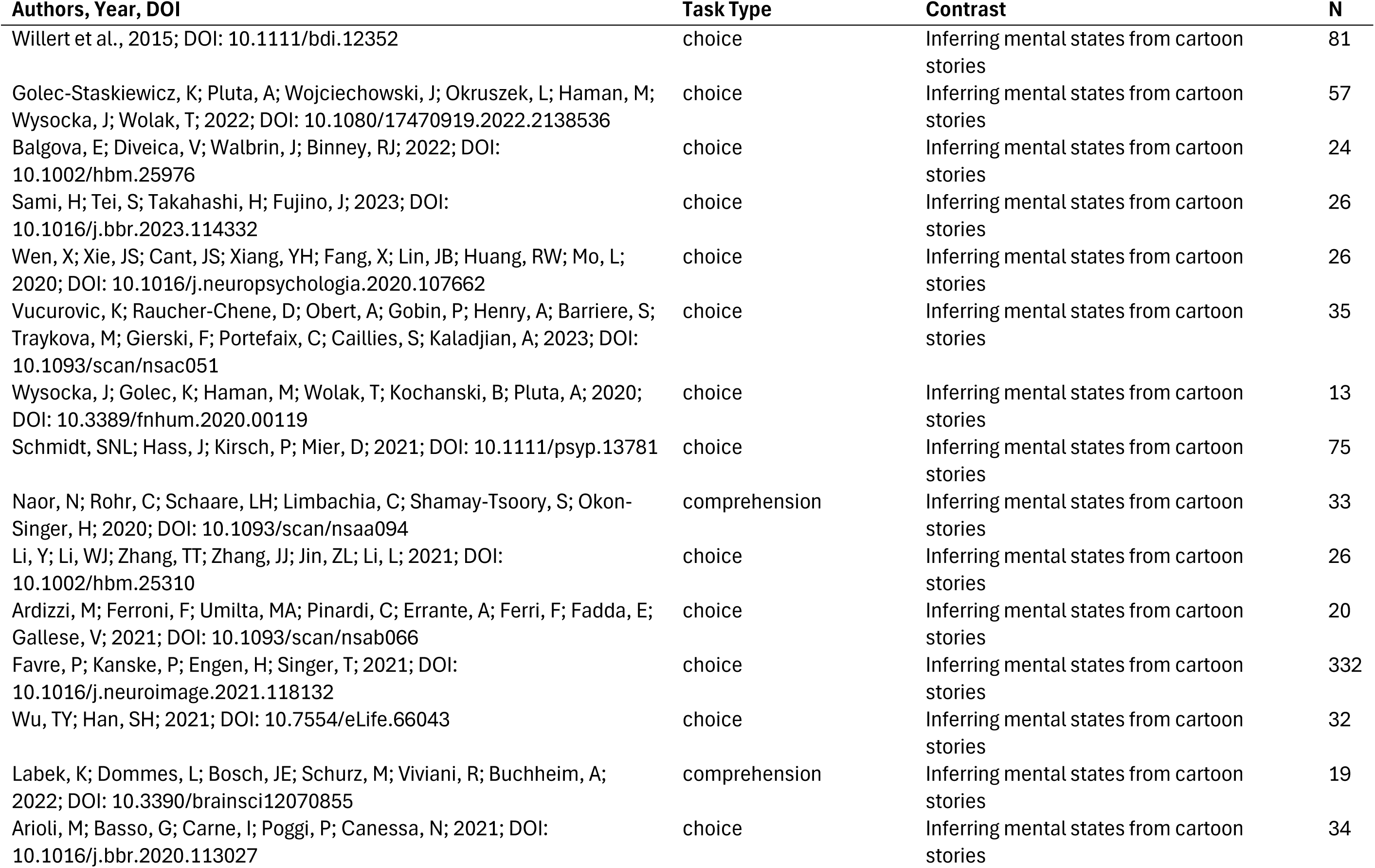
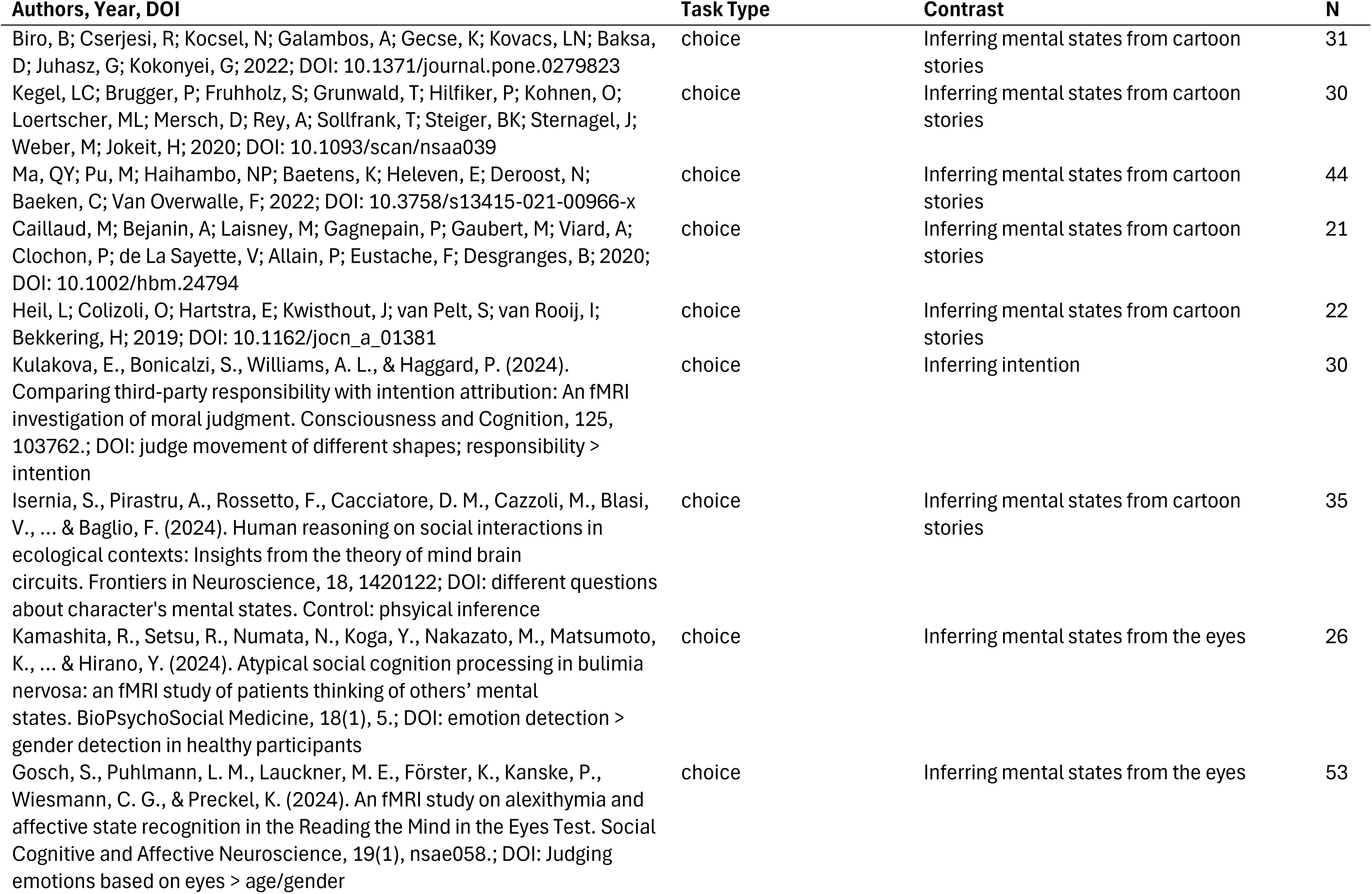
List of studies included in the **Nonverbal Theory of Mind** meta-analysis. Studies have been categorised based **type of task** (choice or comprehension); and the type of Theory of Mind required to perform (**contrast** - inferring intention or mental state, or predicting behaviour, or false belief reasoning). Number of participants is provided (N).

**Supplementary Table 4.**
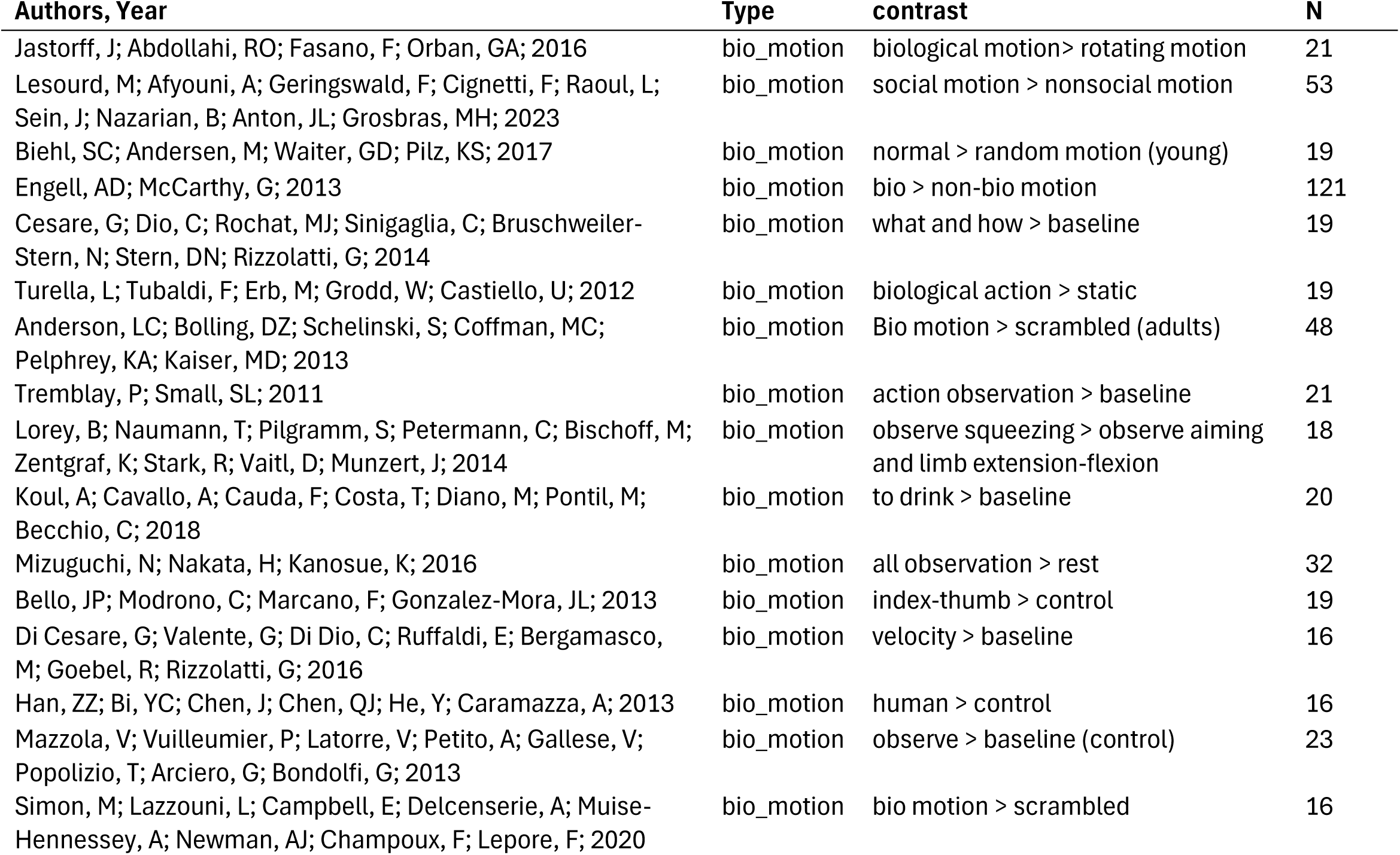
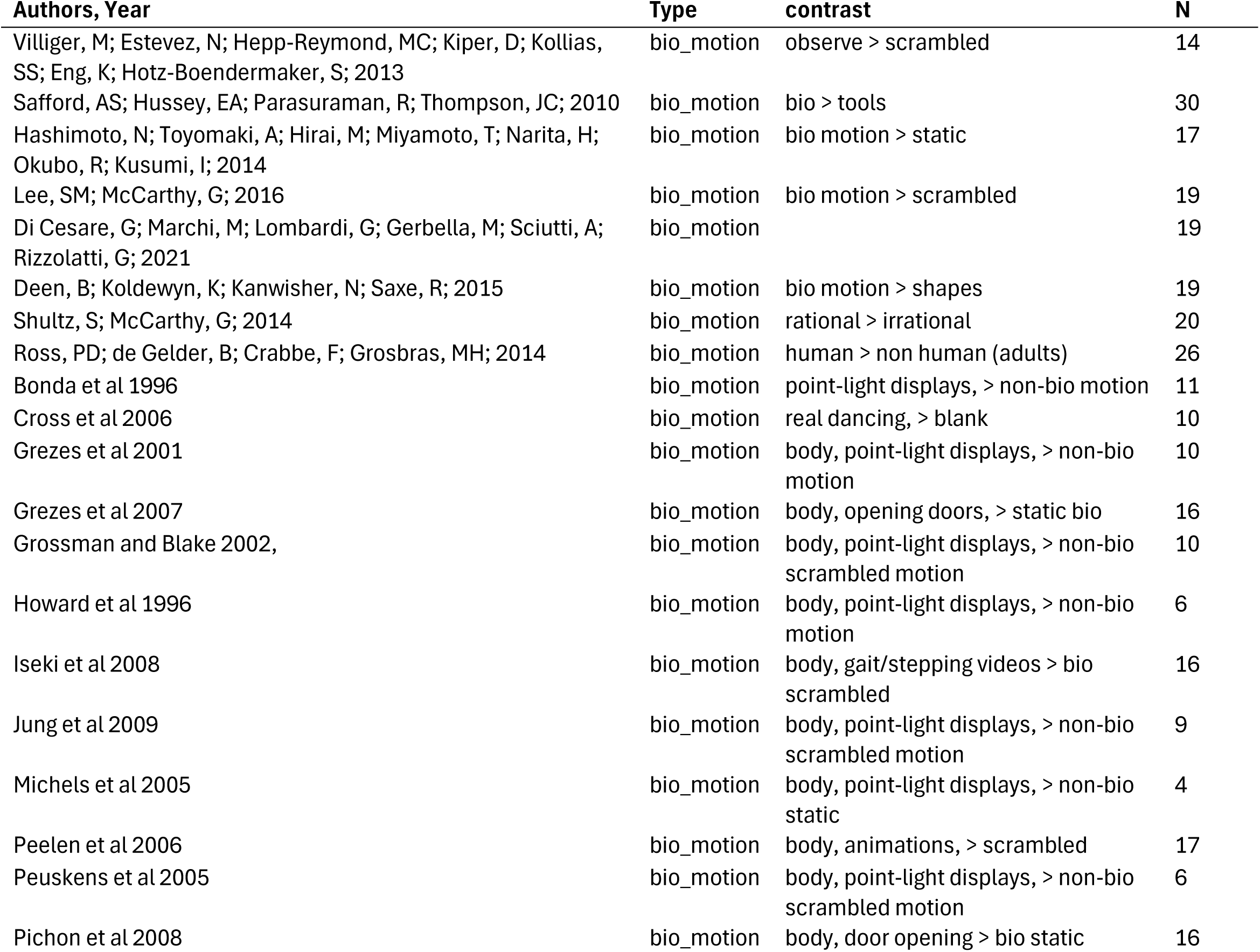
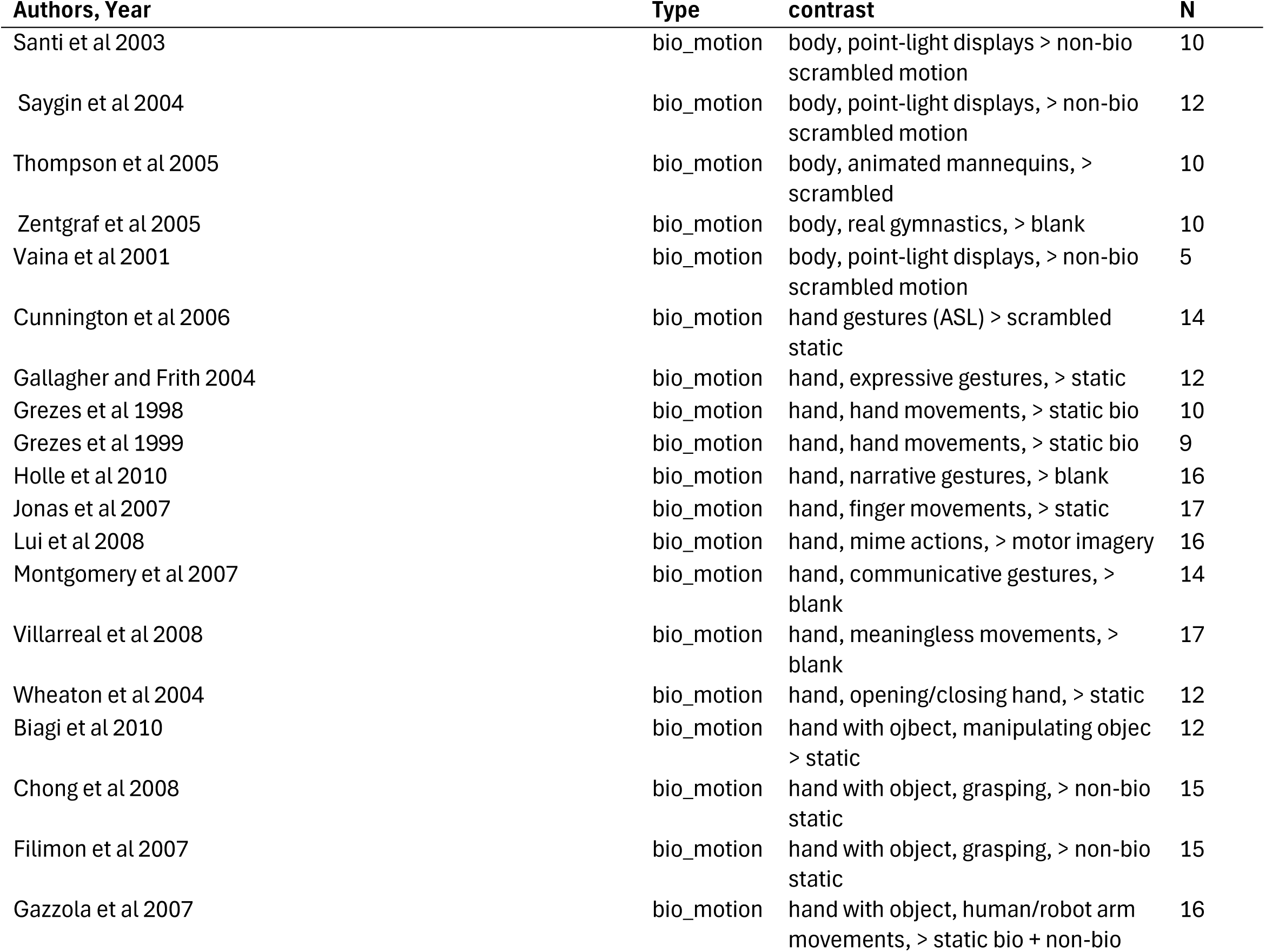
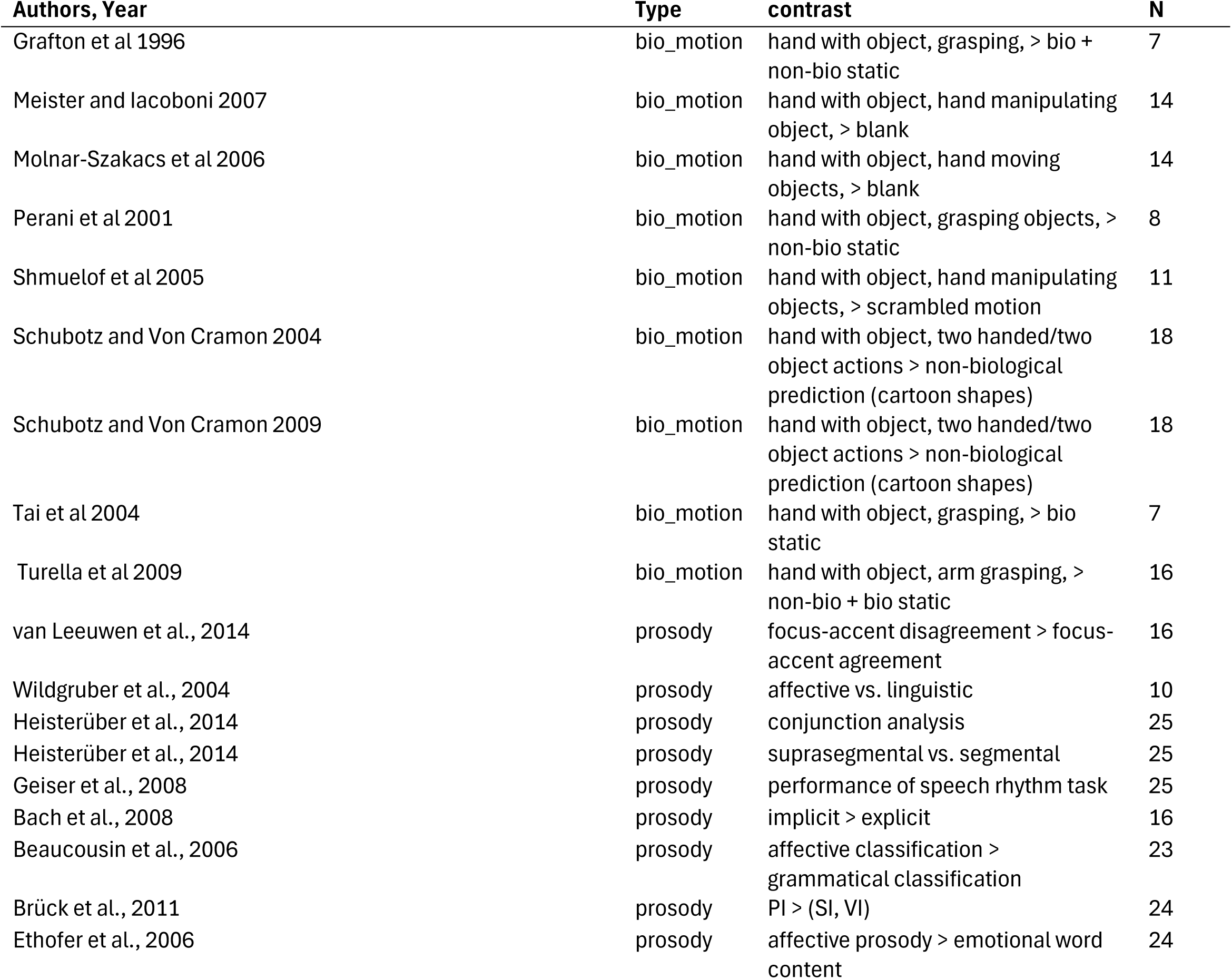
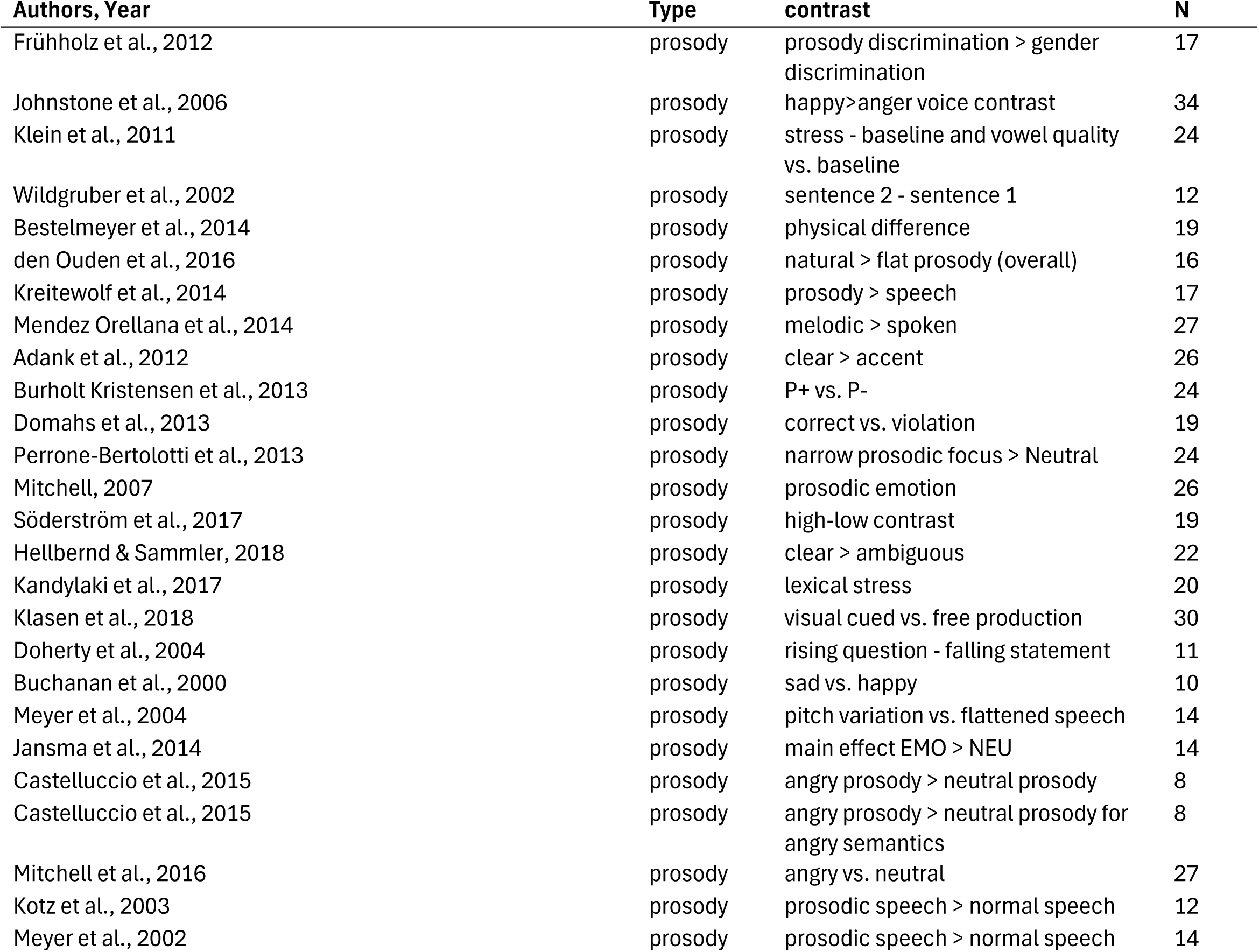
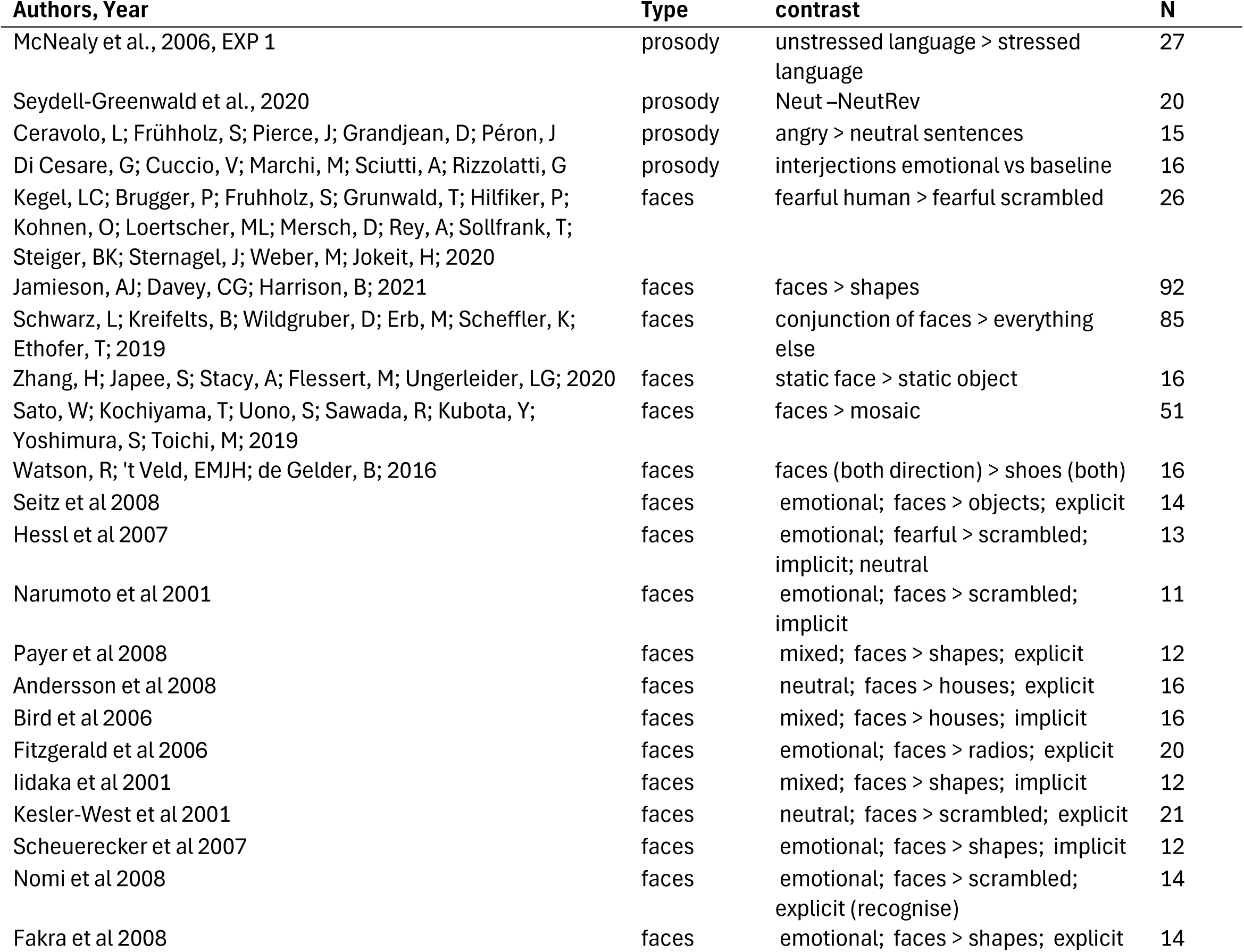
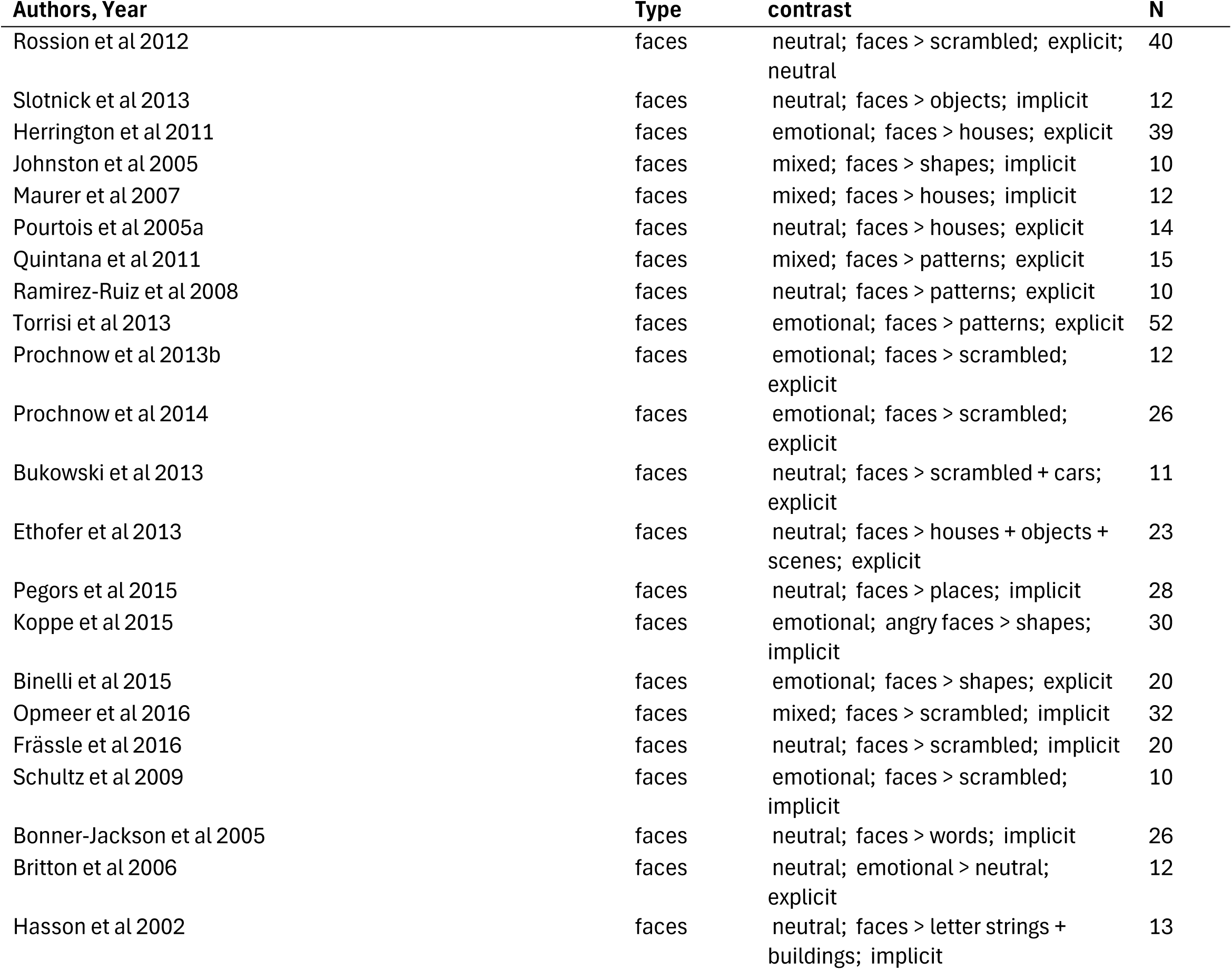
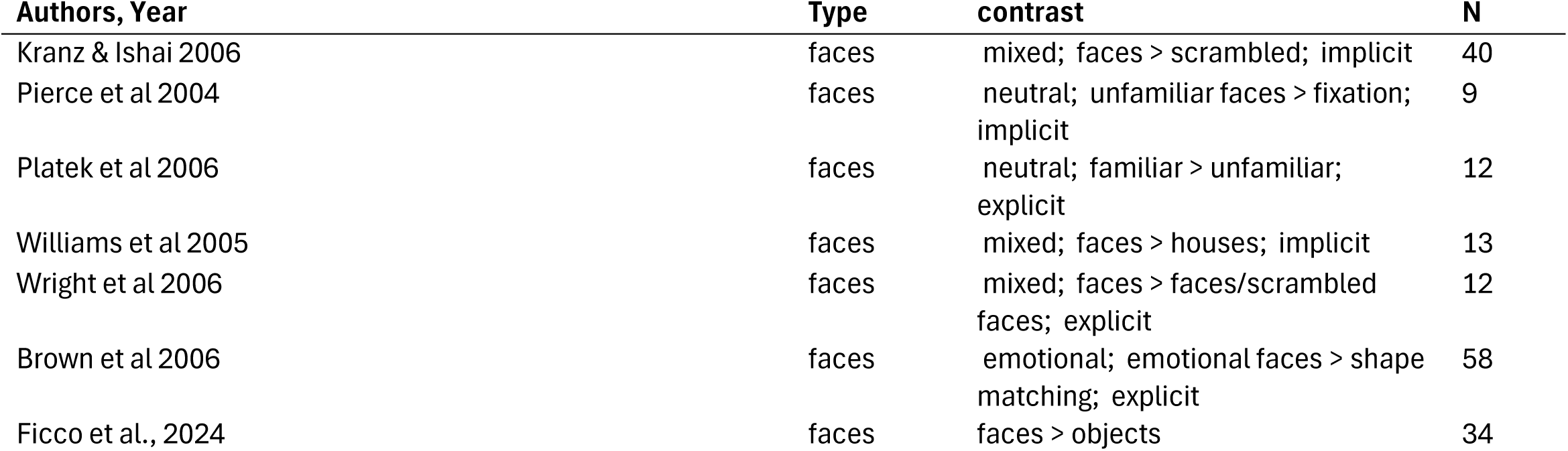
List of studies included in the Social Cues meta-analysis. Studies have been categorised based on type of **social cue** (biological motion or face perception or prosody). The **contrast** taken from each study is specified. Number of participants is provided (N).

| Authors, Year | Type | contrast | N |
| --- | --- | --- | --- |
| Jastorff, J; Abdollahi, RO; Fasano, F; Orban, GA; 2016 | bio_motion | biological motion> rotating motion | 21 |
| Lesourd, M; Afyouni, A; Geringswald, F; Cignetti, F; Raoul, L; Sein, J; Nazarian, B; Anton, JL; Grosbras, MH; 2023 | bio_motion | social motion > nonsocial motion | 53 |
| Biehl, SC; Andersen, M; Waiter, GD; Pilz, KS; 2017 | bio_motion | normal > random motion (young) | 19 |
| Engell, AD; McCarthy, G; 2013 | bio_motion | bio > non-bio motion | 121 |
| Cesare, G; Dio, C; Rochat, MJ; Sinigaglia, C; Bruschweiler-Stern, N; Stern, DN; Rizzolatti, G; 2014 | bio_motion | what and how > baseline | 19 |
| Turella, L; Tubaldi, F; Erb, M; Grodd, W; Castiello, U; 2012 | bio_motion | biological action > static | 19 |
| Anderson, LC; Bolling, DZ; Schelinski, S; Coffman, MC; Pelphrey, KA; Kaiser, MD; 2013 | bio_motion | Bio motion > scrambled (adults) | 48 |
| Tremblay, P; Small, SL; 2011 | bio_motion | action observation > baseline | 21 |
| Lorey, B; Naumann, T; Pilgramm, S; Petermann, C; Bischoff, M; Zentgraf, K; Stark, R; Vaitl, D; Munzert, J; 2014 | bio_motion | observe squeezing > observe aiming and limb extension-flexion | 18 |
| Koul, A; Cavallo, A; Cauda, F; Costa, T; Diano, M; Pontil, M; Becchio, C; 2018 | bio_motion | to drink > baseline | 20 |
| Mizuguchi, N; Nakata, H; Kanosue, K; 2016 | bio_motion | all observation > rest | 32 |
| Bello, JP; Modrono, C; Marcano, F; Gonzalez-Mora, JL; 2013 | bio_motion | index-thumb > control | 19 |
| Di Cesare, G; Valente, G; Di Dio, C; Ruffaldi, E; Bergamasco, M; Goebel, R; Rizzolatti, G; 2016 | bio_motion | velocity > baseline | 16 |
| Han, ZZ; Bi, YC; Chen, J; Chen, QJ; He, Y; Caramazza, A; 2013 | bio_motion | human > control | 16 |
| Mazzola, V; Vuilleumier, P; Latorre, V; Petito, A; Gallese, V; Popolizio, T; Arciero, G; Bondolfi, G; 2013 | bio_motion | observe > baseline (control) | 23 |
| Simon, M; Lazzouni, L; Campbell, E; Delcenserie, A; Muise-Hennessey, A; Newman, AJ; Champoux, F; Lepore, F; 2020 | bio_motion | bio motion > scrambled | 16 |
| Villiger, M; Estevez, N; Hepp-Reymond, MC; Kiper, D; Kollias, SS; Eng, K; Hotz-Boendermaker, S; 2013 | bio_motion | observe > scrambled | 14 |
| Safford, AS; Hussey, EA; Parasuraman, R; Thompson, JC; 2010 | bio_motion | bio > tools | 30 |
| Hashimoto, N; Toyomaki, A; Hirai, M; Miyamoto, T; Narita, H; Okubo, R; Kusumi, I; 2014 | bio_motion | bio motion > static | 17 |
| Lee, SM; McCarthy, G; 2016 | bio_motion | bio motion > scrambled | 19 |
| Di Cesare, G; Marchi, M; Lombardi, G; Gerbella, M; Sciutti, A; Rizzolatti, G; 2021 | bio_motion |  | 19 |
| Deen, B; Koldewyn, K; Kanwisher, N; Saxe, R; 2015 | bio_motion | bio motion > shapes | 19 |
| Shultz, S; McCarthy, G; 2014 | bio_motion | rational > irrational | 20 |
| Ross, PD; de Gelder, B; Crabbe, F; Grosbras, MH; 2014 | bio_motion | human > non human (adults) | 26 |
| Bonda et al 1996 | bio_motion | point-light displays, > non-bio motion | 11 |
| Cross et al 2006 | bio_motion | real dancing, > blank | 10 |
| Grezes et al 2001 | bio_motion | body, point-light displays, > non-bio motion | 10 |
| Grezes et al 2007 | bio_motion | body, opening doors, > static bio | 16 |
| Grossman and Blake 2002, | bio_motion | body, point-light displays, > non-bio scrambled motion | 10 |
| Howard et al 1996 | bio_motion | body, point-light displays, > non-bio motion | 6 |
| Iseki et al 2008 | bio_motion | body, gait/stepping videos > bio scrambled | 16 |
| Jung et al 2009 | bio_motion | body, point-light displays, > non-bio scrambled motion | 9 |
| Michels et al 2005 | bio_motion | body, point-light displays, > non-bio static | 4 |
| Peelen et al 2006 | bio_motion | body, animations, > scrambled | 17 |
| Peuskens et al 2005 | bio_motion | body, point-light displays, > non-bio scrambled motion | 6 |
| Pichon et al 2008 | bio_motion | body, door opening > bio static | 16 |
| Santi et al 2003 | bio_motion | body, point-light displays > non-bio scrambled motion | 10 |
| Saygin et al 2004 | bio_motion | body, point-light displays, > non-bio scrambled motion | 12 |
| Thompson et al 2005 | bio_motion | body, animated mannequins, > scrambled | 10 |
| Zentgraf et al 2005 | bio_motion | body, real gymnastics, > blank | 10 |
| Vaina et al 2001 | bio_motion | body, point-light displays, > non-bio scrambled motion | 5 |
| Cunnington et al 2006 | bio_motion | hand gestures (ASL) > scrambled static | 14 |
| Gallagher and Frith 2004 | bio_motion | hand, expressive gestures, > static | 12 |
| Grezes et al 1998 | bio_motion | hand, hand movements, > static bio | 10 |
| Grezes et al 1999 | bio_motion | hand, hand movements, > static bio | 9 |
| Holle et al 2010 | bio_motion | hand, narrative gestures, > blank | 16 |
| Jonas et al 2007 | bio_motion | hand, finger movements, > static | 17 |
| Lui et al 2008 | bio_motion | hand, mime actions, > motor imagery | 16 |
| Montgomery et al 2007 | bio_motion | hand, communicative gestures, > blank | 14 |
| Villarreal et al 2008 | bio_motion | hand, meaningless movements, > blank | 17 |
| Wheaton et al 2004 | bio_motion | hand, opening/closing hand, > static | 12 |
| Biagi et al 2010 | bio_motion | hand with object, manipulating object > static | 12 |
| Chong et al 2008 | bio_motion | hand with object, grasping, > non-bio static | 15 |
| Filimon et al 2007 | bio_motion | hand with object, grasping, > non-bio static | 15 |
| Gazzola et al 2007 | bio_motion | hand with object, human/robot arm movements, > static bio + non-bio | 16 |
| Grafton et al 1996 | bio_motion | hand with object, grasping, > bio + non-bio static | 7 |
| Meister and Iacoboni 2007 | bio_motion | hand with object, hand manipulating object, > blank | 14 |
| Molnar-Szakacs et al 2006 | bio_motion | hand with object, hand moving objects, > blank | 14 |
| Perani et al 2001 | bio_motion | hand with object, grasping objects, > non-bio static | 8 |
| Shmuelof et al 2005 | bio_motion | hand with object, hand manipulating objects, > scrambled motion | 11 |
| Schubotz and Von Cramon 2004 | bio_motion | hand with object, two handed/two object actions > non-biological prediction (cartoon shapes) | 18 |
| Schubotz and Von Cramon 2009 | bio_motion | hand with object, two handed/two object actions > non-biological prediction (cartoon shapes) | 18 |
| Tai et al 2004 | bio_motion | hand with object, grasping, > bio static | 7 |
| Turella et al 2009 | bio_motion | hand with object, arm grasping, > non-bio + bio static | 16 |
| van Leeuwen et al., 2014 | prosody | focus-accent disagreement > focus-accent agreement | 16 |
| Wildgruber et al., 2004 | prosody | affective vs. linguistic | 10 |
| Heisterüber et al., 2014 | prosody | conjunction analysis | 25 |
| Heisterüber et al., 2014 | prosody | suprasegmental vs. segmental | 25 |
| Geiser et al., 2008 | prosody | performance of speech rhythm task | 25 |
| Bach et al., 2008 | prosody | implicit > explicit | 16 |
| Beaucousin et al., 2006 | prosody | affective classification > grammatical classification | 23 |
| Brück et al., 2011 | prosody | PI > (SI, VI) | 24 |
| Ethofer et al., 2006 | prosody | affective prosody > emotional word content | 24 |
| Frühholz et al., 2012 | prosody | prosody discrimination > gender discrimination | 17 |
| Johnstone et al., 2006 | prosody | happy>anger voice contrast | 34 |
| Klein et al., 2011 | prosody | stress - baseline and vowel quality vs. baseline | 24 |
| Wildgruber et al., 2002 | prosody | sentence 2 - sentence 1 | 12 |
| Bestelmeyer et al., 2014 | prosody | physical difference | 19 |
| den Ouden et al., 2016 | prosody | natural > flat prosody (overall) | 16 |
| Kreitewolf et al., 2014 | prosody | prosody > speech | 17 |
| Mendez Orellana et al., 2014 | prosody | melodic > spoken | 27 |
| Adank et al., 2012 | prosody | clear > accent | 26 |
| Burholt Kristensen et al., 2013 | prosody | P+ vs. P- | 24 |
| Domahs et al., 2013 | prosody | correct vs. violation | 19 |
| Perrone-Bertolotti et al., 2013 | prosody | narrow prosodic focus > Neutral | 24 |
| Mitchell, 2007 | prosody | prosodic emotion | 26 |
| Söderström et al., 2017 | prosody | high-low contrast | 19 |
| Hellbernd & Sammler, 2018 | prosody | clear > ambiguous | 22 |
| Kandylaki et al., 2017 | prosody | lexical stress | 20 |
| Klasen et al., 2018 | prosody | visual cued vs. free production | 30 |
| Doherty et al., 2004 | prosody | rising question - falling statement | 11 |
| Buchanan et al., 2000 | prosody | sad vs. happy | 10 |
| Meyer et al., 2004 | prosody | pitch variation vs. flattened speech | 14 |
| Jansma et al., 2014 | prosody | main effect EMO > NEU | 14 |
| Castelluccio et al., 2015 | prosody | angry prosody > neutral prosody | 8 |
| Castelluccio et al., 2015 | prosody | angry prosody > neutral prosody for angry semantics | 8 |
| Mitchell et al., 2016 | prosody | angry vs. neutral | 27 |
| Kotz et al., 2003 | prosody | prosodic speech > normal speech | 12 |
| Meyer et al., 2002 | prosody | prosodic speech > normal speech | 14 |
| McNealy et al., 2006, EXP 1 | prosody | unstressed language > stressed language | 27 |
| Seydell-Greenwald et al., 2020 | prosody | Neut –NeutRev | 20 |
| Ceravolo, L; Frühholz, S; Pierce, J; Grandjean, D; Péron, J | prosody | angry > neutral sentences | 15 |
| Di Cesare, G; Cuccio, V; Marchi, M; Sciutti, A; Rizzolatti, G | prosody | interjections emotional vs baseline | 16 |
| Kegel, LC; Brugger, P; Fruhhholz, S; Grunwald, T; Hilfiker, P; Kohnen, O; Loertscher, ML; Mersch, D; Rey, A; Sollfrank, T; Steiger, BK; Sternagel, J; Weber, M; Jokeit, H; 2020 | faces | fearful human > fearful scrambled | 26 |
| Jamieson, AJ; Davey, CG; Harrison, B; 2021 | faces | faces > shapes | 92 |
| Schwarz, L; Kreifelts, B; Wildgruber, D; Erb, M; Scheffler, K; Ethofer, T; 2019 | faces | conjunction of faces > everything else | 85 |
| Zhang, H; Japee, S; Stacy, A; Flessert, M; Ungerleider, LG; 2020 | faces | static face > static object | 16 |
| Sato, W; Kochiyama, T; Uono, S; Sawada, R; Kubota, Y; Yoshimura, S; Toichi, M; 2019 | faces | faces > mosaic | 51 |
| Watson, R; 't Veld, EMJH; de Gelder, B; 2016 | faces | faces (both direction) > shoes (both) | 16 |
| Seitz et al 2008 | faces | emotional; faces > objects; explicit | 14 |
| Hessl et al 2007 | faces | emotional; fearful > scrambled; implicit; neutral | 13 |
| Narumoto et al 2001 | faces | emotional; faces > scrambled; implicit | 11 |
| Payer et al 2008 | faces | mixed; faces > shapes; explicit | 12 |
| Andersson et al 2008 | faces | neutral; faces > houses; explicit | 16 |
| Bird et al 2006 | faces | mixed; faces > houses; implicit | 16 |
| Fitzgerald et al 2006 | faces | emotional; faces > radios; explicit | 20 |
| Iidaka et al 2001 | faces | mixed; faces > shapes; implicit | 12 |
| Kesler-West et al 2001 | faces | neutral; faces > scrambled; explicit | 21 |
| Scheuerecker et al 2007 | faces | emotional; faces > shapes; implicit | 12 |
| Nomi et al 2008 | faces | emotional; faces > scrambled; explicit (recognise) | 14 |
| Fakra et al 2008 | faces | emotional; faces > shapes; explicit | 14 |

| **Authors, Year** | **Type** | **contrast** | **N** |
| --- | --- | --- | --- |
| Rossion et al 2012 | faces | neutral; faces > scrambled; explicit; neutral | 40 |
| Slotnick et al 2013 | faces | neutral; faces > objects; implicit | 12 |
| Herrington et al 2011 | faces | emotional; faces > houses; explicit | 39 |
| Johnston et al 2005 | faces | mixed; faces > shapes; implicit | 10 |
| Maurer et al 2007 | faces | mixed; faces > houses; implicit | 12 |
| Pourtois et al 2005a | faces | neutral; faces > houses; explicit | 14 |
| Quintana et al 2011 | faces | mixed; faces > patterns; explicit | 15 |
| Ramirez-Ruiz et al 2008 | faces | neutral; faces > patterns; explicit | 10 |
| Torrisi et al 2013 | faces | emotional; faces > patterns; explicit | 52 |
| Prochnow et al 2013b | faces | emotional; faces > scrambled; explicit | 12 |
| Prochnow et al 2014 | faces | emotional; faces > scrambled; explicit | 26 |
| Bukowski et al 2013 | faces | neutral; faces > scrambled + cars; explicit | 11 |
| Ethofer et al 2013 | faces | neutral; faces > houses + objects + scenes; explicit | 23 |
| Pegors et al 2015 | faces | neutral; faces > places; implicit | 28 |
| Koppe et al 2015 | faces | emotional; angry faces > shapes; implicit | 30 |
| Binelli et al 2015 | faces | emotional; faces > shapes; explicit | 20 |
| Opmeer et al 2016 | faces | mixed; faces > scrambled; implicit | 32 |
| Frässle et al 2016 | faces | neutral; faces > scrambled; implicit | 20 |
| Schultz et al 2009 | faces | emotional; faces > scrambled; implicit | 10 |
| Bonner-Jackson et al 2005 | faces | neutral; faces > words; implicit | 26 |
| Britton et al 2006 | faces | neutral; emotional > neutral; explicit | 12 |
| Hasson et al 2002 | faces | neutral; faces > letter strings + buildings; implicit | 13 |
| Kranz & Ishai 2006 | faces | mixed; faces > scrambled; implicit | 40 |
| Pierce et al 2004 | faces | neutral; unfamiliar faces > fixation; implicit | 9 |
| Platek et al 2006 | faces | neutral; familiar > unfamiliar; explicit | 12 |
| Williams et al 2005 | faces | mixed; faces > houses; implicit | 13 |
| Wright et al 2006 | faces | mixed; faces > faces/scrambled faces; explicit | 12 |
| Brown et al 2006 | faces | emotional; emotional faces > shape matching; explicit | 58 |
| Ficco et al., 2024 | faces | faces > objects | 34 |

**Supplementary Table 5.**
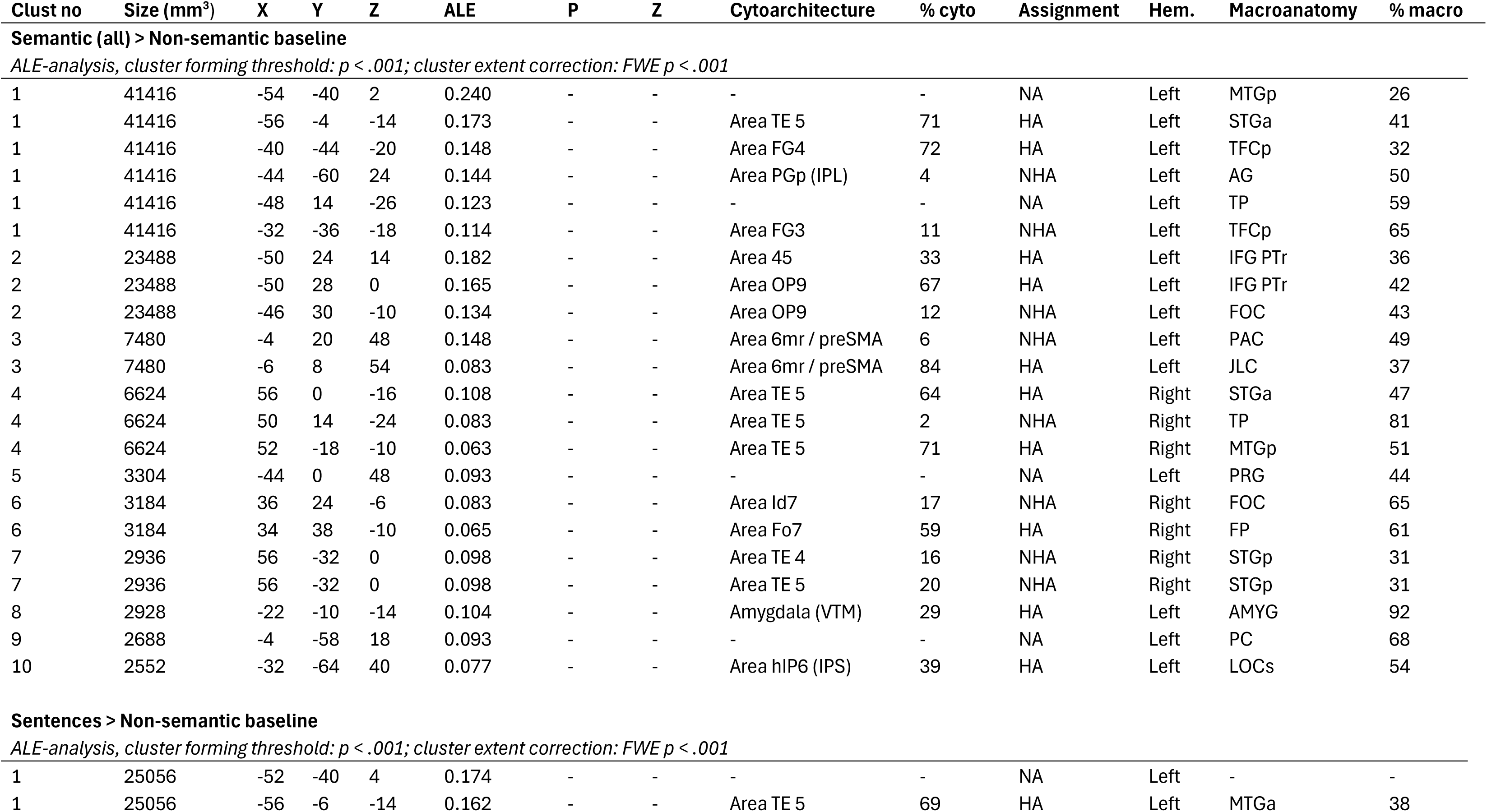
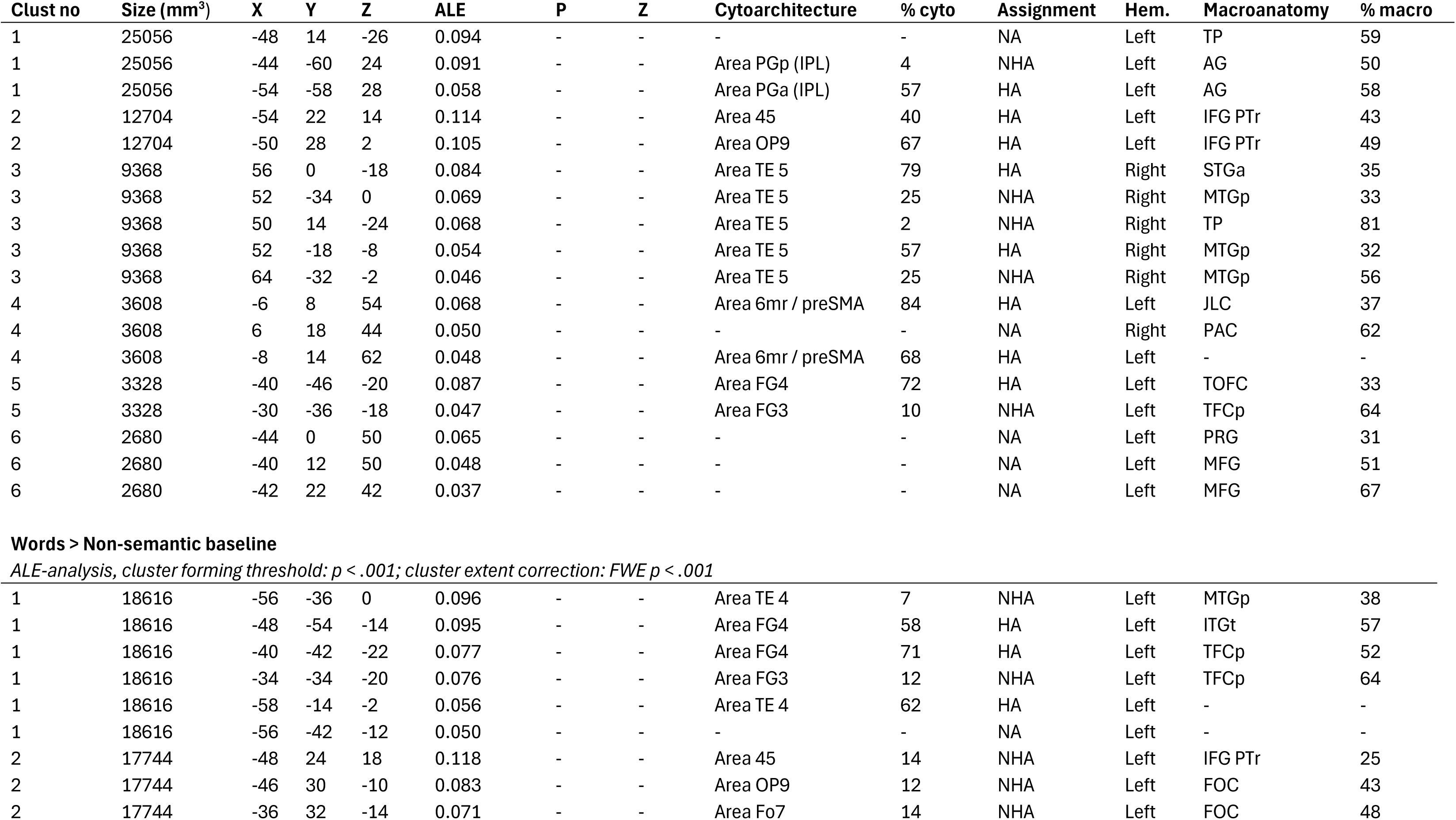
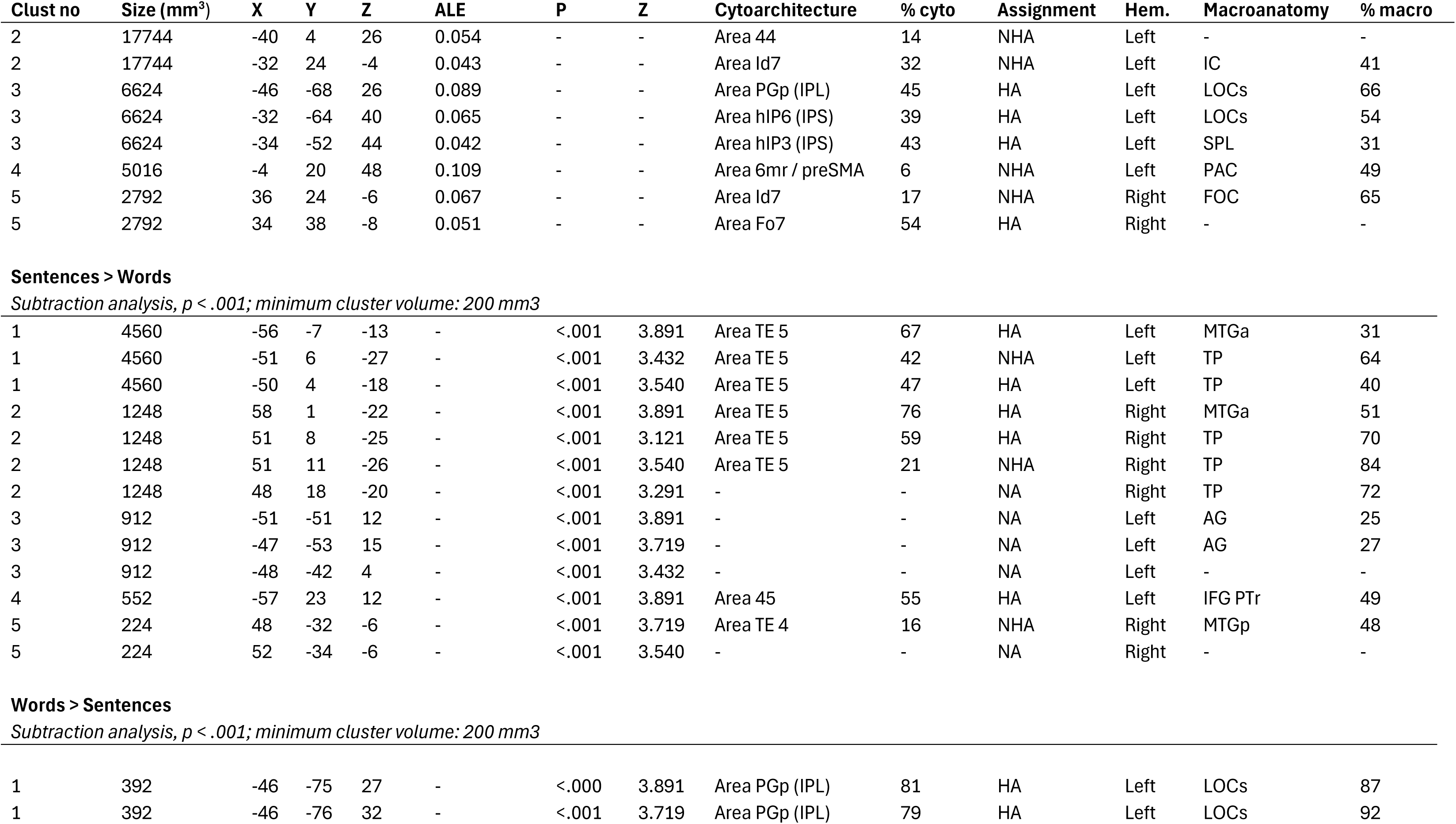
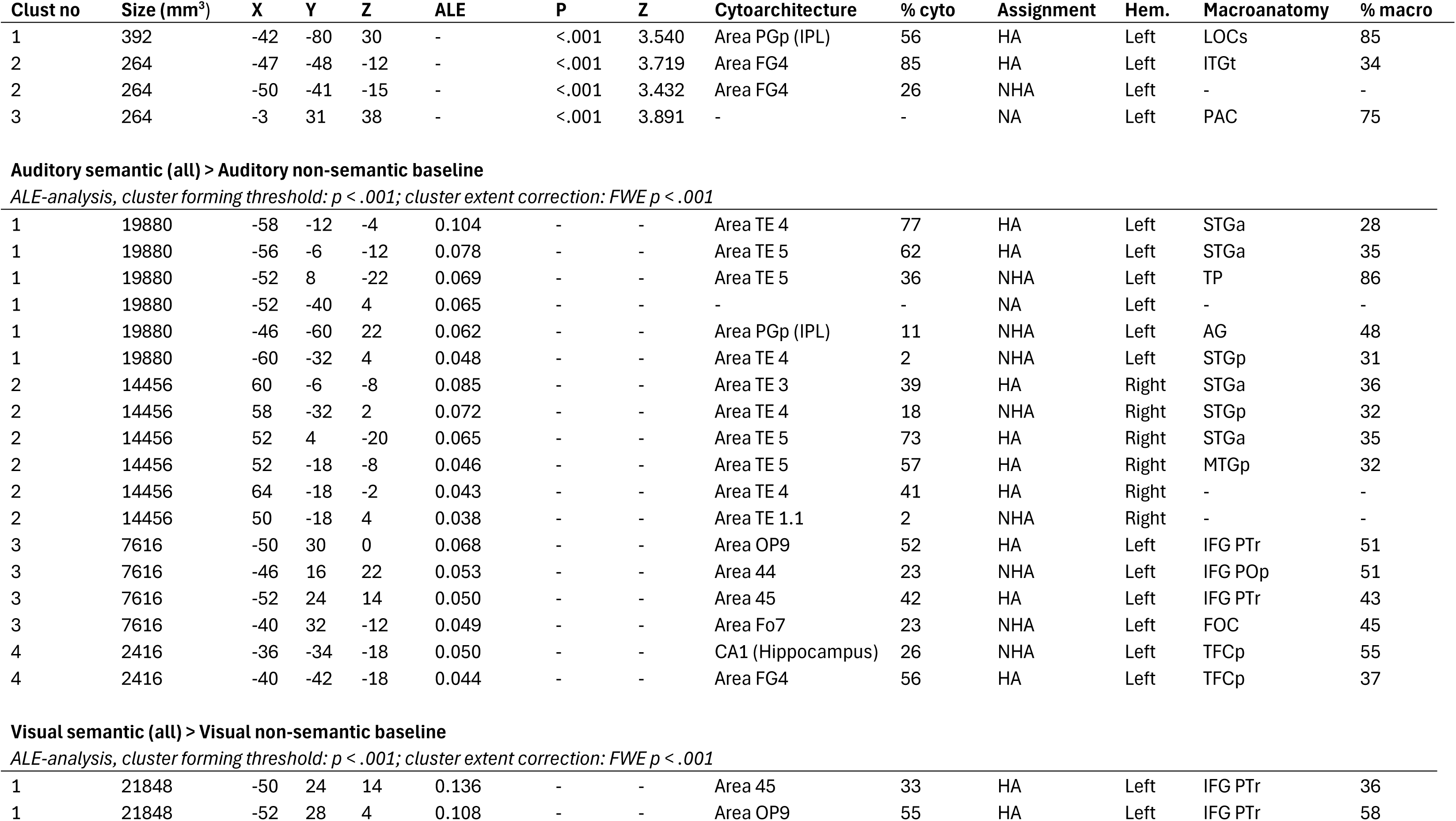
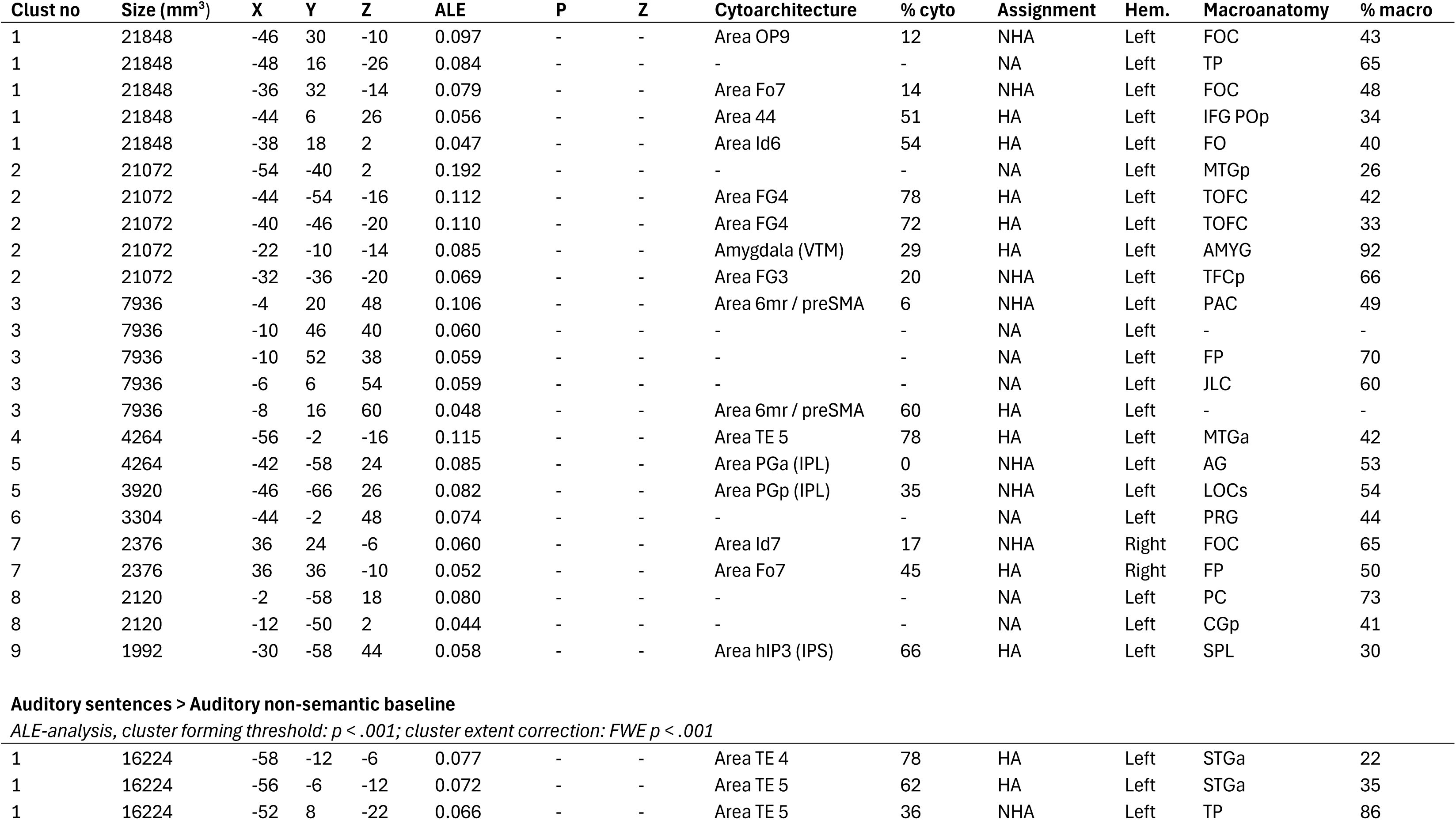
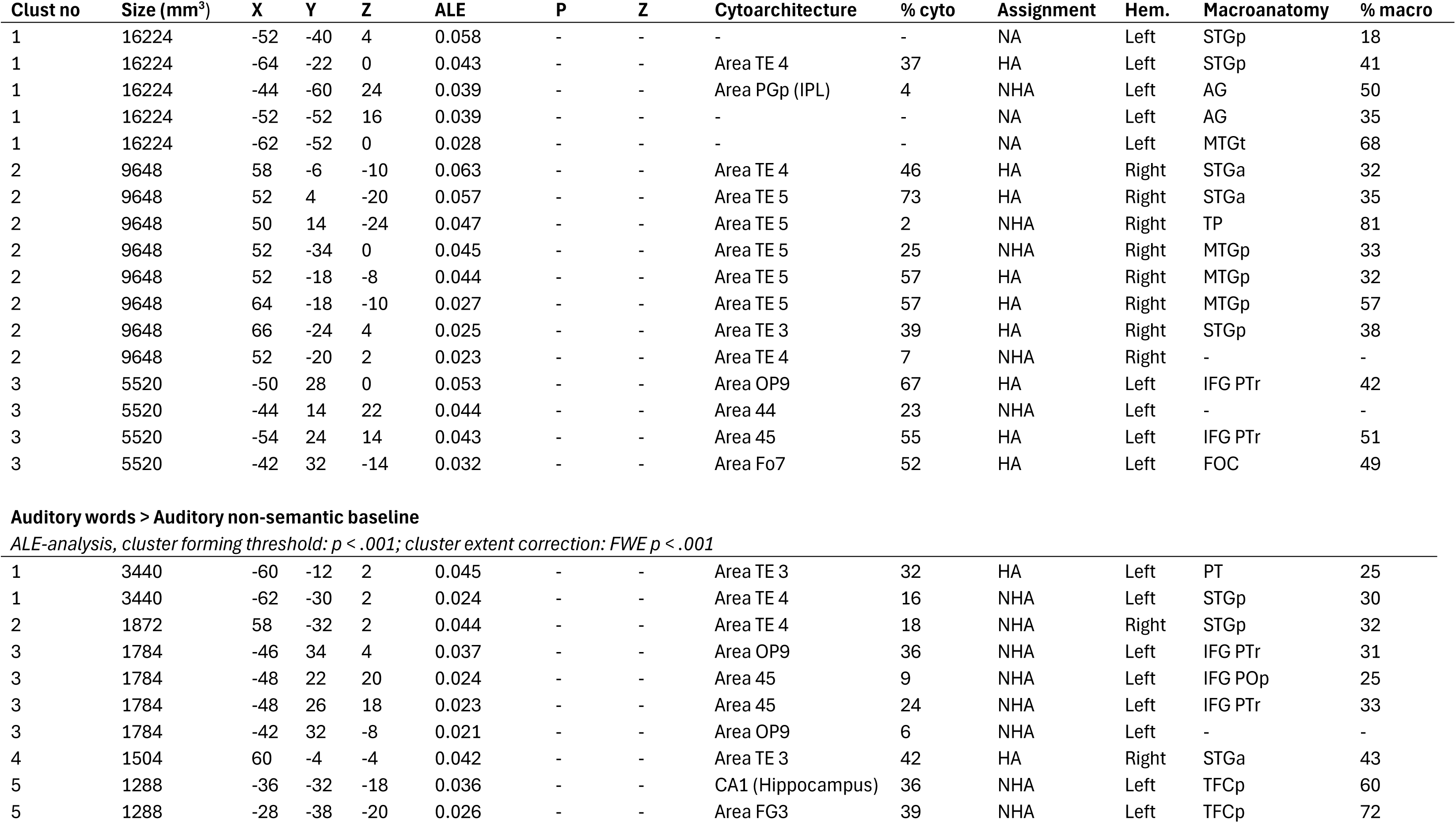
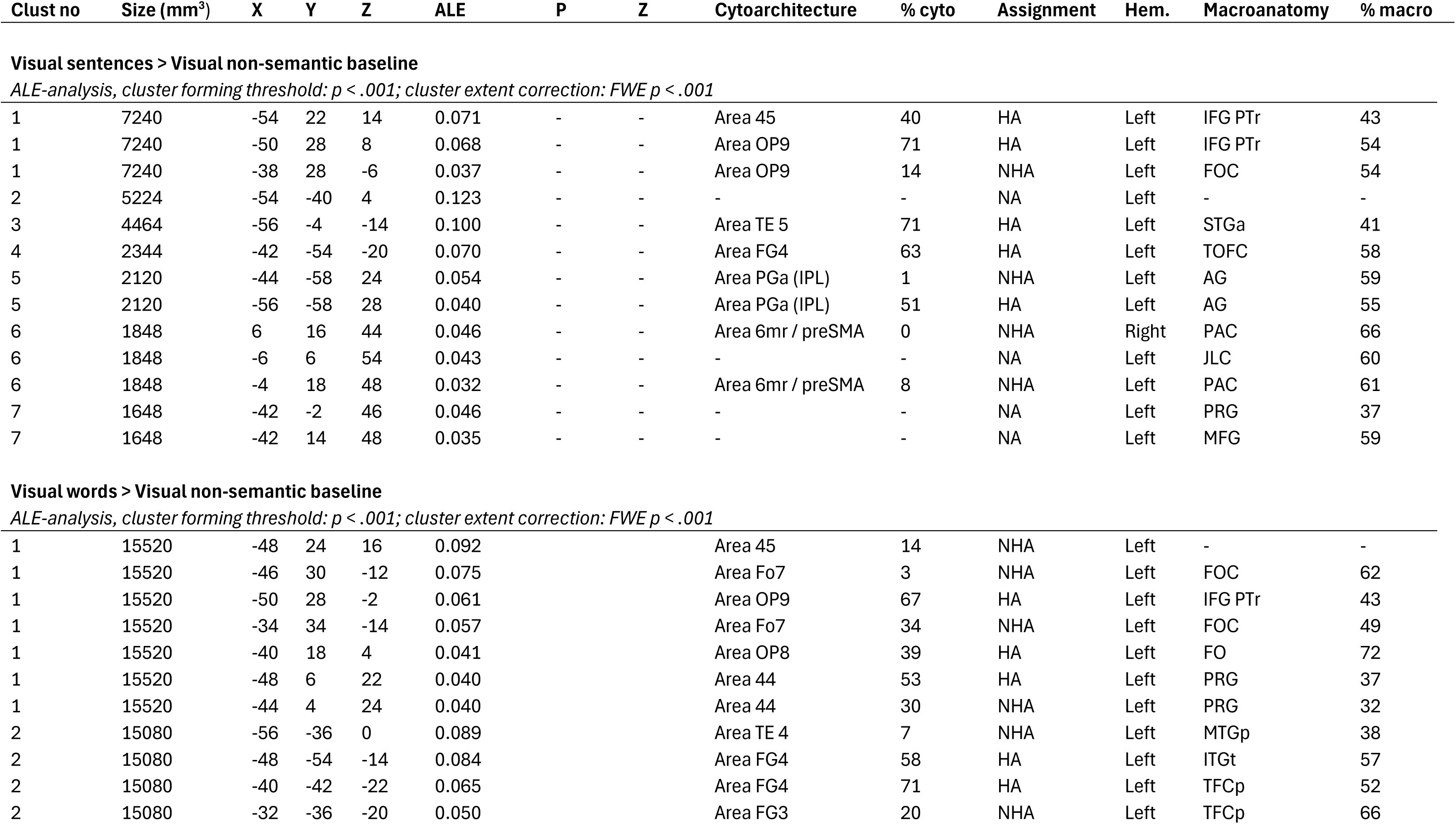
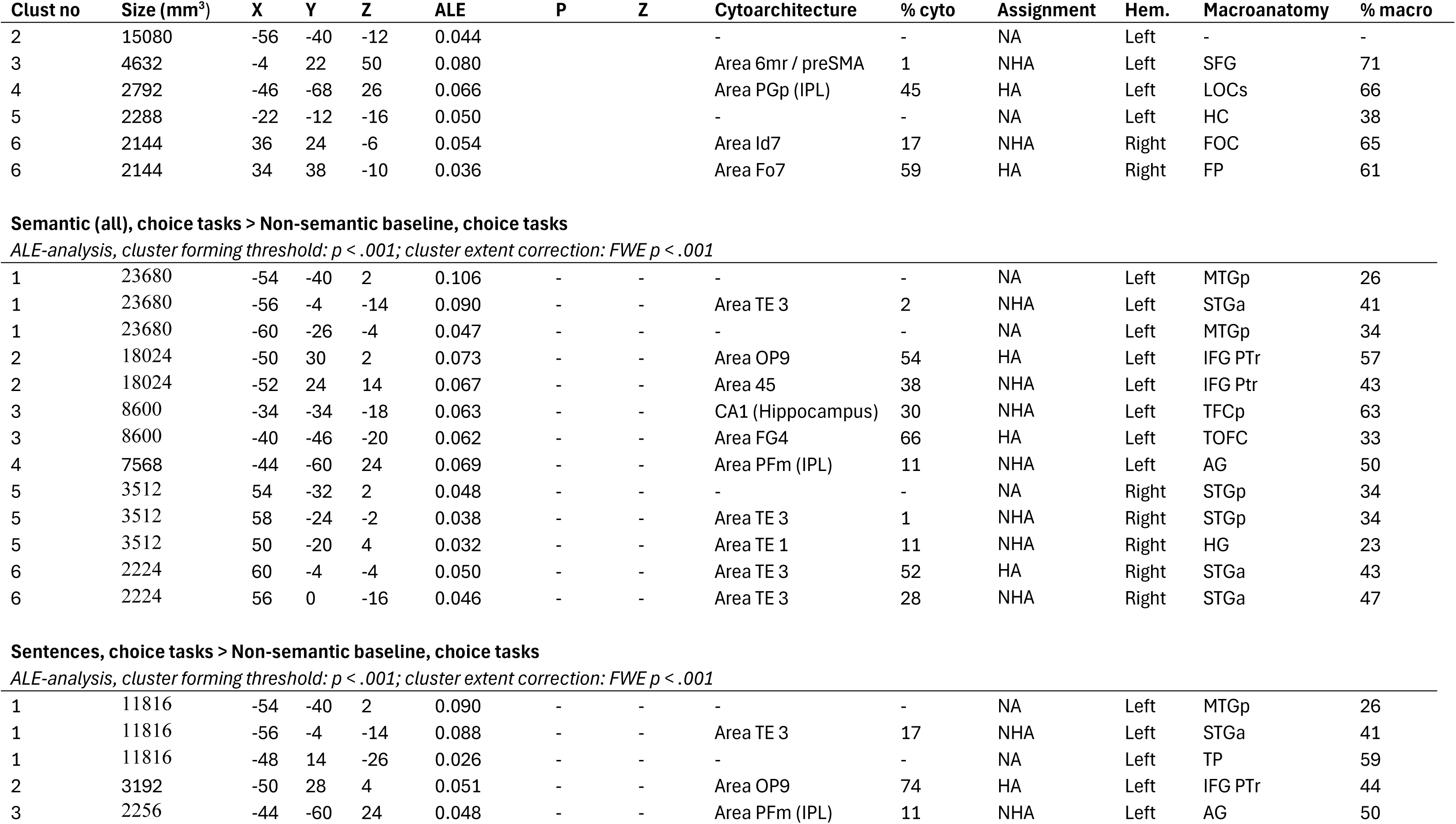
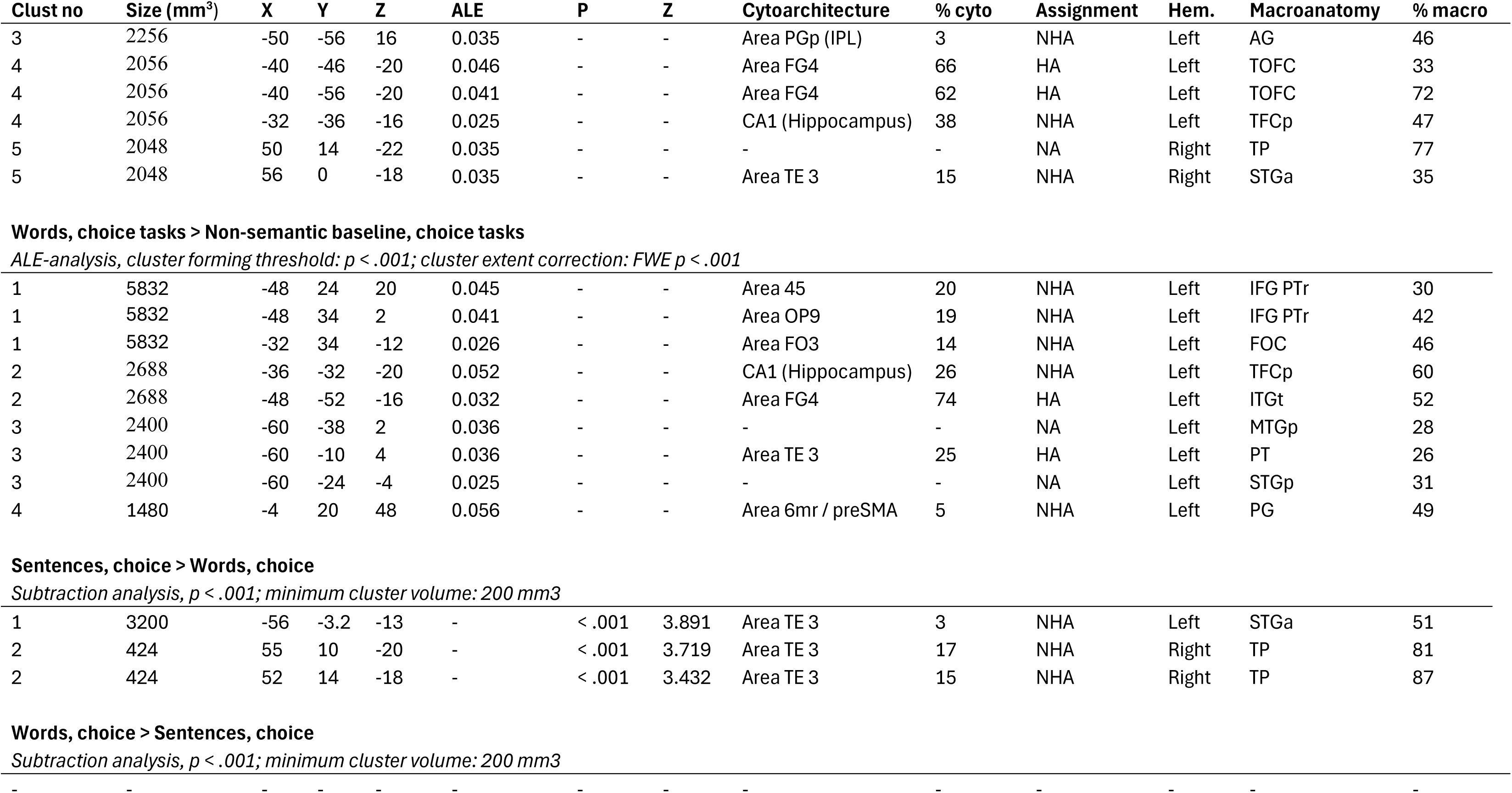
All activation clusters and local maxima for verbal semantic cognition. Coordinates x, y and z reported in the MNI coordinate system; “Cluster”: Cluster number in the individual contrast; “ALE”: Ale values output from Ginger ALE, along with P and Z values; “Cytoarchitecture”: cytoarchitectonic information for foci assigned by the JuBrain Anatomy Toolbox (SPM), based on the Maximum Probability Map; “% cyto”: probability of the coordinate falling into the specified Cytoarchitecture, as an output of the Anatomy Toolbox; “Assignment”: Type of assignment of coordinate into the specified Cytoarchitecture, as an output of the Anatomy Toolbox – HA: Hard Assignment, NHA: No Hard Assignment, NA: No Assignment “Hem”: hemisphere; “Macroanatomy”: Assignment of the foci and to the Harvard-Oxford microanatomical atlas; “% macro”: probability of the coordinate falling into the assigned region by the Harvard-Oxford microanatomical atlas; “AG”: Angular Gyrus; “AMYG”: Amygdala; “CGa”: Cingulate Gyrus, anterior; “CGp”: Cingulate Gyrus, posterior; “COP”: Central Opercular Cortex; “CRcr-I”: Cerebellum Crus I; “CRcr-II”: Cerebellum Crus II; “FMC”: Frontal Medial Cortex; “FO”: Frontal Operculum Cortex; “FOC”: Frontal Orbital Cortex; “FP”: Frontal Pole; “HC”: Hippocampus; “HG”: Heschl’s Gyrus; “IC”: Insular Cortex; “IFG POp”: Inferior Frontal Gyrus, pars opercularis; “IFG PTr”: Inferior Frontal Gyrus, pars triangularis; “IFGt”: Inferior Frontal Gyrus, temporooccipital; “ITGp”: Inferior Temporal Gyrus, posterior; “ITGt”: Inferior Temporal Gyrus, temporooccipital; “JLC”: Juxtapositional Lobule Cortex; “LOCi”: Lateral Occipital Cortex, inferior; “LOCs”: Lateral Occipital Cortex, superior; “MFG”: Middle Frontal Gyrus; “MTGa”: Middle Temporal Gyrus, anterior; “MTGp”: Middle Temporal Gyrus, posterior; “MTGt”: Middle Temporal Gyrus, temporooccipital; “OFC”: Occipital Fusiform Gyrus; “OP”: Occipital Pole; “PAC”: Paracingulate Gyrus; “PC”: Precuneous Cortex; “PGp”: Parahippocampal Gyrus, posterior; “POC”: Parietal Operculum Cortex; “PP”: Planum Polare; “PRG”: Precentral Gyrus; “PT”: Planum Temporale; “RC”: Right Caudate; “SFG”: Superior Frontal Gyrus; “SGp”: Supramarginal Gyrus, posterior; “SPL”: Superior Parietal Lobule; “STGa”: Superior Temporal Gyrus, anterior; “STGp”: Superior Temporal Gyrus, posterior; “STGs”: Superior Temporal Gyrus, superior; “TFCp”: Temporal Fusiform Cortex, posterior; “TOFC”: Temporal Occipital Fusiform Cortex; “TP”: Temporal Pole

| Clust no | Size (mm<sup>3</sup>) | X | Y | Z | ALE | P | Z | Cytoarchitecture | % cyto | Assignment | Hem. | Macroanatomy | % macro |
| --- | --- | --- | --- | --- | --- | --- | --- | --- | --- | --- | --- | --- | --- |
| **Semantic (all) > Non-semantic baseline** |  |  |  |  |  |  |  |  |  |  |  |  |  |
| *ALE-analysis, cluster forming threshold: $p < .001$; cluster extent correction: FWE $p < .001$* |  |  |  |  |  |  |  |  |  |  |  |  |  |
| 1 | 41416 | -54 | -40 | 2 | 0.240 | - | - | - | - | NA | Left | MTGp | 26 |
| 1 | 41416 | -56 | -4 | -14 | 0.173 | - | - | Area TE 5 | 71 | HA | Left | STGa | 41 |
| 1 | 41416 | -40 | -44 | -20 | 0.148 | - | - | Area FG4 | 72 | HA | Left | TFCp | 32 |
| 1 | 41416 | -44 | -60 | 24 | 0.144 | - | - | Area PGp (IPL) | 4 | NHA | Left | AG | 50 |
| 1 | 41416 | -48 | 14 | -26 | 0.123 | - | - | - | - | NA | Left | TP | 59 |
| 1 | 41416 | -32 | -36 | -18 | 0.114 | - | - | Area FG3 | 11 | NHA | Left | TFCp | 65 |
| 2 | 23488 | -50 | 24 | 14 | 0.182 | - | - | Area 45 | 33 | HA | Left | IFG PTr | 36 |
| 2 | 23488 | -50 | 28 | 0 | 0.165 | - | - | Area OP9 | 67 | HA | Left | IFG PTr | 42 |
| 2 | 23488 | -46 | 30 | -10 | 0.134 | - | - | Area OP9 | 12 | NHA | Left | FOC | 43 |
| 3 | 7480 | -4 | 20 | 48 | 0.148 | - | - | Area 6mr / preSMA | 6 | NHA | Left | PAC | 49 |
| 3 | 7480 | -6 | 8 | 54 | 0.083 | - | - | Area 6mr / preSMA | 84 | HA | Left | JLC | 37 |
| 4 | 6624 | 56 | 0 | -16 | 0.108 | - | - | Area TE 5 | 64 | HA | Right | STGa | 47 |
| 4 | 6624 | 50 | 14 | -24 | 0.083 | - | - | Area TE 5 | 2 | NHA | Right | TP | 81 |
| 4 | 6624 | 52 | -18 | -10 | 0.063 | - | - | Area TE 5 | 71 | HA | Right | MTGp | 51 |
| 5 | 3304 | -44 | 0 | 48 | 0.093 | - | - | - | - | NA | Left | PRG | 44 |
| 6 | 3184 | 36 | 24 | -6 | 0.083 | - | - | Area Id7 | 17 | NHA | Right | FOC | 65 |
| 6 | 3184 | 34 | 38 | -10 | 0.065 | - | - | Area Fo7 | 59 | HA | Right | FP | 61 |
| 7 | 2936 | 56 | -32 | 0 | 0.098 | - | - | Area TE 4 | 16 | NHA | Right | STGp | 31 |
| 7 | 2936 | 56 | -32 | 0 | 0.098 | - | - | Area TE 5 | 20 | NHA | Right | STGp | 31 |
| 8 | 2928 | -22 | -10 | -14 | 0.104 | - | - | Amygdala (VTM) | 29 | HA | Left | AMYG | 92 |
| 9 | 2688 | -4 | -58 | 18 | 0.093 | - | - | - | - | NA | Left | PC | 68 |
| 10 | 2552 | -32 | -64 | 40 | 0.077 | - | - | Area hIP6 (IPS) | 39 | HA | Left | LOCs | 54 |
| **Sentences > Non-semantic baseline** |  |  |  |  |  |  |  |  |  |  |  |  |  |
| *ALE-analysis, cluster forming threshold: $p < .001$; cluster extent correction: FWE $p < .001$* |  |  |  |  |  |  |  |  |  |  |  |  |  |
| 1 | 25056 | -52 | -40 | 4 | 0.174 | - | - | - | - | NA | Left | - | - |
| 1 | 25056 | -56 | -6 | -14 | 0.162 | - | - | Area TE 5 | 69 | HA | Left | MTGa | 38 |
| 1 | 25056 | -48 | 14 | -26 | 0.094 | - | - | - | - | NA | Left | TP | 59 |
| 1 | 25056 | -44 | -60 | 24 | 0.091 | - | - | Area PGp (IPL) | 4 | NHA | Left | AG | 50 |
| 1 | 25056 | -54 | -58 | 28 | 0.058 | - | - | Area PGa (IPL) | 57 | HA | Left | AG | 58 |
| 2 | 12704 | -54 | 22 | 14 | 0.114 | - | - | Area 45 | 40 | HA | Left | IFG PTr | 43 |
| 2 | 12704 | -50 | 28 | 2 | 0.105 | - | - | Area OP9 | 67 | HA | Left | IFG PTr | 49 |
| 3 | 9368 | 56 | 0 | -18 | 0.084 | - | - | Area TE 5 | 79 | HA | Right | STGa | 35 |
| 3 | 9368 | 52 | -34 | 0 | 0.069 | - | - | Area TE 5 | 25 | NHA | Right | MTGp | 33 |
| 3 | 9368 | 50 | 14 | -24 | 0.068 | - | - | Area TE 5 | 2 | NHA | Right | TP | 81 |
| 3 | 9368 | 52 | -18 | -8 | 0.054 | - | - | Area TE 5 | 57 | HA | Right | MTGp | 32 |
| 3 | 9368 | 64 | -32 | -2 | 0.046 | - | - | Area TE 5 | 25 | NHA | Right | MTGp | 56 |
| 4 | 3608 | -6 | 8 | 54 | 0.068 | - | - | Area 6mr / preSMA | 84 | HA | Left | JLC | 37 |
| 4 | 3608 | 6 | 18 | 44 | 0.050 | - | - | - | - | NA | Right | PAC | 62 |
| 4 | 3608 | -8 | 14 | 62 | 0.048 | - | - | Area 6mr / preSMA | 68 | HA | Left | - | - |
| 5 | 3328 | -40 | -46 | -20 | 0.087 | - | - | Area FG4 | 72 | HA | Left | TOFC | 33 |
| 5 | 3328 | -30 | -36 | -18 | 0.047 | - | - | Area FG3 | 10 | NHA | Left | TFCp | 64 |
| 6 | 2680 | -44 | 0 | 50 | 0.065 | - | - | - | - | NA | Left | PRG | 31 |
| 6 | 2680 | -40 | 12 | 50 | 0.048 | - | - | - | - | NA | Left | MFG | 51 |
| 6 | 2680 | -42 | 22 | 42 | 0.037 | - | - | - | - | NA | Left | MFG | 67 |
| 1 | 18616 | -56 | -36 | 0 | 0.096 | - | - | Area TE 4 | 7 | NHA | Left | MTGp | 38 |
| 1 | 18616 | -48 | -54 | -14 | 0.095 | - | - | Area FG4 | 58 | HA | Left | ITGt | 57 |
| 1 | 18616 | -40 | -42 | -22 | 0.077 | - | - | Area FG4 | 71 | HA | Left | TFCp | 52 |
| 1 | 18616 | -34 | -34 | -20 | 0.076 | - | - | Area FG3 | 12 | NHA | Left | TFCp | 64 |
| 1 | 18616 | -58 | -14 | -2 | 0.056 | - | - | Area TE 4 | 62 | HA | Left | - | - |
| 1 | 18616 | -56 | -42 | -12 | 0.050 | - | - | - | - | NA | Left | - | - |
| 2 | 17744 | -48 | 24 | 18 | 0.118 | - | - | Area 45 | 14 | NHA | Left | IFG PTr | 25 |
| 2 | 17744 | -46 | 30 | -10 | 0.083 | - | - | Area OP9 | 12 | NHA | Left | FOC | 43 |
| 2 | 17744 | -36 | 32 | -14 | 0.071 | - | - | Area Fo7 | 14 | NHA | Left | FOC | 48 |
| 2 | 17744 | -40 | 4 | 26 | 0.054 | - | - | Area 44 | 14 | NHA | Left | - | - |
| 2 | 17744 | -32 | 24 | -4 | 0.043 | - | - | Area Id7 | 32 | NHA | Left | IC | 41 |
| 3 | 6624 | -46 | -68 | 26 | 0.089 | - | - | Area PGp (IPL) | 45 | HA | Left | LOCs | 66 |
| 3 | 6624 | -32 | -64 | 40 | 0.065 | - | - | Area hIP6 (IPS) | 39 | HA | Left | LOCs | 54 |
| 3 | 6624 | -34 | -52 | 44 | 0.042 | - | - | Area hIP3 (IPS) | 43 | HA | Left | SPL | 31 |
| 4 | 5016 | -4 | 20 | 48 | 0.109 | - | - | Area 6mr / preSMA | 6 | NHA | Left | PAC | 49 |
| 5 | 2792 | 36 | 24 | -6 | 0.067 | - | - | Area Id7 | 17 | NHA | Right | FOC | 65 |
| 5 | 2792 | 34 | 38 | -8 | 0.051 | - | - | Area Fo7 | 54 | HA | Right | - | - |
| 1 | 4560 | -56 | -7 | -13 | - | <.001 | 3.891 | Area TE 5 | 67 | HA | Left | MTGa | 31 |
| 1 | 4560 | -51 | 6 | -27 | - | <.001 | 3.432 | Area TE 5 | 42 | NHA | Left | TP | 64 |
| 1 | 4560 | -50 | 4 | -18 | - | <.001 | 3.540 | Area TE 5 | 47 | HA | Left | TP | 40 |
| 2 | 1248 | 58 | 1 | -22 | - | <.001 | 3.891 | Area TE 5 | 76 | HA | Right | MTGa | 51 |
| 2 | 1248 | 51 | 8 | -25 | - | <.001 | 3.121 | Area TE 5 | 59 | HA | Right | TP | 70 |
| 2 | 1248 | 51 | 11 | -26 | - | <.001 | 3.540 | Area TE 5 | 21 | NHA | Right | TP | 84 |
| 2 | 1248 | 48 | 18 | -20 | - | <.001 | 3.291 | - | - | NA | Right | TP | 72 |
| 3 | 912 | -51 | -51 | 12 | - | <.001 | 3.891 | - | - | NA | Left | AG | 25 |
| 3 | 912 | -47 | -53 | 15 | - | <.001 | 3.719 | - | - | NA | Left | AG | 27 |
| 3 | 912 | -48 | -42 | 4 | - | <.001 | 3.432 | - | - | NA | Left | - | - |
| 4 | 552 | -57 | 23 | 12 | - | <.001 | 3.891 | Area 45 | 55 | HA | Left | IFG PTr | 49 |
| 5 | 224 | 48 | -32 | -6 | - | <.001 | 3.719 | Area TE 4 | 16 | NHA | Right | MTGp | 48 |
| 5 | 224 | 52 | -34 | -6 | - | <.001 | 3.540 | - | - | NA | Right | - | - |
| 1 | 392 | -46 | -75 | 27 | - | <.000 | 3.891 | Area PGp (IPL) | 81 | HA | Left | LOCs | 87 |
| 1 | 392 | -46 | -76 | 32 | - | <.001 | 3.719 | Area PGp (IPL) | 79 | HA | Left | LOCs | 92 |
| 1 | 392 | -42 | -80 | 30 | - | <.001 | 3.540 | Area PGp (IPL) | 56 | HA | Left | LOCs | 85 |
| 2 | 264 | -47 | -48 | -12 | - | <.001 | 3.719 | Area FG4 | 85 | HA | Left | ITGt | 34 |
| 2 | 264 | -50 | -41 | -15 | - | <.001 | 3.432 | Area FG4 | 26 | NHA | Left | - | - |
| 3 | 264 | -3 | 31 | 38 | - | <.001 | 3.891 | - | - | NA | Left | PAC | 75 |
| 1 | 19880 | -58 | -12 | -4 | 0.104 | - | - | Area TE 4 | 77 | HA | Left | STGa | 28 |
| 1 | 19880 | -56 | -6 | -12 | 0.078 | - | - | Area TE 5 | 62 | HA | Left | STGa | 35 |
| 1 | 19880 | -52 | 8 | -22 | 0.069 | - | - | Area TE 5 | 36 | NHA | Left | TP | 86 |
| 1 | 19880 | -52 | -40 | 4 | 0.065 | - | - | - | - | NA | Left | - | - |
| 1 | 19880 | -46 | -60 | 22 | 0.062 | - | - | Area PGp (IPL) | 11 | NHA | Left | AG | 48 |
| 1 | 19880 | -60 | -32 | 4 | 0.048 | - | - | Area TE 4 | 2 | NHA | Left | STGp | 31 |
| 2 | 14456 | 60 | -6 | -8 | 0.085 | - | - | Area TE 3 | 39 | HA | Right | STGa | 36 |
| 2 | 14456 | 58 | -32 | 2 | 0.072 | - | - | Area TE 4 | 18 | NHA | Right | STGp | 32 |
| 2 | 14456 | 52 | 4 | -20 | 0.065 | - | - | Area TE 5 | 73 | HA | Right | STGa | 35 |
| 2 | 14456 | 52 | -18 | -8 | 0.046 | - | - | Area TE 5 | 57 | HA | Right | MTGp | 32 |
| 2 | 14456 | 64 | -18 | -2 | 0.043 | - | - | Area TE 4 | 41 | HA | Right | - | - |
| 2 | 14456 | 50 | -18 | 4 | 0.038 | - | - | Area TE 1.1 | 2 | NHA | Right | - | - |
| 3 | 7616 | -50 | 30 | 0 | 0.068 | - | - | Area OP9 | 52 | HA | Left | IFG PTr | 51 |
| 3 | 7616 | -46 | 16 | 22 | 0.053 | - | - | Area 44 | 23 | NHA | Left | IFG POp | 51 |
| 3 | 7616 | -52 | 24 | 14 | 0.050 | - | - | Area 45 | 42 | HA | Left | IFG PTr | 43 |
| 3 | 7616 | -40 | 32 | -12 | 0.049 | - | - | Area Fo7 | 23 | NHA | Left | FOC | 45 |
| 4 | 2416 | -36 | -34 | -18 | 0.050 | - | - | CA1 (Hippocampus) | 26 | NHA | Left | TFCp | 55 |
| 4 | 2416 | -40 | -42 | -18 | 0.044 | - | - | Area FG4 | 56 | HA | Left | TFCp | 37 |
| 1 | 21848 | -50 | 24 | 14 | 0.136 | - | - | Area 45 | 33 | HA | Left | IFG PTr | 36 |
| 1 | 21848 | -52 | 28 | 4 | 0.108 | - | - | Area OP9 | 55 | HA | Left | IFG PTr | 58 |
| 1 | 21848 | -46 | 30 | -10 | 0.097 | - | - | Area OP9 | 12 | NHA | Left | FOC | 43 |
| 1 | 21848 | -48 | 16 | -26 | 0.084 | - | - | - | - | NA | Left | TP | 65 |
| 1 | 21848 | -36 | 32 | -14 | 0.079 | - | - | Area Fo7 | 14 | NHA | Left | FOC | 48 |
| 1 | 21848 | -44 | 6 | 26 | 0.056 | - | - | Area 44 | 51 | HA | Left | IFG POp | 34 |
| 1 | 21848 | -38 | 18 | 2 | 0.047 | - | - | Area Id6 | 54 | HA | Left | FO | 40 |
| 2 | 21072 | -54 | -40 | 2 | 0.192 | - | - | - | - | NA | Left | MTGp | 26 |
| 2 | 21072 | -44 | -54 | -16 | 0.112 | - | - | Area FG4 | 78 | HA | Left | TOFC | 42 |
| 2 | 21072 | -40 | -46 | -20 | 0.110 | - | - | Area FG4 | 72 | HA | Left | TOFC | 33 |
| 2 | 21072 | -22 | -10 | -14 | 0.085 | - | - | Amygdala (VTM) | 29 | HA | Left | AMYG | 92 |
| 2 | 21072 | -32 | -36 | -20 | 0.069 | - | - | Area FG3 | 20 | NHA | Left | TFCp | 66 |
| 3 | 7936 | -4 | 20 | 48 | 0.106 | - | - | Area 6mr / preSMA | 6 | NHA | Left | PAC | 49 |
| 3 | 7936 | -10 | 46 | 40 | 0.060 | - | - | - | - | NA | Left | - | - |
| 3 | 7936 | -10 | 52 | 38 | 0.059 | - | - | - | - | NA | Left | FP | 70 |
| 3 | 7936 | -6 | 6 | 54 | 0.059 | - | - | - | - | NA | Left | JLC | 60 |
| 3 | 7936 | -8 | 16 | 60 | 0.048 | - | - | Area 6mr / preSMA | 60 | HA | Left | - | - |
| 4 | 4264 | -56 | -2 | -16 | 0.115 | - | - | Area TE 5 | 78 | HA | Left | MTGa | 42 |
| 5 | 4264 | -42 | -58 | 24 | 0.085 | - | - | Area PGa (IPL) | 0 | NHA | Left | AG | 53 |
| 5 | 3920 | -46 | -66 | 26 | 0.082 | - | - | Area PGp (IPL) | 35 | NHA | Left | LOCs | 54 |
| 6 | 3304 | -44 | -2 | 48 | 0.074 | - | - | - | - | NA | Left | PRG | 44 |
| 7 | 2376 | 36 | 24 | -6 | 0.060 | - | - | Area Id7 | 17 | NHA | Right | FOC | 65 |
| 7 | 2376 | 36 | 36 | -10 | 0.052 | - | - | Area Fo7 | 45 | HA | Right | FP | 50 |
| 8 | 2120 | -2 | -58 | 18 | 0.080 | - | - | - | - | NA | Left | PC | 73 |
| 8 | 2120 | -12 | -50 | 2 | 0.044 | - | - | - | - | NA | Left | CGp | 41 |
| 9 | 1992 | -30 | -58 | 44 | 0.058 | - | - | Area hIP3 (IPS) | 66 | HA | Left | SPL | 30 |
| 1 | 16224 | -58 | -12 | -6 | 0.077 | - | - | Area TE 4 | 78 | HA | Left | STGa | 22 |
| 1 | 16224 | -56 | -6 | -12 | 0.072 | - | - | Area TE 5 | 62 | HA | Left | STGa | 35 |
| 1 | 16224 | -52 | 8 | -22 | 0.066 | - | - | Area TE 5 | 36 | NHA | Left | TP | 86 |
| 1 | 16224 | -52 | -40 | 4 | 0.058 | - | - | - | - | NA | Left | STGp | 18 |
| 1 | 16224 | -64 | -22 | 0 | 0.043 | - | - | Area TE 4 | 37 | HA | Left | STGp | 41 |
| 1 | 16224 | -44 | -60 | 24 | 0.039 | - | - | Area PGp (IPL) | 4 | NHA | Left | AG | 50 |
| 1 | 16224 | -52 | -52 | 16 | 0.039 | - | - | - | - | NA | Left | AG | 35 |
| 1 | 16224 | -62 | -52 | 0 | 0.028 | - | - | - | - | NA | Left | MTGt | 68 |
| 2 | 9648 | 58 | -6 | -10 | 0.063 | - | - | Area TE 4 | 46 | HA | Right | STGa | 32 |
| 2 | 9648 | 52 | 4 | -20 | 0.057 | - | - | Area TE 5 | 73 | HA | Right | STGa | 35 |
| 2 | 9648 | 50 | 14 | -24 | 0.047 | - | - | Area TE 5 | 2 | NHA | Right | TP | 81 |
| 2 | 9648 | 52 | -34 | 0 | 0.045 | - | - | Area TE 5 | 25 | NHA | Right | MTGp | 33 |
| 2 | 9648 | 52 | -18 | -8 | 0.044 | - | - | Area TE 5 | 57 | HA | Right | MTGp | 32 |
| 2 | 9648 | 64 | -18 | -10 | 0.027 | - | - | Area TE 5 | 57 | HA | Right | MTGp | 57 |
| 2 | 9648 | 66 | -24 | 4 | 0.025 | - | - | Area TE 3 | 39 | HA | Right | STGp | 38 |
| 2 | 9648 | 52 | -20 | 2 | 0.023 | - | - | Area TE 4 | 7 | NHA | Right | - | - |
| 3 | 5520 | -50 | 28 | 0 | 0.053 | - | - | Area OP9 | 67 | HA | Left | IFG PTr | 42 |
| 3 | 5520 | -44 | 14 | 22 | 0.044 | - | - | Area 44 | 23 | NHA | Left | - | - |
| 3 | 5520 | -54 | 24 | 14 | 0.043 | - | - | Area 45 | 55 | HA | Left | IFG PTr | 51 |
| 3 | 5520 | -42 | 32 | -14 | 0.032 | - | - | Area Fo7 | 52 | HA | Left | FOC | 49 |
| 1 | 3440 | -60 | -12 | 2 | 0.045 | - | - | Area TE 3 | 32 | HA | Left | PT | 25 |
| 1 | 3440 | -62 | -30 | 2 | 0.024 | - | - | Area TE 4 | 16 | NHA | Left | STGp | 30 |
| 2 | 1872 | 58 | -32 | 2 | 0.044 | - | - | Area TE 4 | 18 | NHA | Right | STGp | 32 |
| 3 | 1784 | -46 | 34 | 4 | 0.037 | - | - | Area OP9 | 36 | NHA | Left | IFG PTr | 31 |
| 3 | 1784 | -48 | 22 | 20 | 0.024 | - | - | Area 45 | 9 | NHA | Left | IFG POp | 25 |
| 3 | 1784 | -48 | 26 | 18 | 0.023 | - | - | Area 45 | 24 | NHA | Left | IFG PTr | 33 |
| 3 | 1784 | -42 | 32 | -8 | 0.021 | - | - | Area OP9 | 6 | NHA | Left | - | - |
| 4 | 1504 | 60 | -4 | -4 | 0.042 | - | - | Area TE 3 | 42 | HA | Right | STGa | 43 |
| 5 | 1288 | -36 | -32 | -18 | 0.036 | - | - | CA1 (Hippocampus) | 36 | NHA | Left | TFCp | 60 |
| 5 | 1288 | -28 | -38 | -20 | 0.026 | - | - | Area FG3 | 39 | NHA | Left | TFCp | 72 |
| **Visual sentences > Visual non-semantic baseline** |  |  |  |  |  |  |  |  |  |  |  |  |  |
| *ALE-analysis, cluster forming threshold: $p < .001$; cluster extent correction: FWE $p < .001$* |  |  |  |  |  |  |  |  |  |  |  |  |  |
| 1 | 7240 | -54 | 22 | 14 | 0.071 | - | - | Area 45 | 40 | HA | Left | IFG PTr | 43 |
| 1 | 7240 | -50 | 28 | 8 | 0.068 | - | - | Area OP9 | 71 | HA | Left | IFG PTr | 54 |
| 1 | 7240 | -38 | 28 | -6 | 0.037 | - | - | Area OP9 | 14 | NHA | Left | FOC | 54 |
| 2 | 5224 | -54 | -40 | 4 | 0.123 | - | - | - | - | NA | Left | - | - |
| 3 | 4464 | -56 | -4 | -14 | 0.100 | - | - | Area TE 5 | 71 | HA | Left | STGa | 41 |
| 4 | 2344 | -42 | -54 | -20 | 0.070 | - | - | Area FG4 | 63 | HA | Left | TOFC | 58 |
| 5 | 2120 | -44 | -58 | 24 | 0.054 | - | - | Area PGa (IPL) | 1 | NHA | Left | AG | 59 |
| 5 | 2120 | -56 | -58 | 28 | 0.040 | - | - | Area PGa (IPL) | 51 | HA | Left | AG | 55 |
| 6 | 1848 | 6 | 16 | 44 | 0.046 | - | - | Area 6mr / preSMA | 0 | NHA | Right | PAC | 66 |
| 6 | 1848 | -6 | 6 | 54 | 0.043 | - | - | - | - | NA | Left | JLC | 60 |
| 6 | 1848 | -4 | 18 | 48 | 0.032 | - | - | Area 6mr / preSMA | 8 | NHA | Left | PAC | 61 |
| 7 | 1648 | -42 | -2 | 46 | 0.046 | - | - | - | - | NA | Left | PRG | 37 |
| 7 | 1648 | -42 | 14 | 48 | 0.035 | - | - | - | - | NA | Left | MFG | 59 |
| 1 | 15520 | -48 | 24 | 16 | 0.092 |  |  | Area 45 | 14 | NHA | Left | - | - |
| 1 | 15520 | -46 | 30 | -12 | 0.075 |  |  | Area Fo7 | 3 | NHA | Left | FOC | 62 |
| 1 | 15520 | -50 | 28 | -2 | 0.061 |  |  | Area OP9 | 67 | HA | Left | IFG PTr | 43 |
| 1 | 15520 | -34 | 34 | -14 | 0.057 |  |  | Area Fo7 | 34 | NHA | Left | FOC | 49 |
| 1 | 15520 | -40 | 18 | 4 | 0.041 |  |  | Area OP8 | 39 | HA | Left | FO | 72 |
| 1 | 15520 | -48 | 6 | 22 | 0.040 |  |  | Area 44 | 53 | HA | Left | PRG | 37 |
| 1 | 15520 | -44 | 4 | 24 | 0.040 |  |  | Area 44 | 30 | NHA | Left | PRG | 32 |
| 2 | 15080 | -56 | -36 | 0 | 0.089 |  |  | Area TE 4 | 7 | NHA | Left | MTGp | 38 |
| 2 | 15080 | -48 | -54 | -14 | 0.084 |  |  | Area FG4 | 58 | HA | Left | ITGt | 57 |
| 2 | 15080 | -40 | -42 | -22 | 0.065 |  |  | Area FG4 | 71 | HA | Left | TFCp | 52 |
| 2 | 15080 | -32 | -36 | -20 | 0.050 |  |  | Area FG3 | 20 | NHA | Left | TFCp | 66 |
| 2 | 15080 | -56 | -40 | -12 | 0.044 |  |  | - | - | NA | Left | - | - |
| 3 | 4632 | -4 | 22 | 50 | 0.080 |  |  | Area 6mr / preSMA | 1 | NHA | Left | SFG | 71 |
| 4 | 2792 | -46 | -68 | 26 | 0.066 |  |  | Area PGp (IPL) | 45 | HA | Left | LOCs | 66 |
| 5 | 2288 | -22 | -12 | -16 | 0.050 |  |  | - | - | NA | Left | HC | 38 |
| 6 | 2144 | 36 | 24 | -6 | 0.054 |  |  | Area Id7 | 17 | NHA | Right | FOC | 65 |
| 6 | 2144 | 34 | 38 | -10 | 0.036 |  |  | Area Fo7 | 59 | HA | Right | FP | 61 |
| 1 | 23680 | -54 | -40 | 2 | 0.106 | - | - | - | - | NA | Left | MTGp | 26 |
| 1 | 23680 | -56 | -4 | -14 | 0.090 | - | - | Area TE 3 | 2 | NHA | Left | STGa | 41 |
| 1 | 23680 | -60 | -26 | -4 | 0.047 | - | - | - | - | NA | Left | MTGp | 34 |
| 2 | 18024 | -50 | 30 | 2 | 0.073 | - | - | Area OP9 | 54 | HA | Left | IFG PTr | 57 |
| 2 | 18024 | -52 | 24 | 14 | 0.067 | - | - | Area 45 | 38 | NHA | Left | IFG Ptr | 43 |
| 3 | 8600 | -34 | -34 | -18 | 0.063 | - | - | CA1 (Hippocampus) | 30 | NHA | Left | TFCp | 63 |
| 3 | 8600 | -40 | -46 | -20 | 0.062 | - | - | Area FG4 | 66 | HA | Left | TOFC | 33 |
| 4 | 7568 | -44 | -60 | 24 | 0.069 | - | - | Area PFm (IPL) | 11 | NHA | Left | AG | 50 |
| 5 | 3512 | 54 | -32 | 2 | 0.048 | - | - | - | - | NA | Right | STGp | 34 |
| 5 | 3512 | 58 | -24 | -2 | 0.038 | - | - | Area TE 3 | 1 | NHA | Right | STGp | 34 |
| 5 | 3512 | 50 | -20 | 4 | 0.032 | - | - | Area TE 1 | 11 | NHA | Right | HG | 23 |
| 6 | 2224 | 60 | -4 | -4 | 0.050 | - | - | Area TE 3 | 52 | HA | Right | STGa | 43 |
| 6 | 2224 | 56 | 0 | -16 | 0.046 | - | - | Area TE 3 | 28 | NHA | Right | STGa | 47 |
| 1 | 11816 | -54 | -40 | 2 | 0.090 | - | - | - | - | NA | Left | MTGp | 26 |
| 1 | 11816 | -56 | -4 | -14 | 0.088 | - | - | Area TE 3 | 17 | NHA | Left | STGa | 41 |
| 1 | 11816 | -48 | 14 | -26 | 0.026 | - | - | - | - | NA | Left | TP | 59 |
| 2 | 3192 | -50 | 28 | 4 | 0.051 | - | - | Area OP9 | 74 | HA | Left | IFG PTr | 44 |
| 3 | 2256 | -44 | -60 | 24 | 0.048 | - | - | Area PFm (IPL) | 11 | NHA | Left | AG | 50 |
| 3 | 2256 | -50 | -56 | 16 | 0.035 | - | - | Area PGp (IPL) | 3 | NHA | Left | AG | 46 |
| 4 | 2056 | -40 | -46 | -20 | 0.046 | - | - | Area FG4 | 66 | HA | Left | TOFC | 33 |
| 4 | 2056 | -40 | -56 | -20 | 0.041 | - | - | Area FG4 | 62 | HA | Left | TOFC | 72 |
| 4 | 2056 | -32 | -36 | -16 | 0.025 | - | - | CA1 (Hippocampus) | 38 | NHA | Left | TFCp | 47 |
| 5 | 2048 | 50 | 14 | -22 | 0.035 | - | - | - | - | NA | Right | TP | 77 |
| 5 | 2048 | 56 | 0 | -18 | 0.035 | - | - | Area TE 3 | 15 | NHA | Right | STGa | 35 |
| 1 | 5832 | -48 | 24 | 20 | 0.045 | - | - | Area 45 | 20 | NHA | Left | IFG PTr | 30 |
| 1 | 5832 | -48 | 34 | 2 | 0.041 | - | - | Area OP9 | 19 | NHA | Left | IFG PTr | 42 |
| 1 | 5832 | -32 | 34 | -12 | 0.026 | - | - | Area FO3 | 14 | NHA | Left | FOC | 46 |
| 2 | 2688 | -36 | -32 | -20 | 0.052 | - | - | CA1 (Hippocampus) | 26 | NHA | Left | TFCp | 60 |
| 2 | 2688 | -48 | -52 | -16 | 0.032 | - | - | Area FG4 | 74 | HA | Left | ITGt | 52 |
| 3 | 2400 | -60 | -38 | 2 | 0.036 | - | - | - | - | NA | Left | MTGp | 28 |
| 3 | 2400 | -60 | -10 | 4 | 0.036 | - | - | Area TE 3 | 25 | HA | Left | PT | 26 |
| 3 | 2400 | -60 | -24 | -4 | 0.025 | - | - | - | - | NA | Left | STGp | 31 |
| 4 | 1480 | -4 | 20 | 48 | 0.056 | - | - | Area 6mr / preSMA | 5 | NHA | Left | PG | 49 |
| 1 | 3200 | -56 | -3.2 | -13 | - | < .001 | 3.891 | Area TE 3 | 3 | NHA | Left | STGa | 51 |
| 2 | 424 | 55 | 10 | -20 | - | < .001 | 3.719 | Area TE 3 | 17 | NHA | Right | TP | 81 |
| 2 | 424 | 52 | 14 | -18 | - | < .001 | 3.432 | Area TE 3 | 15 | NHA | Right | TP | 87 |
| - | - | - | - | - | - | - | - | - | - | - | - | - | - |

**Supplementary Table 6.**
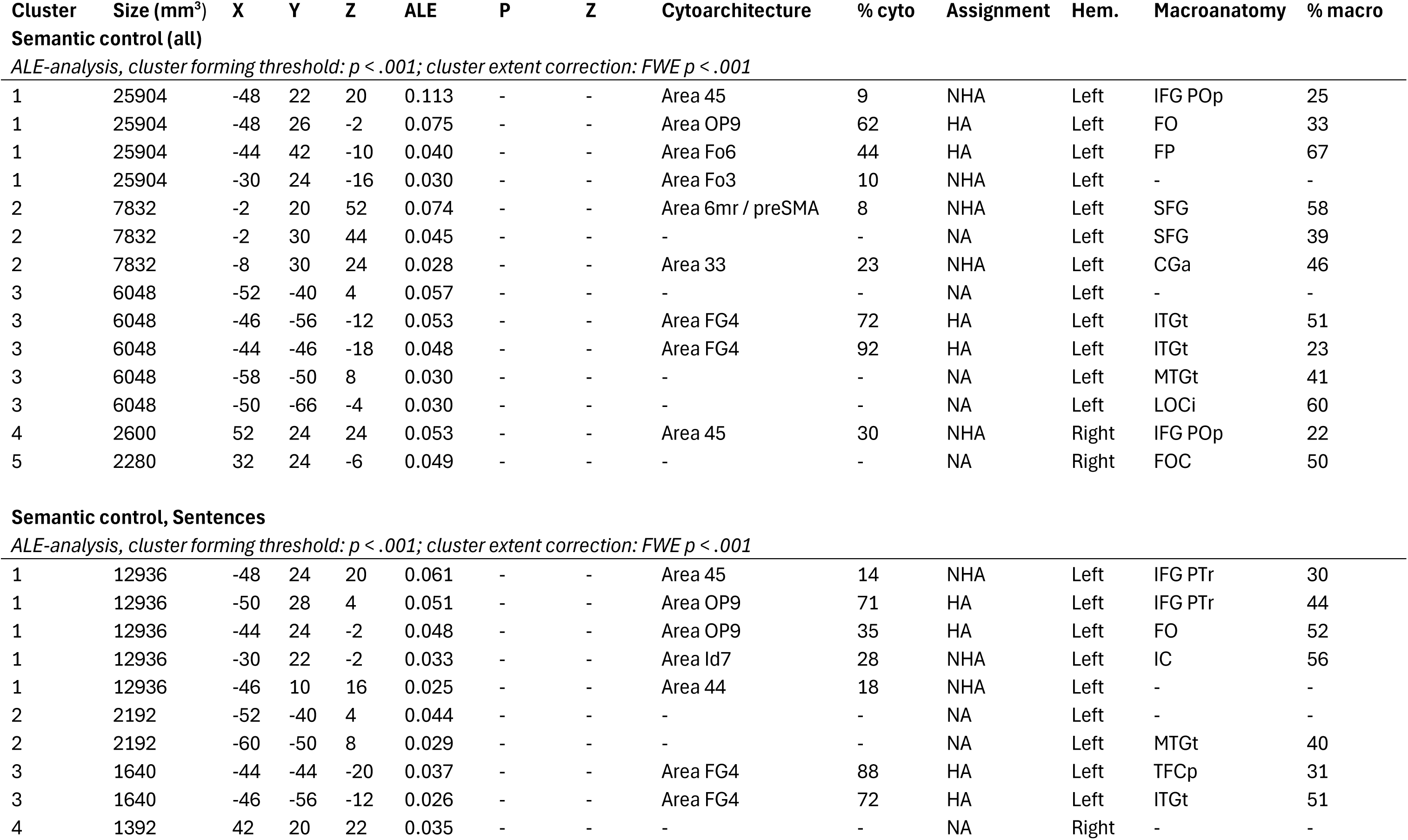
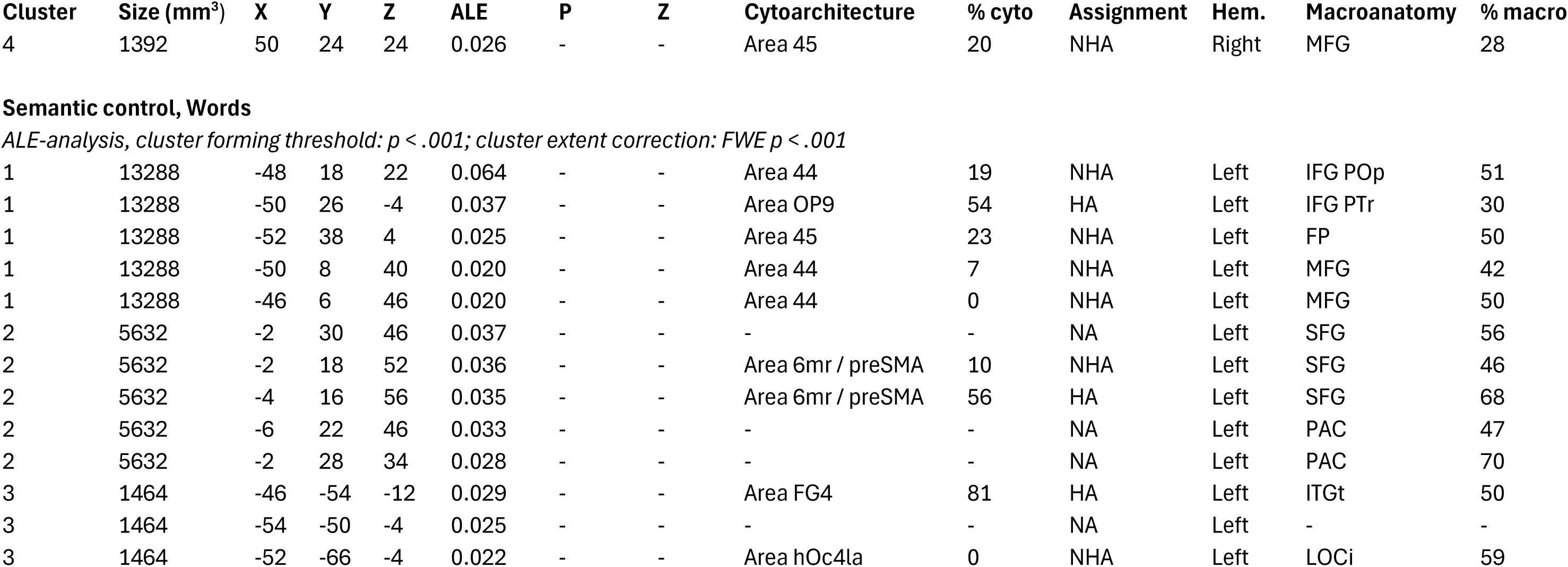
All activation clusters and local maxima for verbal semantic control. Coordinates x, y and z reported in the MNI coordinate system; “Cluster”: Cluster number in the individual contrast; “ALE”: Ale values output from Ginger ALE, along with P and Z values; “Cytoarchitecture”: cytoarchitectonic information for foci assigned by the JuBrain Anatomy Toolbox (SPM), based on the Maximum Probability Map; “% cyto”: probability of the coordinate falling into the specified Cytoarchitecture, as an output of the Anatomy Toolbox; “Assignment”: Type of assignment of coordinate into the specified Cytoarchitecture, as an output of the Anatomy Toolbox – HA: Hard Assignment, NHA: No Hard Assignment, NA: No Assignment “Hem”: hemisphere; “Macroanatomy”: Assignment of the foci and to the Harvard-Oxford microanatomical atlas; “% macro”: probability of the coordinate falling into the assigned region by the Harvard-Oxford microanatomical atlas; “AG”: Angular Gyrus; “AMYG”: Amygdala; “CGa”: Cingulate Gyrus, anterior; “CGp”: Cingulate Gyrus, posterior; “COP”: Central Opercular Cortex; “CRcr-I”: Cerebellum Crus I; “CRcr-II”: Cerebellum Crus II; “FMC”: Frontal Medial Cortex; “FO”: Frontal Operculum Cortex; “FOC”: Frontal Orbital Cortex; “FP”: Frontal Pole; “HC”: Hippocampus; “HG”: Heschl’s Gyrus; “IC”: Insular Cortex; “IFG POp”: Inferior Frontal Gyrus, pars opercularis; “IFG PTr”: Inferior Frontal Gyrus, pars triangularis; “IFGt”: Inferior Frontal Gyrus, temporooccipital; “ITGp”: Inferior Temporal Gyrus, posterior; “ITGt”: Inferior Temporal Gyrus, temporooccipital; “JLC”: Juxtapositional Lobule Cortex; “LOCi”: Lateral Occipital Cortex, inferior; “LOCs”: Lateral Occipital Cortex, superior; “MFG”: Middle Frontal Gyrus; “MTGa”: Middle Temporal Gyrus, anterior; “MTGp”: Middle Temporal Gyrus, posterior; “MTGt”: Middle Temporal Gyrus, temporooccipital; “OFC”: Occipital Fusiform Gyrus; “OP”: Occipital Pole; “PAC”: Paracingulate Gyrus; “PC”: Precuneous Cortex; “PGp”: Parahippocampal Gyrus, posterior; “POC”: Parietal Operculum Cortex; “PP”: Planum Polare; “PRG”: Precentral Gyrus; “PT”: Planum Temporale; “RC”: Right Caudate; “SFG”: Superior Frontal Gyrus; “SGp”: Supramarginal Gyrus, posterior; “SPL”: Superior Parietal Lobule; “STGa”: Superior Temporal Gyrus, anterior; “STGp”: Superior Temporal Gyrus, posterior; “STGs”: Superior Temporal Gyrus, superior; “TFCp”: Temporal Fusiform Cortex, posterior; “TOFC”: Temporal Occipital Fusiform Cortex; “TP”: Temporal Pole

| Cluster | Size (mm<sup>3</sup>) | X | Y | Z | ALE | P | Z | Cytoarchitecture | % cyto | Assignment | Hem. | Macroanatomy | % macro |
| --- | --- | --- | --- | --- | --- | --- | --- | --- | --- | --- | --- | --- | --- |
| **Semantic control (all)** |  |  |  |  |  |  |  |  |  |  |  |  |  |
| *ALE-analysis, cluster forming threshold: $p < .001$; cluster extent correction: FWE $p < .001$* |  |  |  |  |  |  |  |  |  |  |  |  |  |
| 1 | 25904 | -48 | 22 | 20 | 0.113 | - | - | Area 45 | 9 | NHA | Left | IFG POp | 25 |
| 1 | 25904 | -48 | 26 | -2 | 0.075 | - | - | Area OP9 | 62 | HA | Left | FO | 33 |
| 1 | 25904 | -44 | 42 | -10 | 0.040 | - | - | Area Fo6 | 44 | HA | Left | FP | 67 |
| 1 | 25904 | -30 | 24 | -16 | 0.030 | - | - | Area Fo3 | 10 | NHA | Left | - | - |
| 2 | 7832 | -2 | 20 | 52 | 0.074 | - | - | Area 6mr / preSMA | 8 | NHA | Left | SFG | 58 |
| 2 | 7832 | -2 | 30 | 44 | 0.045 | - | - | - | - | NA | Left | SFG | 39 |
| 2 | 7832 | -8 | 30 | 24 | 0.028 | - | - | Area 33 | 23 | NHA | Left | CGa | 46 |
| 3 | 6048 | -52 | -40 | 4 | 0.057 | - | - | - | - | NA | Left | - | - |
| 3 | 6048 | -46 | -56 | -12 | 0.053 | - | - | Area FG4 | 72 | HA | Left | ITGt | 51 |
| 3 | 6048 | -44 | -46 | -18 | 0.048 | - | - | Area FG4 | 92 | HA | Left | ITGt | 23 |
| 3 | 6048 | -58 | -50 | 8 | 0.030 | - | - | - | - | NA | Left | MTGt | 41 |
| 3 | 6048 | -50 | -66 | -4 | 0.030 | - | - | - | - | NA | Left | LOCi | 60 |
| 4 | 2600 | 52 | 24 | 24 | 0.053 | - | - | Area 45 | 30 | NHA | Right | IFG POp | 22 |
| 5 | 2280 | 32 | 24 | -6 | 0.049 | - | - | - | - | NA | Right | FOC | 50 |
| 1 | 12936 | -48 | 24 | 20 | 0.061 | - | - | Area 45 | 14 | NHA | Left | IFG PTr | 30 |
| 1 | 12936 | -50 | 28 | 4 | 0.051 | - | - | Area OP9 | 71 | HA | Left | IFG PTr | 44 |
| 1 | 12936 | -44 | 24 | -2 | 0.048 | - | - | Area OP9 | 35 | HA | Left | FO | 52 |
| 1 | 12936 | -30 | 22 | -2 | 0.033 | - | - | Area Id7 | 28 | NHA | Left | IC | 56 |
| 1 | 12936 | -46 | 10 | 16 | 0.025 | - | - | Area 44 | 18 | NHA | Left | - | - |
| 2 | 2192 | -52 | -40 | 4 | 0.044 | - | - | - | - | NA | Left | - | - |
| 2 | 2192 | -60 | -50 | 8 | 0.029 | - | - | - | - | NA | Left | MTGt | 40 |
| 3 | 1640 | -44 | -44 | -20 | 0.037 | - | - | Area FG4 | 88 | HA | Left | TFCp | 31 |
| 3 | 1640 | -46 | -56 | -12 | 0.026 | - | - | Area FG4 | 72 | HA | Left | ITGt | 51 |
| 4 | 1392 | 42 | 20 | 22 | 0.035 | - | - | - | - | NA | Right | - | - |
| 4 | 1392 | 50 | 24 | 24 | 0.026 | - | - | Area 45 | 20 | NHA | Right | MFG | 28 |
| 1 | 13288 | -48 | 18 | 22 | 0.064 | - | - | Area 44 | 19 | NHA | Left | IFG POp | 51 |
| 1 | 13288 | -50 | 26 | -4 | 0.037 | - | - | Area OP9 | 54 | HA | Left | IFG PTr | 30 |
| 1 | 13288 | -52 | 38 | 4 | 0.025 | - | - | Area 45 | 23 | NHA | Left | FP | 50 |
| 1 | 13288 | -50 | 8 | 40 | 0.020 | - | - | Area 44 | 7 | NHA | Left | MFG | 42 |
| 1 | 13288 | -46 | 6 | 46 | 0.020 | - | - | Area 44 | 0 | NHA | Left | MFG | 50 |
| 2 | 5632 | -2 | 30 | 46 | 0.037 | - | - | - | - | NA | Left | SFG | 56 |
| 2 | 5632 | -2 | 18 | 52 | 0.036 | - | - | Area 6mr / preSMA | 10 | NHA | Left | SFG | 46 |
| 2 | 5632 | -4 | 16 | 56 | 0.035 | - | - | Area 6mr / preSMA | 56 | HA | Left | SFG | 68 |
| 2 | 5632 | -6 | 22 | 46 | 0.033 | - | - | - | - | NA | Left | PAC | 47 |
| 2 | 5632 | -2 | 28 | 34 | 0.028 | - | - | - | - | NA | Left | PAC | 70 |
| 3 | 1464 | -46 | -54 | -12 | 0.029 | - | - | Area FG4 | 81 | HA | Left | ITGt | 50 |
| 3 | 1464 | -54 | -50 | -4 | 0.025 | - | - | - | - | NA | Left | - | - |
| 3 | 1464 | -52 | -66 | -4 | 0.022 | - | - | Area hOc4la | 0 | NHA | Left | LOCi | 59 |

**Supplementary Table 7.**
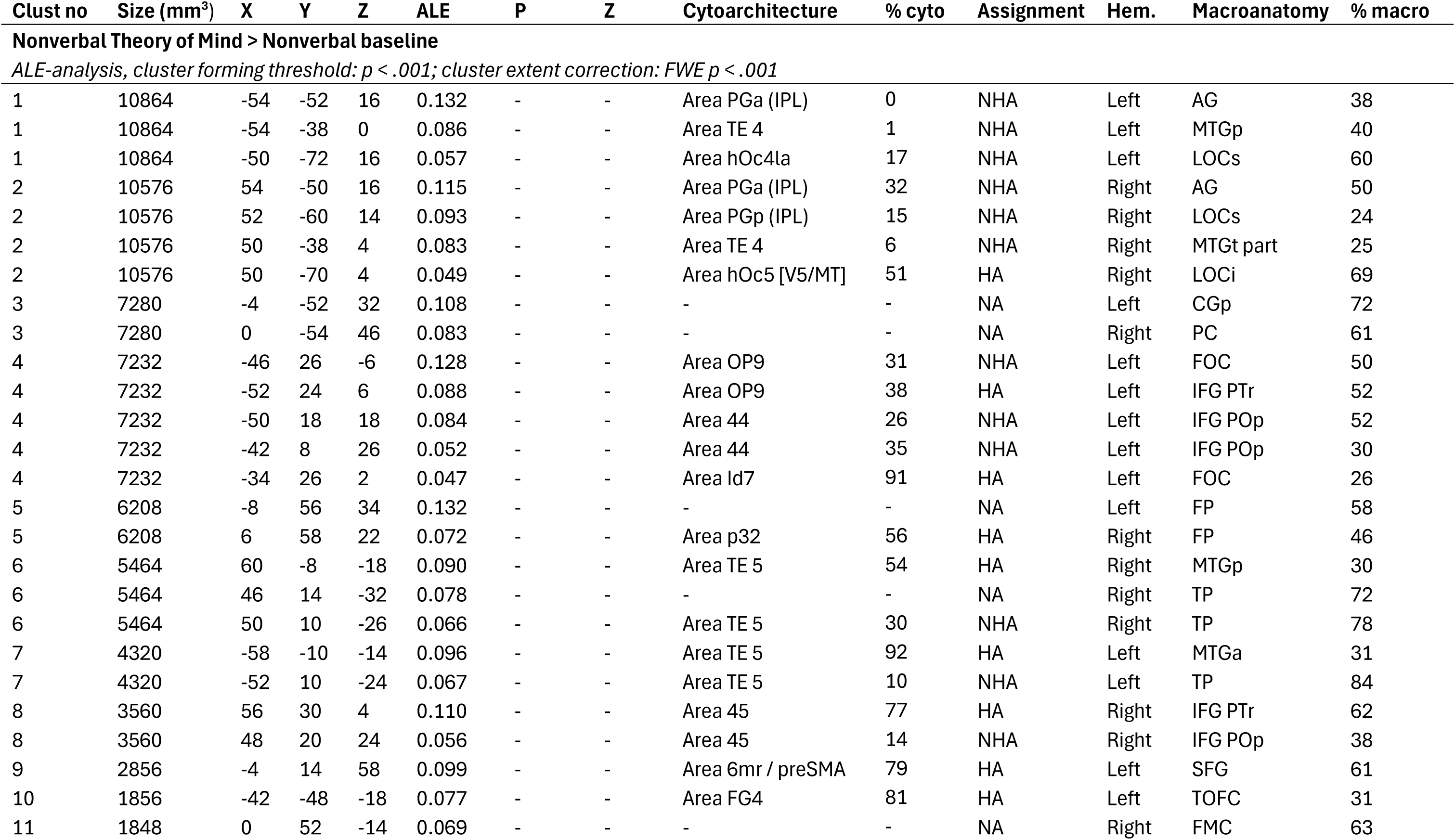
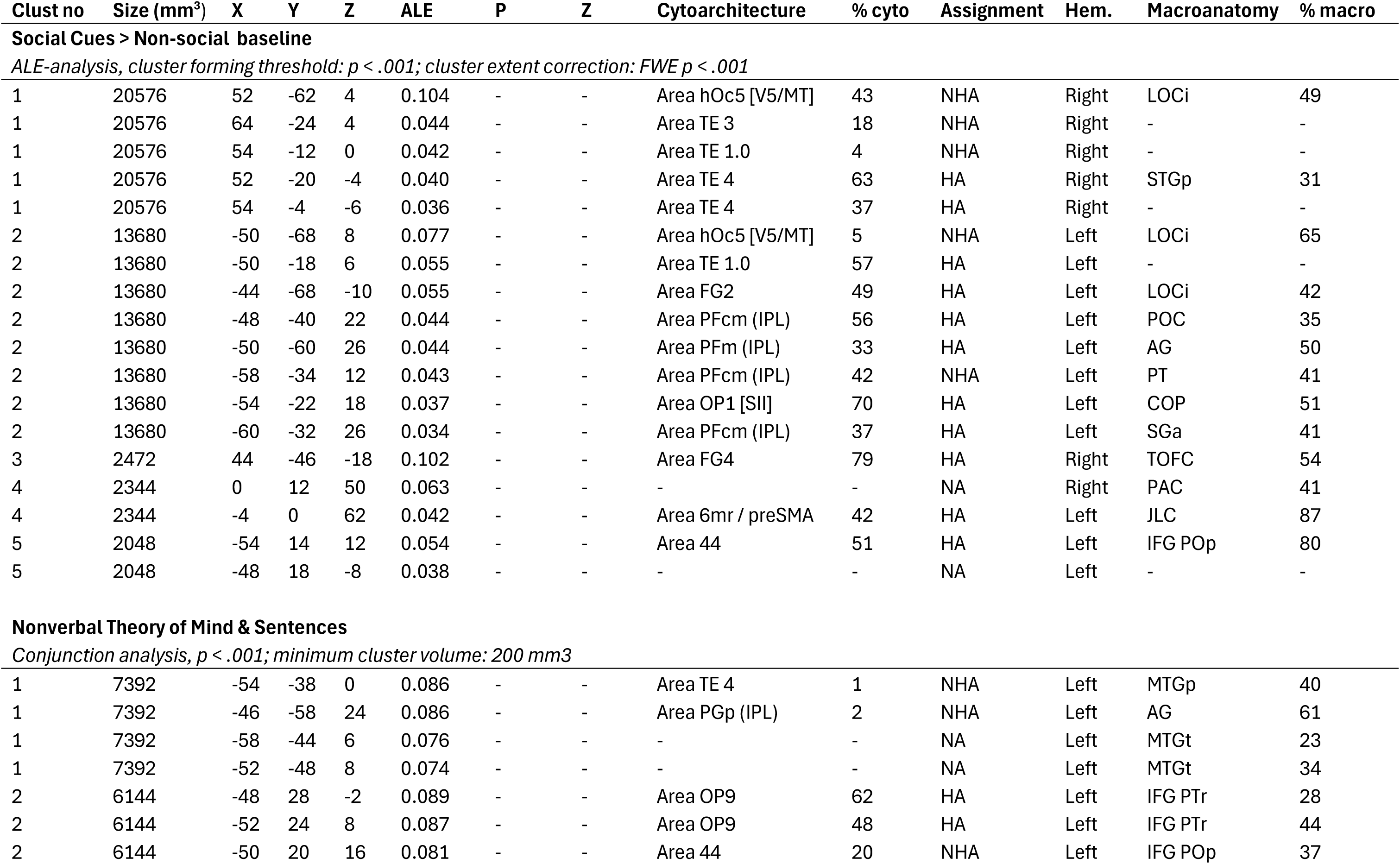
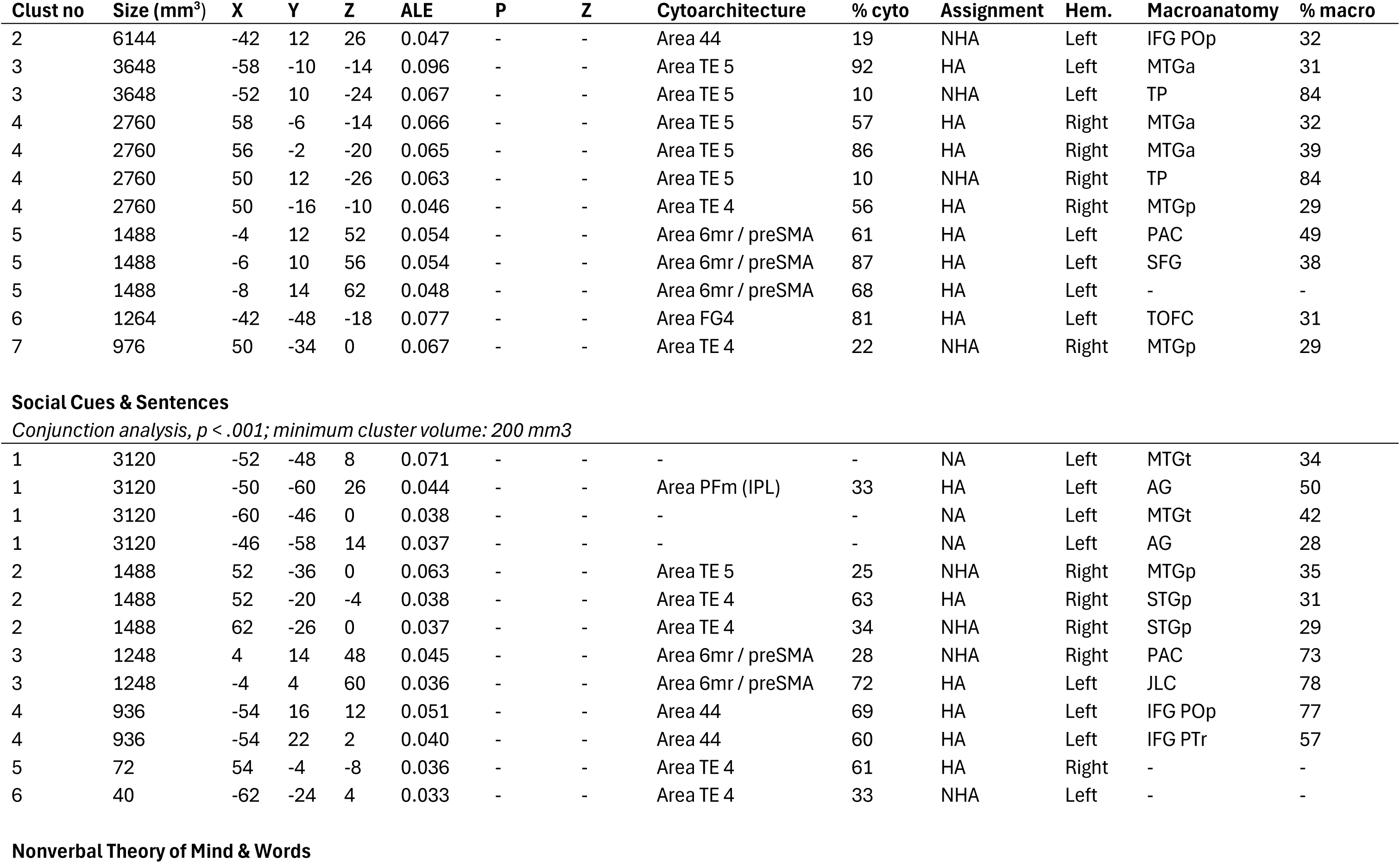
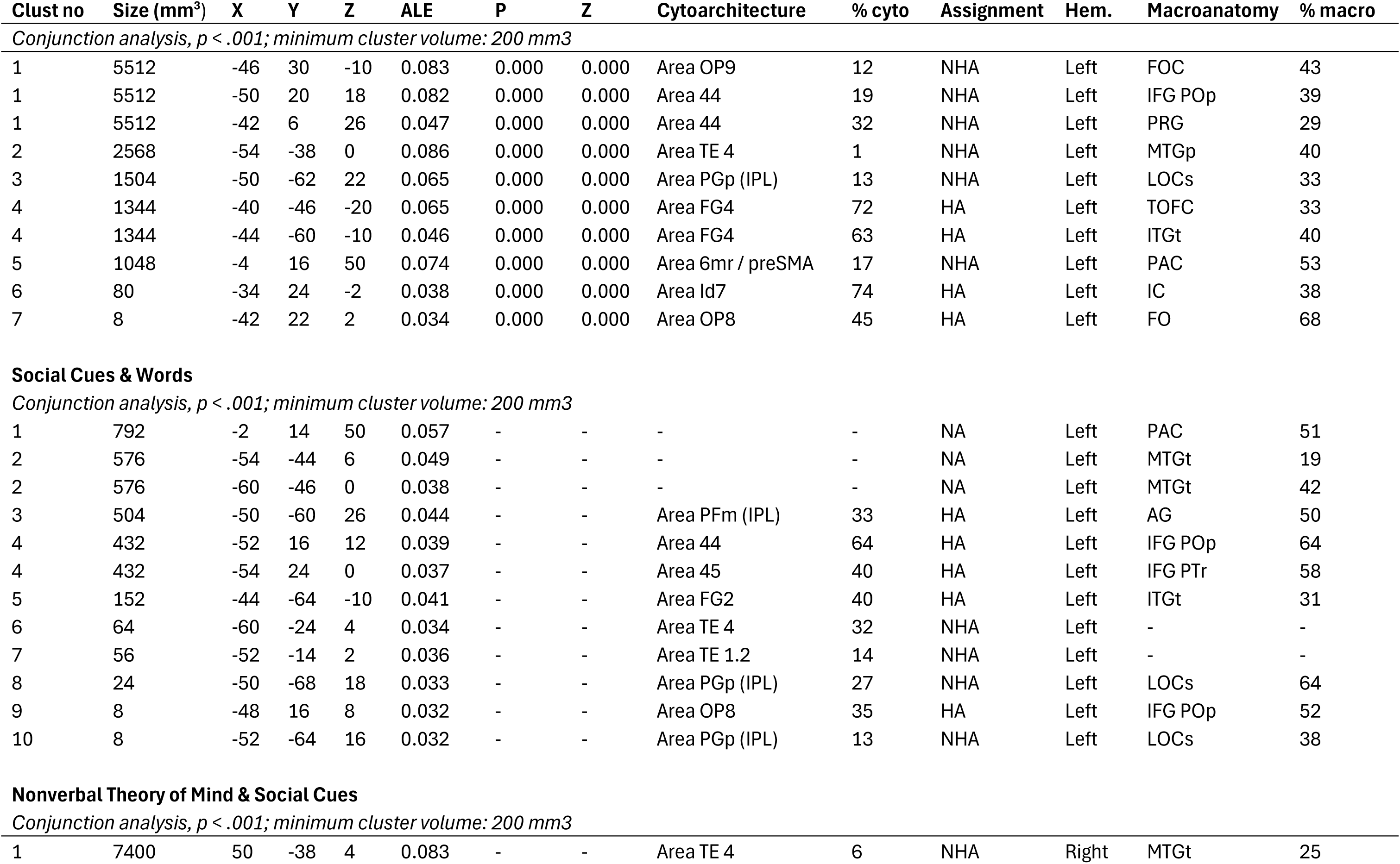
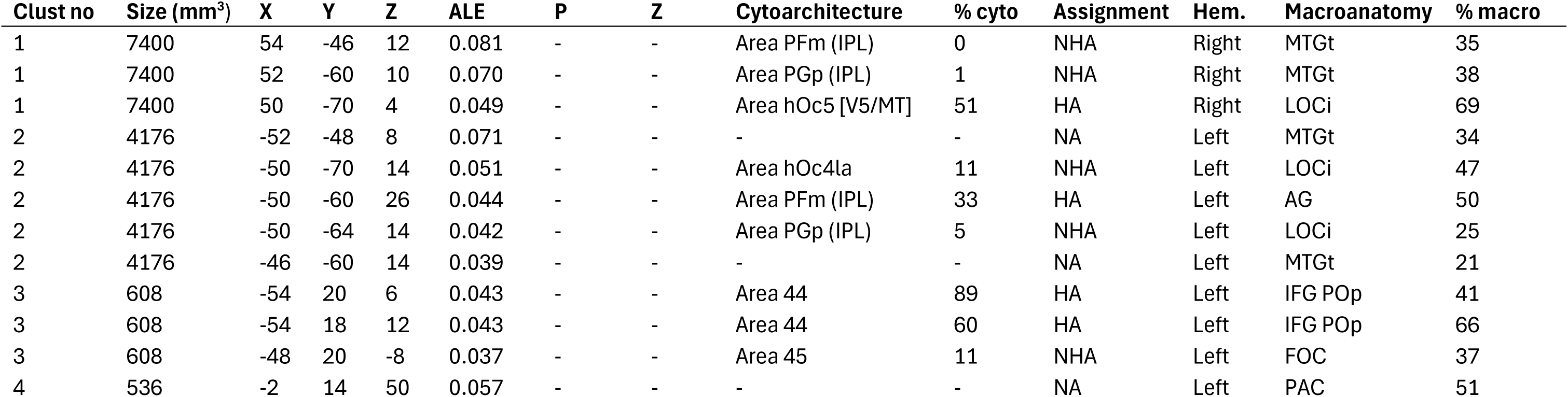
All activation clusters and local maxima for social processing, and specified verbal semantics. Coordinates x, y and z reported in the MNI coordinate system; “Cluster”: Cluster number in the individual contrast; “ALE”: Ale values output from Ginger ALE, along with P and Z values; “Cytoarchitecture”: cytoarchitectonic information for foci assigned by the JuBrain Anatomy Toolbox (SPM), based on the Maximum Probability Map; “% cyto”: probability of the coordinate falling into the specified Cytoarchitecture, as an output of the Anatomy Toolbox; “Assignment”: Type of assignment of coordinate into the specified Cytoarchitecture, as an output of the Anatomy Toolbox – HA: Hard Assignment, NHA: No Hard Assignment, NA: No Assignment “Hem”: hemisphere; “Macroanatomy”: Assignment of the foci and to the Harvard-Oxford microanatomical atlas; “% macro”: probability of the coordinate falling into the assigned region by the Harvard-Oxford microanatomical atlas; “aCG”: Cingulate Gyrus, anterior; “AG”: Angular Gyrus; “AMYG”: Amygdala; “COP”: Central Opercular Cortex; “CRcr-I”: Cerebellum Crus I; “CRcr-II”: Cerebellum Crus II; “FMC”: Frontal Medial Cortex; “FO”: Frontal Operculum Cortex; “FOC”: Frontal Orbital Cortex; “FP”: Frontal Pole; “HC”: Hippocampus; “HG”: Heschl’s Gyrus; “IC”: Insular Cortex; “IFG POp”: Inferior Frontal Gyrus, pars opercularis; “IFG PTr”: Inferior Frontal Gyrus, pars triangularis; “IFGt”: Inferior Frontal Gyrus, temporooccipital; “ITGp”: Inferior Temporal Gyrus, posterior; “ITGt”: Inferior Temporal Gyrus, temporooccipital; “JLC”: Juxtapositional Lobule Cortex; “LOCi”: Lateral Occipital Cortex, inferior; “LOCs”: Lateral Occipital Cortex, superior; “MFG”: Middle Frontal Gyrus; “MTGa”: Middle Temporal Gyrus, anterior; “MTGp”: Middle Temporal Gyrus, posterior; “MTGt”: Middle Temporal Gyrus, temporooccipital; “OFC”: Occipital Fusiform Gyrus; “OP”: Occipital Pole; “PAC”: Paracingulate Gyrus; “PC”: Precuneous Cortex; “pCG”: Cingulate Gyrus, posterior; “PGp”: Parahippocampal Gyrus, posterior; “POC”: Parietal Operculum Cortex; “PP”: Planum Polare; “PRG”: Precentral Gyrus; “PT”: Planum Temporale; “RC”: Right Caudate; “SFG”: Superior Frontal Gyrus; “SGp”: Supramarginal Gyrus, posterior; “SPL”: Superior Parietal Lobule; “STGa”: Superior Temporal Gyrus, anterior; “STGp”: Superior Temporal Gyrus, posterior; “STGs”: Superior Temporal Gyrus, superior; “TFCp”: Temporal Fusiform Cortex, posterior; “TOFC”: Temporal Occipital Fusiform Cortex; “TP”: Temporal Pole

| Clust no | Size (mm<sup>3</sup>) | X | Y | Z | ALE | P | Z | Cytoarchitecture | % cyto | Assignment | Hem. | Macroanatomy | % macro |
| --- | --- | --- | --- | --- | --- | --- | --- | --- | --- | --- | --- | --- | --- |
| **Nonverbal Theory of Mind > Nonverbal baseline** |  |  |  |  |  |  |  |  |  |  |  |  |  |
| *ALE-analysis, cluster forming threshold: $p < .001$; cluster extent correction: FWE $p < .001$* |  |  |  |  |  |  |  |  |  |  |  |  |  |
| 1 | 10864 | -54 | -52 | 16 | 0.132 | - | - | Area PGa (IPL) | 0 | NHA | Left | AG | 38 |
| 1 | 10864 | -54 | -38 | 0 | 0.086 | - | - | Area TE 4 | 1 | NHA | Left | MTGp | 40 |
| 1 | 10864 | -50 | -72 | 16 | 0.057 | - | - | Area hOc4la | 17 | NHA | Left | LOCs | 60 |
| 2 | 10576 | 54 | -50 | 16 | 0.115 | - | - | Area PGa (IPL) | 32 | NHA | Right | AG | 50 |
| 2 | 10576 | 52 | -60 | 14 | 0.093 | - | - | Area PGp (IPL) | 15 | NHA | Right | LOCs | 24 |
| 2 | 10576 | 50 | -38 | 4 | 0.083 | - | - | Area TE 4 | 6 | NHA | Right | MTGt part | 25 |
| 2 | 10576 | 50 | -70 | 4 | 0.049 | - | - | Area hOc5 [V5/MT] | 51 | HA | Right | LOCi | 69 |
| 3 | 7280 | -4 | -52 | 32 | 0.108 | - | - | - | - | NA | Left | CGp | 72 |
| 3 | 7280 | 0 | -54 | 46 | 0.083 | - | - | - | - | NA | Right | PC | 61 |
| 4 | 7232 | -46 | 26 | -6 | 0.128 | - | - | Area OP9 | 31 | NHA | Left | FOC | 50 |
| 4 | 7232 | -52 | 24 | 6 | 0.088 | - | - | Area OP9 | 38 | HA | Left | IFG PTr | 52 |
| 4 | 7232 | -50 | 18 | 18 | 0.084 | - | - | Area 44 | 26 | NHA | Left | IFG POp | 52 |
| 4 | 7232 | -42 | 8 | 26 | 0.052 | - | - | Area 44 | 35 | NHA | Left | IFG POp | 30 |
| 4 | 7232 | -34 | 26 | 2 | 0.047 | - | - | Area Id7 | 91 | HA | Left | FOC | 26 |
| 5 | 6208 | -8 | 56 | 34 | 0.132 | - | - | - | - | NA | Left | FP | 58 |
| 5 | 6208 | 6 | 58 | 22 | 0.072 | - | - | Area p32 | 56 | HA | Right | FP | 46 |
| 6 | 5464 | 60 | -8 | -18 | 0.090 | - | - | Area TE 5 | 54 | HA | Right | MTGp | 30 |
| 6 | 5464 | 46 | 14 | -32 | 0.078 | - | - | - | - | NA | Right | TP | 72 |
| 6 | 5464 | 50 | 10 | -26 | 0.066 | - | - | Area TE 5 | 30 | NHA | Right | TP | 78 |
| 7 | 4320 | -58 | -10 | -14 | 0.096 | - | - | Area TE 5 | 92 | HA | Left | MTGa | 31 |
| 7 | 4320 | -52 | 10 | -24 | 0.067 | - | - | Area TE 5 | 10 | NHA | Left | TP | 84 |
| 8 | 3560 | 56 | 30 | 4 | 0.110 | - | - | Area 45 | 77 | HA | Right | IFG PTr | 62 |
| 8 | 3560 | 48 | 20 | 24 | 0.056 | - | - | Area 45 | 14 | NHA | Right | IFG POp | 38 |
| 9 | 2856 | -4 | 14 | 58 | 0.099 | - | - | Area 6mr / preSMA | 79 | HA | Left | SFG | 61 |
| 10 | 1856 | -42 | -48 | -18 | 0.077 | - | - | Area FG4 | 81 | HA | Left | TOFC | 31 |
| 11 | 1848 | 0 | 52 | -14 | 0.069 | - | - | - | - | NA | Right | FMC | 63 |
| **Social Cues > Non-social baseline** |  |  |  |  |  |  |  |  |  |  |  |  |  |
| *ALE-analysis, cluster forming threshold: $p < .001$; cluster extent correction: FWE $p < .001$* |  |  |  |  |  |  |  |  |  |  |  |  |  |
| 1 | 20576 | 52 | -62 | 4 | 0.104 | - | - | Area hOc5 [V5/MT] | 43 | NHA | Right | LOCi | 49 |
| 1 | 20576 | 64 | -24 | 4 | 0.044 | - | - | Area TE 3 | 18 | NHA | Right | - | - |
| 1 | 20576 | 54 | -12 | 0 | 0.042 | - | - | Area TE 1.0 | 4 | NHA | Right | - | - |
| 1 | 20576 | 52 | -20 | -4 | 0.040 | - | - | Area TE 4 | 63 | HA | Right | STGp | 31 |
| 1 | 20576 | 54 | -4 | -6 | 0.036 | - | - | Area TE 4 | 37 | HA | Right | - | - |
| 2 | 13680 | -50 | -68 | 8 | 0.077 | - | - | Area hOc5 [V5/MT] | 5 | NHA | Left | LOCi | 65 |
| 2 | 13680 | -50 | -18 | 6 | 0.055 | - | - | Area TE 1.0 | 57 | HA | Left | - | - |
| 2 | 13680 | -44 | -68 | -10 | 0.055 | - | - | Area FG2 | 49 | HA | Left | LOCi | 42 |
| 2 | 13680 | -48 | -40 | 22 | 0.044 | - | - | Area PFcm (IPL) | 56 | HA | Left | POC | 35 |
| 2 | 13680 | -50 | -60 | 26 | 0.044 | - | - | Area PFm (IPL) | 33 | HA | Left | AG | 50 |
| 2 | 13680 | -58 | -34 | 12 | 0.043 | - | - | Area PFcm (IPL) | 42 | NHA | Left | PT | 41 |
| 2 | 13680 | -54 | -22 | 18 | 0.037 | - | - | Area OP1 [SII] | 70 | HA | Left | COP | 51 |
| 2 | 13680 | -60 | -32 | 26 | 0.034 | - | - | Area PFcm (IPL) | 37 | HA | Left | SGa | 41 |
| 3 | 2472 | 44 | -46 | -18 | 0.102 | - | - | Area FG4 | 79 | HA | Right | TOFC | 54 |
| 4 | 2344 | 0 | 12 | 50 | 0.063 | - | - | - | - | NA | Right | PAC | 41 |
| 4 | 2344 | -4 | 0 | 62 | 0.042 | - | - | Area 6mr / preSMA | 42 | HA | Left | JLC | 87 |
| 5 | 2048 | -54 | 14 | 12 | 0.054 | - | - | Area 44 | 51 | HA | Left | IFG POp | 80 |
| 5 | 2048 | -48 | 18 | -8 | 0.038 | - | - | - | - | NA | Left | - | - |
| **Nonverbal Theory of Mind & Sentences** |  |  |  |  |  |  |  |  |  |  |  |  |  |
| *Conjunction analysis, $p < .001$; minimum cluster volume: 200 mm<sup>3</sup>* |  |  |  |  |  |  |  |  |  |  |  |  |  |
| 1 | 7392 | -54 | -38 | 0 | 0.086 | - | - | Area TE 4 | 1 | NHA | Left | MTGp | 40 |
| 1 | 7392 | -46 | -58 | 24 | 0.086 | - | - | Area PGp (IPL) | 2 | NHA | Left | AG | 61 |
| 1 | 7392 | -58 | -44 | 6 | 0.076 | - | - | - | - | NA | Left | MTGt | 23 |
| 1 | 7392 | -52 | -48 | 8 | 0.074 | - | - | - | - | NA | Left | MTGt | 34 |
| 2 | 6144 | -48 | 28 | -2 | 0.089 | - | - | Area OP9 | 62 | HA | Left | IFG PTr | 28 |
| 2 | 6144 | -52 | 24 | 8 | 0.087 | - | - | Area OP9 | 48 | HA | Left | IFG PTr | 44 |
| 2 | 6144 | -50 | 20 | 16 | 0.081 | - | - | Area 44 | 20 | NHA | Left | IFG POp | 37 |
| 2 | 6144 | -42 | 12 | 26 | 0.047 | - | - | Area 44 | 19 | NHA | Left | IFG POp | 32 |
| 3 | 3648 | -58 | -10 | -14 | 0.096 | - | - | Area TE 5 | 92 | HA | Left | MTGa | 31 |
| 3 | 3648 | -52 | 10 | -24 | 0.067 | - | - | Area TE 5 | 10 | NHA | Left | TP | 84 |
| 4 | 2760 | 58 | -6 | -14 | 0.066 | - | - | Area TE 5 | 57 | HA | Right | MTGa | 32 |
| 4 | 2760 | 56 | -2 | -20 | 0.065 | - | - | Area TE 5 | 86 | HA | Right | MTGa | 39 |
| 4 | 2760 | 50 | 12 | -26 | 0.063 | - | - | Area TE 5 | 10 | NHA | Right | TP | 84 |
| 4 | 2760 | 50 | -16 | -10 | 0.046 | - | - | Area TE 4 | 56 | HA | Right | MTGp | 29 |
| 5 | 1488 | -4 | 12 | 52 | 0.054 | - | - | Area 6mr / preSMA | 61 | HA | Left | PAC | 49 |
| 5 | 1488 | -6 | 10 | 56 | 0.054 | - | - | Area 6mr / preSMA | 87 | HA | Left | SFG | 38 |
| 5 | 1488 | -8 | 14 | 62 | 0.048 | - | - | Area 6mr / preSMA | 68 | HA | Left | - | - |
| 6 | 1264 | -42 | -48 | -18 | 0.077 | - | - | Area FG4 | 81 | HA | Left | TOFC | 31 |
| 7 | 976 | 50 | -34 | 0 | 0.067 | - | - | Area TE 4 | 22 | NHA | Right | MTGp | 29 |
| 1 | 3120 | -52 | -48 | 8 | 0.071 | - | - | - | - | NA | Left | MTGt | 34 |
| 1 | 3120 | -50 | -60 | 26 | 0.044 | - | - | Area PFm (IPL) | 33 | HA | Left | AG | 50 |
| 1 | 3120 | -60 | -46 | 0 | 0.038 | - | - | - | - | NA | Left | MTGt | 42 |
| 1 | 3120 | -46 | -58 | 14 | 0.037 | - | - | - | - | NA | Left | AG | 28 |
| 2 | 1488 | 52 | -36 | 0 | 0.063 | - | - | Area TE 5 | 25 | NHA | Right | MTGp | 35 |
| 2 | 1488 | 52 | -20 | -4 | 0.038 | - | - | Area TE 4 | 63 | HA | Right | STGp | 31 |
| 2 | 1488 | 62 | -26 | 0 | 0.037 | - | - | Area TE 4 | 34 | NHA | Right | STGp | 29 |
| 3 | 1248 | 4 | 14 | 48 | 0.045 | - | - | Area 6mr / preSMA | 28 | NHA | Right | PAC | 73 |
| 3 | 1248 | -4 | 4 | 60 | 0.036 | - | - | Area 6mr / preSMA | 72 | HA | Left | JLC | 78 |
| 4 | 936 | -54 | 16 | 12 | 0.051 | - | - | Area 44 | 69 | HA | Left | IFG POp | 77 |
| 4 | 936 | -54 | 22 | 2 | 0.040 | - | - | Area 44 | 60 | HA | Left | IFG PTr | 57 |
| 5 | 72 | 54 | -4 | -8 | 0.036 | - | - | Area TE 4 | 61 | HA | Right | - | - |
| 6 | 40 | -62 | -24 | 4 | 0.033 | - | - | Area TE 4 | 33 | NHA | Left | - | - |

Nonverbal Theory of Mind & Words
| Clust no | Size (mm<sup>3</sup>) | X | Y | Z | ALE | P | Z | Cytoarchitecture | % cyto | Assignment | Hem. | Macroanatomy | % macro |
| --- | --- | --- | --- | --- | --- | --- | --- | --- | --- | --- | --- | --- | --- |
| *Conjunction analysis, $p < .001$; minimum cluster volume: 200 mm<sup>3</sup>* |  |  |  |  |  |  |  |  |  |  |  |  |  |
| 1 | 5512 | -46 | 30 | -10 | 0.083 | 0.000 | 0.000 | Area OP9 | 12 | NHA | Left | FOC | 43 |
| 1 | 5512 | -50 | 20 | 18 | 0.082 | 0.000 | 0.000 | Area 44 | 19 | NHA | Left | IFG POp | 39 |
| 1 | 5512 | -42 | 6 | 26 | 0.047 | 0.000 | 0.000 | Area 44 | 32 | NHA | Left | PRG | 29 |
| 2 | 2568 | -54 | -38 | 0 | 0.086 | 0.000 | 0.000 | Area TE 4 | 1 | NHA | Left | MTGp | 40 |
| 3 | 1504 | -50 | -62 | 22 | 0.065 | 0.000 | 0.000 | Area PGp (IPL) | 13 | NHA | Left | LOCs | 33 |
| 4 | 1344 | -40 | -46 | -20 | 0.065 | 0.000 | 0.000 | Area FG4 | 72 | HA | Left | TOFC | 33 |
| 4 | 1344 | -44 | -60 | -10 | 0.046 | 0.000 | 0.000 | Area FG4 | 63 | HA | Left | ITGt | 40 |
| 5 | 1048 | -4 | 16 | 50 | 0.074 | 0.000 | 0.000 | Area 6mr / preSMA | 17 | NHA | Left | PAC | 53 |
| 6 | 80 | -34 | 24 | -2 | 0.038 | 0.000 | 0.000 | Area Id7 | 74 | HA | Left | IC | 38 |
| 7 | 8 | -42 | 22 | 2 | 0.034 | 0.000 | 0.000 | Area OP8 | 45 | HA | Left | FO | 68 |
| 1 | 792 | -2 | 14 | 50 | 0.057 | - | - | - | - | NA | Left | PAC | 51 |
| 2 | 576 | -54 | -44 | 6 | 0.049 | - | - | - | - | NA | Left | MTGt | 19 |
| 2 | 576 | -60 | -46 | 0 | 0.038 | - | - | - | - | NA | Left | MTGt | 42 |
| 3 | 504 | -50 | -60 | 26 | 0.044 | - | - | Area PFm (IPL) | 33 | HA | Left | AG | 50 |
| 4 | 432 | -52 | 16 | 12 | 0.039 | - | - | Area 44 | 64 | HA | Left | IFG POp | 64 |
| 4 | 432 | -54 | 24 | 0 | 0.037 | - | - | Area 45 | 40 | HA | Left | IFG PTr | 58 |
| 5 | 152 | -44 | -64 | -10 | 0.041 | - | - | Area FG2 | 40 | HA | Left | ITGt | 31 |
| 6 | 64 | -60 | -24 | 4 | 0.034 | - | - | Area TE 4 | 32 | NHA | Left | - | - |
| 7 | 56 | -52 | -14 | 2 | 0.036 | - | - | Area TE 1.2 | 14 | NHA | Left | - | - |
| 8 | 24 | -50 | -68 | 18 | 0.033 | - | - | Area PGp (IPL) | 27 | NHA | Left | LOCs | 64 |
| 9 | 8 | -48 | 16 | 8 | 0.032 | - | - | Area OP8 | 35 | HA | Left | IFG POp | 52 |
| 10 | 8 | -52 | -64 | 16 | 0.032 | - | - | Area PGp (IPL) | 13 | NHA | Left | LOCs | 38 |
| 1 | 7400 | 50 | -38 | 4 | 0.083 | - | - | Area TE 4 | 6 | NHA | Right | MTGt | 25 |
| 1 | 7400 | 54 | -46 | 12 | 0.081 | - | - | Area PFm (IPL) | 0 | NHA | Right | MTGt | 35 |
| 1 | 7400 | 52 | -60 | 10 | 0.070 | - | - | Area PGp (IPL) | 1 | NHA | Right | MTGt | 38 |
| 1 | 7400 | 50 | -70 | 4 | 0.049 | - | - | Area hOc5 [V5/MT] | 51 | HA | Right | LOCi | 69 |
| 2 | 4176 | -52 | -48 | 8 | 0.071 | - | - | - | - | NA | Left | MTGt | 34 |
| 2 | 4176 | -50 | -70 | 14 | 0.051 | - | - | Area hOc4la | 11 | NHA | Left | LOCi | 47 |
| 2 | 4176 | -50 | -60 | 26 | 0.044 | - | - | Area PFm (IPL) | 33 | HA | Left | AG | 50 |
| 2 | 4176 | -50 | -64 | 14 | 0.042 | - | - | Area PGp (IPL) | 5 | NHA | Left | LOCi | 25 |
| 2 | 4176 | -46 | -60 | 14 | 0.039 | - | - | - | - | NA | Left | MTGt | 21 |
| 3 | 608 | -54 | 20 | 6 | 0.043 | - | - | Area 44 | 89 | HA | Left | IFG POp | 41 |
| 3 | 608 | -54 | 18 | 12 | 0.043 | - | - | Area 44 | 60 | HA | Left | IFG POp | 66 |
| 3 | 608 | -48 | 20 | -8 | 0.037 | - | - | Area 45 | 11 | NHA | Left | FOC | 37 |
| 4 | 536 | -2 | 14 | 50 | 0.057 | - | - | - | - | NA | Left | PAC | 51 |

**Supplementary Table 8.**
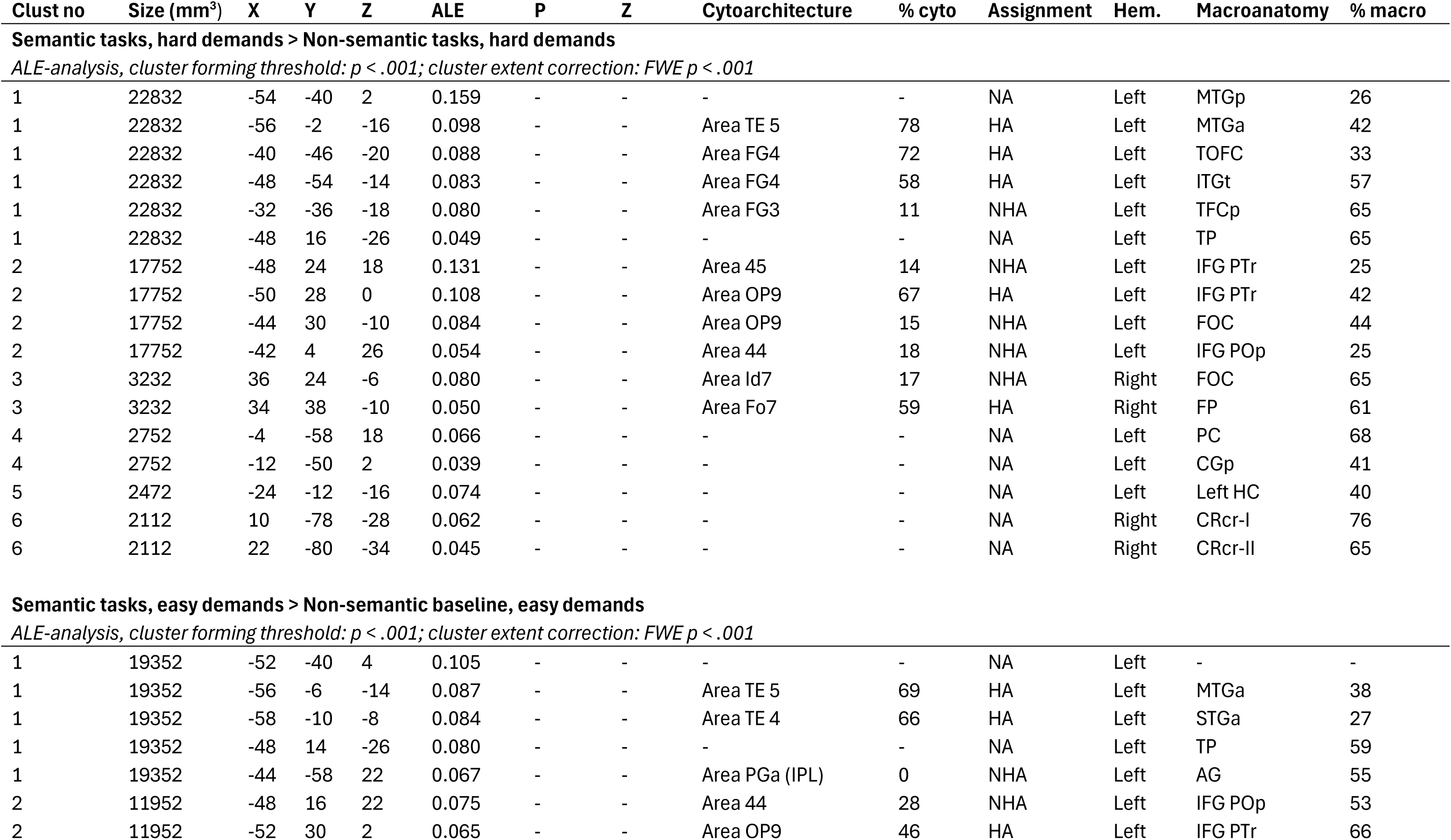
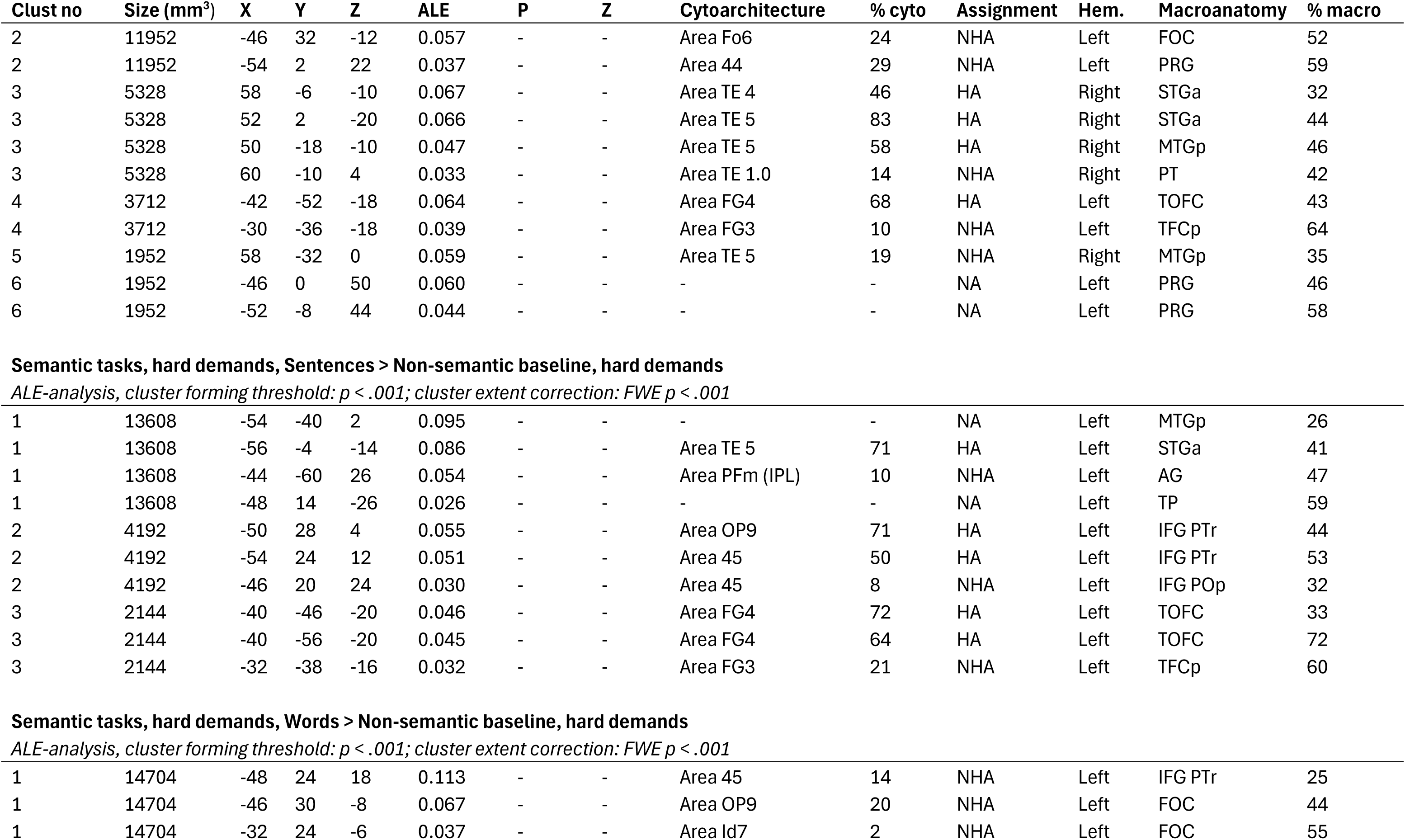
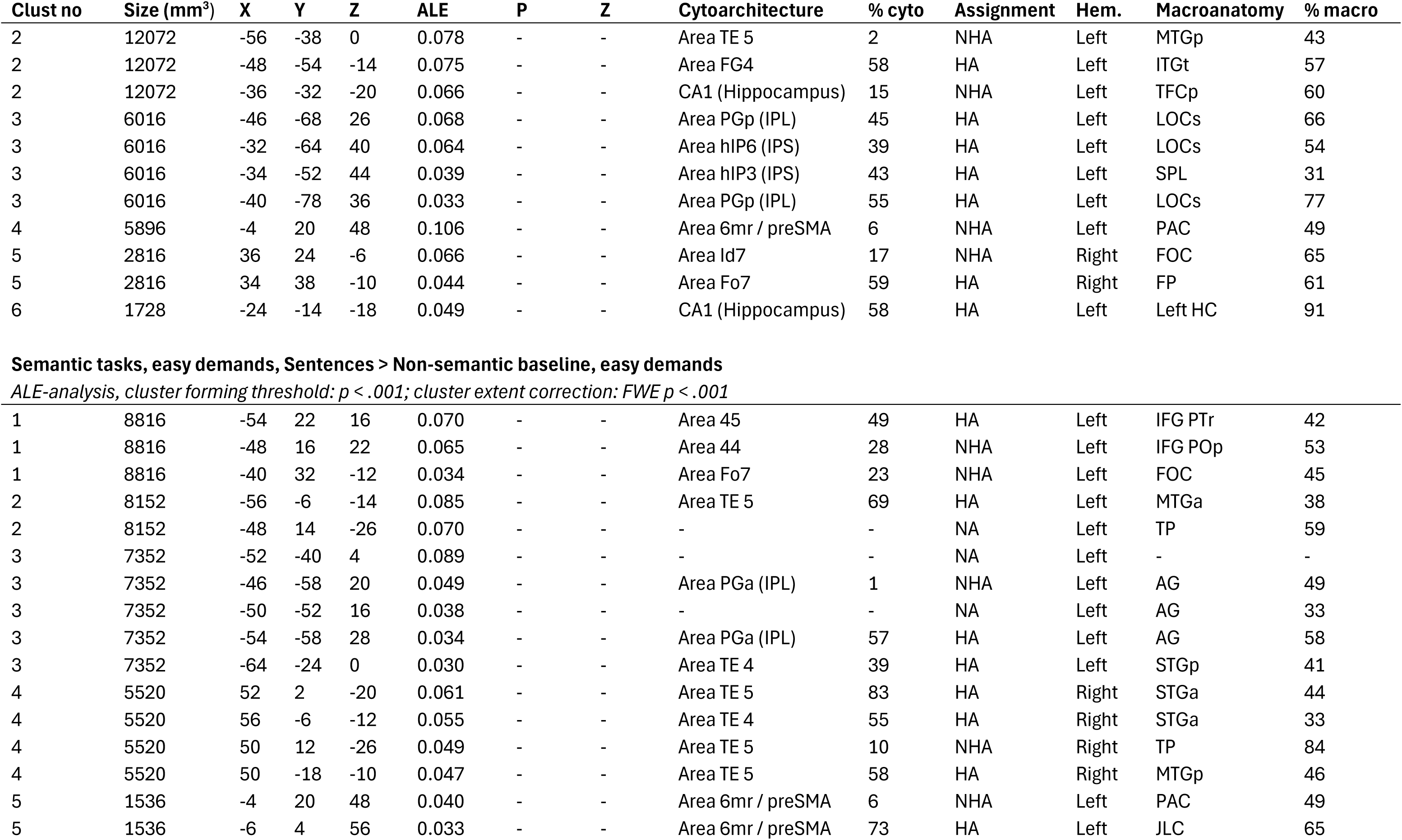
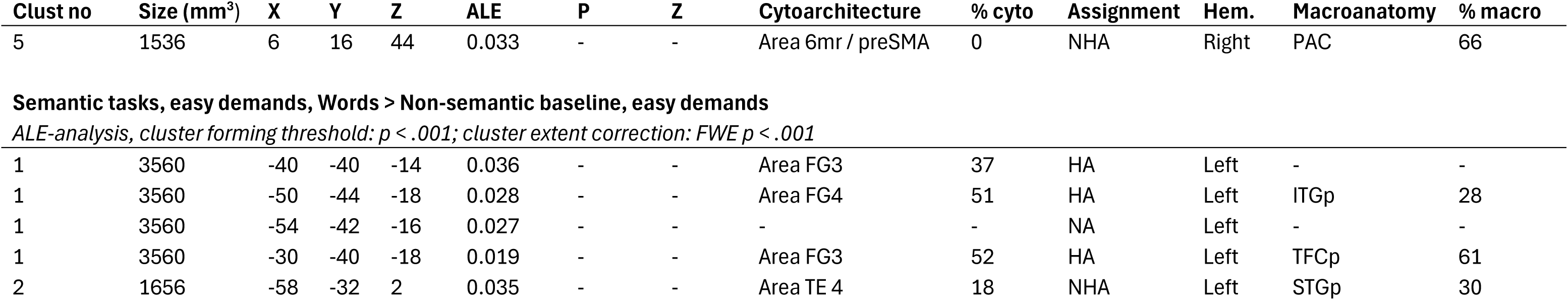
All activation clusters and local maxima for verbal semantic cognition, analysed based on external task demands. Coordinates x, y and z reported in the MNI coordinate system; “Cluster”: Cluster number in the individual contrast; “ALE”: Ale values output from Ginger ALE, along with P and Z values; “Cytoarchitecture”: cytoarchitectonic information for foci assigned by the JuBrain Anatomy Toolbox (SPM), based on the Maximum Probability Map; “% cyto”: probability of the coordinate falling into the specified Cytoarchitecture, as an output of the Anatomy Toolbox; “Assignment”: Type of assignment of coordinate into the specified Cytoarchitecture, as an output of the Anatomy Toolbox – HA: Hard Assignment, NHA: No Hard Assignment, NA: No Assignment “Hem”: hemisphere; “Macroanatomy”: Assignment of the foci and to the Harvard-Oxford microanatomical atlas; “% macro”: probability of the coordinate falling into the assigned region by the Harvard-Oxford microanatomical atlas; “aCG”: Cingulate Gyrus, anterior; “AG”: Angular Gyrus; “AMYG”: Amygdala; “COP”: Central Opercular Cortex; “CRcr-I”: Cerebellum Crus I; “CRcr-II”: Cerebellum Crus II; “FMC”: Frontal Medial Cortex; “FO”: Frontal Operculum Cortex; “FOC”: Frontal Orbital Cortex; “FP”: Frontal Pole; “HC”: Hippocampus; “HG”: Heschl’s Gyrus; “IC”: Insular Cortex; “IFG POp”: Inferior Frontal Gyrus, pars opercularis; “IFG PTr”: Inferior Frontal Gyrus, pars triangularis; “IFGt”: Inferior Frontal Gyrus, temporooccipital; “ITGp”: Inferior Temporal Gyrus, posterior; “ITGt”: Inferior Temporal Gyrus, temporooccipital; “JLC”: Juxtapositional Lobule Cortex; “LOCi”: Lateral Occipital Cortex, inferior; “LOCs”: Lateral Occipital Cortex, superior; “MFG”: Middle Frontal Gyrus; “MTGa”: Middle Temporal Gyrus, anterior; “MTGp”: Middle Temporal Gyrus, posterior; “MTGt”: Middle Temporal Gyrus, temporooccipital; “OFC”: Occipital Fusiform Gyrus; “OP”: Occipital Pole; “PAC”: Paracingulate Gyrus; “PC”: Precuneous Cortex; “pCG”: Cingulate Gyrus, posterior; “PGp”: Parahippocampal Gyrus, posterior; “POC”: Parietal Operculum Cortex; “PP”: Planum Polare; “PRG”: Precentral Gyrus; “PT”: Planum Temporale; “RC”: Right Caudate; “SFG”: Superior Frontal Gyrus; “SGp”: Supramarginal Gyrus, posterior; “SPL”: Superior Parietal Lobule; “STGa”: Superior Temporal Gyrus, anterior; “STGp”: Superior Temporal Gyrus, posterior; “STGs”: Superior Temporal Gyrus, superior; “TFCp”: Temporal Fusiform Cortex, posterior; “TOFC”: Temporal Occipital Fusiform Cortex; “TP”: Temporal Pole

| Clust no | Size (mm<sup>3</sup>) | X | Y | Z | ALE | P | Z | Cytoarchitecture | % cyto | Assignment | Hem. | Macroanatomy | % macro |
| --- | --- | --- | --- | --- | --- | --- | --- | --- | --- | --- | --- | --- | --- |
| **Semantic tasks, hard demands > Non-semantic tasks, hard demands** |  |  |  |  |  |  |  |  |  |  |  |  |  |
| *ALE-analysis, cluster forming threshold: $p < .001$; cluster extent correction: FWE $p < .001$* |  |  |  |  |  |  |  |  |  |  |  |  |  |
| 1 | 22832 | -54 | -40 | 2 | 0.159 | - | - | - | - | NA | Left | MTGp | 26 |
| 1 | 22832 | -56 | -2 | -16 | 0.098 | - | - | Area TE 5 | 78 | HA | Left | MTGa | 42 |
| 1 | 22832 | -40 | -46 | -20 | 0.088 | - | - | Area FG4 | 72 | HA | Left | TOFC | 33 |
| 1 | 22832 | -48 | -54 | -14 | 0.083 | - | - | Area FG4 | 58 | HA | Left | ITGt | 57 |
| 1 | 22832 | -32 | -36 | -18 | 0.080 | - | - | Area FG3 | 11 | NHA | Left | TFCp | 65 |
| 1 | 22832 | -48 | 16 | -26 | 0.049 | - | - | - | - | NA | Left | TP | 65 |
| 2 | 17752 | -48 | 24 | 18 | 0.131 | - | - | Area 45 | 14 | NHA | Left | IFG PTr | 25 |
| 2 | 17752 | -50 | 28 | 0 | 0.108 | - | - | Area OP9 | 67 | HA | Left | IFG PTr | 42 |
| 2 | 17752 | -44 | 30 | -10 | 0.084 | - | - | Area OP9 | 15 | NHA | Left | FOC | 44 |
| 2 | 17752 | -42 | 4 | 26 | 0.054 | - | - | Area 44 | 18 | NHA | Left | IFG POp | 25 |
| 3 | 3232 | 36 | 24 | -6 | 0.080 | - | - | Area Id7 | 17 | NHA | Right | FOC | 65 |
| 3 | 3232 | 34 | 38 | -10 | 0.050 | - | - | Area Fo7 | 59 | HA | Right | FP | 61 |
| 4 | 2752 | -4 | -58 | 18 | 0.066 | - | - | - | - | NA | Left | PC | 68 |
| 4 | 2752 | -12 | -50 | 2 | 0.039 | - | - | - | - | NA | Left | CGp | 41 |
| 5 | 2472 | -24 | -12 | -16 | 0.074 | - | - | - | - | NA | Left | Left HC | 40 |
| 6 | 2112 | 10 | -78 | -28 | 0.062 | - | - | - | - | NA | Right | CRcr-I | 76 |
| 6 | 2112 | 22 | -80 | -34 | 0.045 | - | - | - | - | NA | Right | CRcr-II | 65 |
| **Semantic tasks, easy demands > Non-semantic baseline, easy demands** |  |  |  |  |  |  |  |  |  |  |  |  |  |
| *ALE-analysis, cluster forming threshold: $p < .001$; cluster extent correction: FWE $p < .001$* |  |  |  |  |  |  |  |  |  |  |  |  |  |
| 1 | 19352 | -52 | -40 | 4 | 0.105 | - | - | - | - | NA | Left | - | - |
| 1 | 19352 | -56 | -6 | -14 | 0.087 | - | - | Area TE 5 | 69 | HA | Left | MTGa | 38 |
| 1 | 19352 | -58 | -10 | -8 | 0.084 | - | - | Area TE 4 | 66 | HA | Left | STGa | 27 |
| 1 | 19352 | -48 | 14 | -26 | 0.080 | - | - | - | - | NA | Left | TP | 59 |
| 1 | 19352 | -44 | -58 | 22 | 0.067 | - | - | Area PGa (IPL) | 0 | NHA | Left | AG | 55 |
| 2 | 11952 | -48 | 16 | 22 | 0.075 | - | - | Area 44 | 28 | NHA | Left | IFG POp | 53 |
| 2 | 11952 | -52 | 30 | 2 | 0.065 | - | - | Area OP9 | 46 | HA | Left | IFG PTr | 66 |
| 2 | 11952 | -46 | 32 | -12 | 0.057 | - | - | Area Fo6 | 24 | NHA | Left | FOC | 52 |
| 2 | 11952 | -54 | 2 | 22 | 0.037 | - | - | Area 44 | 29 | NHA | Left | PRG | 59 |
| 3 | 5328 | 58 | -6 | -10 | 0.067 | - | - | Area TE 4 | 46 | HA | Right | STGa | 32 |
| 3 | 5328 | 52 | 2 | -20 | 0.066 | - | - | Area TE 5 | 83 | HA | Right | STGa | 44 |
| 3 | 5328 | 50 | -18 | -10 | 0.047 | - | - | Area TE 5 | 58 | HA | Right | MTGp | 46 |
| 3 | 5328 | 60 | -10 | 4 | 0.033 | - | - | Area TE 1.0 | 14 | NHA | Right | PT | 42 |
| 4 | 3712 | -42 | -52 | -18 | 0.064 | - | - | Area FG4 | 68 | HA | Left | TOFC | 43 |
| 4 | 3712 | -30 | -36 | -18 | 0.039 | - | - | Area FG3 | 10 | NHA | Left | TFCp | 64 |
| 5 | 1952 | 58 | -32 | 0 | 0.059 | - | - | Area TE 5 | 19 | NHA | Right | MTGp | 35 |
| 6 | 1952 | -46 | 0 | 50 | 0.060 | - | - | - | - | NA | Left | PRG | 46 |
| 6 | 1952 | -52 | -8 | 44 | 0.044 | - | - | - | - | NA | Left | PRG | 58 |
| 1 | 13608 | -54 | -40 | 2 | 0.095 | - | - | - | - | NA | Left | MTGp | 26 |
| 1 | 13608 | -56 | -4 | -14 | 0.086 | - | - | Area TE 5 | 71 | HA | Left | STGa | 41 |
| 1 | 13608 | -44 | -60 | 26 | 0.054 | - | - | Area PFm (IPL) | 10 | NHA | Left | AG | 47 |
| 1 | 13608 | -48 | 14 | -26 | 0.026 | - | - | - | - | NA | Left | TP | 59 |
| 2 | 4192 | -50 | 28 | 4 | 0.055 | - | - | Area OP9 | 71 | HA | Left | IFG PTr | 44 |
| 2 | 4192 | -54 | 24 | 12 | 0.051 | - | - | Area 45 | 50 | HA | Left | IFG PTr | 53 |
| 2 | 4192 | -46 | 20 | 24 | 0.030 | - | - | Area 45 | 8 | NHA | Left | IFG POp | 32 |
| 3 | 2144 | -40 | -46 | -20 | 0.046 | - | - | Area FG4 | 72 | HA | Left | TOFC | 33 |
| 3 | 2144 | -40 | -56 | -20 | 0.045 | - | - | Area FG4 | 64 | HA | Left | TOFC | 72 |
| 3 | 2144 | -32 | -38 | -16 | 0.032 | - | - | Area FG3 | 21 | NHA | Left | TFCp | 60 |
| 1 | 14704 | -48 | 24 | 18 | 0.113 | - | - | Area 45 | 14 | NHA | Left | IFG PTr | 25 |
| 1 | 14704 | -46 | 30 | -8 | 0.067 | - | - | Area OP9 | 20 | NHA | Left | FOC | 44 |
| 1 | 14704 | -32 | 24 | -6 | 0.037 | - | - | Area Id7 | 2 | NHA | Left | FOC | 55 |
| 2 | 12072 | -56 | -38 | 0 | 0.078 | - | - | Area TE 5 | 2 | NHA | Left | MTGp | 43 |
| 2 | 12072 | -48 | -54 | -14 | 0.075 | - | - | Area FG4 | 58 | HA | Left | ITGt | 57 |
| 2 | 12072 | -36 | -32 | -20 | 0.066 | - | - | CA1 (Hippocampus) | 15 | NHA | Left | TFCp | 60 |
| 3 | 6016 | -46 | -68 | 26 | 0.068 | - | - | Area PGp (IPL) | 45 | HA | Left | LOCs | 66 |
| 3 | 6016 | -32 | -64 | 40 | 0.064 | - | - | Area hIP6 (IPS) | 39 | HA | Left | LOCs | 54 |
| 3 | 6016 | -34 | -52 | 44 | 0.039 | - | - | Area hIP3 (IPS) | 43 | HA | Left | SPL | 31 |
| 3 | 6016 | -40 | -78 | 36 | 0.033 | - | - | Area PGp (IPL) | 55 | HA | Left | LOCs | 77 |
| 4 | 5896 | -4 | 20 | 48 | 0.106 | - | - | Area 6mr / preSMA | 6 | NHA | Left | PAC | 49 |
| 5 | 2816 | 36 | 24 | -6 | 0.066 | - | - | Area Id7 | 17 | NHA | Right | FOC | 65 |
| 5 | 2816 | 34 | 38 | -10 | 0.044 | - | - | Area Fo7 | 59 | HA | Right | FP | 61 |
| 6 | 1728 | -24 | -14 | -18 | 0.049 | - | - | CA1 (Hippocampus) | 58 | HA | Left | Left HC | 91 |
| 1 | 8816 | -54 | 22 | 16 | 0.070 | - | - | Area 45 | 49 | HA | Left | IFG PTr | 42 |
| 1 | 8816 | -48 | 16 | 22 | 0.065 | - | - | Area 44 | 28 | NHA | Left | IFG POp | 53 |
| 1 | 8816 | -40 | 32 | -12 | 0.034 | - | - | Area Fo7 | 23 | NHA | Left | FOC | 45 |
| 2 | 8152 | -56 | -6 | -14 | 0.085 | - | - | Area TE 5 | 69 | HA | Left | MTGa | 38 |
| 2 | 8152 | -48 | 14 | -26 | 0.070 | - | - | - | - | NA | Left | TP | 59 |
| 3 | 7352 | -52 | -40 | 4 | 0.089 | - | - | - | - | NA | Left | - | - |
| 3 | 7352 | -46 | -58 | 20 | 0.049 | - | - | Area PGa (IPL) | 1 | NHA | Left | AG | 49 |
| 3 | 7352 | -50 | -52 | 16 | 0.038 | - | - | - | - | NA | Left | AG | 33 |
| 3 | 7352 | -54 | -58 | 28 | 0.034 | - | - | Area PGa (IPL) | 57 | HA | Left | AG | 58 |
| 3 | 7352 | -64 | -24 | 0 | 0.030 | - | - | Area TE 4 | 39 | HA | Left | STGp | 41 |
| 4 | 5520 | 52 | 2 | -20 | 0.061 | - | - | Area TE 5 | 83 | HA | Right | STGa | 44 |
| 4 | 5520 | 56 | -6 | -12 | 0.055 | - | - | Area TE 4 | 55 | HA | Right | STGa | 33 |
| 4 | 5520 | 50 | 12 | -26 | 0.049 | - | - | Area TE 5 | 10 | NHA | Right | TP | 84 |
| 4 | 5520 | 50 | -18 | -10 | 0.047 | - | - | Area TE 5 | 58 | HA | Right | MTGp | 46 |
| 5 | 1536 | -4 | 20 | 48 | 0.040 | - | - | Area 6mr / preSMA | 6 | NHA | Left | PAC | 49 |
| 5 | 1536 | -6 | 4 | 56 | 0.033 | - | - | Area 6mr / preSMA | 73 | HA | Left | JLC | 65 |
| 5 | 1536 | 6 | 16 | 44 | 0.033 | - | - | Area 6mr / preSMA | 0 | NHA | Right | PAC | 66 |
| 1 | 3560 | -40 | -40 | -14 | 0.036 | - | - | Area FG3 | 37 | HA | Left | - | - |
| 1 | 3560 | -50 | -44 | -18 | 0.028 | - | - | Area FG4 | 51 | HA | Left | ITGp | 28 |
| 1 | 3560 | -54 | -42 | -16 | 0.027 | - | - | - | - | NA | Left | - | - |
| 1 | 3560 | -30 | -40 | -18 | 0.019 | - | - | Area FG3 | 52 | HA | Left | TFCp | 61 |
| 2 | 1656 | -58 | -32 | 2 | 0.035 | - | - | Area TE 4 | 18 | NHA | Left | STGp | 30 |

## References

Abutalebi, J., & Green, D. (2007). Bilingual language production: The neurocognition of language representation and control. Journal of Neurolinguistics, 20(3), 242–275. 10.1016/j.jneuroling.2006.10.003

Aichhorn, M., Perner, J., Kronbichler, M., Staffen, W., & Ladurner, G. (2006). Do visual perspective tasks need theory of mind? NeuroImage, 30(3), 1059–1068. 10.1016/j.neuroimage.2005.10.026

Aichhorn, M., Perner, J., Weiss, B., Kronbichler, M., Staffen, W., & Ladurner, G. (2009). Temporo-parietal Junction Activity in Theory-of-Mind Tasks: Falseness, Beliefs, or Attention. Journal of Cognitive Neuroscience, 21(6), 1179–1192. 10.1162/jocn.2009.21082

Andrews-Hanna, J. R., Reidler, J. S., Sepulcre, J., Poulin, R., & Buckner, R. L. (2010). Functional-Anatomic Fractionation of the Brain’s Default Network. Neuron, 65(4), 550–562. 10.1016/j.neuron.2010.02.005

Assem, M., Glasser, M. F., Van Essen, D. C., & Duncan, J. (2020). A Domain-General Cognitive Core Defined in Multimodally Parcellated Human Cortex. Cerebral Cortex, 30(8), 4361–4380. 10.1093/cercor/bhaa023

Badre, D., Poldrack, R. A., Paré-Blagoev, E. J., Insler, R. Z., & Wagner, A. D. (2005). Dissociable Controlled Retrieval and Generalized Selection Mechanisms in Ventrolateral Prefrontal Cortex. Neuron, 47(6), 907–918. 10.1016/j.neuron.2005.07.023

Balgova, E., Diveica, V., Jackson, R. L., & Binney, R. J. (2024). Overlapping Neural Correlates Underpin Theory of Mind and Semantic Cognition: Evidence from a Meta-Analysis of 344 Functional Neuroimaging Studies (p. 2023.08.16.553506). bioRxiv. 10.1101/2023.08.16.553506

Balgova, E., Diveica, V., Walbrin, J., & Binney, R. J. (2022). The role of the ventrolateral anterior temporal lobes in social cognition. Human Brain Mapping, 43(15), 4589–4608. 10.1002/hbm.25976

Baron-Cohen, S., Leslie, A. M., & Frith, U. (1985). Does the autistic child have a “theory of mind” ? Cognition, 21(1), 37–46. 10.1016/0010-0277(85)90022-8

Binder, J. R., Desai, R. H., Graves, W. W., & Conant, L. L. (2009). Where Is the Semantic System? A Critical Review and Meta-Analysis of 120 Functional Neuroimaging Studies. Cerebral Cortex, 19(12), 2767–2796. 10.1093/cercor/bhp055

Binney, R. J., & Lambon Ralph, M. A. (2015). Using a combination of fMRI and anterior temporal lobe rTMS to measure intrinsic and induced activation changes across the semantic cognition network. Neuropsychologia, 76, 170–181. 10.1016/j.neuropsychologia.2014.11.009

Binney, R. J., & Ramsey, R. (2020). Social Semantics: The role of conceptual knowledge and cognitive control in a neurobiological model of the social brain. Neuroscience & Biobehavioral Reviews, 112, 28–38. 10.1016/j.neubiorev.2020.01.030

Blank, I. A., & Fedorenko, E. (2017). Domain-General Brain Regions Do Not Track Linguistic Input as Closely as Language-Selective Regions. Journal of Neuroscience, 37(41), 9999–10011. 10.1523/JNEUROSCI.3642-16.2017

Branzi, F. M., Humphreys, G. F., Hoffman, P., & Lambon Ralph, M. A. (2020). Revealing the neural networks that extract conceptual gestalts from continuously evolving or changing semantic contexts. NeuroImage, 220, 116802. 10.1016/j.neuroimage.2020.116802

Branzi, F. M., & Lambon Ralph, M. A. (2023). Semantic-specific and domain-general mechanisms for integration and update of contextual information. Human Brain Mapping, 44(17), 5547– 5566. 10.1002/hbm.26454

Branzi, F. M., Martin, C. D., Carreiras, M., & Paz-Alonso, P. M. (2020). Functional connectivity reveals dissociable ventrolateral prefrontal mechanisms for the control of multilingual word retrieval. Human Brain Mapping, 41(1), 80–94. 10.1002/hbm.24788

Branzi, F. M., Martin, C. D., & Paz-Alonso, P. M. (2022). Task-relevant representations and cognitive control demands modulate functional connectivity from ventral occipito-temporal cortex during object recognition tasks. Cerebral Cortex, 32(14), 3068–3080. 10.1093/cercor/bhab401

Branzi, F. M., Pobric, G., Jung, J., & Lambon Ralph, M. A. (2021). The Left Angular Gyrus Is Causally Involved in Context-dependent Integration and Associative Encoding during Narrative Reading. Journal of Cognitive Neuroscience, 33(6), 1082–1095. 10.1162/jocn_a_01698

Buckner, R. L., Andrews-Hanna, J. R., & Schacter, D. L. (2008). The Brain’s Default Network. Annals of the New York Academy of Sciences, 1124(1), 1–38. 10.1196/annals.1440.011

Button, K. S., Ioannidis, J. P. A., Mokrysz, C., Nosek, B. A., Flint, J., Robinson, E. S. J., & Munafò, M. R. (2013). Power failure: Why small sample size undermines the reliability of neuroscience. Nature Reviews Neuroscience, 14(5), 365–376. 10.1038/nrn3475

Crinion, J., & Price, C. J. (2005). Right anterior superior temporal activation predicts auditory sentence comprehension following aphasic stroke. Brain, 128(Pt 12), 2858–2871. 10.1093/brain/awh659

Diachek, E., Blank, I., Siegelman, M., Affourtit, J., & Fedorenko, E. (2020). The Domain-General Multiple Demand (MD) Network Does Not Support Core Aspects of Language Comprehension: A Large-Scale fMRI Investigation. Journal of Neuroscience, 40(23), 4536– 4550. 10.1523/JNEUROSCI.2036-19.2020

Diveica, V., Koldewyn, K., & Binney, R. J. (2021). Establishing a role of the semantic control network in social cognitive processing: A meta-analysis of functional neuroimaging studies. Neuroimage, 245, 118702. 10.1016/j.neuroimage.2021.118702

Duncan, J. (2010). The multiple-demand (MD) system of the primate brain: Mental programs for intelligent behaviour. Trends in Cognitive Sciences, 14(4), 172–179. 10.1016/j.tics.2010.01.004

Eickhoff, S. B., Bzdok, D., Laird, A. R., Kurth, F., & Fox, P. T. (2012). Activation likelihood estimation meta-analysis revisited. NeuroImage, 59(3), 2349–2361. 10.1016/j.neuroimage.2011.09.017

Eickhoff, S. B., Bzdok, D., Laird, A. R., Roski, C., Caspers, S., Zilles, K., & Fox, P. T. (2011). Co-activation patterns distinguish cortical modules, their connectivity and functional differentiation. NeuroImage, 57(3), 938–949. 10.1016/j.neuroimage.2011.05.021

Eickhoff, S. B., Heim, S., Zilles, K., & Amunts, K. (2006). Testing anatomically specified hypotheses in functional imaging using cytoarchitectonic maps. NeuroImage, 32(2), 570–582. 10.1016/j.neuroimage.2006.04.204

Eickhoff, S. B., Laird, A. R., Grefkes, C., Wang, L. E., Zilles, K., & Fox, P. T. (2009). Coordinate-based activation likelihood estimation meta-analysis of neuroimaging data: A random-effects approach based on empirical estimates of spatial uncertainty. Human Brain Mapping, 30(9), 2907–2926. 10.1002/hbm.20718

Eickhoff, S. B., Paus, T., Caspers, S., Grosbras, M.-H., Evans, A. C., Zilles, K., & Amunts, K. (2007). Assignment of functional activations to probabilistic cytoarchitectonic areas revisited. NeuroImage, 36(3), 511–521. 10.1016/j.neuroimage.2007.03.060

Eickhoff, S. B., Stephan, K. E., Mohlberg, H., Grefkes, C., Fink, G. R., Amunts, K., & Zilles, K. (2005). A new SPM toolbox for combining probabilistic cytoarchitectonic maps and functional imaging data. NeuroImage, 25(4), 1325–1335. 10.1016/j.neuroimage.2004.12.034

Fedorenko, E., Duncan, J., & Kanwisher, N. (2013). Broad domain generality in focal regions of frontal and parietal cortex. Proceedings of the National Academy of Sciences, 110(41), 16616– 16621. 10.1073/pnas.1315235110

Fedorenko, E., Ivanova, A. A., & Regev, T. I. (2024). The language network as a natural kind within the broader landscape of the human brain. Nature Reviews Neuroscience, 25(5), 289–312. 10.1038/s41583-024-00802-4

Fedorenko, E., & Shain, C. (2021). Similarity of Computations Across Domains Does Not Imply Shared Implementation: The Case of Language Comprehension. Current Directions in Psychological Science, 30(6), 526–534. 10.1177/09637214211046955

Fitch, W. T., & Martins, M. D. (2014). Hierarchical processing in music, language, and action: Lashley revisited. Annals of the New York Academy of Sciences, 1316(1), 87–104. 10.1111/nyas.12406

Friederici, A. D., & Gierhan, S. M. (2013). The language network. Current Opinion in Neurobiology, 23(2), 250–254. 10.1016/j.conb.2012.10.002

Gainotti, G. (2015). Is the difference between right and left ATLs due to the distinction between general and social cognition or between verbal and non-verbal representations? Neuroscience & Biobehavioral Reviews, 51, 296–312. 10.1016/j.neubiorev.2015.02.004

Geranmayeh, F., Wise, R. J. S., Mehta, A., & Leech, R. (2014). Overlapping Networks Engaged during Spoken Language Production and Its Cognitive Control. Journal of Neuroscience, 34(26), 8728–8740. 10.1523/JNEUROSCI.0428-14.2014

Graessner, A., Zaccarella, E., & Hartwigsen, G. (2021). Differential contributions of left-hemispheric language regions to basic semantic composition. Brain Structure and Function, 226(2), 501–518. 10.1007/s00429-020-02196-2

Green, D. W., & Abutalebi, J. (2013). Language control in bilinguals: The adaptive control hypothesis. Journal of Cognitive Psychology, 25(5), 515–530. 10.1080/20445911.2013.796377

Grosbras, M.-H., Beaton, S., & Eickhoff, S. B. (2012). Brain regions involved in human movement perception: A quantitative voxel-based meta-analysis. Human Brain Mapping, 33(2), 431–454. 10.1002/hbm.21222

Hartwigsen, G., Saur, D., Price, C. J., Ulmer, S., Baumgaertner, A., & Siebner, H. R. (2013). Perturbation of the left inferior frontal gyrus triggers adaptive plasticity in the right homologous area during speech production. Proceedings of the National Academy of Sciences, 110(41), 16402–16407. 10.1073/pnas.1310190110

Hasson, U., Egidi, G., Marelli, M., & Willems, R. M. (2018). Grounding the neurobiology of language in first principles: The necessity of non-language-centric explanations for language comprehension. Cognition, 180, 135–157. 10.1016/j.cognition.2018.06.018

Hodgson, V. J., Lambon Ralph, M. A., & Jackson, R. L. (2021). Multiple dimensions underlying the functional organization of the language network. NeuroImage, 241, 118444. 10.1016/j.neuroimage.2021.118444

Hodgson, V. J., Lambon Ralph, M. A., & Jackson, R. L. (2022). The cross-domain functional organization of posterior lateral temporal cortex: Insights from ALE meta-analyses of 7 cognitive domains spanning 12,000 participants. Cerebral Cortex, bhac394. 10.1093/cercor/bhac394

Hu, J., Small, H., Kean, H., Takahashi, A., Zekelman, L., Kleinman, D., Ryan, E., Nieto-Castañón, A., Ferreira, V., & Fedorenko, E. (2023). Precision fMRI reveals that the language-selective network supports both phrase-structure building and lexical access during language production. Cerebral Cortex, 33(8), 4384–4404. 10.1093/cercor/bhac350

Humphreys, G. F., Halai, A. D., Branzi, F. M., & Lambon Ralph, M. A. (2024). The left posterior angular gyrus is engaged by autobiographical recall not object-semantics, or event-semantics: Evidence from contrastive propositional speech production. Imaging Neuroscience, 2, 1–19. 10.1162/imag_a_00116

Humphreys, G. F., & Lambon Ralph, M. A. (2015). Fusion and Fission of Cognitive Functions in the Human Parietal Cortex. Cerebral Cortex, 25(10), 3547–3560. 10.1093/cercor/bhu198

Humphreys, G. F., & Lambon Ralph, M. A. (2017). Mapping Domain-Selective and Counterpointed Domain-General Higher Cognitive Functions in the Lateral Parietal Cortex: Evidence from fMRI Comparisons of Difficulty-Varying Semantic Versus Visuo-Spatial Tasks, and Functional Connectivity Analyses. Cerebral Cortex, 27(8), 4199–4212. 10.1093/cercor/bhx107

Jackson, R. L. (2021). The neural correlates of semantic control revisited. NeuroImage, 224, 117444. 10.1016/j.neuroimage.2020.117444

Jung, J., & Lambon Ralph, M. A. (2016). Mapping the Dynamic Network Interactions Underpinning Cognition: A cTBS-fMRI Study of the Flexible Adaptive Neural System for Semantics. Cerebral Cortex, 26(8), 3580–3590. 10.1093/cercor/bhw149

Jung, J. Y., Rice, G. E., & Lambon Ralph, M. A. (2021). The neural bases of resilient semantic system: Evidence of variable neuro-displacement in cognitive systems. Brain Structure and Function, 226(5), 1585–1599. 10.1007/s00429-021-02272-1

Koechlin, E., & Jubault, T. (2006). Broca’s Area and the Hierarchical Organization of Human Behavior. Neuron, 50(6), 963–974. 10.1016/j.neuron.2006.05.017

Konu, D., Turnbull, A., Karapanagiotidis, T., Wang, H.-T., Brown, L. R., Jefferies, E., & Smallwood, J. (2020). A role for the ventromedial prefrontal cortex in self-generated episodic social cognition. NeuroImage, 218, 116977. 10.1016/j.neuroimage.2020.116977

Krieger-Redwood, K., Teige, C., Davey, J., Hymers, M., & Jefferies, E. (2015). Conceptual control across modalities: Graded specialisation for pictures and words in inferior frontal and posterior temporal cortex. Neuropsychologia, 76, 92–107. 10.1016/j.neuropsychologia.2015.02.030

Kuperberg, G. R., Holcomb, P. J., Sitnikova, T., Greve, D., Dale, A. M., & Caplan, D. (2003). Distinct Patterns of Neural Modulation during the Processing of Conceptual and Syntactic Anomalies. Journal of Cognitive Neuroscience, 15(2), 272–293. 10.1162/089892903321208204

Laird, A. R., Robinson, J. L., McMillan, K. M., Tordesillas-Gutiérrez, D., Moran, S. T., Gonzales, S. M., Ray, K. L., Franklin, C., Glahn, D. C., Fox, P. T., & Lancaster, J. L. (2010). Comparison of the disparity between Talairach and MNI coordinates in functional neuroimaging data: Validation of the Lancaster transform. NeuroImage, 51(2), 677–683. 10.1016/j.neuroimage.2010.02.048

Lambon Ralph, M. A. L., Jefferies, E., Patterson, K., & Rogers, T. T. (2017). The neural and computational bases of semantic cognition. Nature Reviews Neuroscience, 18(1), Article 1. 10.1038/nrn.2016.150

Lancaster, J. L., Tordesillas-Gutiérrez, D., Martinez, M., Salinas, F., Evans, A., Zilles, K., Mazziotta, J. C., & Fox, P. T. (2007). Bias between MNI and Talairach coordinates analyzed using the ICBM-152 brain template. Human Brain Mapping, 28(11), 1194–1205. 10.1002/hbm.20345

Lemire-Rodger, S., Lam, J., Viviano, J. D., Stevens, W. D., Spreng, R. N., & Turner, G. R. (2019). Inhibit, switch, and update: A within-subject fMRI investigation of executive control. Neuropsychologia, 132, 107134. 10.1016/j.neuropsychologia.2019.107134

Lindell, A. K. (2006). In Your Right Mind: Right Hemisphere Contributions to Language Processing and Production. Neuropsychology Review, 16(3), 131–148. 10.1007/s11065-006-9011-9

Marinkovic, K., Dhond, R. P., Dale, A. M., Glessner, M., Carr, V., & Halgren, E. (2003). Spatiotemporal Dynamics of Modality-Specific and Supramodal Word Processing. Neuron, 38(3), 487–497. 10.1016/S0896-6273(03)00197-1

Mason, R. A., & Just, M. A. (2007). Lexical ambiguity in sentence comprehension. Brain Research, 1146, 115–127. 10.1016/j.brainres.2007.02.076

Mcmillan, C. T., Coleman, D., Clark, R., Liang, T.-W., Gross, R. G., & Grossman, M. (2013). Converging Evidence for the Processing Costs Associated with Ambiguous Quantifier Comprehension. Frontiers in Psychology, 4. 10.3389/fpsyg.2013.00153

Mollica, F., Siegelman, M., Diachek, E., Piantadosi, S. T., Mineroff, Z., Futrell, R., Kean, H., Qian, P., & Fedorenko, E. (2020). Composition is the Core Driver of the Language-selective Network. Neurobiology of Language, 1(1), 104–134. 10.1162/nol_a_00005

Müller, V. I., Cieslik, E. C., Laird, A. R., Fox, P. T., Radua, J., Mataix-Cols, D., Tench, C. R., Yarkoni, T., Nichols, T. E., Turkeltaub, P. E., Wager, T. D., & Eickhoff, S. B. (2018). Ten simple rules for neuroimaging meta-analysis. Neuroscience & Biobehavioral Reviews, 84, 151–161. 10.1016/j.neubiorev.2017.11.012

Müller, V. I., Höhner, Y., & Eickhoff, S. B. (2018). Influence of task instructions and stimuli on the neural network of face processing: An ALE meta-analysis. Cortex, 103, 240–255. 10.1016/j.cortex.2018.03.011

Nardo, D., Holland, R., Leff, A. P., Price, C. J., & Crinion, J. T. (2017). Less is more: Neural mechanisms underlying anomia treatment in chronic aphasic patients. Brain, 140(11), 3039–3054. 10.1093/brain/awx234

Nichols, T., Brett, M., Andersson, J., Wager, T., & Poline, J.-B. (2005). Valid conjunction inference with the minimum statistic. NeuroImage, 25(3), 653–660. 10.1016/j.neuroimage.2004.12.005

Nieuwland, M. S. (2012). Establishing propositional truth-value in counterfactual and real-world contexts during sentence comprehension: Differential sensitivity of the left and right inferior frontal gyri. NeuroImage, 59(4), 3433–3440. 10.1016/j.neuroimage.2011.11.018

Novais-Santos, S., Gee, J., Shah, M., Troiani, V., Work, M., & Grossman, M. (2007). Resolving sentence ambiguity with planning and working memory resources: Evidence from fMRI. NeuroImage, 37(1), 361–378. 10.1016/j.neuroimage.2007.03.077

Oishi, K., Faria, A. V., Hsu, J., Tippett, D., Mori, S., & Hillis, A. E. (2015). Critical role of the right uncinate fasciculus in emotional empathy. Annals of Neurology, 77(1), 68–74. 10.1002/ana.24300

Olson, I. R., McCoy, D., Klobusicky, E., & Ross, L. A. (2013). Social cognition and the anterior temporal lobes: A review and theoretical framework. Social Cognitive and Affective Neuroscience, 8(2), 123–133. 10.1093/scan/nss119

Olson, I. R., Plotzker, A., & Ezzyat, Y. (2007). The Enigmatic temporal pole: A review of findings on social and emotional processing. Brain, 130(7), 1718–1731. 10.1093/brain/awm052

Papinutto, N., Galantucci, S., Mandelli, M. L., Gesierich, B., Jovicich, J., Caverzasi, E., Henry, R. G., Seeley, W. W., Miller, B. L., Shapiro, K. A., & Gorno-Tempini, M. L. (2016). Structural connectivity of the human anterior temporal lobe: A diffusion magnetic resonance imaging study. Human Brain Mapping, 37(6), 2210–2222. 10.1002/hbm.23167

Pexman, P. M., Diveica, V., & Binney, R., J. (2023). Social semantics: The organization and grounding of abstract concepts | Philosophical Transactions of the Royal Society B: Biological Sciences. https://royalsocietypublishing.org/doi/full/10.1098/rstb.2021.0363

Pobric, G., Lambon Ralph, M. A., & Zahn, R. (2016). Hemispheric Specialization within the Superior Anterior Temporal Cortex for Social and Nonsocial Concepts. Journal of Cognitive Neuroscience, 28(3), 351–360. 10.1162/jocn_a_00902

Price, C. J. (2012). A review and synthesis of the first 20 years of PET and fMRI studies of heard speech, spoken language and reading. NeuroImage, 62(2), 816–847. 10.1016/j.neuroimage.2012.04.062

Quillen, I. A., Yen, M., & Wilson, S. M. (2021). Distinct Neural Correlates of Linguistic and Non-Linguistic Demand. Neurobiology of Language, 2(2), 202–225. 10.1162/nol_a_00031

Raichle, M. E., MacLeod, A. M., Snyder, A. Z., Powers, W. J., Gusnard, D. A., & Shulman, G. L. (2001). A default mode of brain function. Proceedings of the National Academy of Sciences, 98(2), 676–682. 10.1073/pnas.98.2.676

Ranganath, C., & Ritchey, M. (2012). Two cortical systems for memory-guided behaviour. Nature Reviews Neuroscience, 13(10), 713–726. 10.1038/nrn3338

Rankin, K. P., Gorno-Tempini, M. L., Allison, S. C., Stanley, C. M., Glenn, S., Weiner, M. W., & Miller, B. L. (2006). Structural anatomy of empathy in neurodegenerative disease. Brain, 129(11), 2945–2956. 10.1093/brain/awl254

Rice, G. E., Caswell, H., Moore, P., Lambon Ralph, M. A., & Hoffman, P. (2018). Revealing the Dynamic Modulations That Underpin a Resilient Neural Network for Semantic Cognition: An fMRI Investigation in Patients With Anterior Temporal Lobe Resection. Cerebral Cortex, 28(8), 3004–3016. 10.1093/cercor/bhy116

Rice, G. E., Lambon Ralph, M. A., & Hoffman, P. (2015). The Roles of Left Versus Right Anterior Temporal Lobes in Conceptual Knowledge: An ALE Meta-analysis of 97 Functional Neuroimaging Studies. Cerebral Cortex, 25(11), 4374–4391. 10.1093/cercor/bhv024

Rodd, J. M., Davis, M. H., & Johnsrude, I. S. (2005). The Neural Mechanisms of Speech Comprehension: fMRI studies of Semantic Ambiguity. Cerebral Cortex, 15(8), 1261–1269. 10.1093/cercor/bhi009

Rolls, E. T., & Grabenhorst, F. (2008). The orbitofrontal cortex and beyond: From affect to decision-making. Progress in Neurobiology, 86(3), 216–244. 10.1016/j.pneurobio.2008.09.001

Saxe, R., Carey, S., & Kanwisher, N. (2004). Understanding Other Minds: Linking Developmental Psychology and Functional Neuroimaging. Annual Review of Psychology, 55(1), 87–124. 10.1146/annurev.psych.55.090902.142044

Saxe, R., & Wexler, A. (2005). Making sense of another mind: The role of the right temporo-parietal junction. Neuropsychologia, 43(10), 1391–1399. 10.1016/j.neuropsychologia.2005.02.013

Schumacher, R., Halai, A. D., & Lambon Ralph, M. A. (2019). Assessing and mapping language, attention and executive multidimensional deficits in stroke aphasia. Brain, 142(10), 3202– 3216. 10.1093/brain/awz258

Seeley, W. W., Bauer, A. M., Miller, B. L., Gorno-Tempini, M. L., Kramer, J. H., Weiner, M., & Rosen, H. J. (2005). The natural history of temporal variant frontotemporal dementia. Neurology, 64(8), 1384–1390. 10.1212/01.WNL.0000158425.46019.5C

Shain, C., Blank, I. A., van Schijndel, M., Schuler, W., & Fedorenko, E. (2020). fMRI reveals language-specific predictive coding during naturalistic sentence comprehension. Neuropsychologia, 138, 107307. 10.1016/j.neuropsychologia.2019.107307

Silbert, L. J., Honey, C. J., Simony, E., Poeppel, D., & Hasson, U. (2014). Coupled neural systems underlie the production and comprehension of naturalistic narrative speech. Proceedings of the National Academy of Sciences, 111(43), E4687–E4696. 10.1073/pnas.1323812111

Smallwood, J., Bernhardt, B. C., Leech, R., Bzdok, D., Jefferies, E., & Margulies, D. S. (2021). The default mode network in cognition: A topographical perspective. Nature Reviews Neuroscience, 22(8), 503–513. 10.1038/s41583-021-00474-4

Stefaniak, J. D., Alyahya, R. S. W., & Lambon Ralph, M. A. (2021). Language networks in aphasia and health: A 1000 participant activation likelihood estimation meta-analysis. NeuroImage, 233, 117960. 10.1016/j.neuroimage.2021.117960

Stefaniak, J. D., Halai, A. D., & Lambon Ralph, M. A. (2020). The neural and neurocomputational bases of recovery from post-stroke aphasia. Nat Rev Neurol, 16(1), 43–55. 10.1038/s41582-019-0282-1

Turkeltaub, P. E., Eden, G. F., Jones, K. M., & Zeffiro, T. A. (2002). Meta-Analysis of the Functional Neuroanatomy of Single-Word Reading: Method and Validation. NeuroImage, 16(3), 765– 780. 10.1006/nimg.2002.1131

Turkeltaub, P. E., Eickhoff, S. B., Laird, A. R., Fox, M., Wiener, M., & Fox, P. (2012). Minimizing within-experiment and within-group effects in Activation Likelihood Estimation meta-analyses. Human Brain Mapping, 33(1), 1–13. 10.1002/hbm.21186

Turker, S., Kuhnke, P., Eickhoff, S. B., Caspers, S., & Hartwigsen, G. (2023). Cortical, subcortical, and cerebellar contributions to language processing: A meta-analytic review of 403 neuroimaging experiments. Psychological Bulletin. 10.1037/bul0000403

Vatansever, D., Menon, D. K., & Stamatakis, E. A. (2017). Default mode contributions to automated information processing. Proceedings of the National Academy of Sciences, 114(48), 12821– 12826. 10.1073/pnas.1710521114

Visser, M., & Lambon Ralph, M. A. (2011). Differential Contributions of Bilateral Ventral Anterior Temporal Lobe and Left Anterior Superior Temporal Gyrus to Semantic Processes. Journal of Cognitive Neuroscience, 23(10), 3121–3131. 10.1162/jocn_a_00007

Von Der Heide, R. J., Skipper, L. M., Klobusicky, E., & Olson, I. R. (2013). Dissecting the uncinate fasciculus: Disorders, controversies and a hypothesis. Brain, 136(6), 1692–1707. 10.1093/brain/awt094

Whitney, C., Grossman, M., & Kircher, T. T. J. (2009). The Influence of Multiple Primes on Bottom-Up and Top-Down Regulation during Meaning Retrieval: Evidence for 2 Distinct Neural Networks. Cerebral Cortex, 19(11), 2548–2560. 10.1093/cercor/bhp007

Wright, P., Randall, B., Marslen-Wilson, W. D., & Tyler, L. K. (2011). Dissociating Linguistic and Task-related Activity in the Left Inferior Frontal Gyrus. Journal of Cognitive Neuroscience, 23(2), 404–413. 10.1162/jocn.2010.21450

Wu, W., & Hoffman, P. (2024). Functional integration and segregation during semantic cognition: Evidence across age groups. Cortex, 178, 157–173. 10.1016/j.cortex.2024.06.015

Xu, J., Kemeny, S., Park, G., Frattali, C., & Braun, A. (2005). Language in context: Emergent features of word, sentence, and narrative comprehension. NeuroImage, 25(3), 1002–1015. 10.1016/j.neuroimage.2004.12.013

Zahn, R., Moll, J., Krueger, F., Huey, E. D., Garrido, G., & Grafman, J. (2007). Social concepts are represented in the superior anterior temporal cortex. Proceedings of the National Academy of Sciences, 104(15), 6430–6435. 10.1073/pnas.0607061104

Zilles, K., & Amunts, K. (2010). Centenary of Brodmann’s map—Conception and fate. Nature Reviews Neuroscience, 11(2), 139–145. 10.1038/nrn2776

Zilles, K., Schleicher, A., Palomero-Gallagher, N., & Amunts, K. (2002). 21—Quantitative Analysis of Cyto- and Receptor Architecture of the Human Brain. In A. W. Toga & J. C. Mazziotta (Eds.), Brain Mapping: The Methods (Second Edition) (pp. 573–602). Academic Press. 10.1016/B978-012693019-1/50023-X

